# Introducing RELAX (the Reduction of Electroencephalographic Artifacts): A fully automated pre-processing pipeline for cleaning EEG data – Part 2: Application to Event-Related Potentials

**DOI:** 10.1101/2022.03.08.483554

**Authors:** NW Bailey, AT Hill, M Biabani, OW Murphy, NC Rogasch, B McQueen, A Miljevic, PB Fitzgerald

## Abstract

Electroencephalography (EEG) is commonly used to examine neural activity time-locked to the presentation of a stimulus, referred to as an Event-Related Potential (ERP). However, EEG is also influenced by non-neural artifacts, which can confound ERP comparisons. Artifact cleaning can reduce artifacts, but often requires time-consuming manual decisions. Most automated cleaning methods require frequencies <1Hz to be filtered out of the data, so are not recommended for ERPs (which often contain <1Hz frequencies). In our companion article, we introduced RELAX (the Reduction of Electroencephalographic Artifacts), an automated and modular cleaning pipeline that reduces artifacts with Multiple Wiener Filtering (MWF) and/or wavelet enhanced independent component analysis (wICA) applied to artifact components detected with ICLabel (wICA_ICLabel) (Bailey et al., 2022). To evaluate the suitability of RELAX for data cleaning prior to ERP analysis, multiple versions of RELAX were compared to four commonly used EEG cleaning pipelines. Cleaning performance was compared across a range of artifact cleaning metrics and in the amount of variance in ERPs explained by different conditions in a Go-Nogo task. RELAX with MWF and wICA_ICLabel cleaned the data the most effectively and produced amongst the most dependable ERP estimates. RELAX with wICA_ICLabel only or MWF_only may detect experimental effects better for some ERP measures. Importantly, RELAX can high-pass filter data at 0.25Hz, so is applicable to analyses involving ERPs. The pipeline is easy to implement via EEGLAB in MATLAB and is freely available on GitHub. Given its performance, objectivity, and ease of use, we recommend RELAX for EEG data cleaning.

The MATLAB code, the supplementary materials, and a simple instruction manual explaining how to implement the RELAX pipeline can be downloaded from https://github.com/NeilwBailey/RELAX/releases. A condition of use of the pipeline is that the version of the pipeline used is referred to as RELAX_[pipeline], for example “RELAX_MWF_wICA” or “RELAX_wICA_ICLabel”, and that the current paper be cited, as well as the dependencies used. These dependencies are likely to include: EEGLAB (Delorme & Makeig, 2004), fieldtrip (Oostenveld et al., 2011), the MWF toolbox (Somers et al., 2019), fastICA (Hyvarinen, 1999), wICA (Castellanos & Makarov, 2006), ICLabel (Pion-Tonachini et al., 2019), and PREP (Bigdely-Shamlo et al., 2015)

See our companion article for the application of RELAX to the study of oscillatory power (Bailey et al., 2022).

## Introduction

Electroencephalography (EEG) enables investigators to non-invasively measure voltage fluctuations produced by the brain. These voltage fluctuations can be used to obtain information about neural activity related to cognitive processes. To do this, voltage fluctuations following multiple presentations of a stimulus are commonly averaged together, eliminating activity that is not strictly time-locked to the stimulus presentation, which provides an indication of the consistent aspects of brain responses to that stimulus. The outcome of this is process is referred to as an Event-Related Potential (ERP). Unfortunately, in addition to activity produced by the brain, EEG electrodes record the contribution of non-brain related artifacts. Biological artifacts such as blinks, eye movements, and muscle activity generate well-characterised voltage fluctuations, so they are sometimes easy to detect and remove (Fitzgibbon et al., 2016; Kleifges et al., 2017; Muthukumaraswamy, 2013). However, small amplitude examples of these artifacts, and other small amplitude biological artifacts (such as cardiac field artifacts) can be more difficult to distinguish from ongoing EEG activity, and the full extent of their influence is difficult to know (Gerla et al., 2017; Muthukumaraswamy, 2013; Schlögl et al., 2007). Non-biological artifacts are also often present, including voltage drift, 50Hz or 60Hz electrical interference from the alternating current of electrical equipment; and channel noise from poor electrical connections between the electrodes and scalp (Pion-Tonachini et al., 2019). All EEG electrodes record a combination of neural activity and artifacts, and it is not possible to determine with absolute confidence which aspects reflect neural activity in isolation (the “ground truth” of the data). This fact means that conclusions about differences in brain activity between different experimental conditions are vulnerable to being confounded by artifacts (Yuval-Greenberg et al., 2008). To give a concrete example of how this might occur, EEG data could be recorded while an anxious and non-anxious group performed a memory task, with condition 1 containing fear eliciting stimuli, and condition 2 containing neutral stimuli. Results might indicate that anxious participants show larger ERPs in response to fear stimuli. However, if blink artifacts are not removed, this difference might be driven by anxious participants blinking more frequently or with stronger muscle contractions in response to the fear stimuli, particularly if blinks are time-locked and consistent in response to seeing the fear stimuli (which can happen with some experimental designs) (Li & Principe, 2006).

To address the issue of artifacts in EEG data, many EEG cleaning methods have been developed (see Islam et al. (2016) and Ranjan et al. (2021) for reviews, and Barban et al. (2021) and Robbins et al. (2020) for comparisons across multiple cleaning methods). Although these existing EEG pre-processing methods have been demonstrated to meet their aim of reducing artifacts while preserving neural activity, there are still several outstanding issues. We have summarised most of these in detail in our companion article (Bailey et al. 2022). For the purposes of this article, we focus on the following issue: while fully automated pre-processing methods are available (for example, The Harvard Automated Processing Pipeline for Electroencephalography [HAPPE]; Gabard-Durnam et al. (2018)), most existing automated methods apply a 1Hz high-pass filter to the data, because the independent component analysis method (ICA; described later) that most pipelines rely upon to clean the data performs better with data that has been high pass filtered at 1Hz (Winkler et al., 2015). However, filtering out this low frequency data reduces the amplitude of ERPs and distorts their latencies (Rousselet, 2012; Tanner et al., 2015). As such, most automated EEG cleaning pipelines are unsuitable for use in studies involving the analysis of ERPs. Additionally, very few studies that have assessed a new EEG cleaning pipeline have examined the most practically important metric – whether cleaning enhances the detection of ERP experimental effects of interest, and whether the cleaning pipeline produces more reliable ERP estimates (Clayson, Baldwin, et al., 2021). Reliable ERP estimates obtained from clean data may be more important to consider when designing an ERP study than even the sample size of the study, as recent research has suggested the data quality contributes more to study power than sample size (Kappenman & Luck, 2010).

To address these issues, we built the “RELAX” EEG cleaning pipeline (short for “Reduction of Electroencephalographic Artifacts”), in which we combined and adjusted pre-existing approaches to optimise EEG cleaning and ERP outcomes. The pipeline is explained in detail in our companion article (Bailey et al., 2022), but in brief, RELAX: 1) Removes bad electrodes and extreme outlying EEG periods that are unlikely to have recorded meaningful or recoverable brain activity using a combination of algorithms obtained from previous research so that the rejected data matched our expert judgement for all common extreme artifact types; 2) The pipeline then reduces artifacts with Multi-channel Wiener Filters (MWF) (Borowicz, 2018; Somers et al., 2018); and 3) RELAX further reduces any remaining artifacts using wavelet enhanced ICA (wICA) (Castellanos & Makarov, 2006) applied to artifact components identified by the machine learning algorithm ‘ICLabel’ (Pion-Tonachini et al., 2019). This combination approach effectively cleans blinks, muscle activity, horizontal eye movement, voltage drift, and atypical artifacts to produce more dependable ERP estimates that indicate increased signal-to-noise ratios. The pipeline is fully automated but also modular (so it can be adjusted to produce optimal results for a user’s specific experimental design). RELAX is implemented via a graphical user interface in EEGLAB, and is freely available on GitHub (https://github.com/NeilwBailey/RELAX/releases). In this manuscript we report comparisons of both cleaning quality metrics and ERP outcome measures between different versions of the RELAX pipeline and a range of existing pipelines using a large EEG dataset from a typical ERP study. A comprehensive supplementary materials can be found at the GitHub repository, which includes additional exploratory comparisons of certain parameter selections within the RELAX pipeline (Supplementary Materials, section 5, page 72).

## Methods

### RELAX Pipeline

Full details of the RELAX pipeline are reported in our companion article https://github.com/NeilwBailey/RELAX/releases. We tested a number of different versions of RELAX, which were identical to those described in our companion article, except that instead of high-pass filtering at 1Hz, data were high-pass filtered at 0.25Hz which is more appropriate for the analysis of ERPs. An overview of the specific steps included in our recommended RELAX pipeline for ERP analysis is provided in Figure 1.

**Figure 1.**
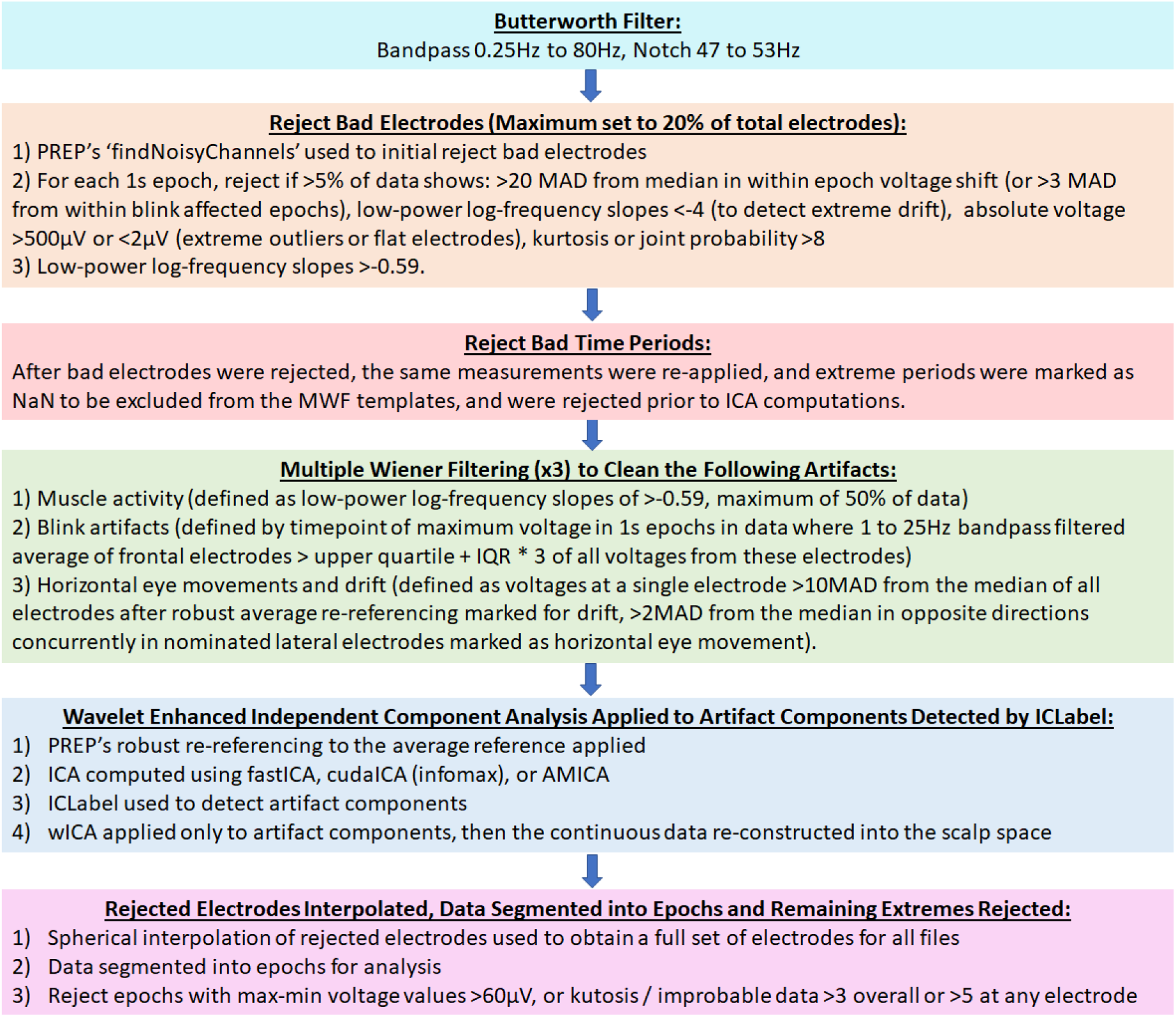
Steps involved in the recommended RELAX pipeline, with the 0.25Hz high-pass filtering option used for ERP analysis. Abbreviations: Hz = hertz; s = seconds; PREP = EEG Preprocessing Pipeline; MAD = median absolute deviation; MWF = multiple Wiener filters; ICA = independent component analysis; IQR = inter-quartile range; AMICA adaptive mixture ICA; cudaICA = ICA performed using the cuda cores of a graphic card; wICA = wavelet enhanced ICA.

### Comparison Pipelines

To comprehensively test the effectiveness of the pipelines for cleaning EEG data in preparation for ERP analysis, we compared seven versions of the RELAX pipeline to four commonly used pipelines (for a total of 11 pipelines included in the comparisons). We present a summary of the steps involved in the pipelines in Tables 1 and 2. All pipelines were implemented on data after the same initial bad electrode and extreme outlying EEG data period rejection (described in full in our companion article). More details for each pipeline are available in the Supplementary Materials (section 2, page 5), and from the citations provided in the following.

**Table 1.** A summary of the steps involved in each variation of the RELAX pipeline. Abbreviations: Hz = hertz; s = seconds; PREP = EEG Preprocessing Pipeline; MAD = median absolute deviation; MWF = multiple Wiener filters; ICA = independent component analysis; IQR = inter-quartile range; AMICA adaptive mixture ICA; cudaICA = ICA performed using the cuda cores of a graphic card; wICA = wavelet enhanced ICA.

|  | MWF_wICA | MWF_wICA_45Hz | MWF_ICA_subtract | MWF_wICA_CCA | wICA_ICLabel |
| --- | --- | --- | --- | --- | --- |
| **Filtering** | 0.25Hz to 80Hz, with 47-53Hz notch filter | 0.25Hz to 80Hz, with 47-53Hz notch filter | 0.25Hz to 80Hz, with 47-53Hz notch filter | 0.25Hz to 80Hz, with 47-53Hz notch filter | 0.25Hz to 80Hz, with 47-53Hz notch filter |
| **Bad Channel Rejection** | PREP's 'findNoisyChannels', then for each 1s epoch, reject if >5% of data shows extreme values (defined in Figure 1) or log-power log-frequency slopes >-0.59 | PREP's 'findNoisyChannels', then for each 1s epoch, reject if >5% of data shows extreme values (defined in Figure 1) or log-power log-frequency slopes >-0.59 | PREP's 'findNoisyChannels', then for each 1s epoch, reject if >5% of data shows extreme values (defined in Figure 1) or log-power log-frequency slopes >-0.59 | PREP's 'findNoisyChannels', then for each 1s epoch, reject if >5% of data shows extreme values (defined in Figure 1) or log-power log-frequency slopes >-0.59 | PREP's 'findNoisyChannels', then for each 1s epoch, reject if >5% of data shows extreme values (defined in Figure 1) or log-power log-frequency slopes >-0.59 |
| **Initial Outlying Data Period Rejection** | After bad channels are rejected, reject remaining 1s epochs that exceed the same thresholds as per the bad channel rejection step | After bad channels are rejected, reject remaining 1s epochs that exceed the same thresholds as per the bad channel rejection step | After bad channels are rejected, reject remaining 1s epochs that exceed the same thresholds as per the bad channel rejection step | After bad channels are rejected, reject remaining 1s epochs that exceed the same thresholds as per the bad channel rejection step | After bad channels are rejected, reject remaining 1s epochs that exceed the same thresholds as per the bad channel rejection step |
| **Initial Artifact Reduction** | 3 sequential MWF, cleaning muscle activity first, then eye blinks, then horizontal eye movement and voltage drift | 3 sequential MWF, cleaning muscle activity first, then eye blinks, then horizontal eye movement and voltage drift | 3 sequential MWF, cleaning muscle activity first, then eye blinks, then horizontal eye movement and voltage drift | 3 sequential MWF, cleaning muscle activity first, then eye blinks, then horizontal eye movement and voltage drift | None |
| **Second Artifact Reduction** | Artifactual ICA components reduced using wICA. ICA computed with fastICA, AMICA, or cudaICA (results only from cudaICA are reported in the main manuscript). Artifacts identified by ICLabel. | 45Hz low pass filtering applied. Artifactual ICA components reduced using wICA. ICA computed with cudaICA. Artifacts identified by ICLabel. | ICA computed using cudaICA. Artifacts identified by ICLabel, and rejected. | ICA computed using cudaICA. Artifactual ICA components except for muscle activity reduced using wICA (identified by ICLabel), then extended CCA used to clean muscle activity. | ICA computed using cudaICA. Artifactual ICA components identified by ICLabel. Artifacts reduced using wICA. |

**Table 2.** A summary of the steps involved in each of the comparison cleaning pipelines. Abbreviations: Hz = hertz; s = seconds; PREP = EEG Preprocessing Pipeline; MAD = median absolute deviation; MWF = multiple Wiener filters; ICA = independent component analysis; IQR = inter-quartile range; AMICA adaptive mixture ICA; cudaICA = ICA performed using the cuda cores of a graphic card; wICA = wavelet enhanced ICA.

|  | ICA_Subtract | MWF_only | MWF_CCA | wICA_all |
| --- | --- | --- | --- | --- |
| **Filtering** | 0.25Hz to 80Hz, with 47-53Hz notch filter | 0.25Hz to 80Hz, with 47-53Hz notch filter | 0.25Hz to 80Hz, with 47-53Hz notch filter | 0.25Hz to 80Hz, with 47-53Hz notch filter |
| **Bad Channel Rejection** | PREP's 'findNoisyChannels', then for each 1s epoch, reject if >5% of data shows extreme values (defined in Figure 1) or log-power log-frequency slopes >-0.59 | PREP's 'findNoisyChannels', then for each 1s epoch, reject if >5% of data shows extreme values (defined in Figure 1) or log-power log-frequency slopes >-0.59 | PREP's 'findNoisyChannels', then for each 1s epoch, reject if >5% of data shows extreme values (defined in Figure 1) or log-power log-frequency slopes >-0.59 | PREP's 'findNoisyChannels', then for each 1s epoch, reject if >5% of data shows extreme values (defined in Figure 1) or log-power log-frequency slopes >-0.59 |
| **Initial Outlying Data Period Rejection** | After bad channels are rejected, reject remaining 1s epochs that exceed the same thresholds as per the bad channel rejection step | After bad channels are rejected, reject remaining 1s epochs that exceed the same thresholds as per the bad channel rejection step | After bad channels are rejected, reject remaining 1s epochs that exceed the same thresholds as per the bad channel rejection step | After bad channels are rejected, reject remaining 1s epochs that exceed the same thresholds as per the bad channel rejection step |
| **Initial Artifact Reduction** | None | 3 sequential MWF, cleaning muscle activity first, then eye blinks, then horizontal eye movement and voltage drift | 3 sequential MWF, cleaning muscle activity first, then eye blinks, then horizontal eye movement and voltage drift | None |
| **Second Artifact Reduction** | ICA computed using cudaICA. Artifacts identified by ICLabel, and rejected. | None | Extended canonical correlation analysis to clean muscle activity (Janani et al., 2018). | ICA computed using cudaICA. All ICA components reduced using wICA |

The different versions of RELAX we tested varied specific parameters to determine the optimal pipeline. We tested three different ICA algorithms: 1) extended-infomax via cudaICA (MWF_wICA_infomax) (Raimondo et al., 2012); 2) fastICA (MWF_wICA_fastICA) (Hyvarinen, 1999); or 3) AMICA (Palmer et al., 2012) (MWF_wICA_AMICA). cudaICA was implemented as the ICA method for all other pipelines, and the different RELAX versions only minimally differed depending on ICA method, so in our main manuscript we only report the results of MWF_wICA_infomax (referred to as MWF_wICA hereafter). We also tested: 4) subtracting artifactual ICA components after the MWF cleaning step instead of using wICA (MWF_ICA_subtract); 5) low-pass filtering EEG data at 45Hz before ICA decomposition (MWF_wICA_45Hz), which has been suggested to improve ICA decomposition (Zakeri, 2017). For the sake of brevity, we only report results involving MWF_wICA_45Hz in the Supplementary Materials (section 4, page 11 onwards), as the 45Hz filtering did not improve performance. We also tested: 6) MWF_wICA_CCA, which excluded muscle artifact cleaning from the wICA step in MWF_wICA, and instead cleaning it with the extended canonical correlation analysis (CCA) (Janani et al., 2018); and finally, 7) not implementing any MWF cleaning, and only using wICA applied to artifactual components identified with ICLabel (Pion-Tonachini et al., 2019). This pipeline is referred to as wICA_ICLabel. It is similar to the methods used by Issa and Juhasz (2019) and Mammone et al. (2011), but we used ICLabel to identify and reduce all artifacts (instead of only identifying eye movements) (Issa & Juhasz, 2019), or identifying artifacts using entropy and kurtosis measures applied to the components (Mammone et al., 2011). To help the reader assess the potential of RELAX for application to their data while reading the results, we note here that our recommendation is to use either “**<u>MWF_wICA</u>**” or “**<u>wICA_ICLabel</u>**”.

In addition to the different versions of RELAX, we also tested four pipelines that have been presented by previous research: 8) MWF_only, which was identical to the initial MWF steps in our RELAX pipeline but did not apply the wICA cleaning (Somers et al. 2018); 9) MWF_CCA, which was identical to the initial MWF steps in our RELAX pipeline but used the extended CCA to reduce any residual muscle artifacts instead of applying wICA (Janani et al., 2018); 10) ICA_subtract, which is probably the most frequently implemented cleaning method.

ICA_subtract computed the ICA decomposition, subtracted the components defined as artifacts from the ICA unmixing matrix using ICLabel, and reconstructed the electrode space data (no MWF step was applied) (Pion-Tonachini et al., 2019). Lastly, 11) wICA_all, which applied wICA to all ICA components, regardless of whether they could be identified as artifacts or brain activity, which is the method initially proposed by Castellanos and Makarov (2006).

### Data

We examined the effectiveness of each cleaning pipeline on a large dataset from healthy participants who had completed a Go-Nogo response inhibition task, which is typically analysed using an ERP approach (N = 127). These EEG files were selected as they were difficult to clean, containing many artifacts reflective of typical EEG data. Data were obtained from participants during cognitive tasks, where participants concentrated (generating frontal and temporal muscle activity and blinks). Bad electrodes were also commonly present in the data, as well as brief electrode disconnections and other atypical artifacts. These data provided an excellent “real world” test for fully automated EEG pre-processing. The Go-Nogo task commonly produces errors of commission, where neural activity time-locked to a mistaken response can be compared to neural activity time-locked to a correct response. Error responses typically produce an event related negativity (ERN), which is a negative voltage deflection from 0 to 150ms post response maximal at fronto-central electrodes and thought to be related to automatic attentional monitoring processes (Bailey, Raj, et al., 2019). Following the ERN, the error positivity (commonly known as the Pe) is commonly analysed, which is a positive voltage deflection from 200 to 400ms post response, maximal at fronto-central electrodes, larger in amplitude and more frontally distributed during error compared to correct responses (Bailey, Raj, et al., 2019). Only participants who provided at least 6 error responses were included in this analysis (N = 73). EEG epochs time-locked to correct responses with the most similar response latency to each individual error response latency within each participant were selected for comparison to the error responses so that the same number of error and correct epochs were submitted to grand averaging for each participant, and results were not confounded by differences in reaction time (Bailey, Raj, et al., 2019). Additionally, analyses of neural activity time-locked to the stimulus for the Go-Nogo task typically focuses on the N2, a negative voltage deflection from 180 to 300ms following stimuli presentation, maximal at fronto-central electrodes and larger for Nogo trials, an ERP which has been proposed to reflect response conflict or conflict monitoring (Bailey, Freedman, et al., 2019). The Go-Nogo task also produces a P3, which is a positive voltage deflection from 300 to 500ms, with a central-parietal maximum during Go trials and a fronto-central maximum during Nogo trials, and has been proposed to reflect sustained attention and response inhibition mechanisms (Bailey, Freedman, et al., 2019). All participants produced >30 correct Go and Nogo responses, so all participants were included in this analysis. All participants provided informed consent prior to participation and the study was approved by the Ethics Committee of the Alfred Hospital and Monash University.

All data were recorded using a Neuroscan amplifier (Compumedics, Melbourne, Australia) with a 64-channel Quickcap (excluding CB1 and CB2 electrodes), a sampling rate of 1000Hz with a 0.01Hz high-pass and 200Hz low-pass filter. The ground electrode was located at AFz, and the reference electrode was located between Cz and CPz. One EEG file was excluded from blink amplitude ratio (BAR) metrics due to not enough blink locked timepoints remaining after cleaning for valid calculation of the BAR. Because we were testing the efficacy of the different cleaning pipelines for application to research involving ERPs, data were high-pass filtered at 0.25Hz as the initial step in all cleaning pipelines.

Various cleaning quality metrics were calculated. Some of the cleaning quality metrics that we assessed were measured from continuous data (for example, the Signal-to-Error Ratios [SER] and Artifact-to-Residue Ratios [ARR], and the proportion of 1 second epochs across all data showing log-power log-frequency slopes reflective of muscle activity after cleaning). However, some cleaning metrics required epoched data, and the measures of each pipeline’s ability to detect differences in ERPs between the conditions’ metrics also required epoched data. As such, after cleaning we interpolated rejected electrodes back into the data (using the EEGLAB ‘pop_interp’ function with the spherical setting) and epoched the data from −400 to 800ms surrounding the stimuli or response of interest. We then applied a typical rejection of remaining bad epochs bad epochs based on max-min voltage values >60 microvolts, or kurtosis / improbable data for all channels >3 or any channel >5, since even the highest performing EEG cleaning pipelines do not completely address all artifacts in all files. For the Go-Nogo data, we baseline corrected data to the pre-stimulus interval using the regression baseline correction method (Alday, 2019). For the error-related epochs, we baseline corrected data using the traditional baseline subtraction method from −400 to −100ms (Bailey, Raj, et al., 2019). This epoching enabled an analysis of differences between the pipelines in the amount of ERP variance explained by differences between commonly analysed experimental conditions (such as between Go and Nogo trials in a response inhibition task), as well as a measure of the proportion of epochs that were removed by the overall cleaning process (including both extreme period rejection and outlier epoch rejection).

### Cleaning Quality Evaluation Metrics

To examine the effectiveness of the RELAX pipeline in cleaning EEG data for ERP analyses, the variants of the RELAX pipeline and comparison pipelines were compared using six different cleaning quality metrics which provided a comprehensive evaluation of various aspects of data integrity and reliability. While these metrics provide a comprehensive and robust assessment of overall cleaning quality (e.g., percentage of muscle activity removed, proportion of epochs removed during cleaning, etc.), these metrics are limited in their ability to provide an estimate of the practical applicability of the pipelines for removing relevant recording artifacts whilst maintaining the underlying neural signal required for scientific research. To address this, the pipelines were also examined using metrics which examined the variance in ERP activity explained by comparisons between different cognitive trial types (using trial types with robust evidence for their differences from the previous literature) and the reliability of typically analysed ERP metrics. We have provided a full description of the cleaning quality evaluation metrics in our companion paper (Bailey et al., 2022) and in our Supplementary Materials (section 3, page 7), and so just provide a brief summary here.

#### EEG Data Cleaning Performance Metrics

The pipelines were compared across six different cleaning quality metrics to provide a comprehensive evaluation of cleaning efficacy, which includes assessment of how each pipeline cleaned the full range of potential artifacts whilst still preserving the neural signal. These metrics have all been used by previous research. Firstly, we included the Signal-to-Error Ratio (SER), which reflects the amount of signal in clean EEG periods that is unaffected by the cleaning process. SER’s contrasting metric is the Artifact-to-Residue Ratio (ARR), which indicates the extent to which all artifacts were reduced. High values for both the SER and ARR indicate effective cleaning, as it is easy but unhelpful to obtain high SER and low ARR values (by not reducing artifacts) or high ARR and low SER (by severely reducing artifacts but not being concerned about preserving neural activity) (Bertrand, 2015; Somers & Bertrand, 2016; Somers et al., 2018).

Next, we measured the ratio of the absolute EEG amplitude in blink affected periods compared to surrounding non-blink periods after cleaning. We did this in two ways: averaged across the frontal electrodes affected by blinks (fBAR) and averaged across all electrodes (allBAR). For this metric, values of ~1 reflect optimal performance, where after cleaning, activity time-locked to blinks is the same amplitude as non-blink activity, while values <1 reflect overcleaning, and values >1 reflect under cleaning (Robbins et al., 2020). Next, in order to assess the efficacy of the pipelines for muscle artifact cleaning, we obtained a measure of the proportion of epochs that showed log-frequency log-power slopes >−0.59, which have been shown by comparisons of EEG recordings taken from paralysed scalps against typical EEG recordings to indicate muscle activity (Fitzgibbon et al., 2016). We also tested the degree to which these slopes exceeded the muscle activity slope threshold within epochs that showed muscle activity remaining after cleaning (for brevity, these results are only reported in the Supplementary Materials, section 4, page 26). Higher values reflect poorer performance for the muscle activity metrics. The final artifact rejection metric assessed was the proportion of epochs rejected by the overall cleaning process. While to some extent the proportion of epochs removed by cleaning is dependent on the raw data quality, less effective cleaning pipelines leave more artifacts remaining after cleaning, which are rejected by the final epoch rejection step, providing less data available for analysis after cleaning. As such, more effective data cleaning should provide more epochs for analysis. Given that all pipelines were tested on the same EEG files, raw data quality did not vary between pipelines, so differences in the proportion of epochs removed by the cleaning process across different pipelines were fully determined by the cleaning performance of the pipelines. Higher quality data with more epochs for analysis has been demonstrated to be more important for statistical power even than a larger overall sample size (Kolossa & Kopp, 2018). As such, within our study, a lower proportion of epochs rejected suggested a better performing pipeline.

#### ERP Trial Type Comparisons - Variance Explained Metrics

In addition to the assessments of the efficacy of each pipeline in reducing artifacts, we compared the amount of variance explained by two common experimental comparisons, across four different ERPs. We chose to analyse the Go-Nogo N2 and P3, and the error processing related ERN and Pe, which are well demonstrated by previous research to differ between the two conditions of interest providing an important assessment of the practical applicability of each cleaning pipeline (Clayson, Baldwin, et al., 2021). For these measures of the variance explained by the difference between experimental conditions, higher values reflect better performance, providing a better chance to detect statistically significant differences within a study. Details of how these ERPs were computed can be found in the Supplementary Materials (section 3, page 11).

#### ERP Amplitude Reliability Metrics

Finally, we performed two tests of outcome measure reliability. The calculation of these measures is explained in detail in the Supplementary Materials (section 3, page 13). In brief, we assessed the number of trials to reach sufficient dependability of the Pe following error responses in the Go-Nogo task (Clayson, Carbine, et al., 2021; Clayson & Miller, 2017). Secondly, we examined the bootstrapped standardised measurement error (bSME) of the single electrode N1, N2 and P3 peak amplitude from the Go-Nogo task. The bSME is a measure that has been proposed to reflect data quality (precision) for an averaged ERP waveform for an individual participant (Luck et al., 2021). This measure has been proposed to indicate the combined impact of all single-trial EEG noise and the number of epochs available for analysis on outcome measures (Luck et al., 2021). Lower values indicate better performance, but bSME values should also be considered in relation to the ERP amplitude (with low bSME values and high amplitude values indicating better performance). The authors of the bSME metric suggest that the measure has the potential to objectively determine which “analysis procedures produce the cleanest data” (Luck et al., 2021). We assessed bSME from peak amplitudes within the N2 window following Nogo trials at FCz, and the P3 window following Go trials at Pz. In order to assess the potential applicability of each pipeline to early sensory processing related ERPs, we additionally analysed the bSME of the N1 at FCz, which is a negative deflection maximal at fronto-central electrodes that follows both Go and Nogo trials equally, and is likely to reflect stimulus processing rather than any response inhibition relevant process (Bokura et al., 2001). Time-periods of interest for the bSME measure were 60 to 180ms for the N1, 180 to 300ms for the N2, and 300 to 500ms for the P3. We used peak amplitudes as they are more vulnerable to artifacts than mean amplitudes within the time-period of interest, and as such peak amplitude was a more sensitive measure of data cleaning efficacy (Clayson et al., 2013).

### Statistics

To compare the different artifact cleaning metrics between the cleaning pipelines, we used the robust repeated measures ANOVA function “rmanova” from the WRS2 package in R (R Core Team v4.0.4) (Mair & Wilcox, 2020). These tests are robust against violations of the normality and homoscedasticity assumptions of parametric statistical tests, while providing equivalent power (Mair & Wilcox, 2020). When omnibus ANOVAs were significant, pairwise comparisons between individual cleaning pipelines were performed using the robust post-hoc t-test function “rmmcp”. This test applies multiple comparison controls across the post-hoc tests using Hochberg’s approach (Mair & Wilcox, 2020). Experiment-wise multiple comparison controls (across the omnibus ANOVAs from each separate metric) were not used – our judgement was that when testing for optimal EEG cleaning methods, it was more important to provide sensitivity to differences in cleaning outcomes than to protect against false positive results. We tested a large number of metrics to demonstrate which pipelines provide the best performance across all artifact cleaning and experimental outcome metrics (rather than for a single artifact type as is more common in the literature). If we applied experiment-wise multiple comparison controls, this large number of comparisons would severely reduce our study’s sensitivity to detect differences between the pipelines, reducing our potential to draw conclusions about the optimal EEG cleaning pipeline, which would have adversely incentivised us to test a smaller range of metrics. As such, we opted to preference sensitivity over applying extensive controls for false positives, as is recommended when sensitivity is the priority for a study (Bender & Lange, 2001).

In addition to the artifact cleaning metrics, we compared the amount of variance (np^2^) explained by ERP comparisons between different conditions after the EEG data was cleaned. To test for differences between pipelines in np^2^ for the experimental manipulations, we performed repeated measures ANOVA design tests of the interaction between every potential pair of pipelines and the two experimental conditions using the randomised graphical user interface (RAGU) global field potential (GFP) and topographical ANOVA (TANOVA) with L2 norm tests. These tests measure differences in the overall neural response amplitudes, and the distribution of neural activity after normalization for amplitude respectively (Habermann et al., 2018; Koenig et al., 2011). We explain these statistics in more detail in the Supplementary Materials (section 3, page 11) and provide heat maps displaying np^2^ values for each interaction in the Supplementary Materials (section 4, page 32-55), with multiple comparison controls applied across the post-hoc tests within each omnibus ANOVA using the false discovery rate (FDR-p) (Benjamini & Hochberg, 1995).

To visualize the data, we have provided raincloud plots for the reader to see the full data with minimal distortion (Allen et al., 2019). These plots include outliers, which importantly indicate individual files where a cleaning pipeline might not have properly reduced an artifact. We have also provided a rank order of the means from each pipeline for each metric with significant differences noted (Table 4). Means and SD tables and specific details of post-hoc tests can be found in our Supplementary Materials (section 4).

**Table 3.** Description of cleaning quality evaluation metrics used for comparing pipelines. Results from pipelines marked by * are not included in the main manuscript but can be viewed in the Supplementary Materials. bSME – bootstrapped standardised measurement error.

| Metric Type | Outcome Metric | Summary of Metric Purpose | Evaluation of Metric |
| --- | --- | --- | --- |
| EEG Data Cleaning Performance Metrics | Signal to Error Ratio (SER) | Reflects the amount of signal in clean EEG periods that is unaffected by the cleaning process | Higher values reflect better cleaning |
|  | Artifact to Residue Ratio (ARR) | Indicates the extent to which artifacts in artifact periods were reduced | Higher values reflect better cleaning |
| | Frontal Electrodes Blink Artifact Ratio (fBAR) | Indicates the ratio of absolute EEG amplitude in blink-affected periods to non-blink periods after cleaning, averaged across frontal electrodes. | Values $\sim 1$ reflect optimal cleaning, values $< 1$ indicate overcleaning, and values $> 1$ reflect under cleaning. |
|  | All Electrodes Blink Artifact Ratio (allBAR)* | Indicates the ratio of absolute EEG amplitude in blink-affected periods to non-blink periods after cleaning, averaged across all electrodes. |  |
|  | Proportion of Epochs Showing Muscle Activity Remaining After Cleaning | Indicates the proportion of epochs which still contain log-power log-frequency slopes indicating muscle activity after cleaning. | Lower values indicate better performance |
|  | Severity of Muscle Activity from Epochs Showing Muscle Activity Remaining After Cleaning* | Indicates how severe the remaining muscle artifact is after cleaning if it has not been completely removed | Lower values indicate better performance |
|  | Proportion of Epochs Removed by Cleaning | Reflects the proportion of epochs removed during the cleaning process. | Lower values indicate better performance |
| ERP Trial Type Comparisons - Variance Explained Metrics | Variance Explained by Error vs Correct Responses | Reflects the degree to which the variance in the pattern of neural activity is explained by differences between error and correct responses when analysing the cleaned data. | Higher values reflect better performance |
|  | Variance Explained by Go vs Nogo trials | Reflects the degree to which the variance in the pattern of neural activity is explained by differences between Go and Nogo stimuli when analysing the cleaned data. | Higher values reflect better performance |
| ERP Amplitude Reliability Metrics | Number of Trials Required for Dependable Analysis of the Pe | Indicates the number of error trials required to achieve a dependable analysis of the Pe ERP | Lower values indicate better performance |
|  | ERP Amplitude and Single Trial Standard Error of the Mean. | Indicates data precision for the averaged ERP waveform, reflected as the bSME for peak amplitudes within the:<br>1. N2 window at FCz for Nogo trials,<br>2. P3 window at Pz for Go trials, and<br>3. N1 window at FCz for both Go and Nogo trials | Lower values indicate better performance |

**Table 4.**
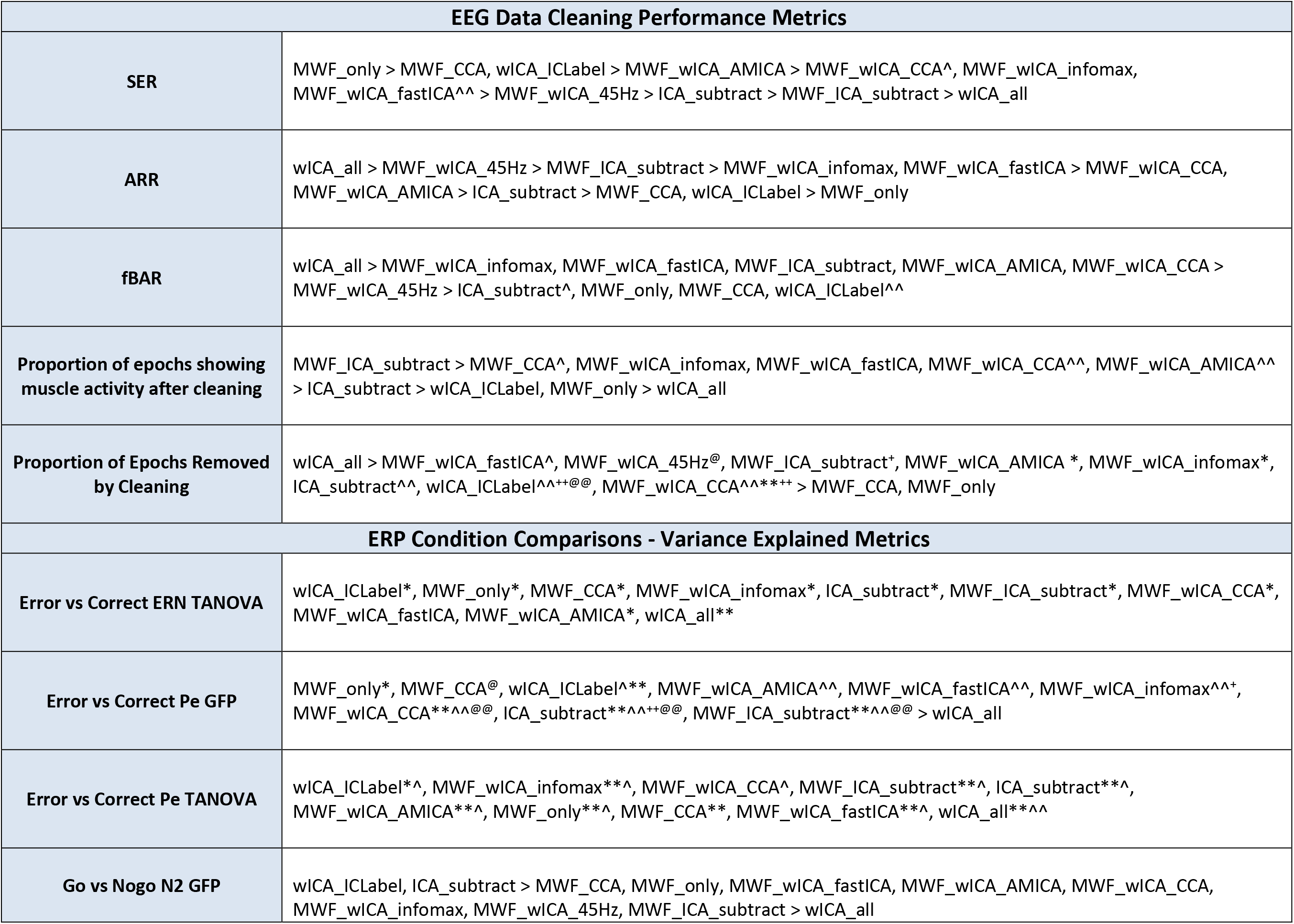
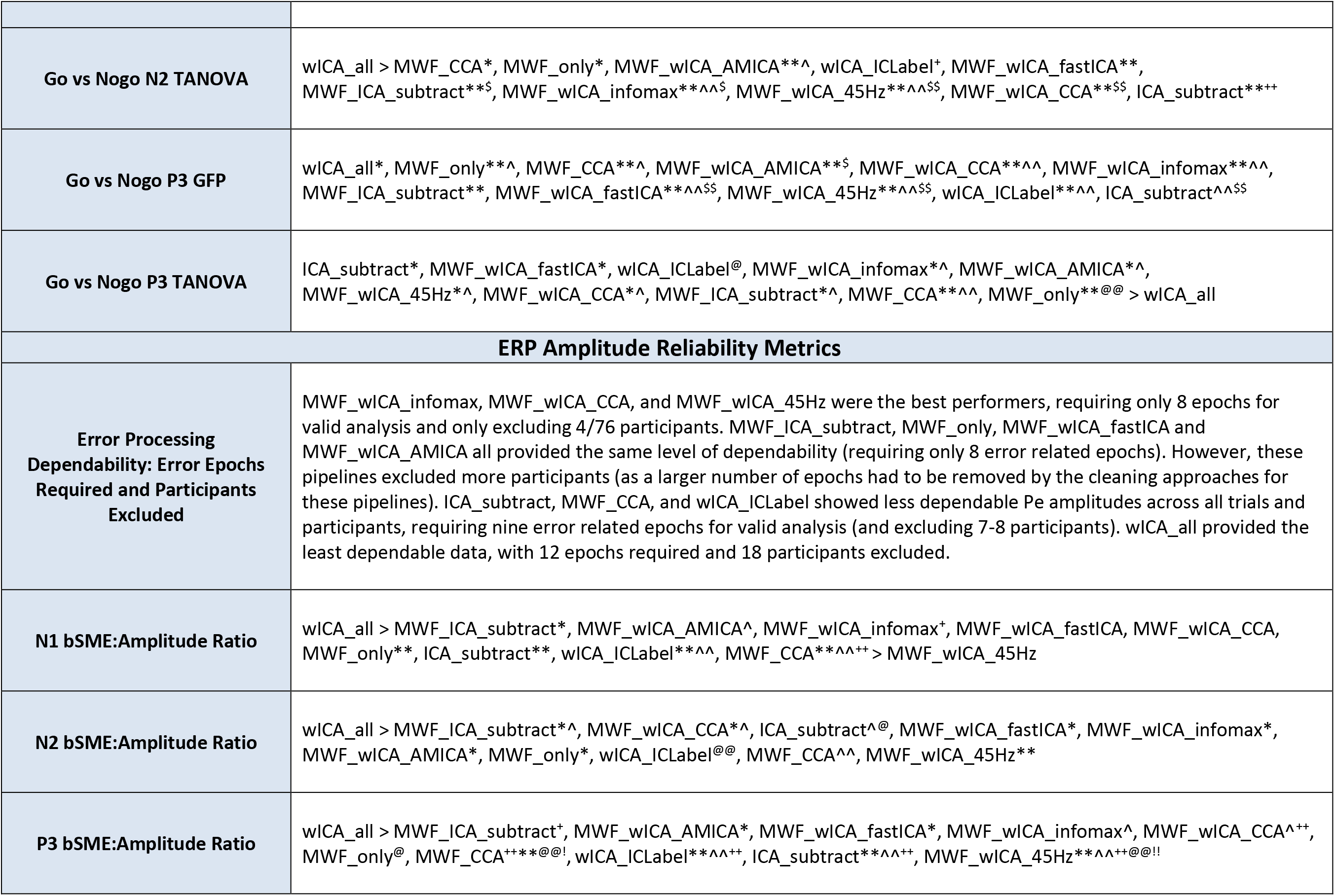
Rank order (by mean) of best performing pipelines to worst performing pipelines for the ERP analysis datasets. Significant differences are highlighted for pipelines that performed significantly better than other pipelines using the following notation for ease of understanding: better performance > worse performance (rather than higher values > lower values). Because sometimes pipeline 1 differed from pipeline 2, but pipeline 3 did not differ from either 1 or 2, we have used the following notation: ^ = significantly higher than the pipeline marked with a ^^ within the same section (while the others in the category are not significantly different from each other). * = significantly higher than the pipeline marked with a ** in the same category, and so on for the following symbols: ^+@$!+^.

## Results

For every metric a significant difference was present in the omnibus ANOVA (all p < 0.01). For brevity, the full statistics are reported in the Supplementary Materials (section 4), and a rank order of the means from each pipeline for each metric with significant post-hoc t-test differences noted can be viewed in Table 4. We provide here a narrative summary of the results most relevant to the evaluation of cleaning efficacy and the selection of an optimal pipeline for EEG cleaning and detection of experimental effects.

### EEG Data Cleaning Performance Metrics

#### Signal-to-Error Ratio and Artifact-to-Residue Ratio

SER and ARR values can be viewed in Figure 2. When SER and ARR values were viewed together (Figure 3), it was apparent that MWF_wICA and MWF_wICA_CCA performed higher than ICA_subtract in the SER metric, while at the same time they performed better than ICA_subtract in the ARR metric. MWF_ICA_subtract provided higher ARR values than MWF_wICA, but at the expense of lower SER values. wICA_all performed the best at removing artifacts (with very high ARR values), but this came at the expense of very low SER values suggesting considerable removal of signal from clean periods of the EEG data. Inversely, MWF_only, MWF_CCA, and wICA_ICLabel showed high SER values but lower ARR values than other pipelines suggesting less effective reduction of artifact periods. MWF_only performed the best of these three pipelines at providing high SER while producing a similar value to MWF_CCA and wICA_ICLabel for ARR. Finally, note that wICA_ICLabel strongly outperformed ICA_subtract in SER, while ICA_subtract showed only a small advantage over wICA_ICLabel in ARR, suggesting wICA_ICLabel may be the superior pipeline when viewing SER and ARR concurrently. This pattern of results is identical to that shown in our companion article, which outlined the application of RELAX for cleaning EEG data for analysis of oscillations (Bailey et al., 2022). The full details of the SER and ARR results can be found in the Supplementary Materials (section 4, pages 14-19).

**Figure 2.**
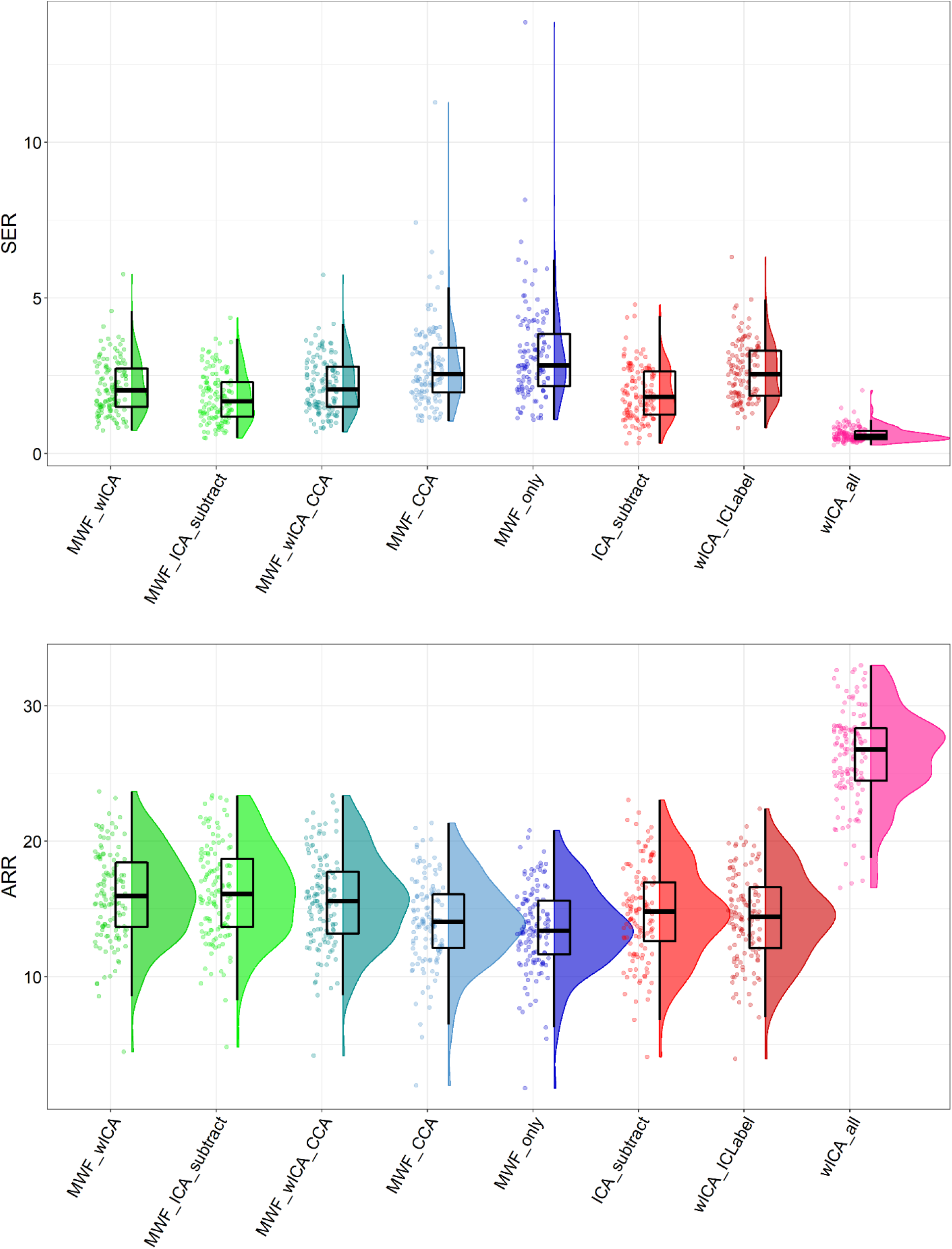
Raincloud plots depicting Signal-to-Error Ratio (SER) and Artifact-to-Residue-Ratio (ARR) values for each of the cleaning pipelines.

**Figure 3.**
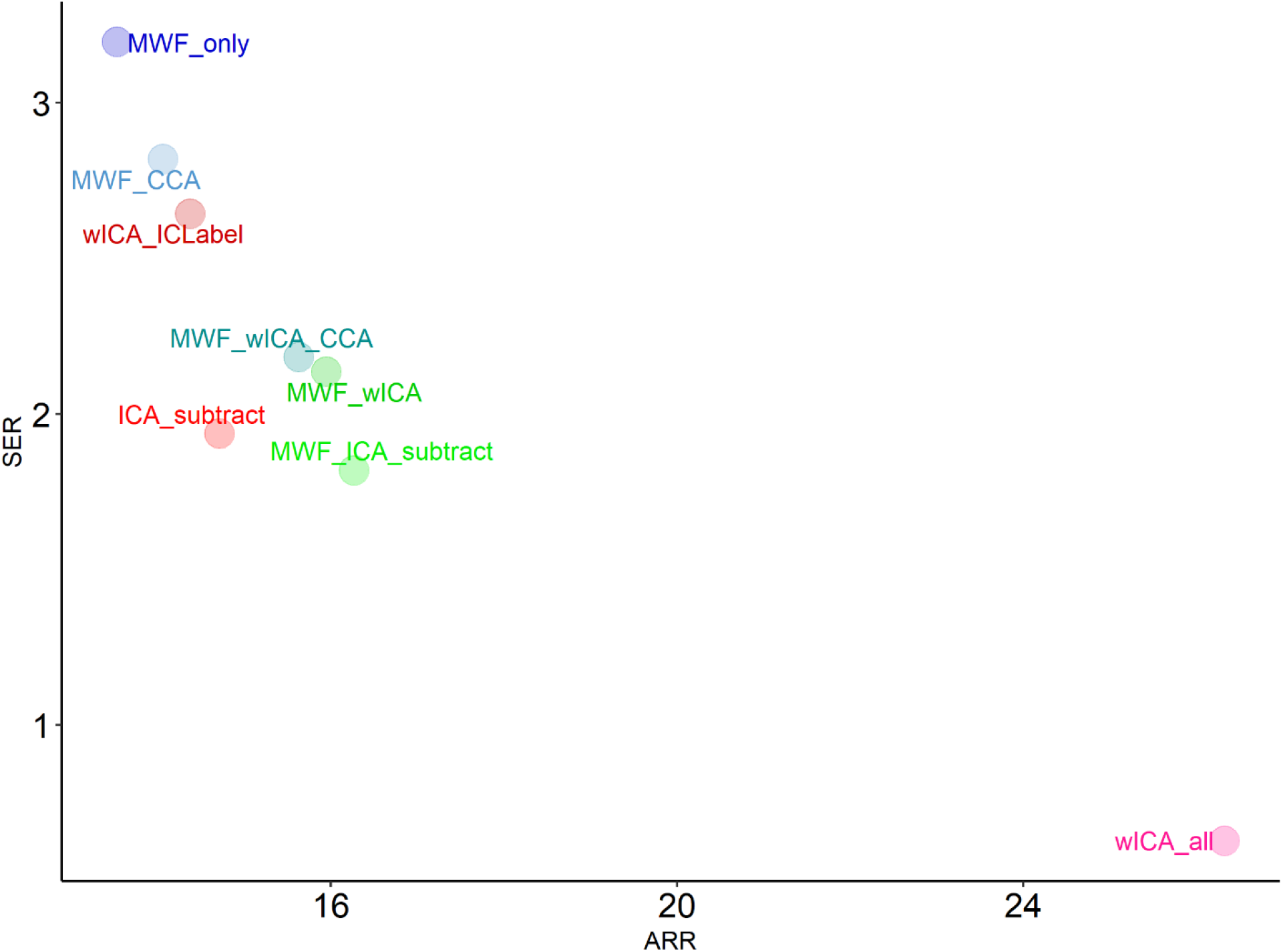
A scatterplot depicting both mean Signal-to-Error Ratio (SER) and Artifact-to-Residue-Ratio (ARR) values from each cleaning pipeline.

#### Blink Amplitude Ratios

Mean fBAR values were between 1.11 and 1.19 for each pipeline, suggesting that after cleaning, periods of EEG data that had originally been contaminated by blink artifacts still showed absolute voltage amplitudes 11-19% larger than the surrounding data (Figure 4). Interestingly, these fBAR values from the 0.25Hz high-pass filtered dataset were larger than when data was 1Hz high-pass filtered for all cleaning pipelines (see our companion paper for details (Bailey et al., 2022), mean values for the 1Hz high-pass filtered datasets were between 1.015 and 1.077 after cleaning by the same pipelines). Additionally, almost all pipelines contained 1-3 EEG files with outlier fBAR values > 2.25. We suspect this reflects the established lower performance of ICA when data is high-pass filtered below 1Hz (Winkler et al., 2015). However, we also note that the MWF_only and MWF_CCA methods (which did not apply ICA) similarly showed worse performance on the 0.25Hz high-pass filtered data. This finding that has not previously been demonstrated as far as we are aware but suggests that MWF cleaning unfortunately does not offer a complete escape from the limitations on effective EEG cleaning imposed by using filter settings that are appropriate for analysis of ERP datasets. In terms of the comparisons between pipelines, wICA_all performed significantly better than all other pipelines for fBAR. MWF_wICA, MWF_ICA_subtract and MWF_wICA_CCA were the next best performers, and ICA_subtract, MWF_only, MWF_CCA and wICA_ICLabel were the worst performing pipelines. A similar pattern of results was present for the allBAR values, with the exception that MWF_only and MWF_CCA outperformed ICA_subtract and wICA_ICLabel, and did not differ from MWF_wICA, MWF_wICA_CCA, or MWF_ICA_subtract (see Supplementary Material Figure S9-10, section 4, page 21-22).

**Figure 4.**
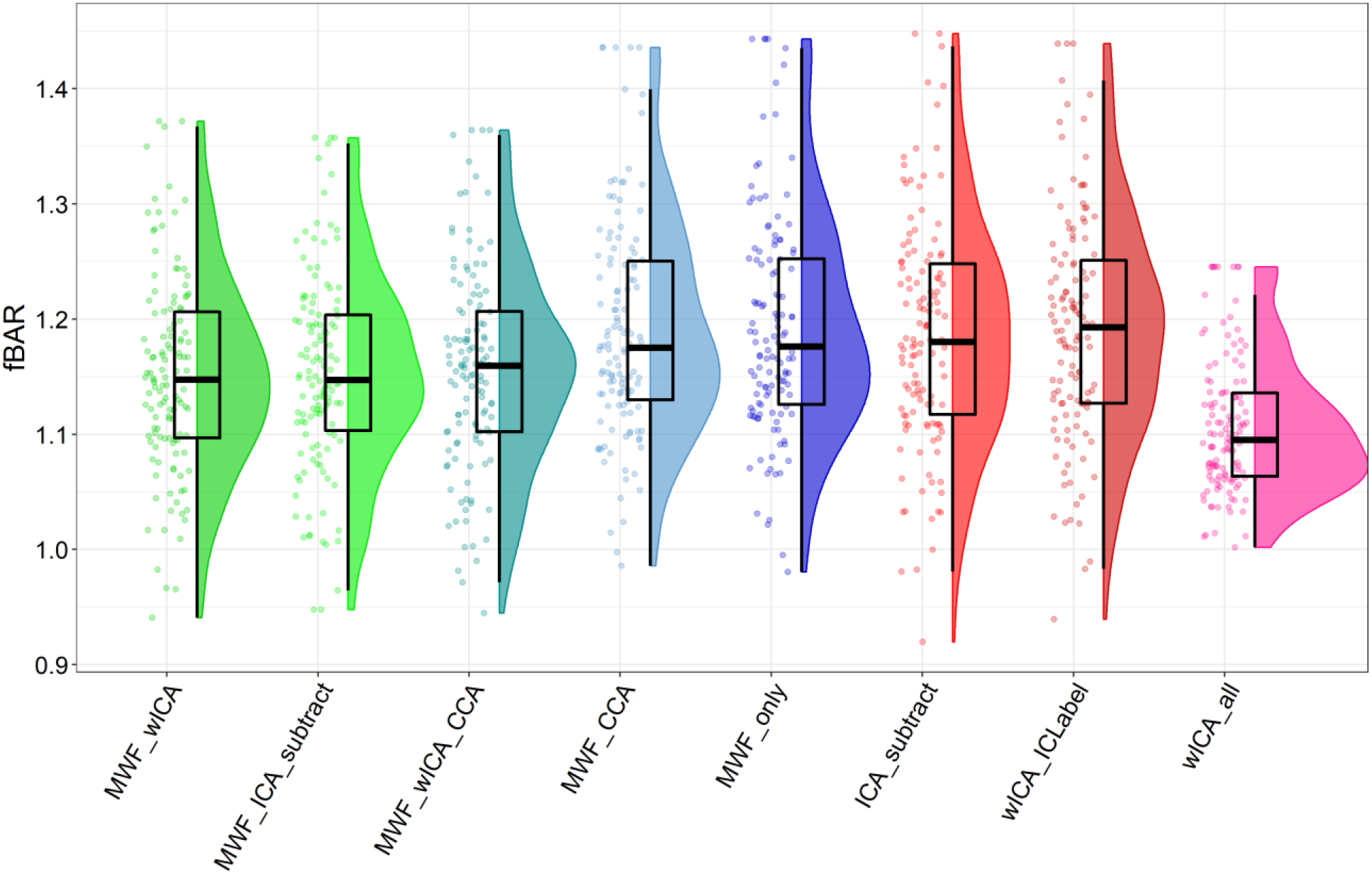
Raincloud plot depicting frontal blink amplitude ratio (fBAR) values for each of the cleaning pipelines. Note that this data has been winsorized to present a scale that enables visualisation of differences between pipelines – almost all pipelines contained 1-3 EEG file outliers with fBAR > 2.25. Plots depicting all unmodified data can be viewed in Supplementary Materials Figure S8.

#### Proportion of Epochs Showing Muscle Activity Remaining After Cleaning

MWF_ICA_subtract outperformed all other pipelines in terms of the proportion of epochs showing muscle activity remaining after cleaning (almost 0 for most EEG files) (Figure 5). MWF_CCA and MWF_wICA performed the next best (with values between 0 and 0.025 for most EEG files, but some outlying files showed more epochs contaminated by muscle activity), followed by MWF_wICA_CCA, then ICA_subtract. wICA_ICLabel and MWF_only performed the worst, except for wICA_all which showed almost all epochs with log-power log-frequency slopes indicating muscle activity after cleaning (post-hoc tests can be viewed in the Supplementary Materials Figure S15, page 26). This may be because wICA_all seems to remove considerable low frequency data, which may flatten the log-power log-frequency slope to the extent that most epochs have a slope suggesting muscle activity remaining. However, the wICA_all cleaned data does also show a bump in power in the beta frequency power ranges, which no other pipeline shows to the same extent (see our companion article for more details (Bailey et al., 2022)). With regards to the severity of muscle slopes that exceeded the threshold after cleaning, the overall rank order of pipeline performance showed that MWF_ICA_subtract, MWF_wICA and MWF_CCA performing the best, not significantly different from each other (further details can be viewed in the Supplementary Materials, section 4, page 26-28).

**Figure 5.**
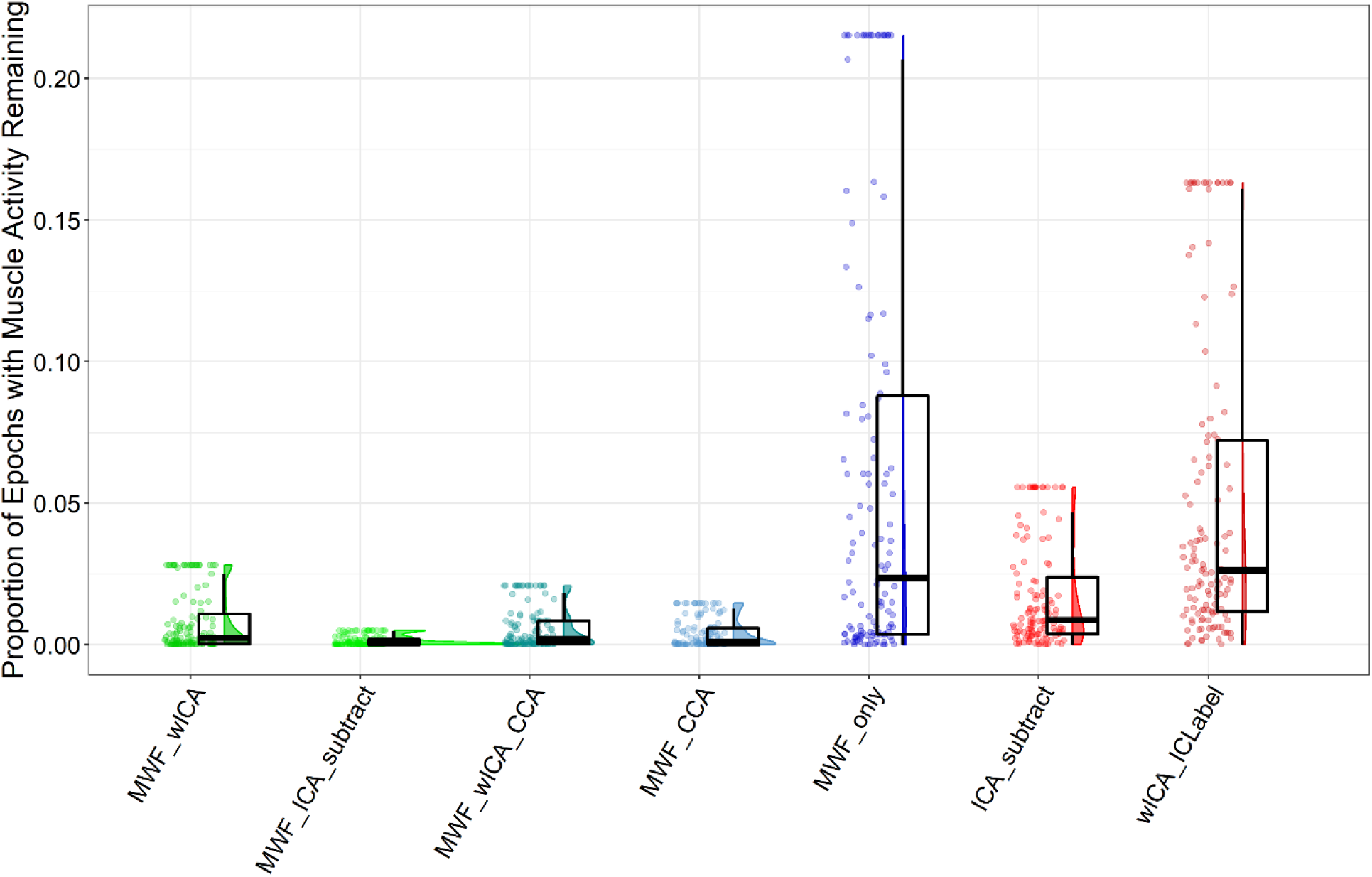
Raincloud plot depicting the proportion of epochs showing log-frequency log-power values above the −0.59 threshold for each of the cleaning pipelines. Note that this figure excludes wICA_all, as this pipeline showed median values > 0.75 and made the scale of the graph such that it was difficult to visualise differences in the other pipelines. Note also that we have winsorized the data in the figure, as the outliers also made the scale such that it was difficult to visualise differences in the other pipelines. The full data can be viewed in the Supplementary Materials Figure S14, page 26.

#### Proportion of EEG Epochs Deleted by the Cleaning Pipeline

wICA_all led to the lowest proportion of epochs removed during cleaning, significantly less than all other pipelines (Figure 6). This was followed by MWF_ICA_subtract, MWF_wICA, ICA_subtract, wICA_ICLabel and MWF_wICA_CCA which did not differ from each other (except for MWF_ICA_subtract, which outperformed wICA_ICLabel and MWF_wICA_CCA, and MWF_wICA which outperformed MWF_wICA_CCA). The previously mentioned pipelines also outperformed MWF_CCA and MWF_only which showed the worst performance. Post-hoc tests for this metric can be viewed in Supplementary Materials Figure S21, page 31.

**Figure 6.**
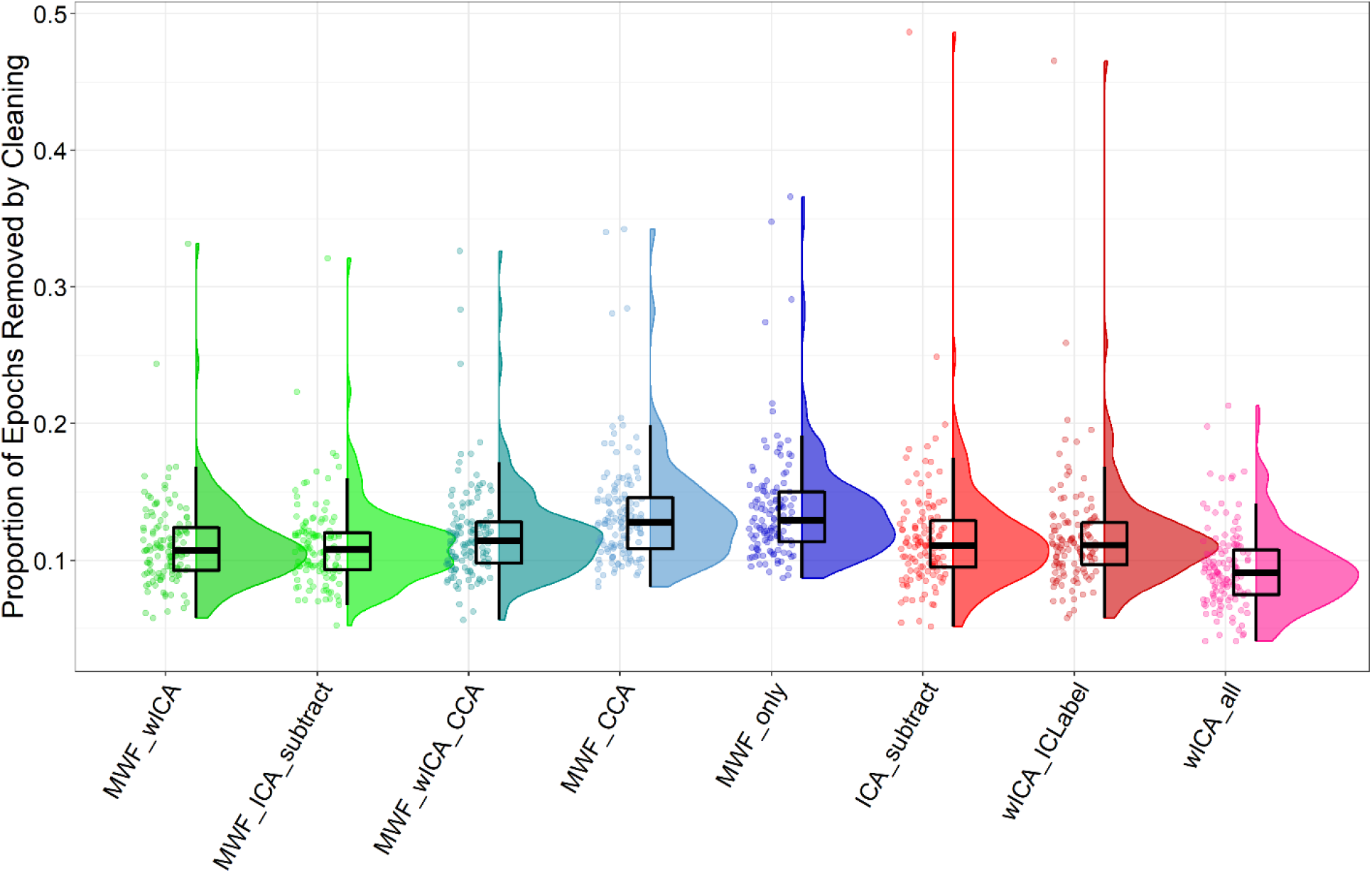
Raincloud plot depicting the proportion of timepoints in the data rejected for each of the cleaning pipelines.

### ERP Condition Comparisons - Variance Explained Metrics

#### Variance Explained by Error vs Correct Responses

ERP waveforms from FCz for error and correct responses can be viewed in Figure 7, and results for each variance explained metric and pipeline can be viewed in Figure 8. With regards to the ERN TANOVA all pipelines provided np^2^ values ~0.30 except for wICA_all (np^2^ = 0.23). Results indicated that none of the pipelines significantly differed from each other, except for wICA_all, which led to significantly less explained variance between errors and correct responses, and showed a different distribution of activity compared to other pipelines (Figures 9–10). With regards to the Pe GFP, all pipelines provided np^2^ values between 0.45 and 0.55, except for wICA_all which provided np^2^ = 0.29. MWF_only and MWF_CCA showed the best performance, followed by wICA_ICLabel, which performed slightly better than MWF_wICA and MWF_wICA_CCA. ICA_subtract and MWF_ICA_subtract performed the worst except for wICA_all. With regards to the Pe TANOVA, all pipelines provided np^2^ values between 0.18 and 0.28. wICA_ICLabel showed the best performance, significantly better than all other pipelines except for MWF_wICA_CCA. All other pipelines showed performance that did not differ from any other pipeline, except for wICA_all, which showed less variance explained than all other pipelines except for MWF_CCA. Visual inspection of the topoplots indicated all pipelines showed a similar pattern for the ERN and for the Pe, with no obvious indication of the reason for the differences in explained variance between the pipelines, with the exception of wICA_all (full details for these metrics can be found in the Supplementary Figure section 4, page 32-41).

**Figure 7.**
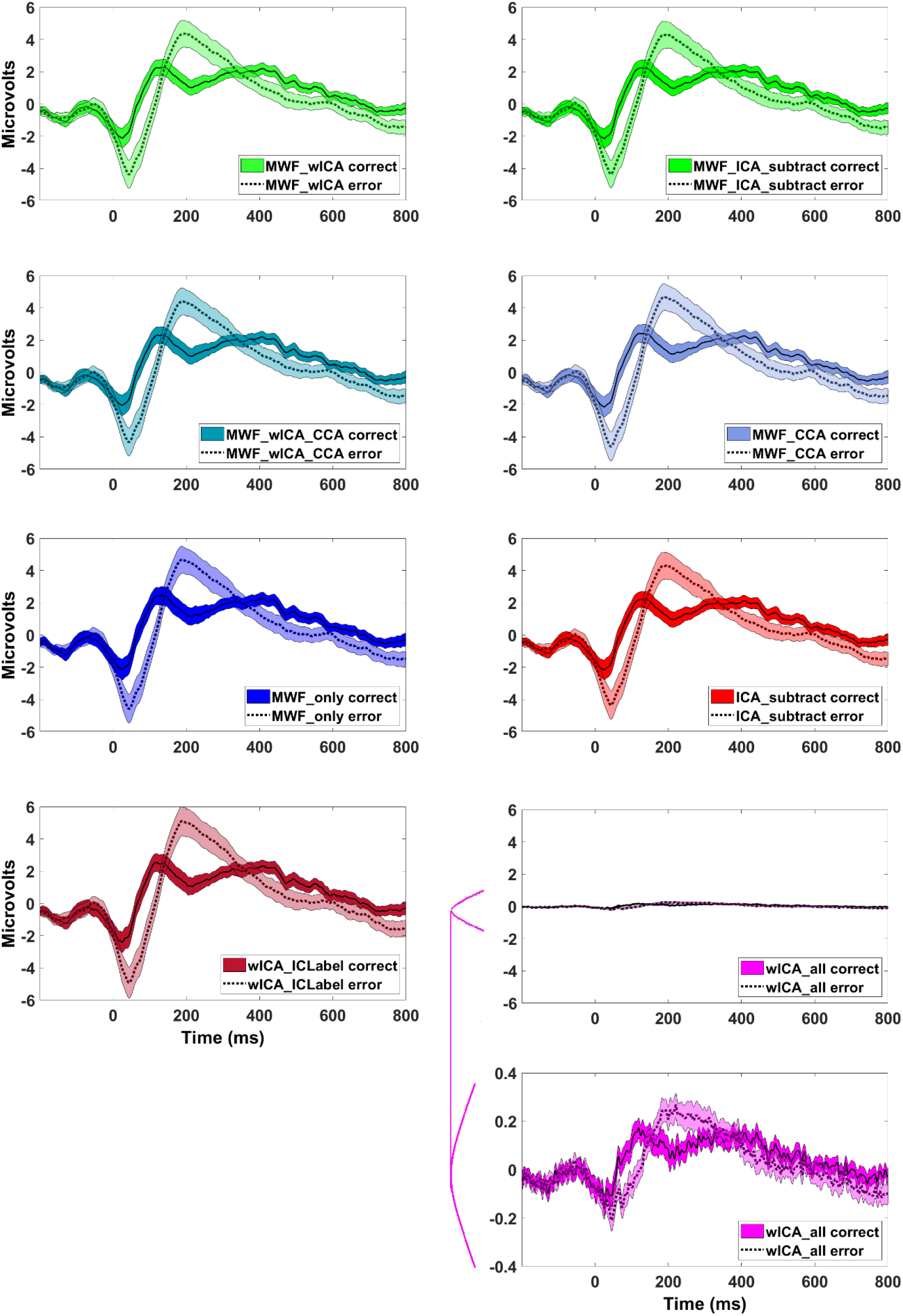
Grand average error and correct related ERP responses from FCz for each pipeline. Shading reflects 95% confidence intervals. Note the different scale required to see the ERP for wICA_all.

**Figure 8.**
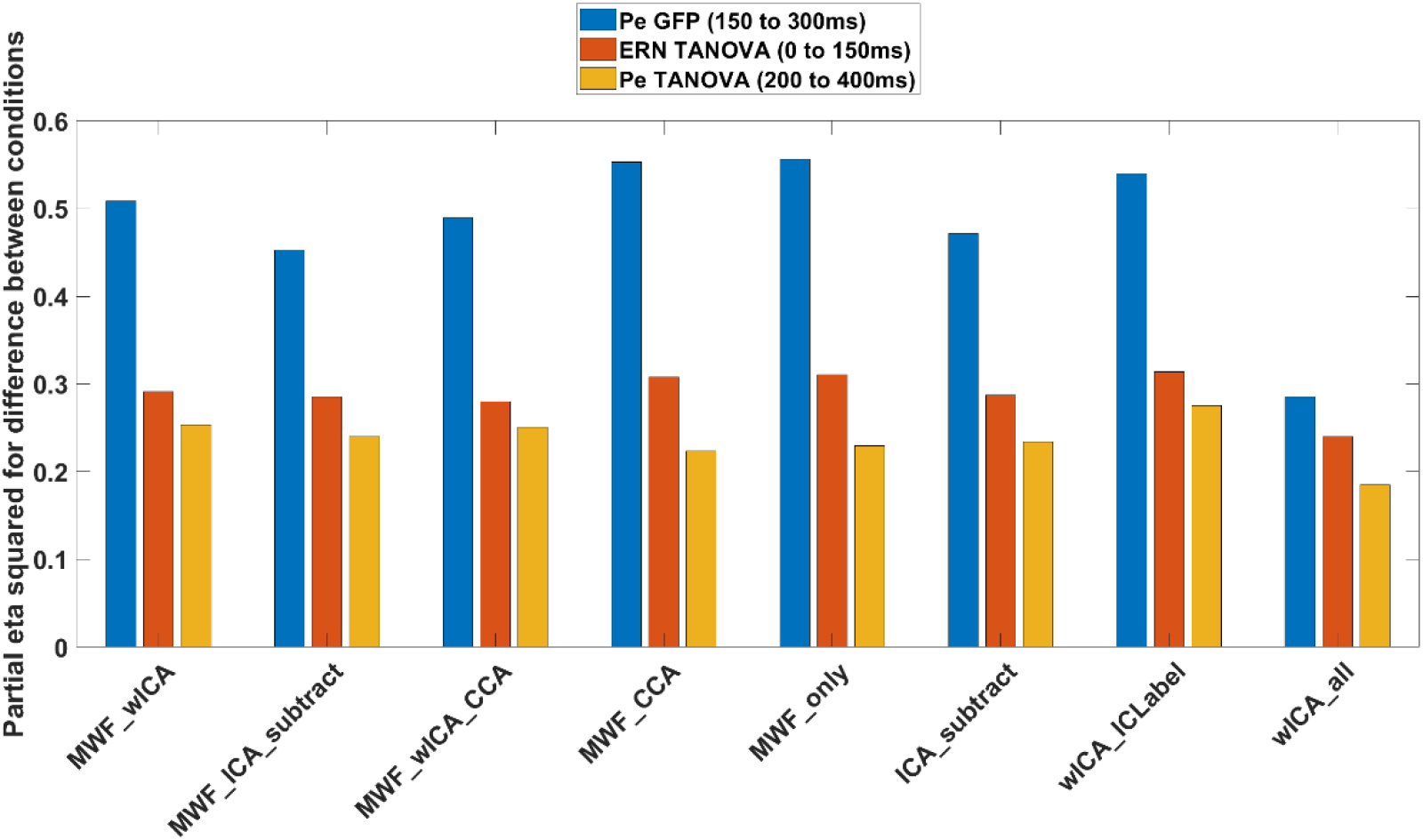
Displays partial eta squared (np^2^) values for each pipeline, indicating the variance explained by the difference between error and correct trials in the distribution of the ERN (0 to 150ms after a response) and the Pe (200 to 400ms after a response), as well as the GFP of the Pe (150 to 300ms after a response) for each of the cleaning pipelines.

**Figure 9.**
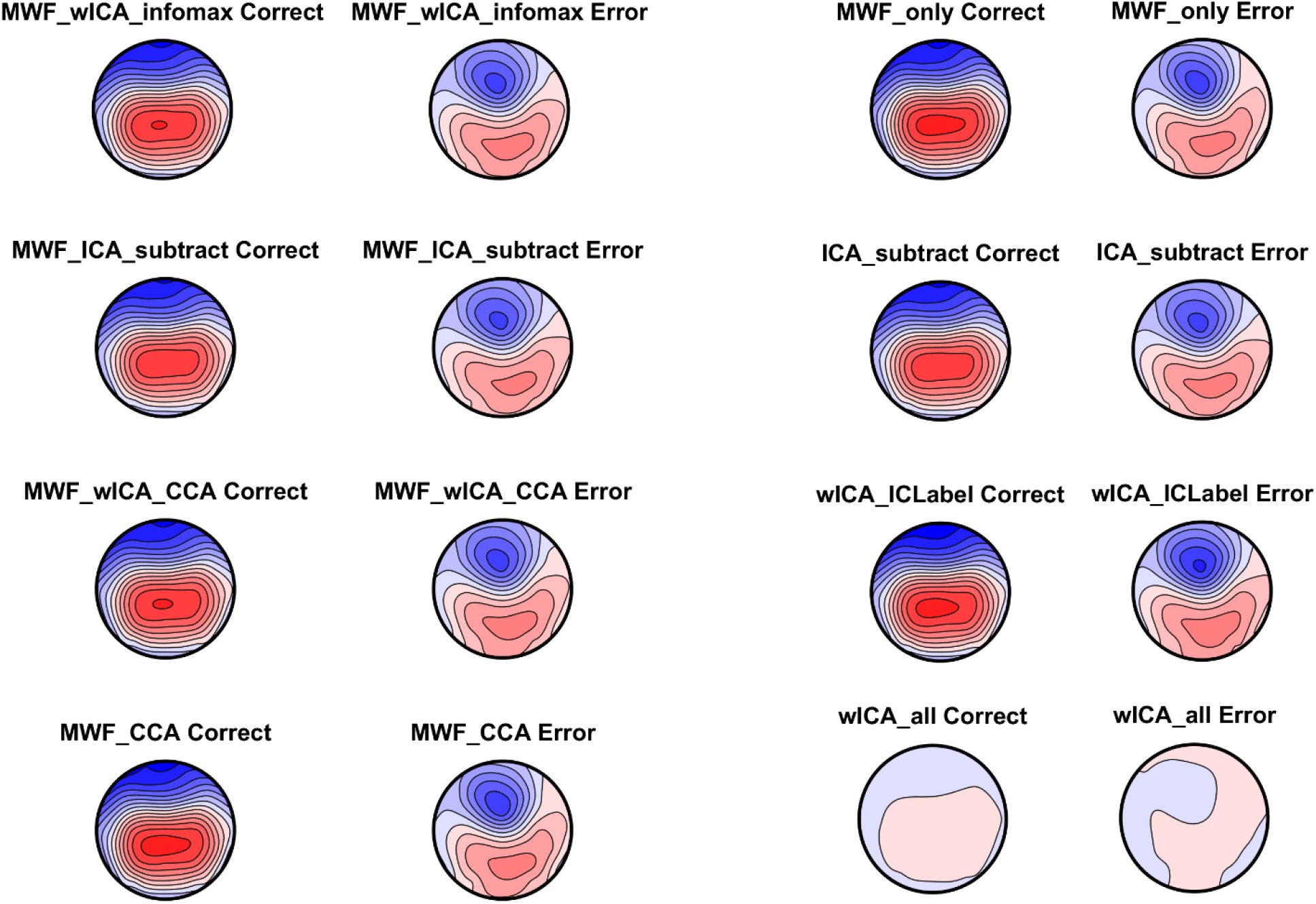
Topoplots of the averaged ERN window for correct and error responses for each pipeline. All plots are on the same scale so comparisons can be made across pipelines / conditions. Note the similarity across the majority of pipelines, with only wICA_all displaying much lower voltage amplitudes across the entire scalp.

**Figure 10.**
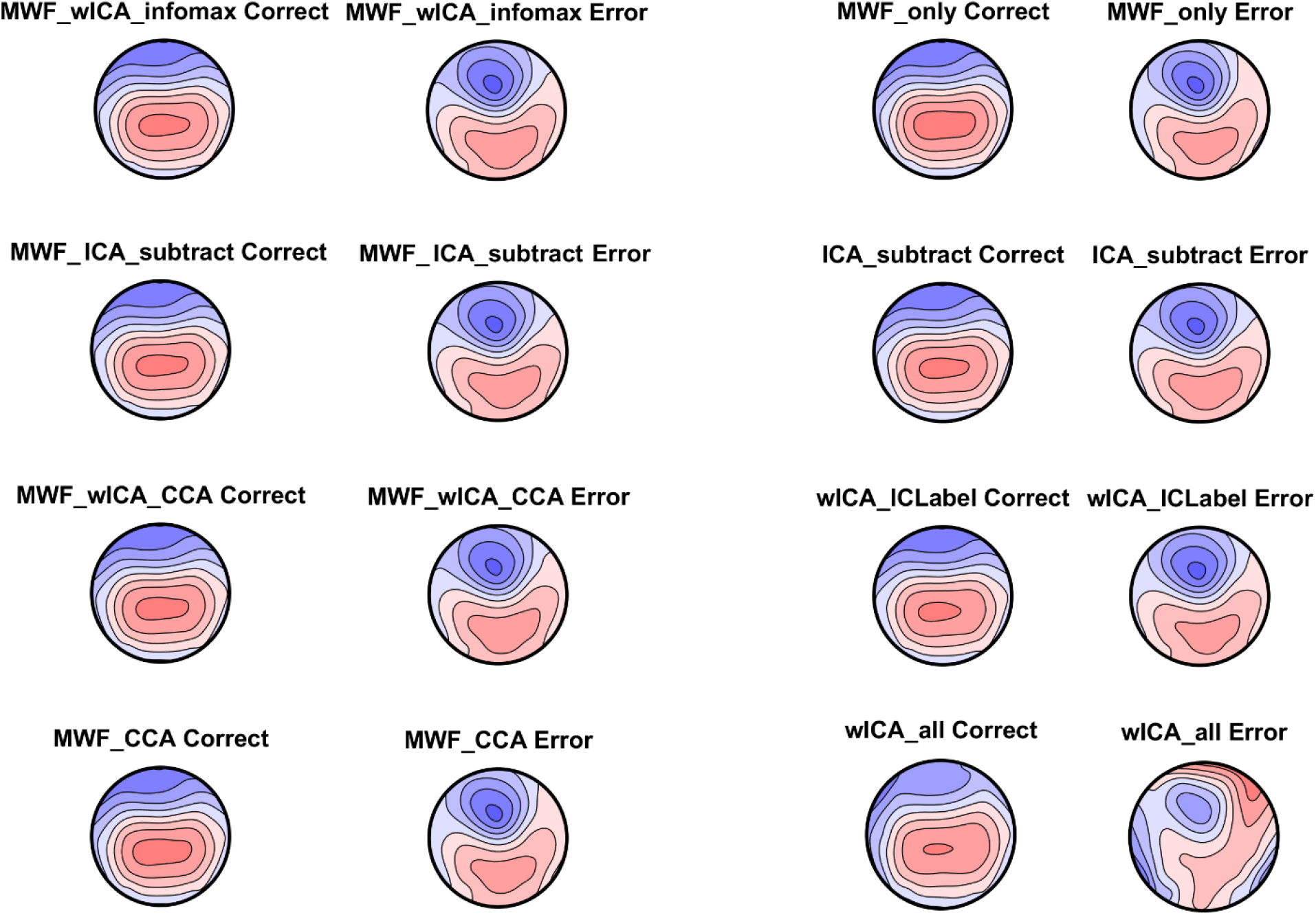
Topoplots of the averaged ERN window for correct and error responses for each pipeline. All plots are on their own scale so the distribution within each pipeline / condition can be viewed. Note the similarity across the majority of pipelines, with only wICA_all showing a different pattern of results.

#### Variance Explained by Go vs Nogo Trials

With regards to the N2 GFP most pipelines provided np^2^ values from 0.28 to 0.4, with wICA_ICLabel and ICA_subtract showing the best performance, followed by MWF_CCA, MWF_only, MWF_wICA, MWF_wICA_CCA, and MWF_ICA_subtract, with wICA_all showing the worst performance (see Figures 11 and 12). With regards to the N2 TANOVA all pipelines provided np^2^ values between 0.15 to 0.23, with wICA_all showing the best performance, followed by MWF_CCA, MWF_only, wICA_ICLabel, MWF_ICA_subtract, MWF_wICA, then MWF_wICA_CCA, and lastly ICA_subtract. It is also worth noting that all methods that combined MWF and wICA showed lower performance than MWF alone.

**Figure 11.**
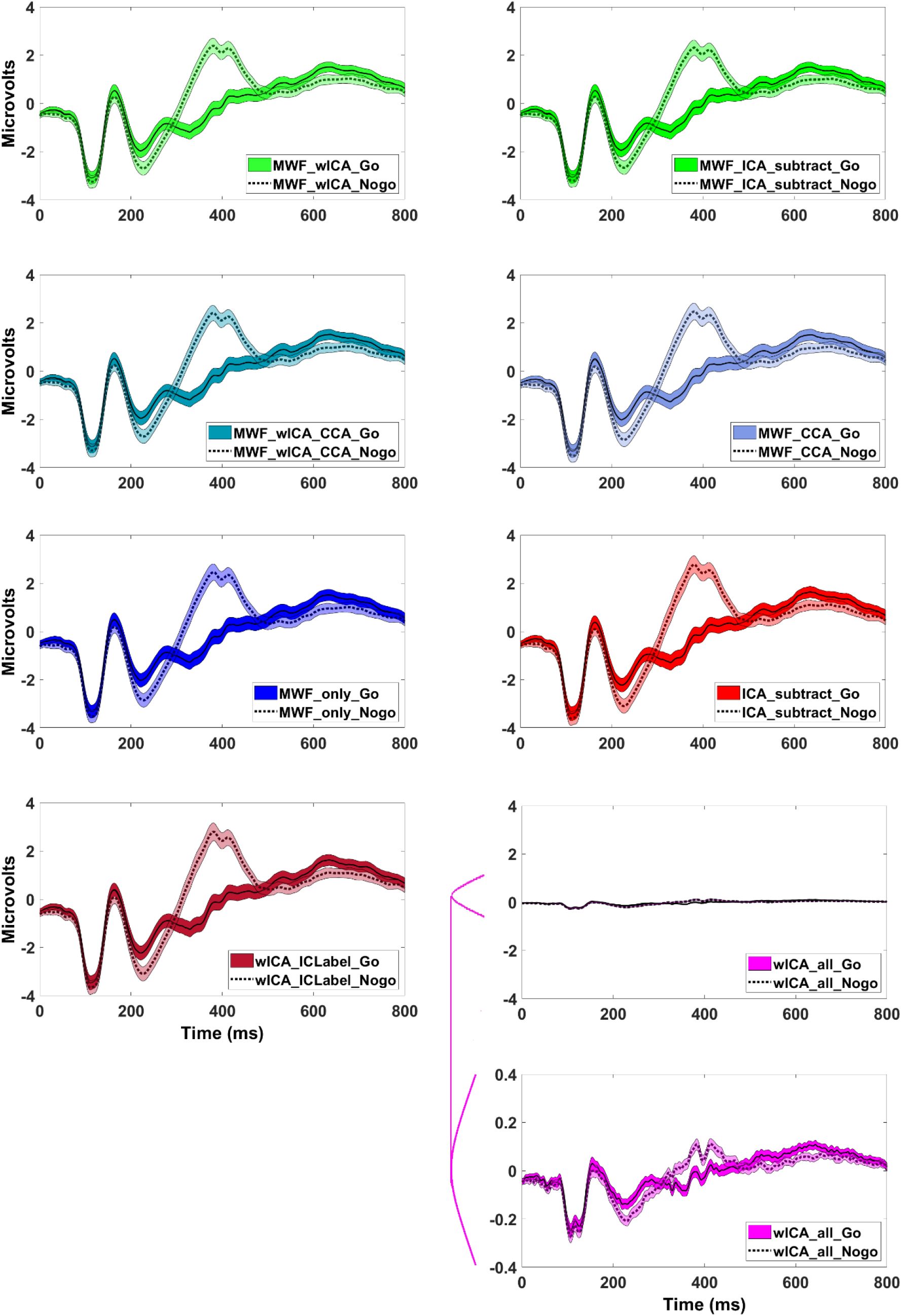
Grand average Go and Nogo related ERP responses from FCz for each pipeline. Error shading reflects 95% confidence intervals. Note the different scale for wICA_all.

**Figure 12.**
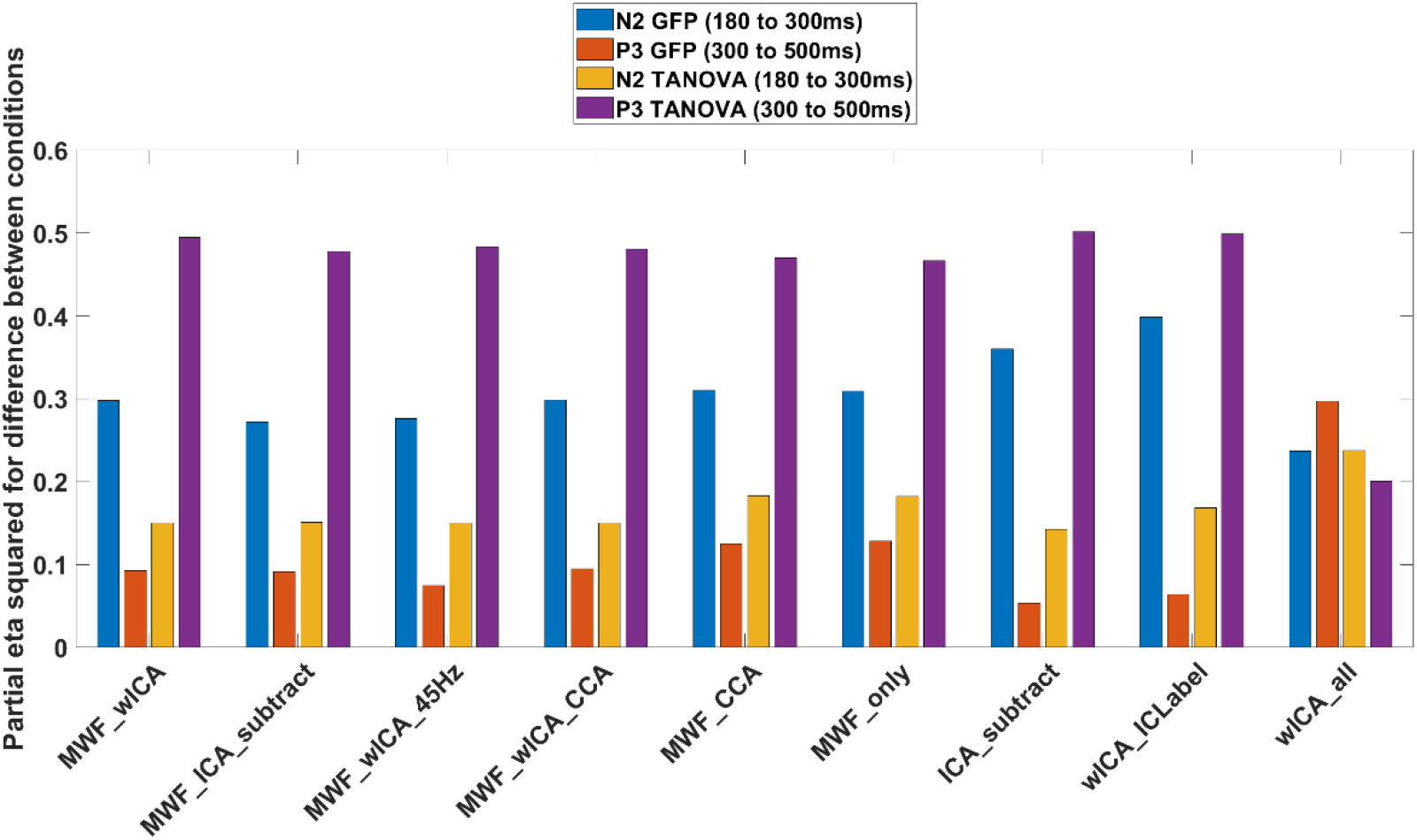
The variance explained (np^2^) by the difference between Go and Nogo trials for the N2 (180 to 300ms) and P3 (300 to 500ms) GFP and TANOVA for each pipeline.

With regards to the P3 GFP most pipelines provided np^2^ values from 0.05 to 0.13 except for wICA_all, which provided a value of 0.30. wICA_all performed the best, followed by MWF_only, MWF_CCA, then MWF_wICA_CCA, MWF_wICA, MWF_ICA_subtract, wICA_ICLabel, and ICA_subtract. While wICA_all seemed to perform the best for the N2 TANOVA and P3 GFP metrics, it produced EEG data with very small amplitudes, suggesting the pipeline is likely to have cleaned neural activity as well as artifacts. Lastly, with regards to the P3 TANOVA, most pipelines provided np^2^ values from 0.46 to 0.50, except for wICA_all which provided np^2^ = 0.20. For this metric, ICA_subtract, wICA_ICLabel, MWF_wICA, MWF_ICA_subtract and MWF_wICA_CCA performed better than MWF_CCA and MWF_only, which in turn performed better than wICA_all. Full details for these metrics can be found in the Supplementary Materials, section 4, pages 41-54.

### ERP Amplitude Reliability Metrics

#### Number of Errors Required for Dependable Analysis of the Pe

With regard to the dependability of the Pe ERP data from error related epochs, MWF_wICA and MWF_wICA_CCA appeared to be the best performers, requiring only eight epochs for valid analysis and only excluding 4/76 participants (Figure 13). MWF_ICA_subtract and MWF_only both provided the same level of dependability (requiring only 8 error related epochs). However, these pipelines excluded more participants (as a larger number of epochs were removed by the cleaning approaches then final epoch rejection step for these pipelines). ICA_subtract, MWF_CCA, and wICA_ICLabel showed less dependable Pe amplitudes across all trials and participants, requiring nine error related epochs for valid analysis (and excluding 7-8 participants). wICA_all provided the least dependable data, with 12 epochs required and 18 participants excluded.

**Figure 13.**
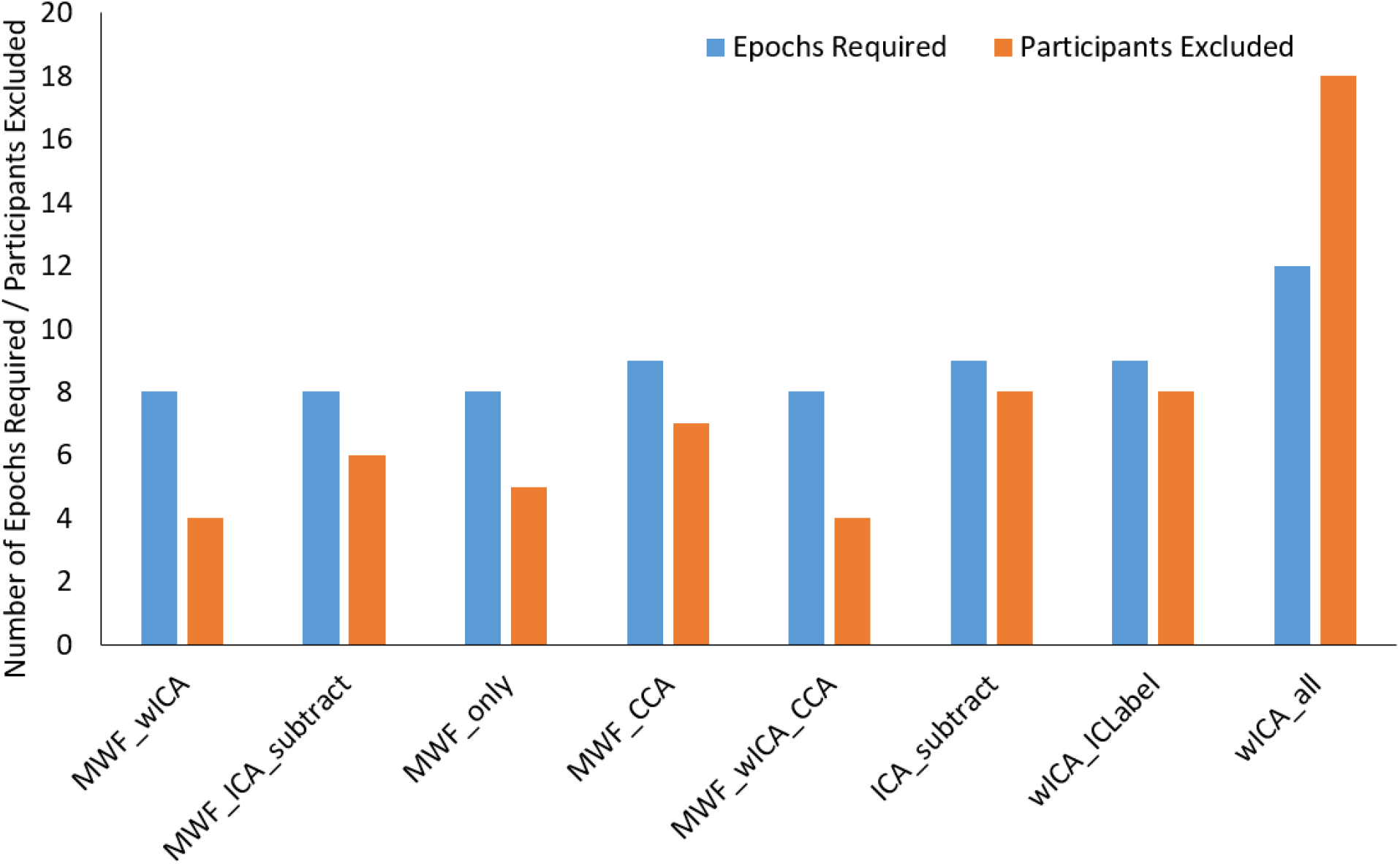
The minimum number of epochs required for a dependability of 0.8 in an analysis of the error related Pe at FCz, and number of participants excluded from error processing analyses based on this dependability threshold for each of the cleaning pipelines.

#### ERP Amplitude and Single Trial Bootstrap Standard Error of the Mean

Lastly, we report here the peak amplitude detection bSME values. Artifacts in the EEG data have a more severe influence on peak amplitude measurements of ERPs than on measurements averaged over the ERP time-period of interest, since the time-period of interest can contain both clean and artifact influenced data, which mitigates the effect of the artifact on the ERP measurement. As such, we measured the bSME using the peak detection method, which is more sensitive to effective cleaning of the EEG data. Since all peak amplitude measures showed positive values (so ratios provide a valid measure for analysis), and since bSME values should be considered in the context of the ERP amplitude (Luck et al., 2021), we analysed the ratio of the bSME to P3 peak amplitude (Figures 14 and 15). wICA_all performed the best, providing very low P3 peak amplitudes relative to the other pipelines (a mean peak amplitude of 1.24μV compared to peaks of 6.6μV to 7.6μV from other pipelines), but bSME values that were proportionally even lower than other pipelines. MWF_ICA_subtract, MWF_wICA, MWF_wICA_CCA performed the next best, with MWF_ICA_subtract performing better than MWF_wICA_CCA, and none of these pipelines differing from MWF_only. MWF_CCA, wICA_ICLabel, ICA_subtract performed the worst. The full details, and analyses of the bSME values independent of ERP amplitudes are reported in the Supplementary Materials (section 4, page 55-70), as well as comparisons of the N1 and N2, but these comparisons show the same pattern to the results (with the exception that the MWF_wICA methods all significantly outperformed wICA_ICLabel, ICA_subtract, MWF_only and MWF_CCA to a larger extent when assessing the N1 and N2 than they did with the P3).

**Figure 14.**
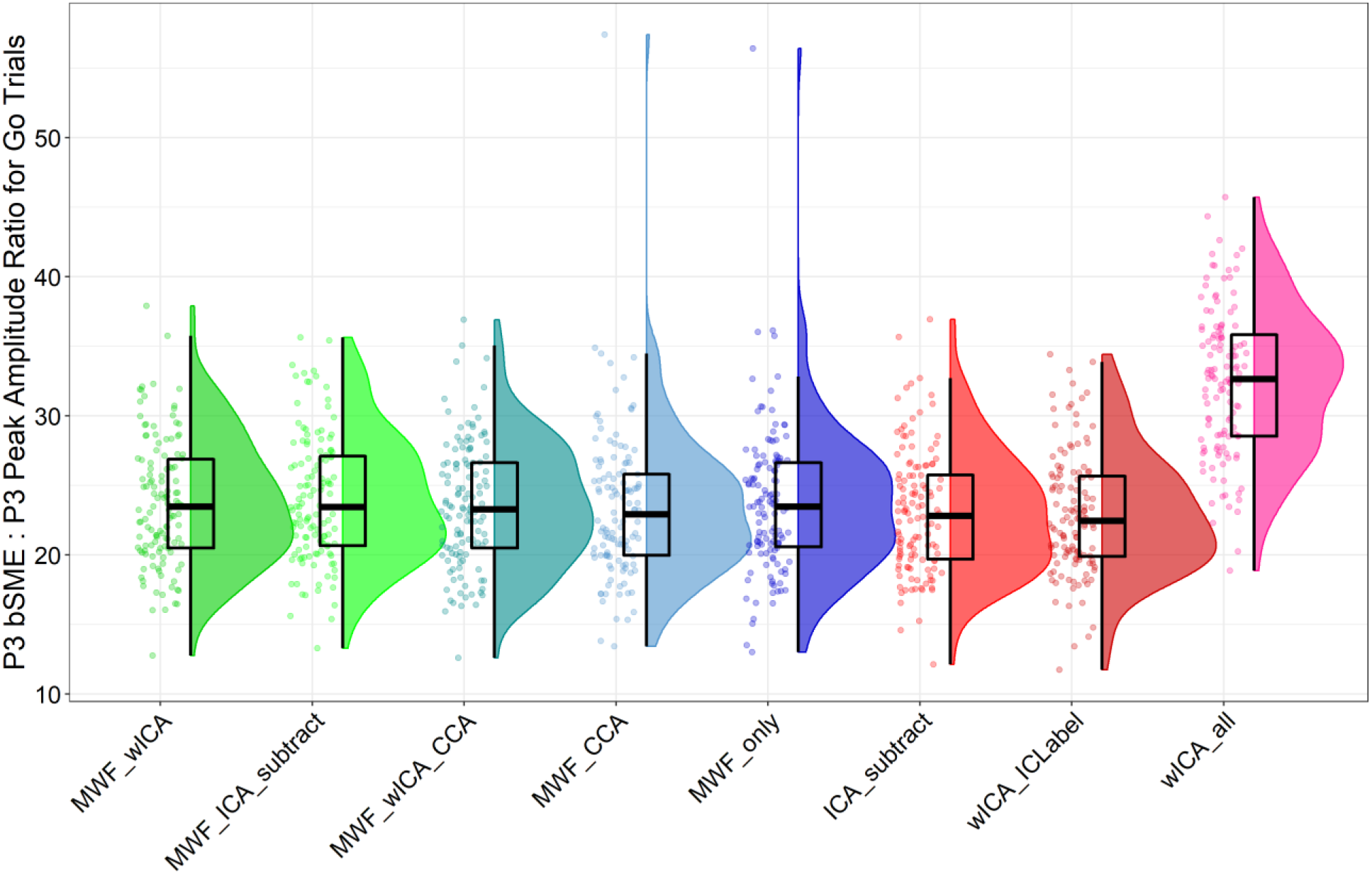
The ratio of peak P3 bSME values to P3 peak amplitude values for peak amplitude detection of the P3 in Go trials across all participants.

**Figure 15.**
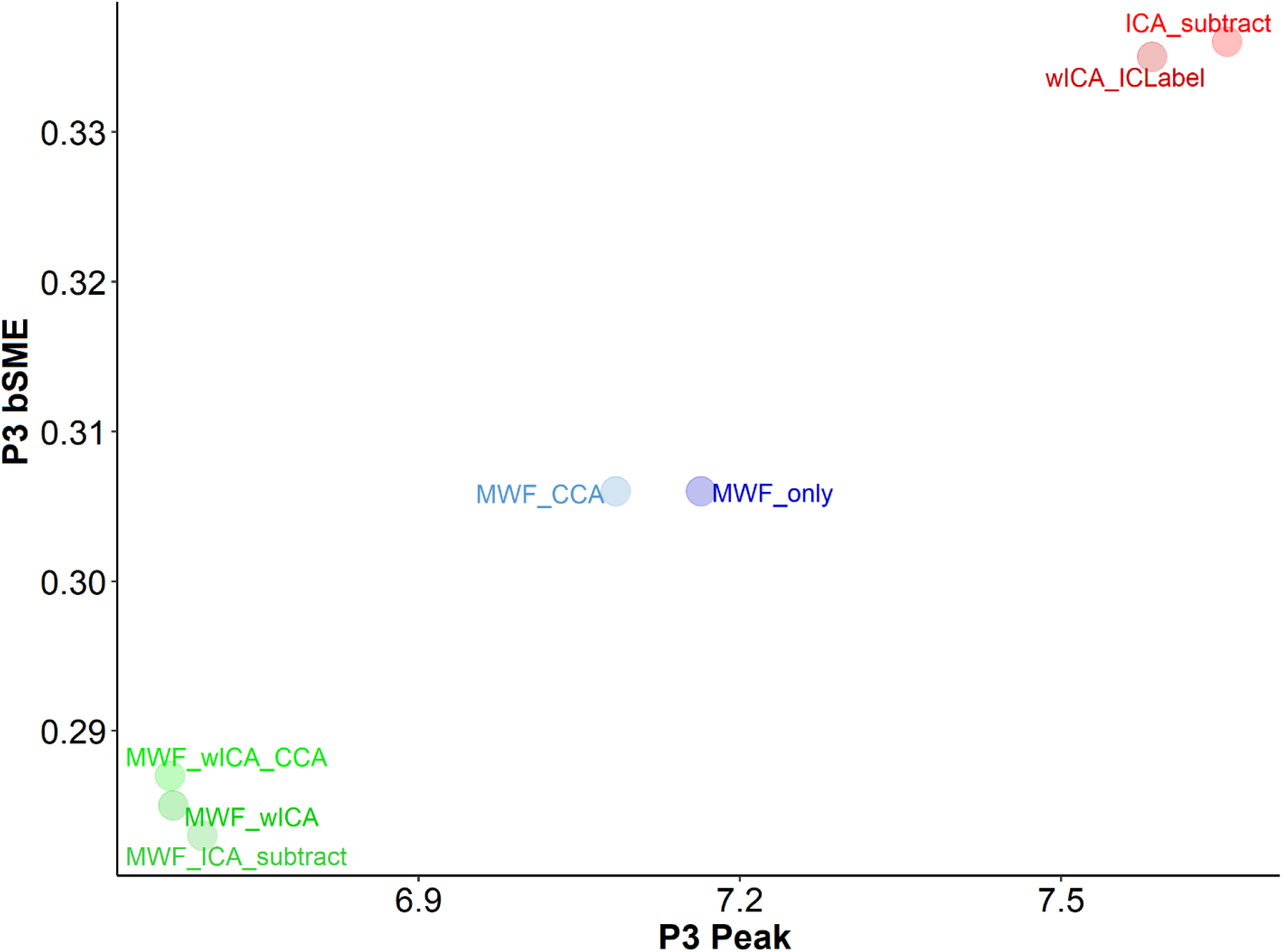
Scatterplot depicting mean bSME values against mean amplitude values for peak amplitude detection of the P3 in Go trials from each cleaning pipeline, excluding wICA_all to provide sufficient resolution to discern differences in the other pipelines (full details can be viewed in Supplementary Materials Figure S55, page 63).

## Discussion

Here we have reported comparisons between several variations of our newly developed RELAX EEG cleaning pipeline and four pipelines that have been presented by previous research. These comparisons were conducted after cleaning data that has been high-pass filtered at 0.25Hz (which enables valid ERP analysis). We have discussed the general cleaning efficacy of RELAX compared to other pipelines in detail in our companion article (Bailey et al., 2022), so here we focus specifically on factors that are relevant to ERP analyses. Similar to the results of our companion paper (which examined data that was high-pass filtered at 1Hz), the comparisons of ERP data indicated that the RELAX MWF_wICA pipeline (with infomax ICA) cleaned artifacts very effectively while successfully preserving the neural signal. Additionally, the RELAX MWF_wICA or wICA_ICLabel methods were either the highest performing pipelines, or nearly equivalent to the highest performing pipeline for most measures of the variance explained by differences in ERP global amplitudes and distributions between Go and Nogo conditions and error and correct responses. The exception to this general finding was that the wICA_all pipeline seemed to produce increased discernment of conditions for the N2 distribution and P3 global amplitude. However, this higher wICA_all performance for detecting differences in the N2 distribution and P3 global amplitude came at the expense of reducing considerable neural signal, and as such reducing the capacity to fully characterise the neural activity. As such, we recommend the use of RELAX MWF_wICA for typical EEG cleaning, or wICA_ICLabel in specific circumstances (discussed below).

Consistent with the results of our companion article, MWF_wICA provided high SER and ARR values concurrently, indicating MWF_wICA showed the best performance at reducing the amplitude of artifact affected periods in the EEG data and preserving neural signal in the clean periods. Importantly, MWF_wICA showed better performance than ICA_subtract in both SER and ARR metrics, and wICA_ICLabel showed better performance than ICA_subtract in SER while also showing very similar performance in ARR. In combination with the other results for ICA_subtract, we suggest this highlights that ICA_subtract is seldom the optimal approach, despite being probably the most commonly used cleaning pipeline in the literature. Similarly, the results showed wICA_all to produce very low SER values, and a reduction by an order of magnitude for the averaged ERP amplitude. The combination of low SER values and very low amplitude ERPs with distorted distributions compared to all other cleaning pipelines suggests that wICA_all overcleaned the data, removing elements of the neural signal from clean periods, and as such we do not recommend its use. MWF_only or wICA_ICLabel preserved more of the signal from the clean EEG periods than other pipelines, indicated by higher SER values, but these methods were less effective at reducing artifacts (indicated by lower ARR values, and higher blink and muscle measures remaining after cleaning). The metrics assessing remaining blink and muscle after cleaning also showed MWF_wICA to be better at cleaning these artifacts than ICA_subtract. The use of MWF_wICA also resulted in fewer epochs being deleted (and as such, more were available for analysis) and produced the most dependable and reliable ERP data (except for wICA_all for the stimulus locked ERPs, which we suspect achieved very reliable data by severely reducing amplitudes). The inclusion of more high-quality epochs for analysis has been suggested by simulations to improve study power even more than increasing sample size (Kolossa & Kopp, 2018). This may be particularly important for error processing research, where often few epochs are available for analysis, and as such we recommend MWF_wICA for studies of error processing and similar “low trial number” applications.

Despite the positive indications noted in the previous paragraph, the primary reason to effectively clean EEG data is to enhance the detection of true experimental outcomes (Clayson, Baldwin, et al., 2021). Based on our findings, we cannot recommend a single best pipeline for all applications. Fortunately, apart from wICA_all, all cleaning pipelines produced similar amounts of variance explained and similar patterns for the ERPs we examined. As such, so long as artifacts are reasonably well addressed, variation in pipeline use may not lead to largely different outcomes in different ERP studies. Having said that, we note that we selected well validated comparisons that provided large differences between ERPs. Studies examining more subtle effect sizes will be more vulnerable to differences in cleaning efficacy, for example, differences between healthy and clinical populations or treatment effects (Rogasch et al., 2020). Effect sizes have also been shown to vary depending on the cleaning pipeline used, suggesting optimal pipeline selection is important (Clayson, Baldwin, et al., 2021; Robbins et al., 2020; Rogasch et al., 2020). Given the benefits of MWF_wICA for cleaning efficacy and its consistently good performance in the detection of differences between experimental conditions, we recommend the use of the RELAX pipeline with MWF_wICA as a default effective cleaning pipeline for ERP research when no rationale exists to prefer another variation of RELAX. With regards to specific experimental manipulations, the amount of variance explained by the experimental manipulations indicated that the MWF_wICA pipeline did not significantly differ from the most effective pipelines when examining the P3 distribution (and MWF_wICA also performed highly for most other measures). However, we note that in specific circumstances, the RELAX pipeline with the wICA_ICLabel setting might be preferred. For example, wICA_ICLabel led to higher values for some of the metrics of variance explained between experimental conditions (for differences in the distribution of activity of the ERN and Pe, and differences in the N2 global amplitude). As such, wICA_ICLabel might be preferred for these ERPs in certain cases. Similarly, MWF_only might be preferred for analyses of the Pe or P3 global response amplitude, or N2 TANOVA.

However, because the wICA_ICLabel and MWF_only pipelines did not clean blinks or muscle as effectively, we recommend they only be used where enough trials are collected (or enough participants) that lower reliability could be acceptable, as research has suggested data quality is more important for experimental power than data quantity (Kolossa & Kopp, 2018). As such, wICA_ICLabel might be recommended for studies examining the distribution of the ERN, Pe, and the N2 global amplitude, and MWF_only for the Pe or P3 global amplitude or differences in the N2 distribution, but only where researchers apply other methods that ensure noisy data will not confound results. For example, robust statistics can protect against noisier data, so studies that use robust statistics and include all single trials in the analyses may prefer to use wICA_ICLabel (Alday & van Paridon, 2021).

In addition to the performance of our comparison pipelines, there were several interesting details revealed by the analyses we reported in our Supplementary Materials (section 5, pages 73-87). First, despite the fact that AMICA has been proposed as a more effective ICA method for decomposing EEG data (Palmer et al., 2012), it was not clearly superior to infomax or fastICA in any metric, and actually displayed poorer performance across several metrics. Although previous research has suggested that AMICA separates components more effectively and provides more dipolar components (Delorme et al., 2012; Leutheuser et al., 2013), as far as we are aware, this is the first time AMICA, infomax and fastICA have been compared across as many ‘outcome oriented’ metrics as in the current study. Given the relatively long processing time required by AMICA, our results suggest that cudaICA (Raimondo et al., 2012) or fastICA (Hyvarinen, 1999) might be preferred options. However, we note that research that examines ICA components as the outcome measure (rather than simply using ICA for artifact cleaning) may still benefit from AMICA’s increased detection of dipolar components (Palmer et al., 2012). Also, it is worth noting that we performed fastICA using the ‘defl’ setting so that non-convergence issues were prevented. However, recent research has suggested that fastICA with the ‘symm’ setting performs better than fastICA with the ‘defl’ setting, and better than infomax (Barban et al., 2021). We tested this and have provided these results in our Supplementary Materials (section 5, pages 73-79). In brief, our results suggested that infomax was the best performer (but by very little) at cleaning blinks, and the ‘defl’ setting outperformed the ‘symm’ setting for muscle activity (again, by very little). These results also suggested infomax was better than ‘defl’ at detecting differences between Go and Nogo trials in the N2 distribution, and ‘symm’ almost outperformed ‘defl’ in this metric also (but also with very small differences, and no differences for any other Go-Nogo ERP metrics). Because there was such little difference between these ICA algorithms, and because fastICA ‘symm’ performs the most quickly (unless cudaICA can be installed), our final version of RELAX provides fastICA with the ‘symm’ setting as the default. We have set RELAX to re-run the ICA decomposition (up to three times) if a non-convergence issue is detected, as these are more common using the ‘symm’ setting (and adversely affect ICA decomposition and cleaning). After the third attempt, the algorithm then switches to the ‘defl’ setting if non-convergence still occurs. Having stated this, note that if cudaICA can be implemented, this is the fastest and best performer. We note cudaICA can be difficult to install. If users are interested, helpful instructions have been provided by Miyakoshi (2018). Finally, while some research has suggested that ICA performance can be enhanced by filtering out data above 45Hz prior to ICA (Zakeri, 2017), we did not find evidence to suggest this improved overall performance.

It is also worth mentioning that ICA decompositions are generally more effective when computed using data that is high-pass filtered at 1Hz (Winkler et al., 2015). This fact is likely to explain the higher fBAR values after cleaning in our ERP dataset (which were high-pass filtered at 0.25Hz) compared to our 1Hz filtered dataset. To address this issue, researchers have often performed ICA decompositions on 1Hz filtered data, then applied the artifactual component rejections to their data that has a lower (ERP compatible) high-pass filter (Debnath et al., 2020; Rodrigues et al., 2021). However, as far as we are aware, until now no research has tested whether this approach improves cleaning performance and the detection of experimental manipulations of interest compared to simply performing the ICA on the ERP compatible high-pass filtered data (data filtered at 0.25Hz). We tested this for both the MWF_ICA_subtract and ICA_subtract pipelines (reported in our Supplementary Materials, section 5, page 83-87). Our results indicated that the approach where “1Hz filtering is performed before ICA decomposition, then the ICA decomposition is copied to the 0.25Hz filtered data” performed worse at both reducing blinks and detecting the experimental manipulation between Go and Nogo trials in N2 GFP values when compared to simply performing the ICA decomposition on the 0.25Hz data. As an informal test to determine why this might be the case, we low-pass filtered our raw data at 1Hz, and inspected this data during blink periods. Voltages became more positive in frontal electrodes during blink periods, indicating blinks contain a contribution from frequencies <1Hz. We suspect the blink cleaning performance is reduced when applying a 1Hz ICA decomposition to the 0.25Hz data because the ICA cleaning does not address this low frequency component of the blink artifact. As such, we recommend simply high-pass filtering data at 0.25Hz, then applying ICA decomposition only to this ERP compatible filtered data (rather than copying 1Hz filtered ICA decompositions to the 0.25Hz ERP compatible filtered data). However, we note that in addition to the reduced ICA decomposition performance when <1Hz data is included, ICLabel was trained on 1Hz high-pass filtered data, so may be mis-identifying some blink influenced components when <1Hz data is included. To address these issues, we recommend that future research explores both ICA methods that are more robust to the influence of <1Hz data, and that an automatic artifactual component identifying algorithm be trained on data after filtering that is appropriate for ERP research.

Given that researchers may prefer to use a different version of RELAX depending on their data, RELAX has been designed to be modular so that different cleaning options can be selected. These options should be justified by reference to empirical results, ideally evidence indicating maximal explained variance from a pipeline using similar experimental design. Best practice would be pre-registration of the pipeline settings to provide confidence that researchers have not repeated their analysis searching for positive results. However, we also note that it may also be useful for the development of optimal analysis methods to report analyses using multiple settings (while indicating which was the primary analysis setting that was a priori selected). This would accelerate knowledge of optimal cleaning methods within the field. Reporting consistency across the results produced by two completely different cleaning pipelines (for example, both MWF_only and wICA_ICLabel) would also provide confidence in the robustness of results to variations in cleaning method, and so demonstrate that a particular result was not simply the product of specific cleaning choices. Lastly, RELAX provides a report of cleaning efficacy metrics, which can be reported to enable reviewers / readers to assess whether remaining artifacts might have influenced results (these could be reported in supplementary materials for the sake of brevity).

### Limitations and Future Directions

We have discussed the general limitations for our conclusions about RELAX and the comparison pipelines in our companion article (Bailey et al., 2022). Here, we focus on the limitations relevant to ERP research. First, the metrics we used to examine the variance explained by the experimental manipulations were based on ERP averages within conditions/participants. While this is a common analysis method, researchers are increasingly using single trial analysis methods to provide improved ability to disentangle relationships between brain activity and perception/behaviour. It may be that averaged ERP analyses masked potential cleaning inefficiencies in some of our tested pipelines, and we think it is likely that including all epochs in the analysis would provide better information about cleaning efficacy. Future research using single trial analyses (for example, using LIMO or the decision decoding toolbox) may provide a stronger ability to differentiate the cleaning pipelines in their ability to produce data that maximally discerns relationships between brain activity and perception/behaviour (Bode et al., 2019; Pernet et al., 2011; Stewart et al., 2014).

Second, it seems that regardless of the cleaning pipeline used, data that is high-pass filtered at 0.25Hz is not as effectively cleaned as data that is high-pass filtered at 1Hz. In particular, it is important to note that approximately 5/127 files contained data that were obviously strongly affected by blinks, with high BAR values even after cleaning regardless of the cleaning approach. Further, all pipelines showed median fBAR values of 1.1 to 1.2 in the 0.25Hz high-pass filtered data, suggesting no pipeline managed to completely clean blink artifacts when filters appropriate to ERP data analysis were used. Note that it is possible that brain regions are differentially activated during a blink compared to non-blink periods, providing different amplitudes of neural activity, which would explain these higher BAR values without the values indicating a remaining contribution of the blink artifact after cleaning. If this is the case, then optimal BAR values might in fact not be exactly 1. However, we are not aware of any evidence that suggests this to be true, and our informal test demonstrated that blinks contain activity from <1Hz frequencies (Supplementary Materials, section 7, page 92). As such, we recommend that studies should implement methods to check for remaining blink affected data after cleaning (BAR is a useful method for this and is supplied as an outcome metric in RELAX) (Robbins et al., 2020). If this check indicates that blinks have still affected the data, and blinks are either time-locked to stimulus presentation or larger/more frequent in one condition/group than a comparison condition/group then we recommend either the use of single trial analysis methods with robust statistics (to prevent the influence of blink outliers), or the deletion of blink affected epochs. However, note that if blinks are commonly time-locked to stimuli (for example, in tasks with bright flashing visual stimuli), even the use of single trial analysis with robust statistics may not provide an adequate control. In future research, we plan to explore cleaning methods that can address these residual artifacts in 0.25Hz high-pass filtered data, and to develop an automated test of whether remaining blink activity might affect statistical comparisons, both of which we will provide with the RELAX toolbox when completed.

Third, research has indicated that for some combinations of EEG systems and temperatures, low frequency drift adversely affects recordings. In these situations, the probability of a significant result was improved by using high-pass filtering cut-offs at 0.5Hz rather than 0.1Hz (Kappenman & Luck, 2010). This low frequency drift is a factor that visual inspection of our raw data suggests has influenced our data. As such, we used fourth-order acausal Butterworth high-pass filtering at 0.25Hz for our ERP data to address the slow voltage drift that affected some of our data (which visual inspection suggested was not to be resolved by 0.1Hz high-pass filtering). We made this choice based on several factors. Firstly, informal testing that showed fourth-order Butterworth high-pass filtering at 0.25Hz led to the best performance across pipelines at eliminating voltage drift, whereas 0.1Hz filtering did not fully eliminate the drift. We also determined that Butterworth filtering worked better with MWF cleaning than EEGLAB’s ‘pop_eegfiltnew’. This might be due to the smooth roll-off and minimisation of ripple filtering artifacts provided by the Butterworth filter, or alternatively, may be related to the extent of the temporal smearing produced by the filters (temporal smearing creates dependence between timepoints, which increases the risk that the MWF cleaning will encounter eigenvectors that are rank deficient at longer delay periods). Our use of Butterworth filtering with these settings is also in agreement with another recently developed ERP cleaning pipeline (Debnath et al., 2020).

Secondly, we selected 0.25Hz as the high-pass filter, as previous research has shown that a Butterworth filter limit of >0.3Hz negatively affected ERP analyses, while 0.2Hz did not (Rousselet, 2012). However, while no research we are aware of has yet demonstrated that 0.25Hz high-pass Butterworth filtering is inferior to 0.1Hz filtering, optimal high-pass filtering is still debated, and experts on the matter have suggested that 0.1Hz is optimal for long latency ERPs (Maess et al., 2016; Tanner et al., 2016). As such, we suggest further research is required, and we have made it simple to adjust high-pass filtering settings between 0.25Hz and 0.1Hz in the RELAX pipeline. We suggest that 0.25Hz is a safe ‘default’, and if 0.1Hz is used, a subset of the data should be inspected for remaining drift artifacts after filtering before statistical analyses are implemented. Finally, to address any potential remaining drift after data cleaning, we recommend the use of regression based baseline correction methods for ERP analyses (rather than subtraction methods) (Alday, 2019), an implementation of which is made available in the epoch selection and rejection script on the RELAX GitHub repository.

Finally, while the MWF_wICA or wICA_ICLabel pipelines showed amongst the most variance explained by the experimental manipulation across most datasets and analysis methods, there is no way to be sure of the ground truth of these comparisons. As such, it is possible that the higher levels of explained variance are due to artifacts rather than actual differences in neural activity. However, there are reasons to believe this is not the explanation for our results. These are explored in detail in our companion paper (Bailey et al., 2022), but in brief, if artifacts explained the result, then artifacts would have to remain after cleaning in only one of the two conditions compared (across two tasks, since we analysed a dataset from the Go-Nogo task and in our companion article the Sternberg task related data). The artifacts would also have to be time-locked to the stimulus onset in that condition and provide differences between conditions that would reflect artifact topographies, factors which were not seen in visual inspections of the data (Supplementary Figures S23-24, S29-30, S36-37, and S42-43). Future research using simulated data or hybrid data with simulated artifacts added to real data that has already been cleaned might help address the unknown ground truth issue, but simulated data comes with its own complications. For example, it is difficult to determine whether the simulated artifacts are analogous enough to real artifacts to provide an effective test (Kumaravel et al., 2022; Mumtaz et al., 2021; Rošťáková & Rosipal, 2021). We chose not to use simulations for this reason, and instead preferred to comprehensively assess real data using previously validated cleaning efficacy metrics and measures of the amount of variance explained by well-established experimental manipulations.

## Conclusion

To conclude, we have reported the results of comparisons of artifact cleaning performance with EEG signal preservation across 11 automated EEG cleaning pipelines. Our results demonstrate that our RELAX pipeline with the default MWF_wICA setting for cleaning EEG provides the following benefits: 1) superior cleaning of artifacts; 2) RELAX is fully automated and implemented simply via an EEGLAB graphical user interface, saving considerable researcher time, and preventing subjectivity from influencing results; and 3) RELAX contains embedded cleaning quality metrics, which can also be easily reported in publications to allow reviewers and readers to see how effectively the data was cleaned, and 4) RELAX MWF_wICA importantly provides the largest number of artifact free epochs for analysis, and amongst the most reliable/dependable ERP measures, minimizing potential biases based on epoch exclusion and increasing statistical power, features that are particularly beneficial for error processing research. However, the RELAX wICA_ICLabel and MWF_only settings may provide superior detection of between condition ERP effects for some studies. Unlike many automated cleaning pipelines, RELAX can be implemented to analyse task-related ERP data, although we note that this does reduce data cleaning efficacy compared to experimental designs that can high-pass filter data at 1Hz. Finally, given the stressful nature of academic research, we hope that the name of our pipeline can also be taken as a suggestion, and that some of the time saved by using RELAX can be spent on researcher well-being, instead of only increasing productivity. RELAX is available for download from GitHub (https://github.com/NeilwBailey/RELAX/releases) and is designed to run within EEGLAB (implemented in MATLAB).

**Recommendations for using RELAX when examining ERPs**

1. High-pass filter data at 0.25Hz
2. Use RELAX_MWF_wICA as the default cleaning pipeline (with fastICA on the symm setting, or cudaICA if it can be installed)
3. Use RELAX_wICA_ICLabel if analysing the Pe distribution or N2 amplitude, but only if data are relatively clean, if using analysis methods that prevent remaining artifacts confounding conclusions (for example single trial analyses with robust statistics) and if a large number of trials will be available for analysis.
4. Use RELAX_MWF_only if analysing the Pe or P3 amplitude, or N2 distribution, but only if data are relatively clean, if using analysis methods that prevent remaining artifacts confounding conclusions (for example single trial analyses with robust statistics) and if a large number of trials will be available for analysis.

See our companion article for recommendations related to other types of analyses (Bailey et al., 2022).

