## Supplementary Materials for "Introducing RELAX (the Reduction of Electroencephalographic Artifacts): A fully automated pre-processing pipeline for cleaning EEG data – Part 2: Application to Event-Related Potentials"

### LIST OF ABBREVIATIONS

| **Acronym** | **Description** |
|---|---|
| allBAR | All electrodes Blink Amplitude Ratios |
| ANOVA | Analysis of Variance |
| ARR | Artifact to Residue Ratio |
| ASR | Artifact Subspace Reconstruction |
| BAR | Blink Amplitude Ratios |
| CCA | Canonical Correlation Analysis |
| EEG | Electroencephalography |
| EOG | Electrooculogram |
| ERN | Event-Related Negativity |
| ERP | Event Related Potentials |
| fBAR | Frontal Electrodes Blink Amplitude Ratios |
| FDR | False Discovery Rate |
| GFP | Global Field Potential |
| HAPPE | Harvard Automated Processing Pipeline for Electroencephalography |
| ICA | Independent Component Analysis |
| MWF | Multiple Wiener Filtering |
| RAGU | Randomised Graphical User Interface |
| RELAX | Reduction of Electroencephalographic Artifacts |
| SER | Signal to Error Ratio |
| wICA | Wavelet Enhanced Independent Component Analysis |
| TANOVA | Topographic Analysis of Variance |

### Supplementary Materials

### SECTION ONE

#### Supplementary Background Points

The multi-channel Weiner filter (MWF) approach performs well at reducing temporary artifacts that can be identified in limited time windows, such as muscle activity, eye movement / blinks, and electrode drift. After cleaning with the MWF, the data primarily contains only smaller artifacts and most of the brain activity is preserved, allowing for optimal application of the ICA algorithm. The MWF method is also not adversely affected by frequencies <1Hz (unlike ICA methods), requiring only that data be zero mean overall [3], making the RELAX MWF_wICA pipeline compatible with event related potential (ERP) analyses. The use of wavelet ICA (wICA) instead of the typical approach of subtracting independent components means that reducing artifact components with wICA has a reduced chance of removing probable brain activity as well as the artifact. While ICA is commonly used to address blink activity, it is worth noting that non-stationary electrooculogram (EOG) artifacts have been suggested to be not fully addressed by ICA, which does not take into account temporal information in its modelling [1]. In contrast, wICA has the additional advantage of not requiring artifacts to be stationary [2].

Each step in our cleaning pipeline allows for the selection of multiple parameters, which can affect cleaning outcomes. During the design stage of our pipeline, we varied the selection of each of the parameters across the spectrum of potential values via considerable informal testing, to narrow down to the optimal outcomes in terms of metrics showing artifact reduction, the signal of identified non-artifact periods being maximally retained, and the variance explained by the experimental design being optimized across multiple large datasets and experimental designs. As such, we recommend use of the default parameters, but if future research demonstrates other parameters are superior, it is simple to adjust the selected parameters.

### SECTION TWO

#### Additional Comparison Pipeline Description

MWF_CCA and MWF_wICA_CCA used the sequential MWF cleaned data, but instead of applying wICA to clean the muscle activity as per the RELAX methods, they used the extended canonical correlation analysis (CCA) to further clean any remaining muscle artifacts [4]. CCA separates the EEG data into components that are not correlated with each other but are maximally autocorrelated at a lag of one datapoint. Muscle activity is characterised by a similar pattern to white noise, with a low autocorrelation (in contrast to neural activity, which shows voltage fluctuations at a slower rate with higher autocorrelation). As such, CCA is an effective method for identifying and removing muscle activity from EEG and has been suggested to be superior to ICA methods [5, 6]. Recently CCA has been improved through the use of a log-power log-frequency slope thresholds to detect muscle activity. The optimal threshold was identified by the comparison of paralysed and non-paralysed scalp EEG recordings to detect probable muscle contaminated components for removal [4]. This approach removes components with a one timepoint-lag autocorrelation of less than 0.19 and log-power log-frequency slopes of more than -0.48 [4]. We used this extended CCA after the initial MWF cleaning, and refer to this method throughout as MWF_CCA. Similarly, MWF_wICA_CCA was identical to RELAX, except that muscle components were not cleaned in the wICA cleaning step. Instead, CCA was implemented after the wICA step in order to address any remaining muscle components. This method was referred to throughout as MWF_wICA_CCA.

We also tested four pipelines that have been presented by previous research. ICA_subtract is perhaps one of the most commonly implemented: simply rejecting outlying data first (as per the approach used in the initial steps of our RELAX pipeline), computing ICA, and subtracting the components identified as artifacts, then reconstructing the electrode space data [7]. We implemented this using ICLabel to identify artifactual components [7]. wICA_all was identical to ICA_subtract, but instead of simple ICA subtraction on artifact components, it applied the wICA approach to all components (as per [2]). We refer to this pipeline throughout as wICA_all. A similar approach was used in wICA_ICLabel, but instead of applying wICA to all components, wICA was applied only to components identified as artifacts by ICLabel. Although a similar approach of applying wICA to only artifact components has been previously implemented [8, 9], as far as we are aware, this is the first time it has been tested by selecting components with ICLabel. MWF_only implemented only a sequential MWF cleaning identical to the MWF cleaning steps in our RELAX pipeline but did not apply any additional cleaning after the MWF stage (no wICA, unlike the RELAX methods). This is similar to the approach used by [3], with the extension of applying their suggested sequential MWF cleaning to clean multiple different categories of artifacts. We refer to this pipeline throughout as MWF_only.

We also tested a few modifications of our RELAX pipeline. These included using different ICA algorithms, namely, infomax (implemented with cudaICA, referred to as MWF_wICA_infomax) [10], fastica (referred to as MWF_wICA_fastICA) [11], or AMICA (referred to as MWF_wICA_AMICA) [12]. Another modified version of RELAX involved subtracting artifactual ICA components instead of using wICA (referred to as

MWF_ICA_subtract), and low pass filtering at 45Hz prior to implementation of the ICA algorithm (which has been suggested to improve ICA decomposition [13], referred to as MWF_wICA_45Hz).

Lastly, we tested a limited number of our cleaning metrics on a few different parameters that could be set within the RELAX pipeline. Firstly, we tested whether fastICA using the deflation method, or symmetrical method was superior (compared against our typical cudaICA method), when using the wICA_ICLabel setting. Secondly, it has been demonstrated that ICA decompositions are adversely affected by high-pass filtering <1Hz. A typical method to address this for ERP research has been to compute the ICA on 1Hz filtered data, then to copy the ICA decompositions to the data that were filtered appropriate for ERP analysis (<1Hz). As far as we are aware, the merits of doing this have never been empirically compared to simply computing the ICA on the data after it has been filtered <1Hz. As such, we tested two approaches using ICA_subtract as our test pipeline: 1) computing the ICA on 1Hz filtered data, ascertaining which components were artifacts within this 1Hz filtered data using ICLabel, then copying the ICA decomposition to the 0.25Hz data and rejecting the artifacts, and; 2) we tested computing the ICA on 1Hz filtered data, then copying the ICA decompositions to the 0.25Hz filtered data, before using ICLabel to detect the artifactual components and remove them from this data. We also tested approach 1 against simply computing ICA on the 0.25Hz filtered data using MWF_ICA_subtract. These parameter explorations are reported in Section 5 of the supplementary materials.

### SECTION THREE

To examine the effectiveness of the RELAX pipeline in cleaning EEG data for ERP analyses, the variants of the RELAX pipeline and comparison pipelines were compared using a number of different cleaning quality metrics which provide a comprehensive evaluation of various aspects of data integrity and reliability. The pipelines were also examined using metrics which examined the variance in ERP activity explained by comparisons between different cognitive trial types (using trial types with robust evidence for their differences from the previous literature) and the reliability of typically analysed ERP metrics. A detailed explanation of each metric is provided below.

#### EEG Data Cleaning Performance Metrics

The pipelines were compared across six different cleaning quality metrics to provide a comprehensive evaluation of cleaning efficacy, which includes assessment of how each pipeline cleaned the full range of potential artifacts whilst still preserving the neural signal. All of the metrics we used have been used in previous research. These metrics included a measure of the amount of signal left unaffected after the cleaning (Signal to Error Ratio - SER), and the extent to which all artifacts identified by our MWF template were reduced (Amplitude to Residue Ratio – ARR), for which higher values indicate good performance [3, 14, 15]. Note that the SER and ARR values should be considered together. Low SER values and high ARR values are likely to indicate effective artifact removal but also removal of data during the clean signal periods, and low ARR values and high SER values are likely to indicate ineffective artifact removal. A cleaning approach that obtained very high SER values by very effectively removing artifact, but also removing large amounts of the signal would not be helpful, and similarly, very high SER values that were produced by not cleaning artifacts at all would also not be helpful. As such, higher values for both the SER and ARR concurrently indicated better performance.

The metrics also included the ratio of blink amplitudes compared to the amplitudes of surrounding non-blink periods after cleaning, both averaged across frontal electrodes affected by blinks (fBAR) and across all electrodes (allBAR). For fBAR and allBAR, values of 1 reflect optimal performance, values <1 reflect overcleaning, and values >1 reflect under cleaning [16]. Next, the number of epochs showing log-power log-frequency slopes indicative of muscle activity, and the amount by which these slopes exceeded the muscle threshold were assessed, where higher values reflect poorer performance [17]. We assessed the percentage of overall variance explained by brain activity after cleaning (measured by ICLabel), where values closer to 100% reflect good performance. We assessed the proportion of epochs that were rejected through the cleaning process (against the total epochs in the raw data), where lower values reflect better performance.

##### *The Signal-to-Error-Ratio*

The SER was calculated from segments of the data marked as free of artifacts by the automatic artifact detection approaches implemented in the RELAX pipeline. The SER is calculated first on each electrode (*i*) by obtaining the expected value operator (which is analogous to the weighted average, where more probable values given stronger weights when computing the average) of the square of the signal in the “raw” (not yet cleaned) data across all periods marked as clean ($y_i$), then dividing this value by the expected value operator of the square of the signal that was removed by the cleaning pipeline across the periods marked as clean, then multiplying this value by the log10 of 10 ($\hat{d}_i$) (see Equation S1) [3, 14, 15]. Note that the “raw” data we used in the calculation of the SER was obtained after data had been filtered and extreme outlying electrodes and periods had been rejected (and before any of the MWF cleaning steps were applied). This was done because all filtering and application of extreme outlying electrode and period rejections were the same across all pipelines, and were not of interest to this study. In order to obtain a single measure for each cleaned dataset, the SER from each electrode is combined by weighted averaging over all electrodes (Equation S2) [3, 14, 15], with the weighting performed by the proportion of artifact power an electrode produces relative to the artifact power from all other electrodes ($p_i$) (estimated by subtracting the power in the clean segments from the power in the artifact segments, Equation S3) [3, 14, 15]. This has the effect that the electrodes that contained the most artifact contribute the most to the final SER value for that dataset. This approach ensured SER values appropriately reflect the contribution of noisier electrodes and makes the SER robust against high SER values being produced by mostly clean data with a single electrode which is very noisy in artifact periods, and distorted by the cleaning pipeline in the clean periods.

Equation S1:

$$SER_i = 10log_{10} \frac{E\{(y_i)^2\}Ho}{E\{(\hat{d}_i)^2\}} \ (for\ clean\ segments)$$

Equation S2:

$$SER = \sum_{i=1}^{M} \ p_i \ SER_i$$

Equation S3:

$$p_i = \frac{E\{(y_i)^2\}\ (artifact\ segments) - \ E\{(y_i)^2\}\ (clean\ segments)}{\sum_{i=1}^{M} \quad (E\{(y_i)^2\}\ (artifact\ segments) - \ E\{(y_i)^2\}\ (clean\ segments))}$$

The EEG periods that are marked as clean by the automated MWF template construction approach implemented by RELAX do not include blinks, muscle activity, horizontal eye movements and drift (nor do they include extreme artifacts, which were marked as NaNs in the clean/artifact template). As such these “clean” segments should be minimally modified by the cleaning pipelines. Because of this, high SER values are expected if cleaning has left the non-artifact periods undistorted, so high values indicate good performance [3, 14, 15].

***The-Artifact-to-Residue-Ratio***

The ARR was calculated from the periods of the data marked as artifact by the automatic artifact detection approaches implemented in the RELAX pipeline. As with the SER, the calculation of this measure was first performed on individual electrodes by obtaining the expected value operator of the square of the removed artifact ($d_i$), divided by the expected value of the square of the total signal from the artifact periods ($\hat{y}_i$) from the “raw” (not cleaned) data ($y_i$) (when ARR is calculated on real data where the true artifact signal is not known), then multiplying this total by the log10 of 10 (Equation S4) [3, 14, 15]. To obtain a single value for each dataset, the individual electrode values were then combined via weighting in the same manner as the SER (weighting via $p_i$). As such, the ARR provides large values when more artifact is removed relative to the “raw” data (as the denominator of the equation: the “raw” data minus the artifact: $[y_i - \hat{d}_i]$ will be as small as possible). The ARR is valid when artifacts are high in amplitude relative to the clean data (as per the blink, muscle, horizontal eye movement and voltage drift artifacts selected by the MWF artifact template in the current study, which are mostly based on outlying amplitudes or artifacts that are typically large in amplitude). Note that the “raw” data used in the calculation of the ARR was obtained after data had been filtered and extreme outlying electrodes and periods had been rejected (and before any of the MWF cleaning steps were applied).

Equation S4:

$$ARR_i = 10log_{10} \frac{E\{(d_i)^2\}}{E\{\left(y_i - \hat{d}_i\right)^2\}} \; (for\ artifact\ segments)$$

The units for the SER and ARR measures are decibels (dB). It is worth noting that ICA approaches may detect and remove artifacts other than the most common artifacts captured by our MWF cleaning templates. As such, ICA approaches may seem to “distort” clean periods using the SER metric. As such, it is more appropriate to compare across pipelines that apply cleaning to all periods (such as those that implement ICA, CCA, or Artifact Subspace Reconstruction [ASR]), and perhaps not appropriate to compare those pipelines to the MWF_only approach (which does not detect artifacts in the clean periods at all). For this reason, we have used a number of additional metrics to the SER and ARR, in order to fully characterize artifact reduction (with the blink amplitude ratio, artifacts remaining showing muscle activity, and variance explained by brain activity detected by ICLabel after cleaning) and preservation of signal (with the measures of variance explained by the experimental manipulation).

Additionally, because the SER and ARR metrics are based on the variance in the clean and artifact periods, it is possible for the metrics to be biased by low-powered brain signals during the artifact periods more commonly than the clean periods. For example, at times the clean periods may have showed more alpha activity for example (and thus high variance), while the muscle affected periods show less alpha activity and only low powered muscle activity. In this case, sometimes the variance of the artifact periods may have been less than the variance of the clean periods, leading to very low ARR and very high SER values. In fact, because individual electrodes within this metric are scaled by the amount that the artifact period variance exceeds the clean period variance, with electrodes showing higher clean variance than artifact variance set to zero before the weighting based on amount of variance in each electrode (by dividing electrodes variance by the total of all electrodes), it is possible for all electrodes to be set to zero, and the SER and ARR to produce NaN values. As such, this metric is perhaps less ideal for evaluating files where only small amplitude muscle artifacts are present (but is well suited to evaluating blink activity or high-power muscle activity, which is almost always higher in amplitude than the non-blink/non-muscle periods).

In order to address this issue, we have also used muscle activity artifact specific metrics (described in the following sections).

***Blink Amplitude Ratio***

The BAR metric [16] provides a ratio of the absolute amplitude within periods marked as blinks to the periods on either side of the blink. When applied to cleaned data, the measure provides a good indication of whether the cleaning pipeline has effectively cleaned the blink (leading to values ~=1). Alternatively, the metric indicates if the blink has been under-cleaned (leading to values of >1), or the subtraction of a blink artifact component has included the influence of brain activity as well as blink related activity, so that the subtraction creates a negative deflection where the blink was previously (also leading to values >1 due to the absolute transform). The metric also indicates if blinks are over-cleaned so that both positive and negative signals have been reduced towards zero (leading to values <1). To compute BAR, we epoched data for 4 seconds centered on the blink maximum, excluding epochs that included more than one blink within this 4 second epoch (to prevent these additional blinks from influencing the baseline period). We baseline corrected the epochs by subtracting the average of the first 500ms and last 500ms of the epoch. We then performed an absolute transform on all data in the epoch, then divided the mean of the 1 second centred on the blink maximum by the mean of the first 500ms and last 500ms of the epoch. For analysis, we examined both the frontal BAR (fBAR), which was the average BAR across electrodes FP1, FPz, FP2, AF3 and AF4, and the average BAR over all electrodes (allBAR).

***Log-frequency Log-power Slopes Indicating Muscle Activity***

We examined the proportion of epochs that contained any electrode with likely muscle activity remaining after cleaning, using the log-power log-frequency slope threshold of -0.59 [17]. We also examined the amount by which epochs showing remaining muscle activity exceeded the threshold, by subtracting -0.59 from the log-power log-frequency slope values from all epoch / electrode datapoints that exceeded this threshold, then averaged across all these remaining values (providing a value reflecting the average amount the slopes exceeded the log-power log-frequency slope threshold of -0.59 in the epochs and electrodes that showed muscle activity remaining after cleaning). It is worth noting that this the amount by which epochs showing remaining muscle activity exceed the threshold could be a misleading metric of the impact of muscle related artifacts a minority cleaned files. The metric is calculated only from epochs that show muscle slopes above the threshold. As such, if only a single epoch is still affected, but that one epoch shows a very severe muscle artifact, the metric will provide a very high score for that file, but the impact of the artifact on experimental measures may be very low. However, across the large number of files included in our analysis, the effect of such outliers is minimal (particularly since we used robust statistics, which excluded these outliers when calculating statistical effects).

***ICA Variance Categorized by ICLabel***

We examined the amount of ICA variance attributed to components categorized as brain activity by ICLabel. This was computed by summing the amount of variance in the EEG data explained by components categorized as brain by ICLabel (after cudaICA). Variance was calculated for each component individually using `compvar` (EEGLAB). An absolute transform was performed on the value of variance for each component to ensure all components provided a positive value for the amount of variance the component contributed to the data. This was performed because `compvar` provides negative values if a component influences the data in the opposite direction to the overall trend. However, for our purposes we were only interested in the percentage of total variance of the data that was influenced by brain activity. As such, negative variance values were made positive with this absolute transform so that their influence would not reduce the sum of brain activity component variance or artifact activity component variance, and the total values from all components would be equivalent to 100%. Following this, the variance from all brain components was summed and the variance from all artifact components was summed. The summed variance for brain activity was divided by the sum of the total brain variance and total artifact variance, and multiplied by 100 to obtain a percentage of the variance explained by brain activity (as determined by ICLabel). Note that methods that subtract components such as ICA_subtract and CCA were excluded from this measure, as component subtraction completely removed any variability from that artifactual component, so the contribution to variance from artifactual components would be 0 for these artifacts from these methods. Since this metric is only applicable for some pipelines, we only report the results of these analyses in the supplementary materials. It should be noted that infomax (with cudaICA) was used to compute the ICA artifact components for selection by ICLabel. Given the use of a common ICA method for this metric across all pipelines, this approach may have biased the metric towards the pipelines using infomax and against the fastICA / AMICA pipelines.

***Proportion of Epochs Rejected***

We examined the proportion of epochs that were rejected by the cleaning pipeline, after both excluding outlying data in the initial pre-cleaning steps and rejecting outlying epochs in the final stage prior to data analysis. With regards to the rejection of outlying epochs after cleaning, we used an approach typical in the literature, applying an automated algorithm rejected epochs that still showed potential artifacts, as defined by kurtosis or improbable data with a value higher than 5 SD from the mean at any electrode or 3 SD from the mean for all electrodes (using the relevant EEGLAB functions `pop_rejkurt` and `pop_jointprob`), or epochs showing values outside of a -60 to 60 microvolt window.

***ERP Trial Type Comparisons - Variance Explained Metrics***

Perhaps most importantly, we assessed the amount of variance explained by a variety of experimental manipulations that are well established to provide differentiation of neural activity in the comparison of two conditions after cleaning by the pipelines. This assessed the real-world applicability of each cleaning pipeline [18]. Ideally, effectively cleaned EEG data should lead to data that still contains all of the brain signal, and none of the artifact. Data cleaned this effectively should in theory produce the largest amount of variance explained by different experimental manipulations. This is because we assume that non-neural artifacts are unrelated to different experimental manipulations (so their inclusion in the data would contribute noise to a comparison between two experimental conditions, reducing the variance explained), and neural activity *is* related to the experimental manipulation (so maintaining more of the neural activity leads to increased detection of the effect of the experimental manipulation on brain activity and thus more variance explained by the experimental manipulation). As such, the best cleaning pipelines should provide the maximal between condition effects, with the largest amount of variance explained by the experimental manipulation [18].

In order to test the amount of variance explained by the different experimental manipulations, we used the randomization graphical user interface (RAGU) [19, 20], which compared differences across all electrodes available from averaged ERP data (within experimental condition and across each pipeline) using randomisation statistics (while controlling for multiple comparisons across both the spatial and temporal dimension). This toolbox additionally provides the ability to test for differences in overall neural response strength (using the global field potential (GFP) or root mean squared test) and separately to test for differences in the distribution of neural activity (using the global field potential dissimilarity between conditions after the recommended L2 normalisation for differences in global field potential or root mean squared). More details on this toolbox can be found in [19, 20]. Given the potential computation time when including all statistical tests, 1000 permutations were used for all tests within RAGU.

We computed the explained variance for between condition comparisons from each pipeline across two real world experimental designs – the Go-Nogo effect and the difference between error response and correct responses (more details provided below). We selected these experimental effect related neural activities because they have been well validated by previous research, showing differences between the conditions. The specific conditions were also selected because of they are commonly of interest in EEG research. To compute the explained variance, we averaged the explained variance across time periods where the different conditions have been suggested to show the strongest or most robust differences in neural activity by previous literature. As such, effective cleaning should produce larger effect sizes, with higher levels of explained variance produced by more effective cleaning pipelines. We have provided statistical comparisons of the ability of the different pipelines to differentiate these experimental conditions by examining the interactions between different pairs of pipelines and the condition of interest (for example between the Go and Nogo trials). We provide heat maps depicting the variance explained by this interaction for each pair of pipelines, marking the interactions that were significant. Significance values were corrected for multiple comparisons across all interaction comparisons within each metric using the Benjamini-Hochberg [21] false discovery rate (* indicates FDR-p < 0.05). We also provide indication of which pipeline provided larger values for variance explained using – and + symbols, which can be interpreted as the pipeline listed on the left of the heatmap having shown less (-) or more (+) variance explained in the comparison between the two experimental conditions than the pipeline listed at the bottom of the heatmap.

First, we examined the amount of variance explained by the difference between correct and error responses within the Go-Nogo task. Out of the full dataset of 127 files, a total of 76 participants provided a minimum of 6 artifact free error related epochs available for analysis. These error response locked epochs were averaged into ERPs for analysis after cleaning by each pipeline. A matched number and matched reaction time set of correct response epochs were also averaged for analysis after cleaning by each pipeline. We baseline corrected these epochs to the -400 to -100ms period. Out of the full dataset of 127 files, a total of 76 participants provided a minimum of 6 artifact free error related epochs available for analysis. We examined the variance explained in the difference between error and correct response trials in the Pe GFP from 150 to 300ms, the ERN TANOVA from 0 to 150ms (note that the ERN did not show a difference between errors and corrects in the GFP), and the Pe TANOVA (200 to 400ms).

Second, we examined ERPs from the Go-Nogo task (N = 127). The inclusion of comparisons across two separate studies provided an extra indicator of the consistency of the results. All Go and Nogo epochs available after cleaning were baseline corrected using a regression baseline correction to the -200 to 0ms period [24]. After this, the Go and Nogo trials were separately averaged to create ERPs. We made comparisons between Go and Nogo trials in the N2 GFP (180 to 300ms), the P3 GFP (300 to 500ms), then N2 TANOVA (180 to 300ms) and the P3 TANOVA (300 to 500ms).

***ERP Amplitude Reliability Metrics***
In order to assess how consistent and reliable error related ERP data were after cleaning, we examined the dependability (a generalisation statistics measure) of the error positivity (Pe) following error responses in a Go-Nogo task. We extracted the Pe on each error trial from each participant as the average amplitude at electrode FCz from 200 to 400ms post error, after baseline correction to the -400 to -100ms period (a typical analysis approach for the Pe) and submitted these values to the ERP Reliability Analysis (ERA) toolbox [22, 23] for each pipeline. We report the minimum number of trials from each participant required for dependability value of 0.8, and given this number, the number of excluded participants (out of a possible 76) who had fewer than this number of epochs remaining from each pre-processing approach. Note that the number of participants excluded could vary between pipelines that produced the same minimum number of trials for dependability, if one of the cleaning pipelines required more epochs or data to be rejected as an outlier (leaving fewer trials available for analysis).

Second, we calculated metrics to examine the reliability of ERPs amplitudes after cleaning. We examined the standardized measurement error of the voltage peaks within Nogo N2 and Go P3 time windows from single electrodes of interest [25]. The SME is defined as “the standard error of measurement for an ERP amplitude or latency score, assuming that the score is obtained from a single participant’s averaged ERP waveform. Formally, this means that the SME is an estimate of the standard deviation of the sampling distribution for a given participant’s amplitude or latency score” [25]. This measure represents the precision of the output values used and quantifies the extent to which noise impacts the outcomes. We used the ERPLAB Toolbox to calculate the SME [26] (https://erpinfo.org/erplab). The analytic SME is calculated by computing the standard deviation of the outcome measure within single participants and dividing this value by the square root of the number of epochs the individual provided and is recommended for window of interest analyses. This analytic SME is on average equal to the empirical SME and the bootstrapped SME (bSME) [25]. The bSME is constructed from re-sampling from replacement, and is recommended for peak amplitude approaches. We ran the analysis on peak detections (rather than averaged windows of interest), as peak detection methods of measuring ERPs are more vulnerable to artifacts, since high frequency muscle artifacts can result in a spike in a small number of timepoints (which can be averaged out by average window ERP measures). As such, we used the bSME for our peak amplitude estimates, with 1000 bootstraps performed. We computed the bSME for the N2 peak amplitudes (voltage minimum between 180 and 300ms after the stimuli) from Nogo trials at FCz, and the P3 peak amplitudes (voltage maximum between 300 and 500ms after the stimuli) from Go trials at Pz. Because the N1 was prominent after both Go and Nogo trials, we also computed the bSME for the N1 peak amplitudes (voltage minimum between 60 and 180ms after the stimuli) from both trial types at FCz.

### SECTION FOUR

#### Supplementary Results

In this section, we have provided a rank order (by mean) of the best performing pipelines to worst performing pipelines, interpreted from the post-hoc tests which can be visualised in heatmap figures. Significant differences are highlighted for pipelines that performed significantly better than other pipelines using the following notation for ease of understanding: better performance > worse performance (***it is important to note that we have used this better performance > worse performance approach rather than a higher values > lower values approach, as we hope that the consistency will help the reader understand each of the results in the context of all other results***). Because sometimes pipeline 1 differed from pipeline 2, but pipeline 3 did not differ from either 1 or 2, we have used the following notation: ^ = significantly higher than the pipeline marked with a ^^ within the same section (while the others in the category are not significantly different from each other). * = significantly higher than the pipeline marked with a ** in the same category, and so on for the following symbols: $^{+@\$!+}$. For each post-hoc figure, values reflect the 95% confidence intervals for the comparison between each pipeline listed on the left, and each pipeline listed along the bottom. Asterix's indicate significant results after multiple comparison controls were applied using the robust post-hoc t-test function "rmmcp", which uses Hochberg's approach to control for the FWE ($p < 0.05$). Note that because the post-hoc t-test significant values were derived from the robust statistics, and the 95% confidence intervals were calculated in the usual parametric manner, sometimes the confidence intervals overlapped with 0 while at the same time the comparison was marked as significant. We interpreted significant differences from the p-value rather than the confidence intervals, but both can be visualised in the figures if the reader would prefer to interpret significance from the confidence intervals.

##### *EEG Data Cleaning Performance Metrics*

###### *Signal to Error Ratio*

In the Go-Nogo dataset, the robust ANOVA showed that there was a significant difference in SER between the pipelines for the Go-Nogo data: $F(2.73, 207.17) = 246.8577$, $p < 0.0001$. The rank order from best performing pipeline to worst performing pipeline of significant differences between individual cleaning pipelines from post-hoc t-tests was as follows: MWF_only > MWF_CCA, wICA_ICLabel > MWF_wICA_AMICA > MWF_wICA_CCA^, MWF_wICA_infomax, MWF_wICA_fastICA^^ > MWF_wICA_45Hz > ICA_subtract > MWF_ICA_subtract > wICA_all (Figure S1-3).

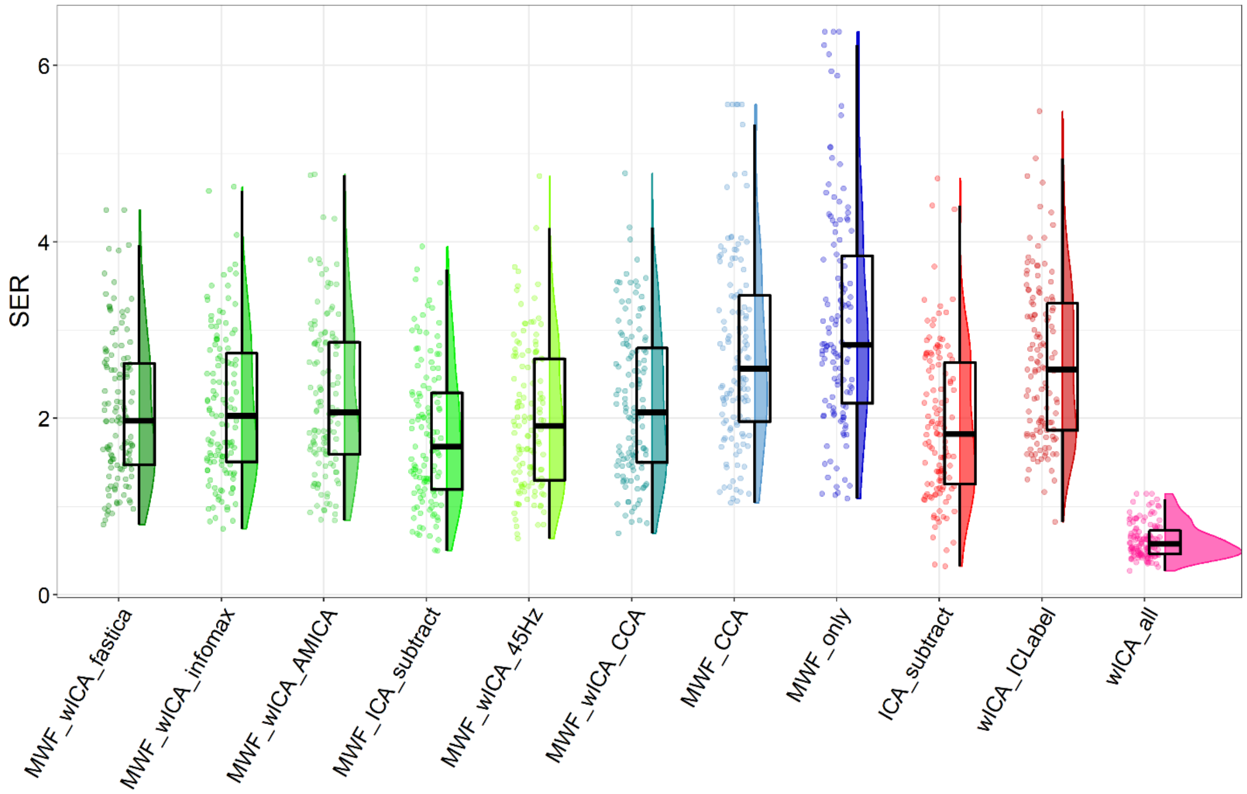

Figure S1. Raincloud plot depicting Signal to Error Ratio (SER) values from the Go-Nogo data (N = 127) for each of the cleaning pipelines. Note that this figure reflects winsorized data to enable easier visualization with a reduced data spread and smaller scale, so that the pipelines can be more easily discriminated. The full data is depicted in the Figure S2.

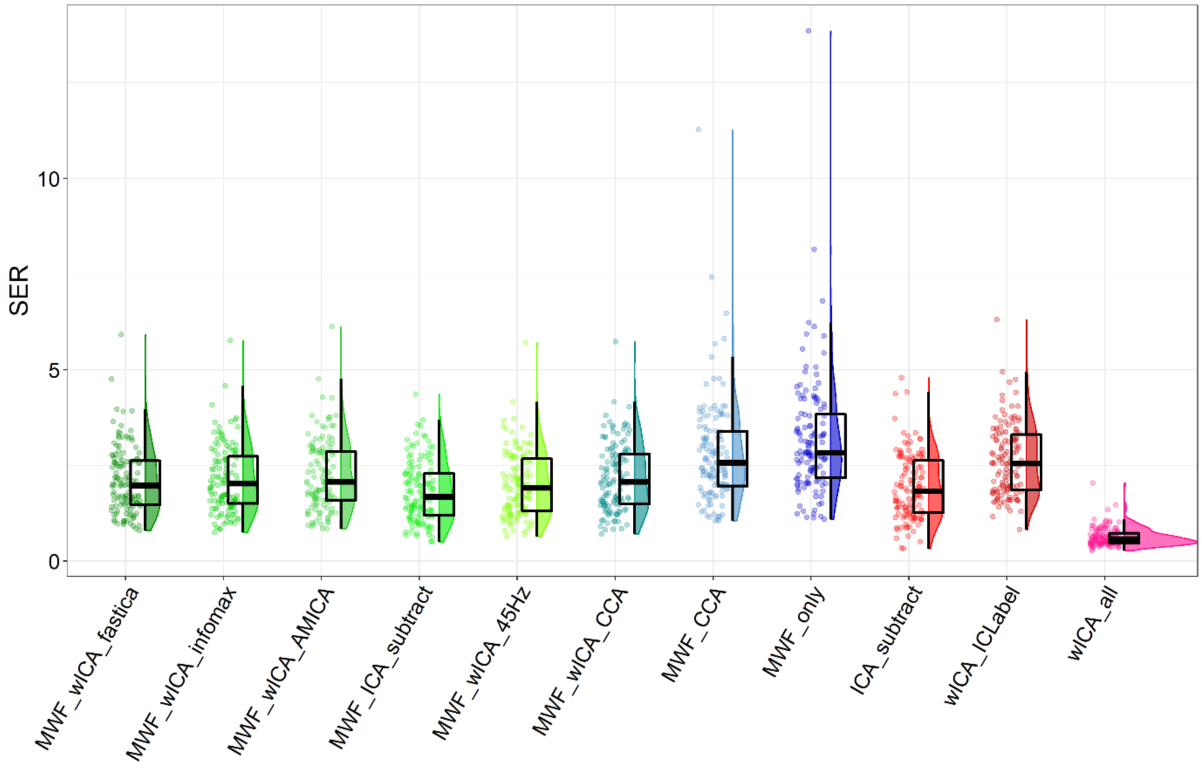

Figure S2. Raincloud plot of Signal to Error Ratio (SER) values without winsorizing outliers so the full spread of data can be visualized for the Go-Nogo dataset.

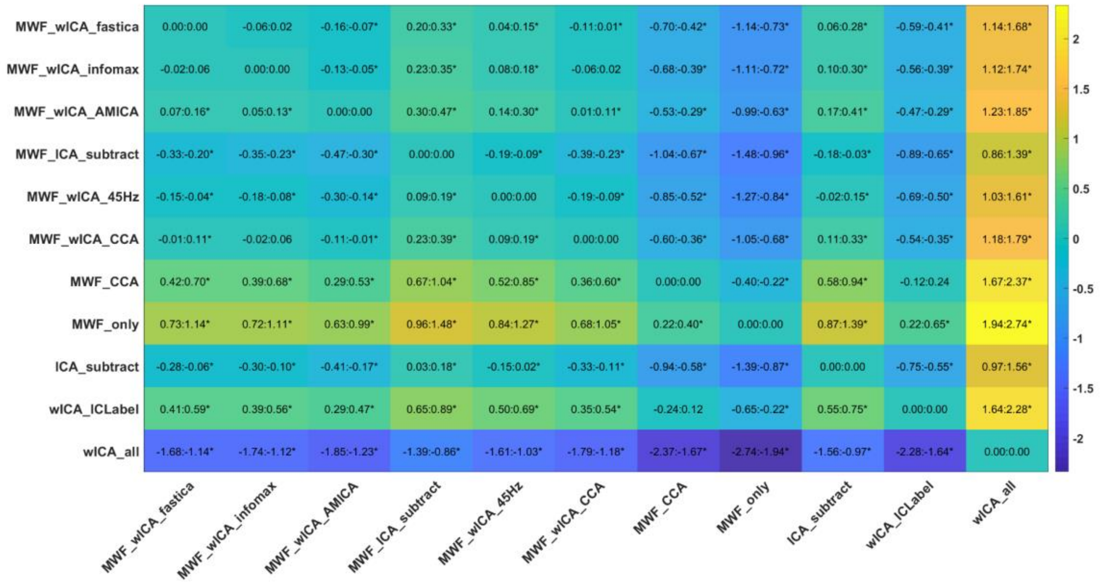

Figure S3. Post-hoc test of Signal to Error Ratio (SER) values for the Go-Nogo dataset.

#### *Artifact to Residue Ratio*

There was a significant difference in ARR between the pipelines for the Go-Nogo data: $F(2.44, 185.64) = 1015.293$, $p < 0.0001$. The rank order from best performing pipeline to worst performing pipeline of significant differences between individual cleaning pipelines from post-hoc t-tests was as follows: wICA_all > MWF_wICA_45Hz > MWF_ICA_subtract > MWF_wICA_infomax, MWF_wICA_fastICA > MWF_wICA_CCA, MWF_wICA_AMICA > ICA_subtract > MWF_CCA, wICA_ICLabel > MWF_only (Figure S4-6). When SER and ARR values were viewed together in a scatterplot, the Go-Nogo datasets showed an almost identical pattern to the combined Sternberg and resting dataset in our companion article, with the MWF_wICA methods showing a higher combination of SER and ARR at the same time than ICA_subtract (which showed lower ARR values) and higher ARR values than wICA_all, MWF_only and MWF_CCA, but slightly lower SER values (Figure S5).

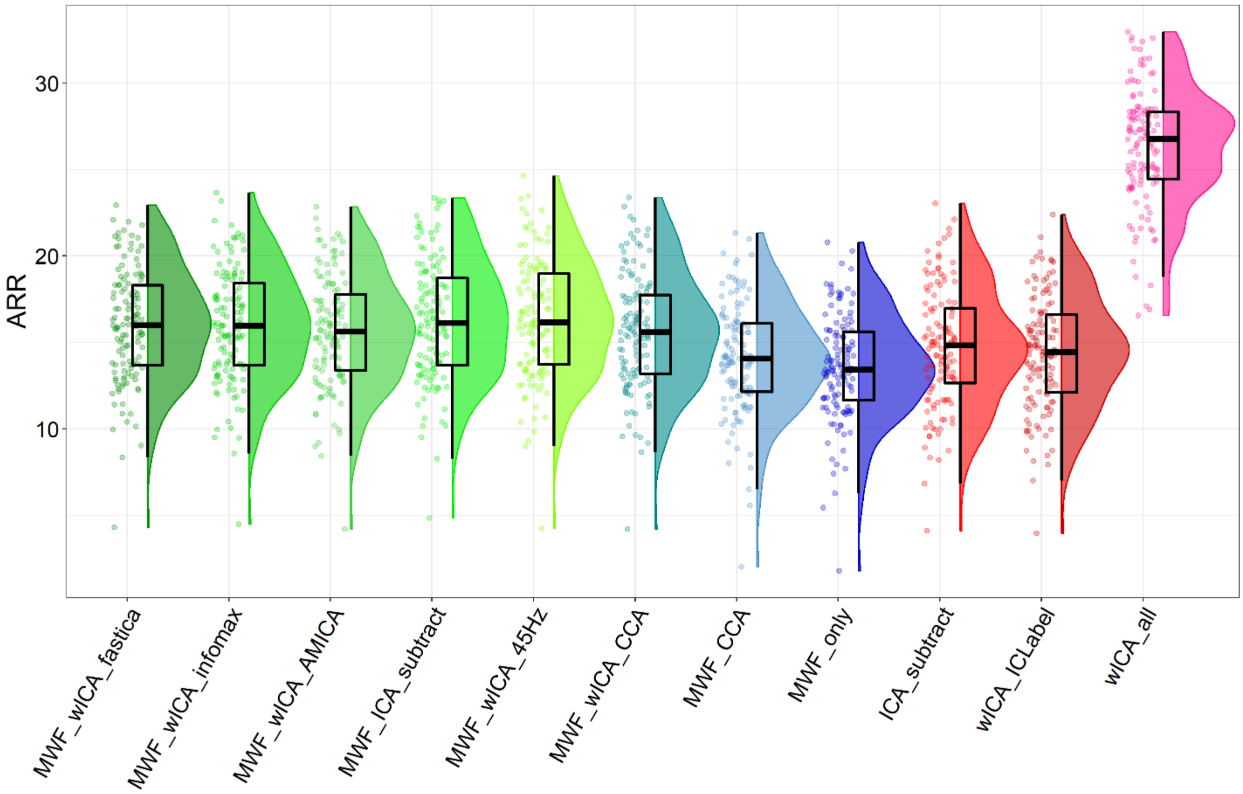

Figure S4. Raincloud plot depicting Artifact to Residue Ratio (ARR) values from the Go-Nogo data (N = 127) for each of the cleaning pipelines.

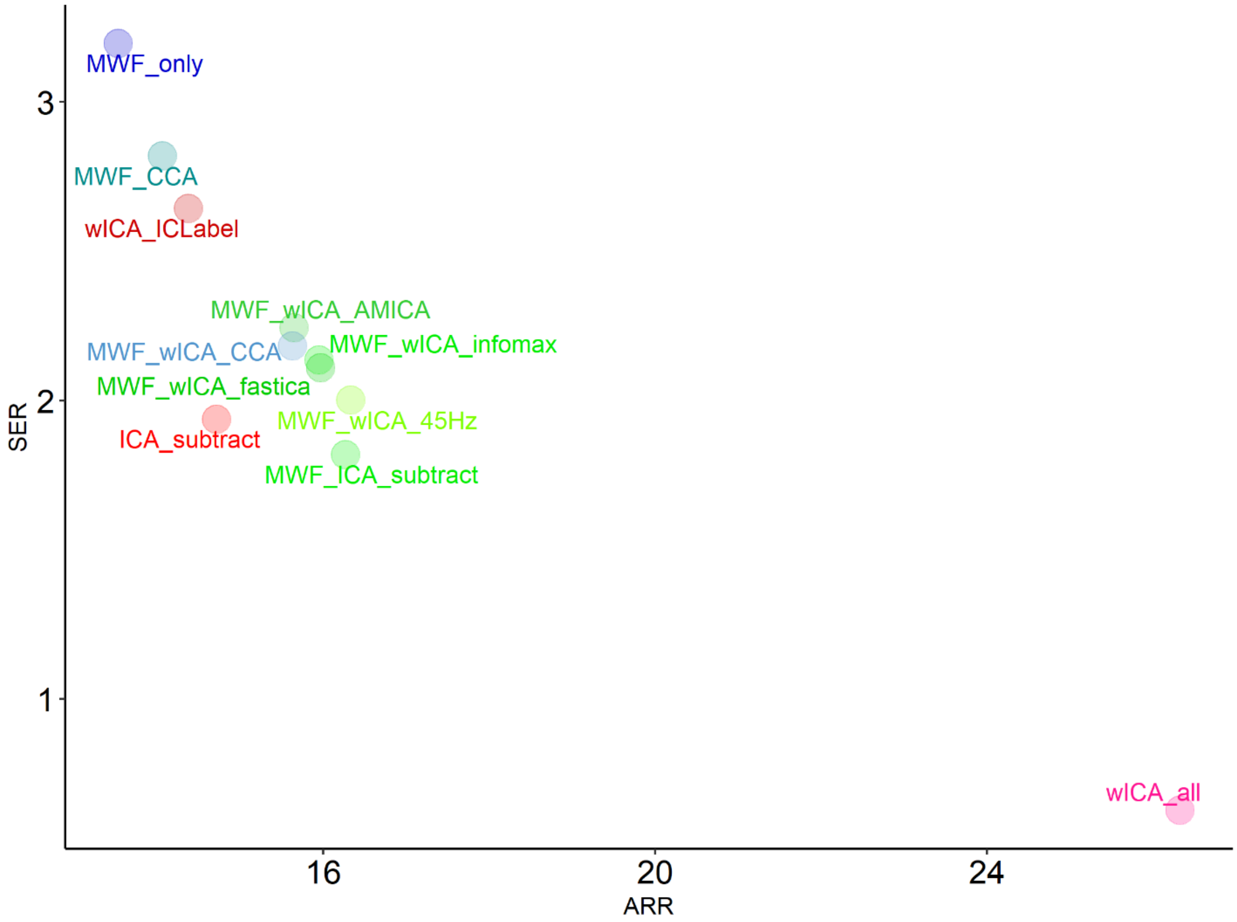

Figure S5. A scatterplot depicting both SER and ARR values for the Go-Nogo dataset from each cleaning pipeline.

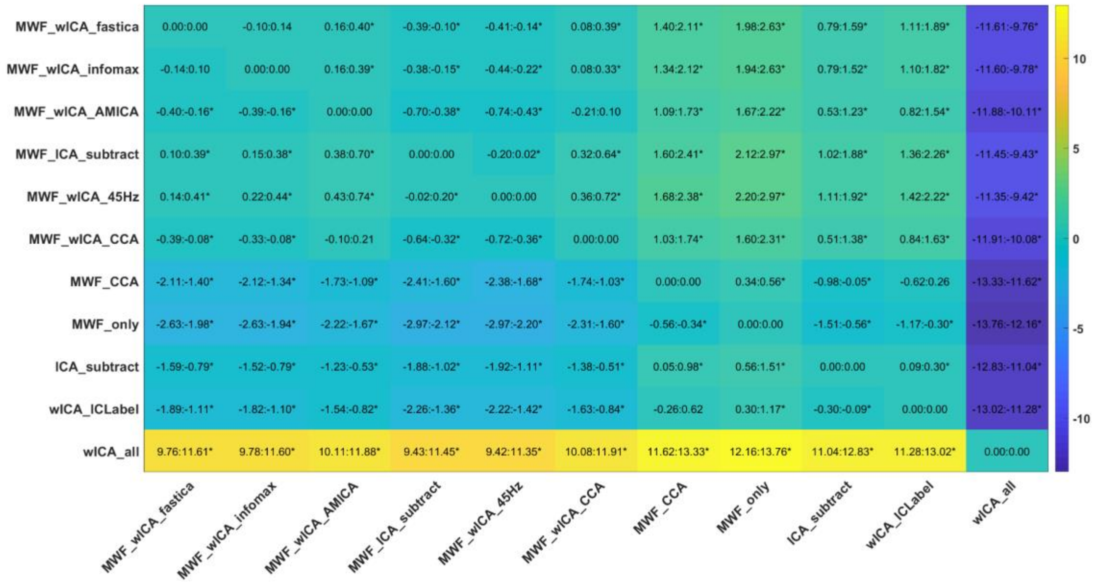

Figure S6. Post-hoc test of ARR values for the Go-Nogo dataset.

| | SER | | ARR | |
|---|---|---|---|---|
| **Pipeline** | **Mean** | **SD** | **Mean** | **SD** |
| **ICA_subtract** | 1.937 | 0.854 | 14.711 | 3.460 |
| **MWF_CCA** | 2.819 | 1.366 | 14.058 | 3.183 |
| **MWF_wICA_CCA** | 2.183 | 0.883 | 15.628 | 3.376 |
| **MWF_wICA_AMICA** | 2.243 | 0.926 | 15.648 | 3.262 |
| **MWF_wICA_fastICA** | 2.109 | 0.873 | 15.968 | 3.276 |
| **MWF_ICA_subtract** | 1.819 | 0.798 | 16.265 | 3.496 |
| **MWF_wICA_infomax** | 2.136 | 0.873 | 15.947 | 3.307 |
| **MWF_wICA_45Hz** | 2.001 | 0.848 | 16.331 | 3.480 |
| **MWF_only** | 3.196 | 1.590 | 13.529 | 3.087 |
| **wICA_all** | 0.627 | 0.242 | 26.321 | 3.427 |
| **wICA_ICLabel** | 2.644 | 0.931 | 14.372 | 3.296 |

Table S1. Means and SDs for Signal to Error Ratio (SER) and Artifact to Residue Ratio (ARR) values.

#### *Frontal Electrode Blink Amplitude Ratio*

One file was excluded from the blink analyses due to no blink epochs being available that did not also contain another blink in the baseline period for calculation of BAR. The robust ANOVA showed a significant difference in blink amplitude ratio in frontal electrodes between the pipelines: $F(4.26, 319.74) = 36.4994$, $p < 0.0001$. The rank order from best performing pipeline to worst performing pipeline of significant differences between individual cleaning pipelines from post-hoc t-tests was as follows: wICA_all > MWF_wICA_infomax, MWF_wICA_fastICA, MWF_ICA_subtract, MWF_wICA_AMICA, MWF_wICA_CCA > MWF_wICA_45Hz > ICA_subtract^, MWF_only, MWF_CCA, wICA_ICLabel^^ (Figure S7-9). Some data showed outliers suggesting inadequate cleaning of the blink artifact for these files (see Figure S8). As such, we recommend that studies should implement methods to check the severity of remaining blinks after cleaning (such as BAR) and if necessary, exclude remaining blink affected data after cleaning.

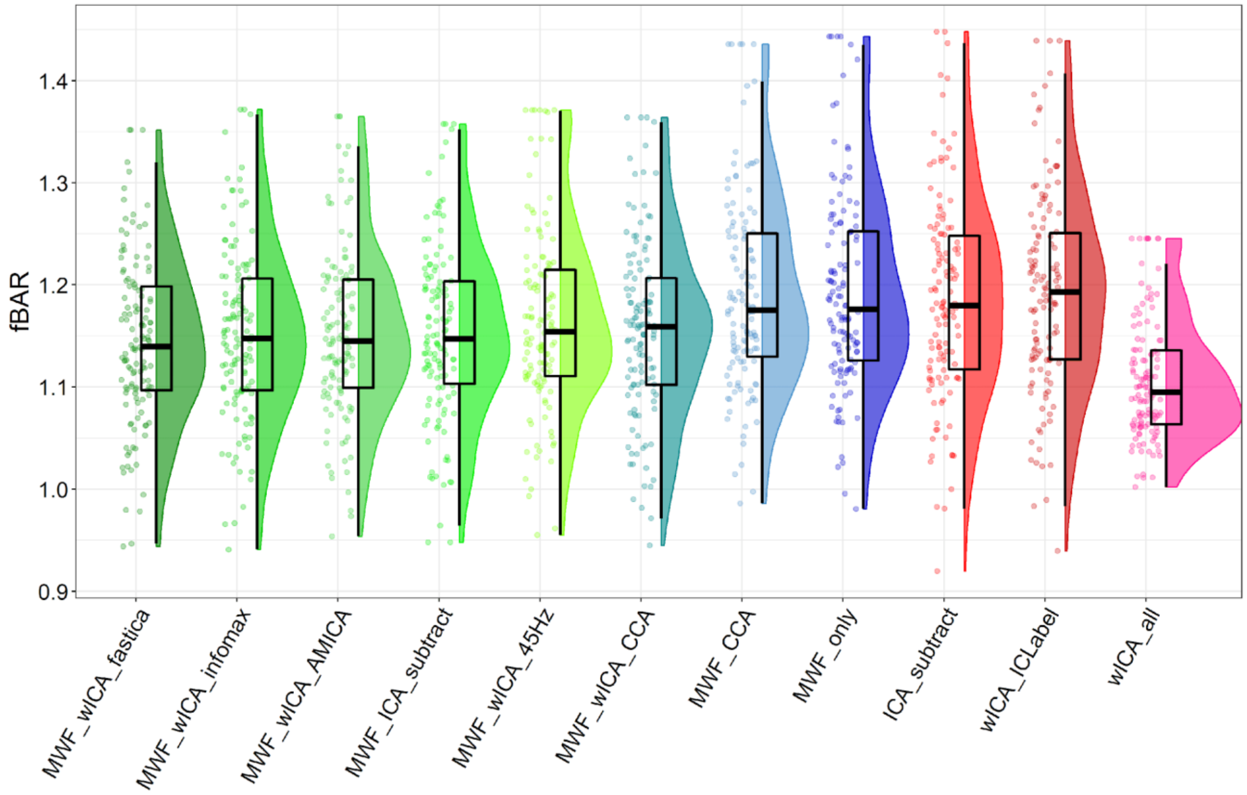

Figure S7. Raincloud plot depicting frontal blink amplitude ratio (fBAR) values from the Go-Nogo data (N = 126) for each of the cleaning pipelines. Note that this data has been winsorized to present a scale that enables visualisation of differences between pipelines – almost all datasets contained 1-3 outliers with fBARs > 2.25. Note also that one participant was excluded from this analysis for not showing any blink periods without a blink within the BAR baseline period. Plots depicting all unmodified data can be viewed in Figure S8.

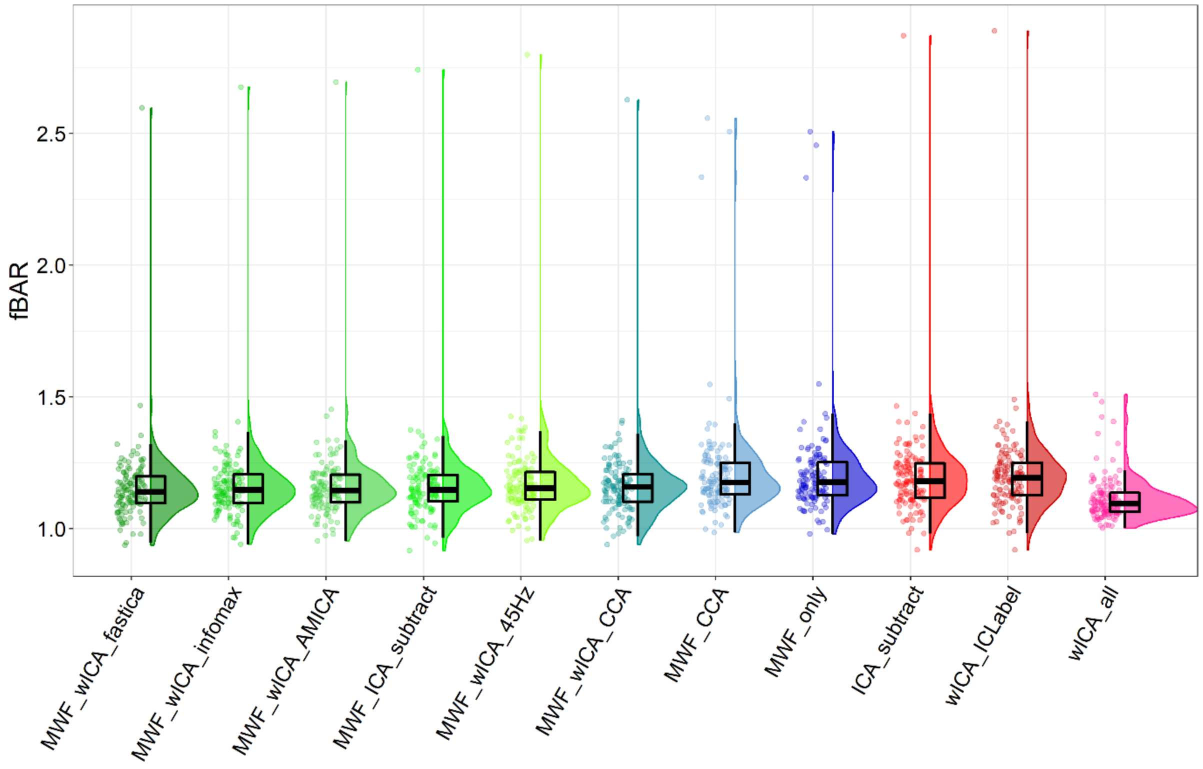

Figure S8. Frontal blink amplitude ratios, without winsorizing outliers for the Go-Nogo dataset.

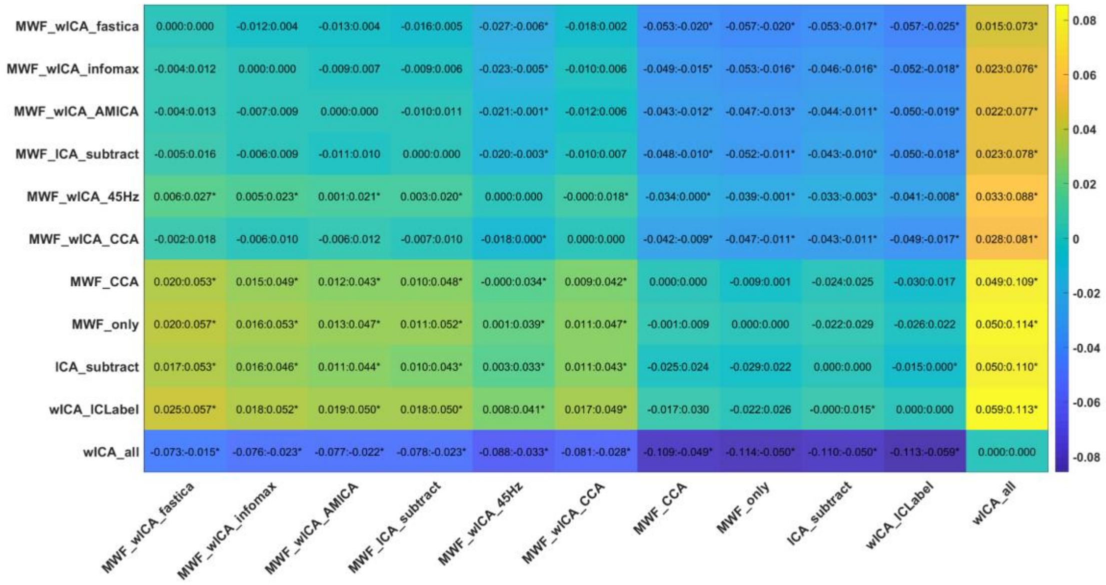

Figure S9. Post-hoc test for fBAR values for the Go-Nogo dataset.

#### *Statistical Comparisons of the Blink Amplitude Ratio Averaged Across All Electrodes*

The robust ANOVA showed a significant difference in blink amplitude ratio averaged across all electrodes between the pipelines: $F(3.76, 281.95) = 194.3259$, $p > 0.00001$. The rank order from best performing pipeline to worst performing pipeline of significant differences between individual cleaning pipelines from post-hoc t-tests was as follows: wICA_all > MWF_wICA_fastICA^, MWF_wICA_CCA^, MWF_wICA_infomax, MWF_ICA_subtract, MWF_wICA_AMICA, MWF_only, MWF_CCA^^ > MWF_wICA_45Hz > ICA_subtract, wICA_ICLabel. See Figure S10 for a raincloud plot depicting the distribution of the allBAR data. Note that this data has been winsorized to present a scale that enables visualisation of differences between pipelines – almost all datasets contained 1-3 outliers with allBARs > 2.25. Raincloud plots depicting all unmodified data can be viewed in supplementary materials Figure S11, and post-hoc comparisons can be viewed in Figure S12.

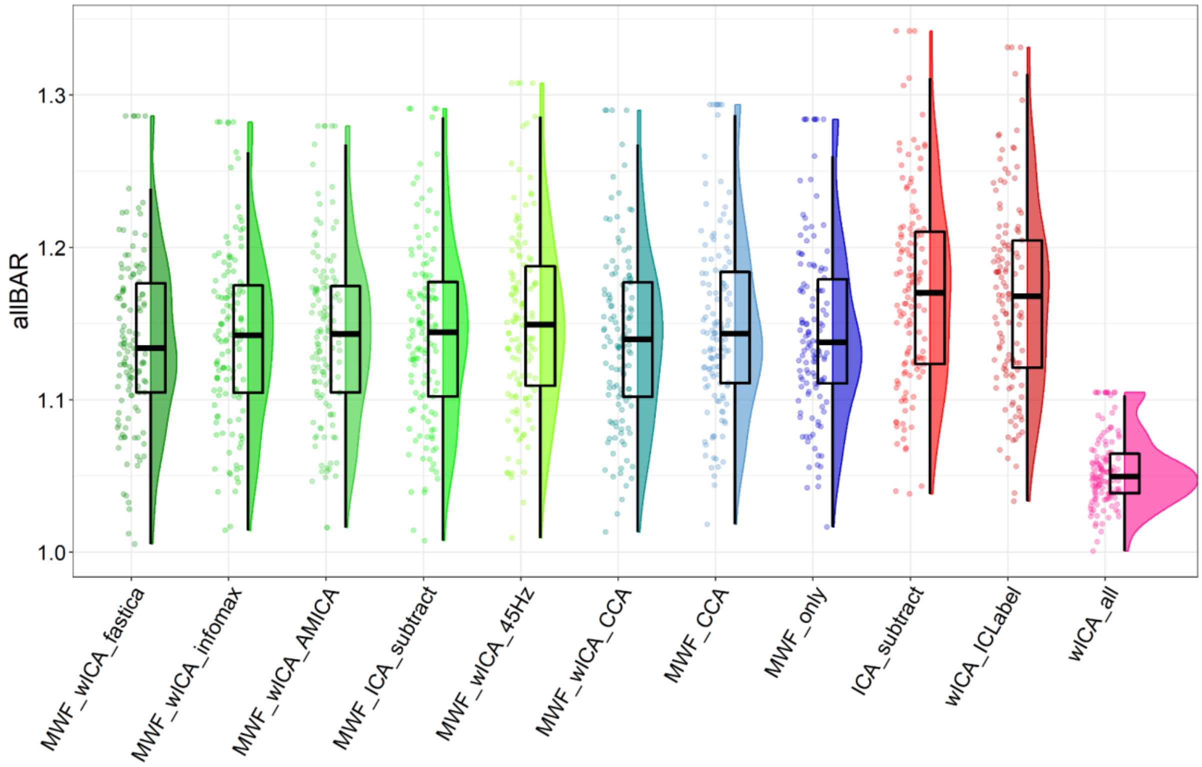

Figure S10. Raincloud plot depicting allBAR values from the Go-Nogo data (N = 126) for each of the cleaning pipelines. Note that this data has been winsorized to present a scale that enables visualisation of differences between pipelines – almost all datasets contained 1-3 outliers with allBARs > 2.25. Note also that one participant was excluded from this analysis for not showing any blink periods without a blink within the BAR baseline period. Raincloud plots depicting all unmodified data can be viewed in supplementary materials Figure S11.

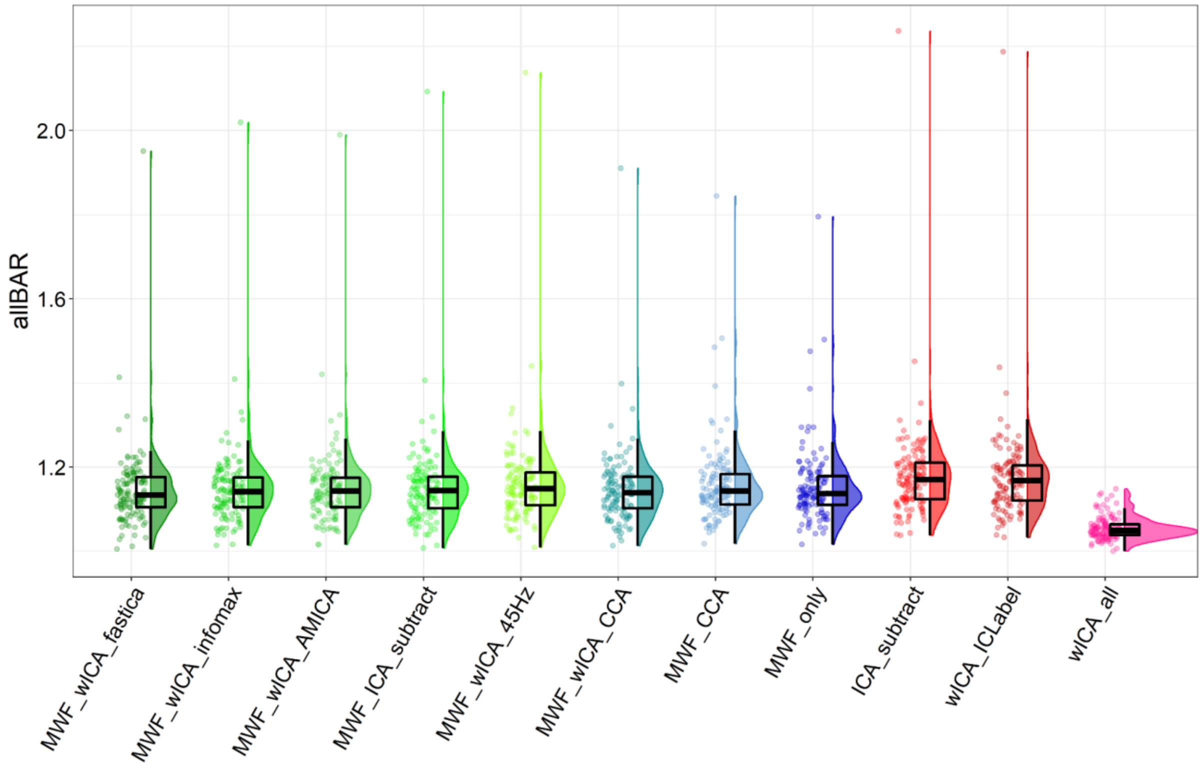

Figure S11. Raincloud plot depicting allBAR values from the Go-Nogo data (N = 126) for each of the cleaning pipelines.

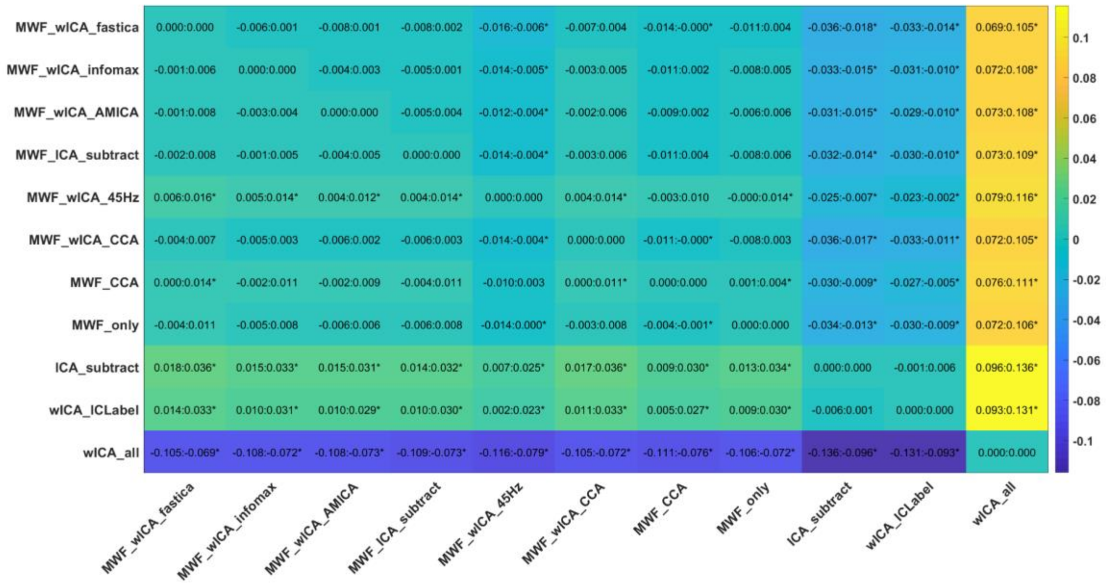

Figure S12. Post-hoc tests for allBAR values from the Go-Nogo dataset.

| | Frontal Blink Amplitude Ratio | | All Electrode Blink Amplitude Ratio | |
|---|---|---|---|---|
| **Pipeline** | **Mean** | **SD** | **Mean** | **SD** |
| **ICA_subtract** | 1.187 | 0.1 | 1.18 | 0.116 |
| **MWF_CCA** | 1.191 | 0.096 | 1.159 | 0.096 |
| **MWF_wICA_CCA** | 1.156 | 0.088 | 1.147 | 0.092 |
| **MWF_wICA_AMICA** | 1.152 | 0.084 | 1.151 | 0.097 |
| **MWF_wICA_fastICA** | 1.145 | 0.083 | 1.146 | 0.095 |
| **MWF_ICA_subtract** | 1.153 | 0.086 | 1.152 | 0.105 |
| **MWF_wICA_infomax** | 1.152 | 0.086 | 1.15 | 0.1 |
| **MWF_wICA_45Hz** | 1.165 | 0.091 | 1.16 | 0.109 |
| **MWF_only** | 1.194 | 0.098 | 1.155 | 0.093 |
| **wICA_all** | 1.107 | 0.059 | 1.055 | 0.027 |
| **wICA_ICLabel** | 1.193 | 0.099 | 1.176 | 0.112 |

Table S2. Blink amplitude ratio means and standard deviations.

#### *Proportion of Epochs Showing Muscle Activity After Cleaning*

The pipelines significantly differed in the number of epochs with log-power log-frequency slopes indicating muscle activity remaining after cleaning, with the robust ANOVA showing a significant effect: $F(1.84, 139.94) = 16398.28$, $p < 0.0001$. The rank order of significant differences between individual cleaning pipelines from post-hoc t-tests was as follows: MWF_ICA_subtract > MWF_CCA^, MWF_wICA_infomax, MWF_wICA_fastICA, MWF_wICA_CCA^^, MWF_wICA_AMICA^^ > ICA_subtract > wICA_ICLabel, MWF_only > wICA_all (Figure S13-15).

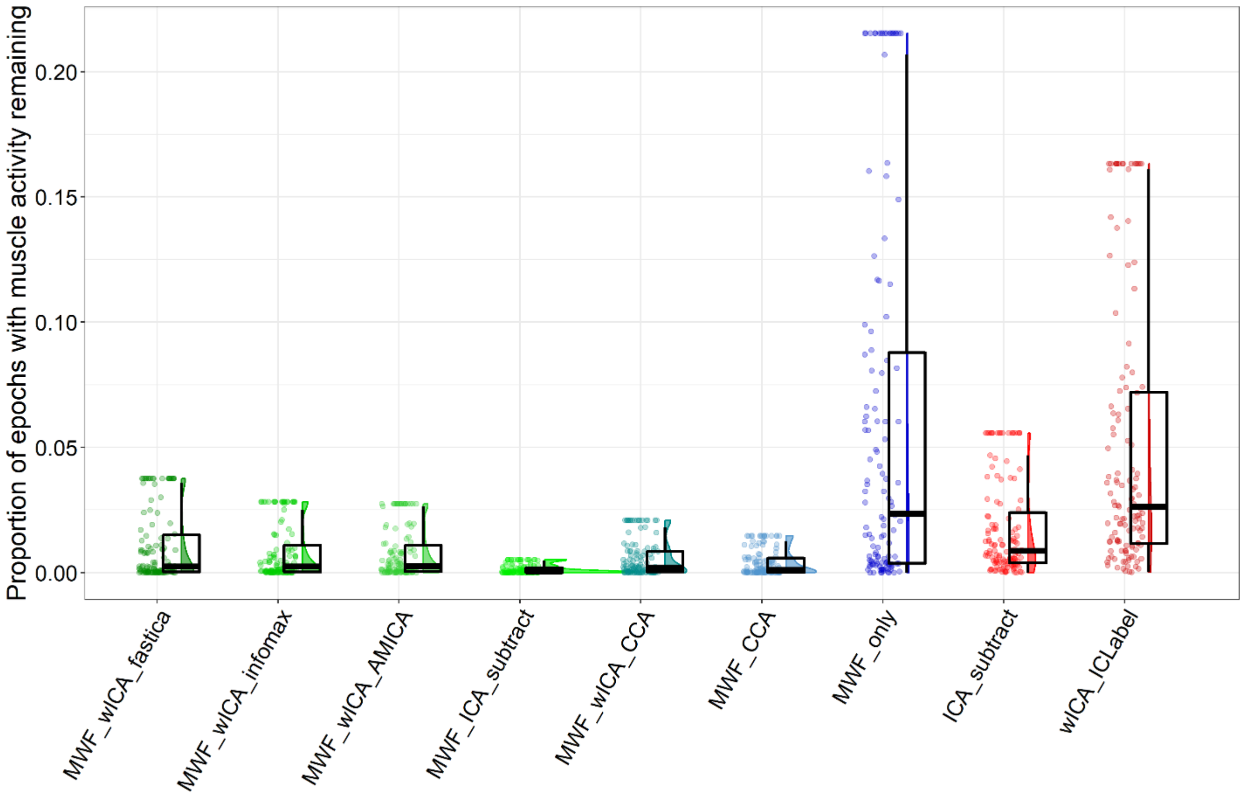

Figure S13. Raincloud plot depicting the proportion of epochs showing log-power log-frequency values above the -0.59 threshold for each of the cleaning pipelines. Note that this figure excludes wICA_all, as this pipeline showed median values > 0.75 and made the scale of the graph such that it was difficult to visualise differences in the other pipelines. Note also that we have winsorized the data in the figure, as the outliers also made the scale such that it was difficult to visualise differences in the other pipelines. The full data can be viewed in the supplementary materials Figure S14.

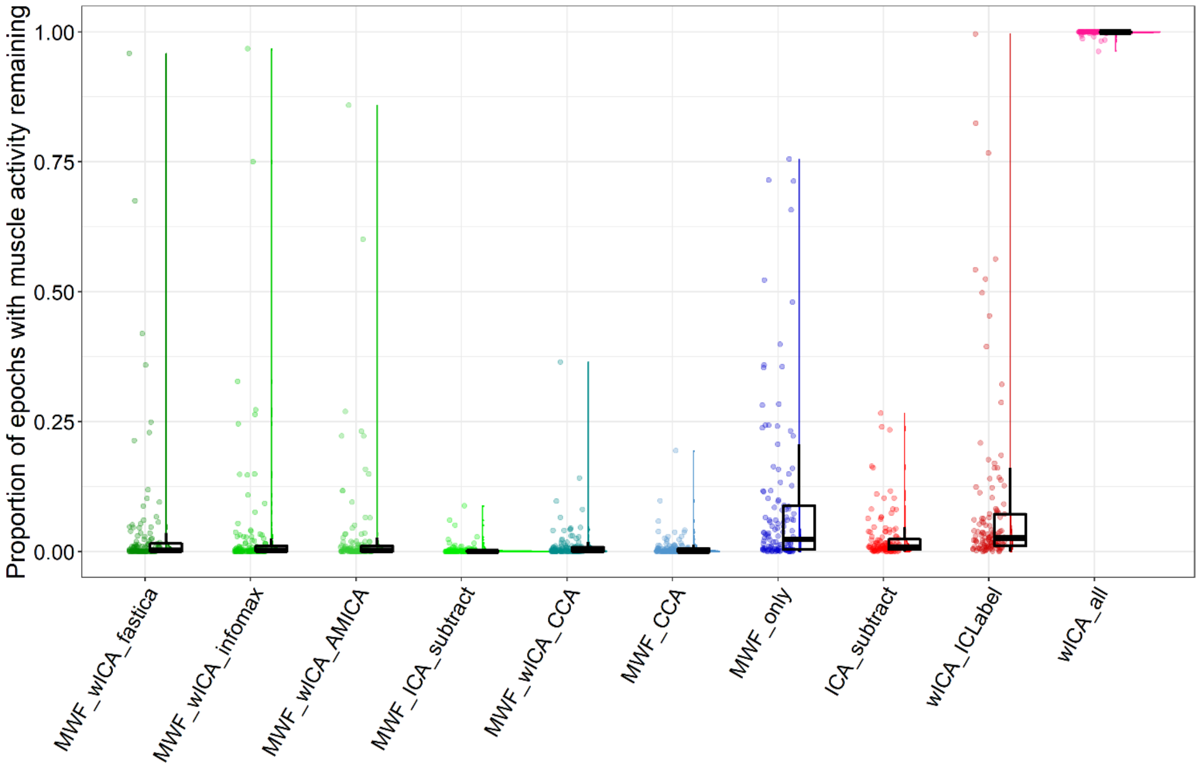

Figure S14. Proportion of epochs showing muscle slopes after cleaning including all pipelines and without winsorizing data.

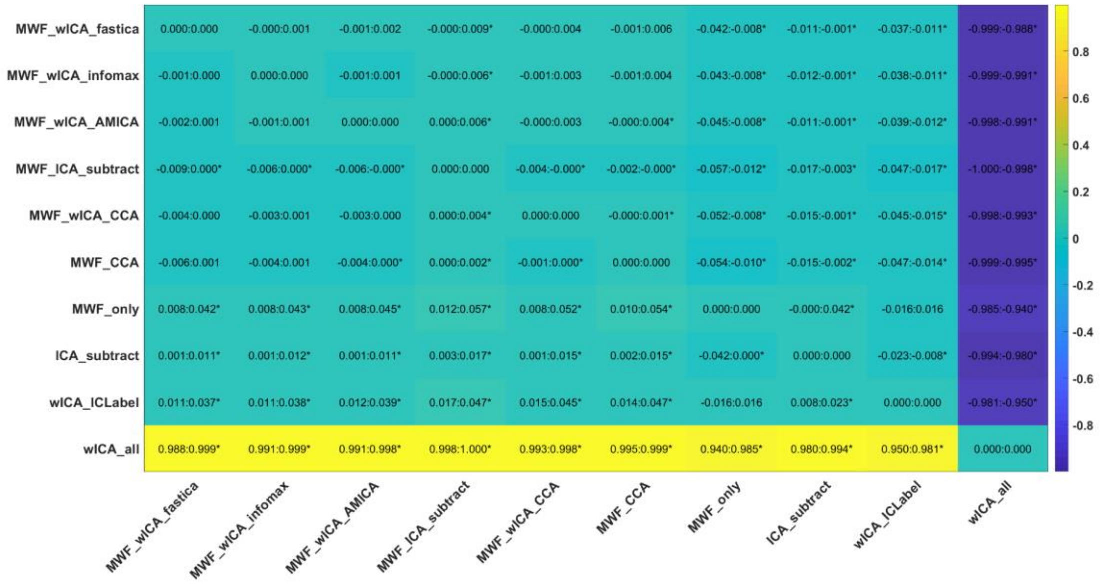

Figure S15. Post-hoc tests for the proportion of epochs showing muscle slopes after cleaning.

#### ***Severity of Muscle Slope Values from Epochs that Exceeded the Threshold***

There was a significant difference between the pipelines in the amount by which the mean slope exceeded the log-power log-frequency threshold from epochs and electrodes that showed muscle activity remaining: $F(5.54, 421.27) = 270.6692$, $p < 0.0001$. The rank order from best performing pipeline to worst performing pipeline of significant differences between individual cleaning pipelines from post-hoc t-tests was as follows: MWF_ICA_subtract^, MWF_wICA_infomax, MWF_CCA, MWF_wICA_fastICA^^, MWF_wICA_CCA^^, MWF_wICA_AMICA^^ > wICA_ICLabel, MWF_only, ICA_subtract, > wICA_all (Figure S16-17).

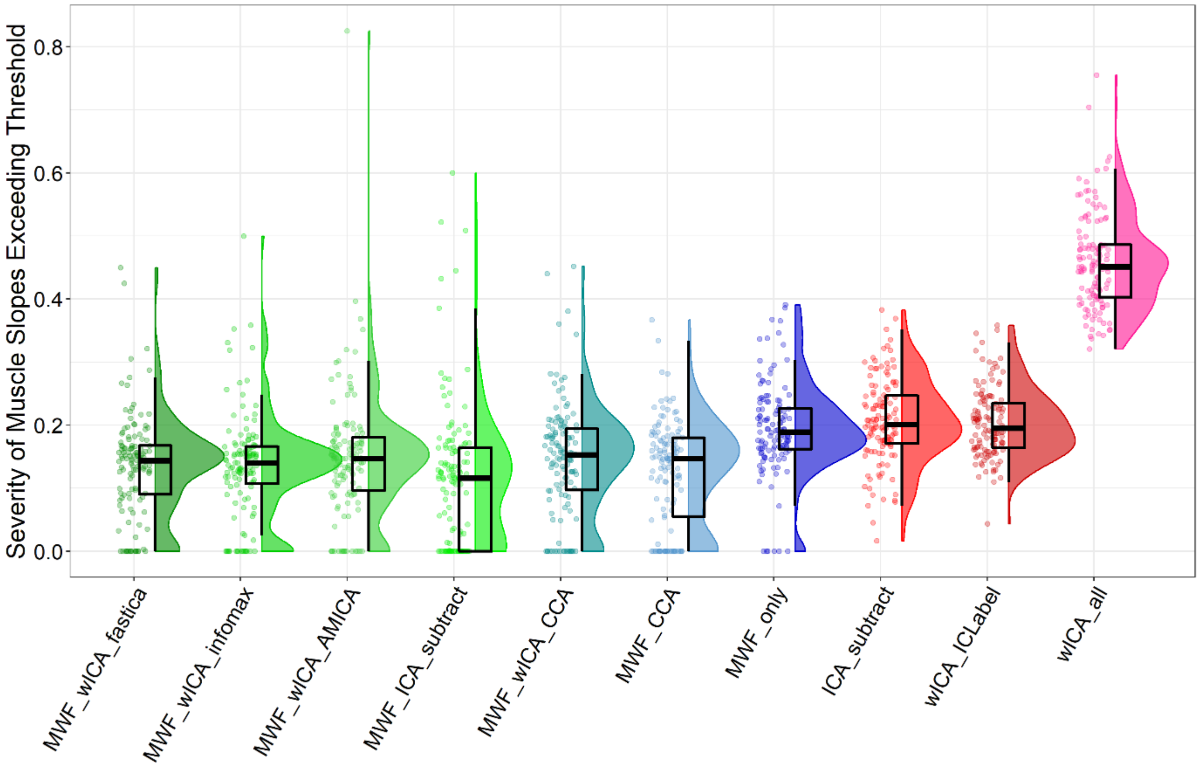

Figure S16. Raincloud plot depicting the amount by which log-power log-frequency slopes exceeded the -0.59 threshold, when values were averaged across super-threshold epochs and electrodes from the Go-Nogo data (N = 127) for each of the cleaning pipelines.

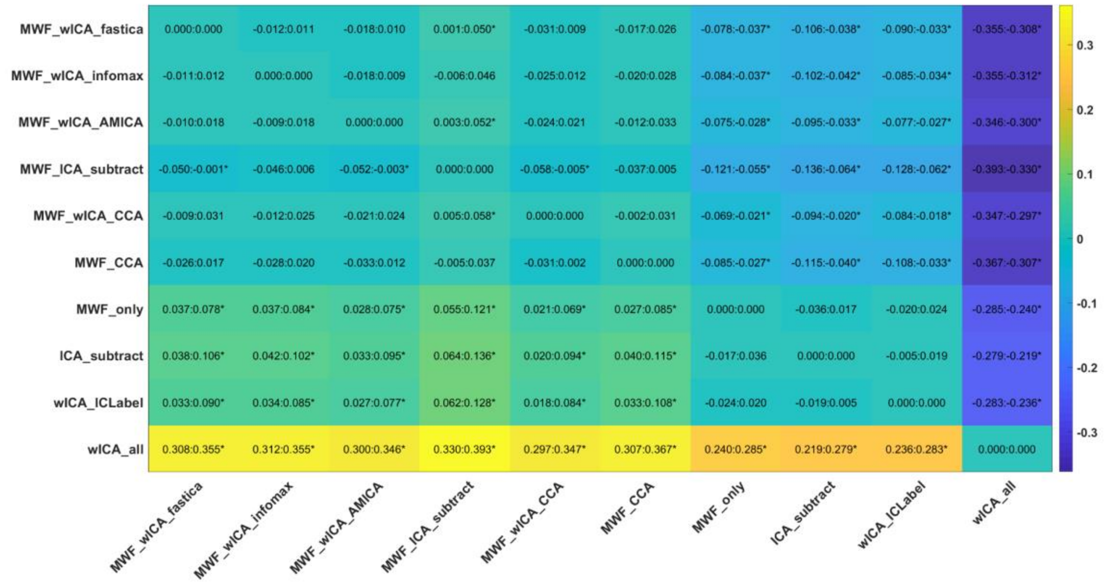

Figure S17. Post-hoc tests for the amount by which log-power log-frequency slopes exceeded the -0.59 threshold, when values were averaged across super-threshold epochs and electrodes.

| | **Proportion of epochs showing muscle slopes after cleaning** | | **Slope steepness over muscle slope threshold in epochs showing muscle slopes after cleaning** | |
|---|---|---|---|---|
| **Pipeline** | **Mean** | **SD** | **Mean** | **SD** |
| **ICA_subtract** | 0.017 | 0.019 | 0.207 | 0.067 |
| **MWF_CCA** | 0.004 | 0.005 | 0.125 | 0.084 |
| **MWF_wICA_CCA** | 0.006 | 0.007 | 0.147 | 0.09 |
| **MWF_wICA_AMICA** | 0.008 | 0.01 | 0.148 | 0.103 |
| **MWF_wICA_fastICA** | 0.011 | 0.014 | 0.135 | 0.081 |
| **MWF_ICA_subtract** | 0.001 | 0.002 | 0.116 | 0.115 |
| **MWF_wICA_infomax** | 0.008 | 0.011 | 0.136 | 0.083 |
| **MWF_only** | 0.061 | 0.075 | 0.194 | 0.073 |
| **wICA_all** | 1 | 0 | 0.459 | 0.076 |
| **wICA_ICLabel** | 0.051 | 0.054 | 0.203 | 0.055 |

Table S3. Means and SDs for muscle related metrics.

#### *ICA Variance Explained by Neural Components*

There was a significant difference in the percentage of variance explained by neural activity between the pipelines with the robust ANOVA $F(2.38, 180.98) = 2403.582$, $p < 0.0001$. The rank order from best performing pipeline to worst performing pipeline of significant differences between individual cleaning pipelines from post-hoc t-tests was as follows: MWF_wICA_45Hz > MWF_wICA_infomax > MWF_wICA_fastICA, wICA_ICLabel > MWF_wICA_AMICA > MWF_only > wICA_all (note that pipelines using ICA subtraction were excluded from this metric) (Figure S18-19).

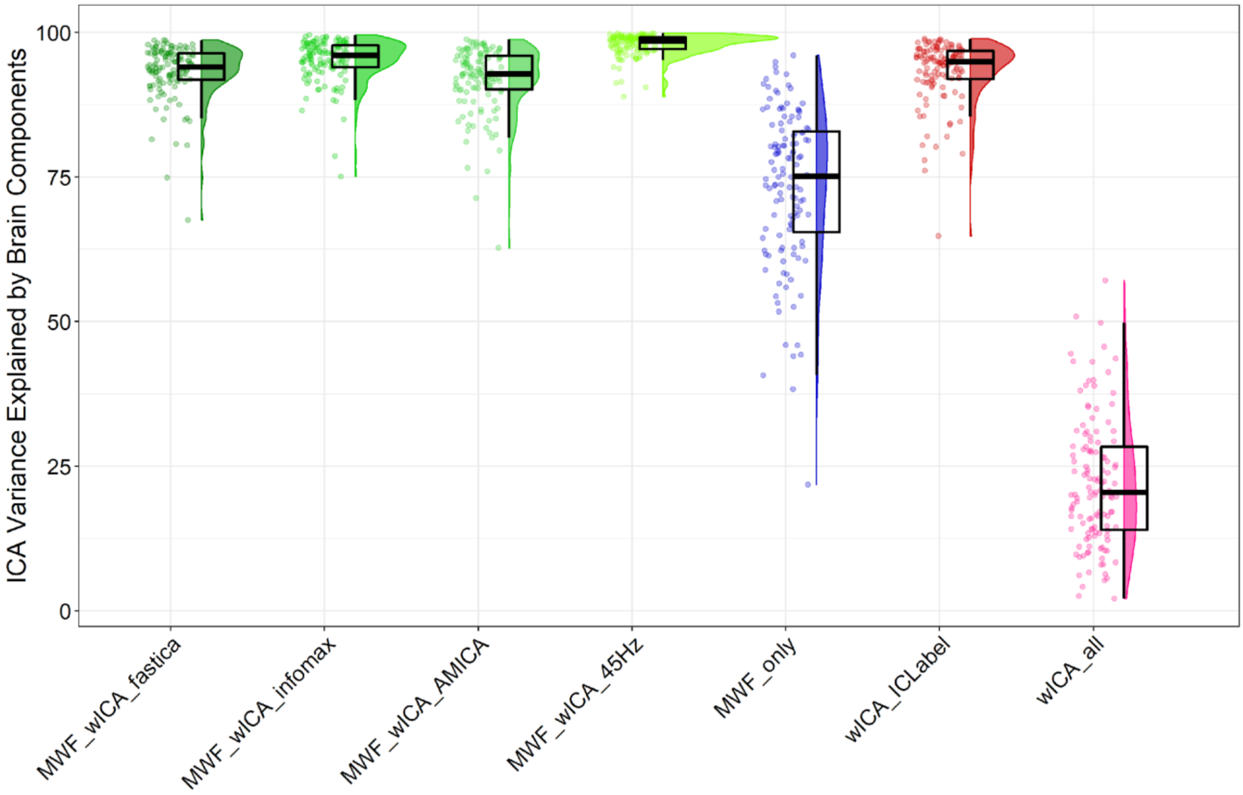

Figure S18. Raincloud plot depicting the amount of ICA variance explained by neural activity for each of the cleaning pipelines.

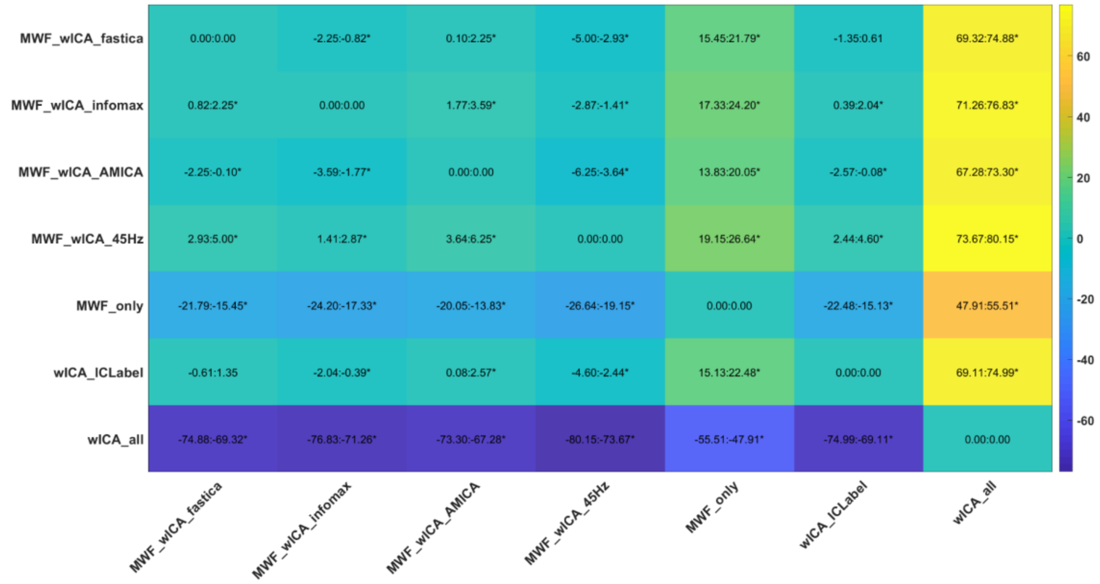

Figure S19. Post-hoc tests for variance explained by brain activity after cleaning detected by ICLabel.

| | Amount of variance explained by brain activity detected by ICLabel after cleaning | |
|---|---|---|
| **Pipeline** | **Mean** | **SD** |
| **MWF_wICA_AMICA** | 91.843 | 5.78 |
| **MWF_wICA_fastICA** | 93.265 | 4.688 |
| **MWF_wICA_infomax** | 95.148 | 3.767 |
| **MWF_wICA_45Hz** | 97.863 | 1.894 |
| **MWF_only** | 73.437 | 13.382 |
| **wICA_all** | 22.099 | 11.131 |
| **wICA_ICLabel** | 93.268 | 5.389 |

Table S4. Means and SDs for the variance explained by brain activity after cleaning detected by ICLabel.

#### *Proportion of EEG Epochs Deleted by the Cleaning Pipeline*

There was a significant difference between the pipelines in the proportion epochs in the data rejected by the cleaning process: $F(6.35, 482.9) = 54.0523$, $p < 0.0001$. The rank order from best performing pipeline to worst performing pipeline of significant differences between individual cleaning pipelines from post-hoc t-tests was as follows: wICA_all > MWF_wICA_fastICA^, MWF_wICA_45Hz[@], MWF_ICA_subtract[+], MWF_wICA_AMICA*,

MWF_wICA_infomax*, ICA_subtract^^, wICA_ICLabel^^++@@, MWF_wICA_CCA^^****++ > MWF_CCA, MWF_only (Figure S20-22).

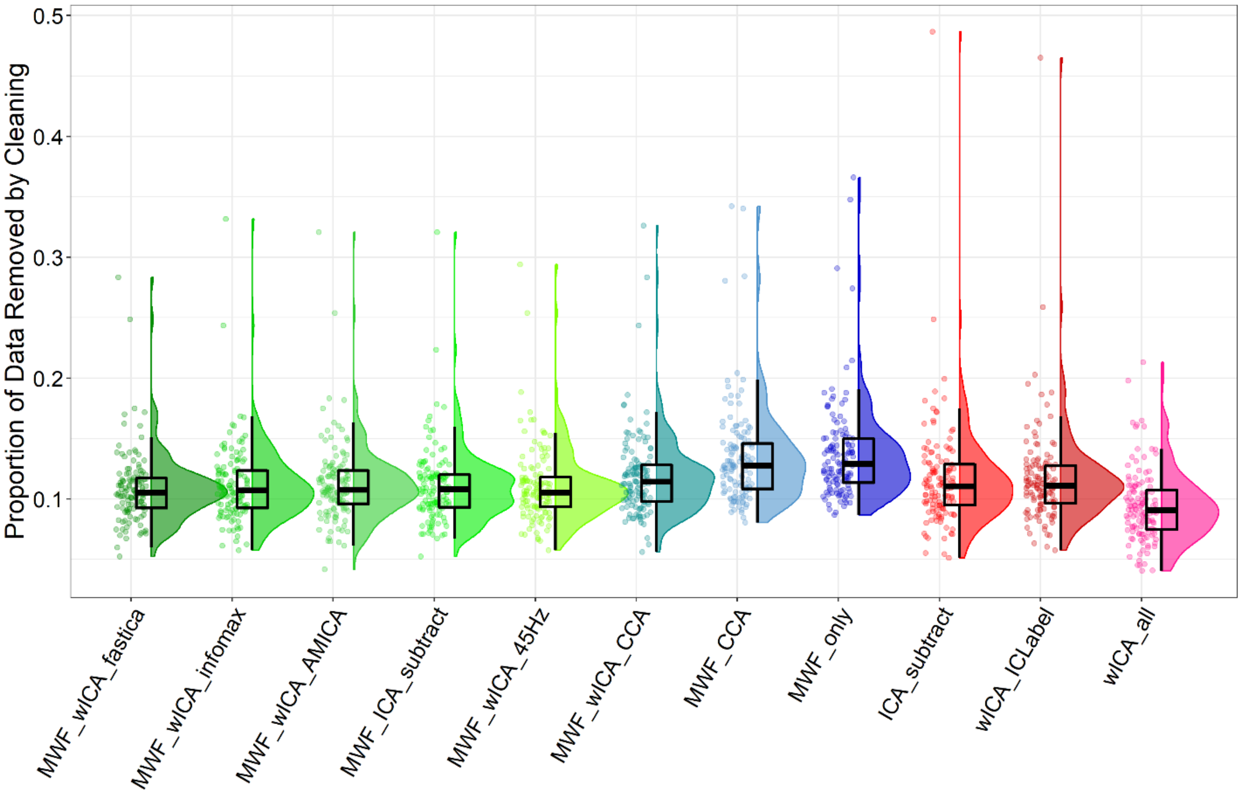

Figure S20. Raincloud plot depicting the proportion of epochs in the data removed by the cleaning process (N = 127) for each of the cleaning pipelines.

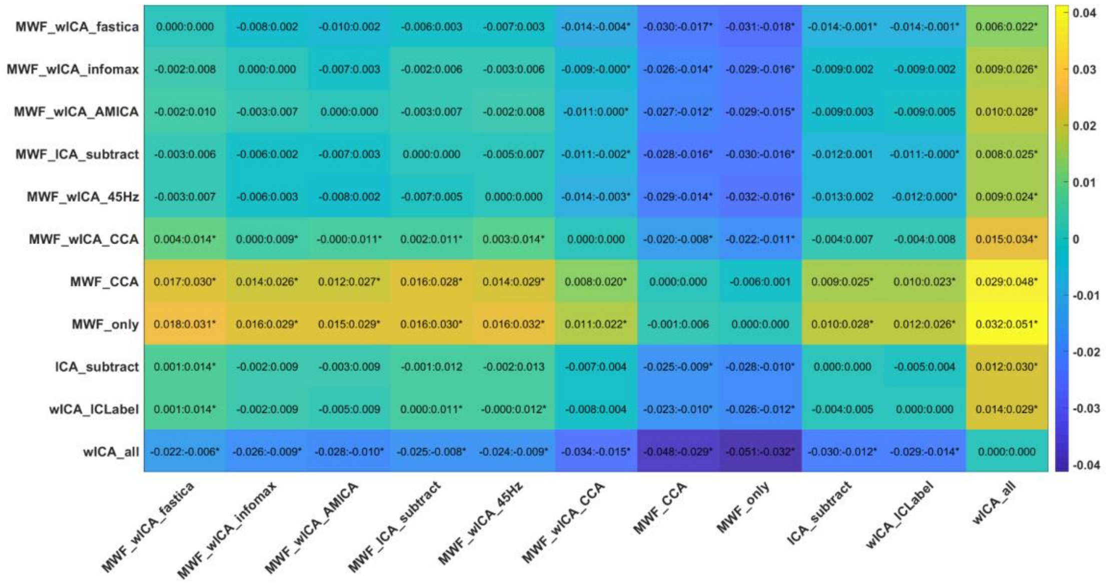

Figure S21. Post-hoc tests for the proportion of epochs deleted by the cleaning pipelines.

| | Proportion of data removed by the pipeline | |
|---|---|---|
| **Pipeline** | **Mean** | **SD** |
| **ICA_subtract** | 0.113 | 0.029 |
| **MWF_CCA** | 0.132 | 0.03 |
| **MWF_wICA_CCA** | 0.117 | 0.026 |
| **MWF_wICA_AMICA** | 0.11 | 0.025 |
| **MWF_wICA_fastICA** | 0.107 | 0.022 |
| **MWF_ICA_subtract** | 0.109 | 0.024 |
| **MWF_wICA_infomax** | 0.11 | 0.024 |
| **MWF_wICA_45Hz** | 0.109 | 0.023 |
| **MWF_only** | 0.134 | 0.029 |
| **wICA_all** | 0.093 | 0.027 |
| **wICA_ICLabel** | 0.114 | 0.026 |

Table S5. Means and SDs for the proportion of epochs removed by the cleaning pipeline.

#### *ERP Condition Comparisons - Variance Explained Metrics*

##### *Variance Explained by Error vs Correct Responses*

Here we present the post-hoc test of the interaction between each pair of pipelines and correct/error condition for the ERN TANOVA (averaged activity between 0 and 150ms) and Pe GFP (averaged activity between 150 and 300ms) and Pe TANOVA (averaged activity between 200 and 400ms). A bar graph of the explained variance for each pipeline and ERP is presented in the main manuscript, along with the rank order of strength of variance explained across the pipelines.

To save computation time, we did not include MWF_wICA_45Hz in the comparisons of variance explained by the difference between correct and error responses, as this comparison was performed last, and no other metric indicated that MWF_wICA_45Hz was the optimal approach. Statistical comparisons of the overall interaction between pipelines and condition were highly significant for all three measures (ERN TANOVA, Pe GFP, and Pe TANOVA, all $p < 0.001$). With regards to the ERN TANOVA all pipelines provided $np^2$ values between 0.26 and 0.31 except for wICA_all which provided $np^2 = 0.23$. Post-hoc testing of the interaction between each pair of pipelines and the two conditions indicated the following rank order from best performing pipeline to worst performing pipeline of the ability of the pipelines to discriminate between the experimental manipulation: wICA_ICLabel*, MWF_only*, MWF_CCA*, MWF_wICA_infomax*, ICA_subtract*, MWF_ICA_subtract*, MWF_wICA_CCA*, MWF_wICA_fastICA, MWF_wICA_AMICA*, wICA_all**. Results can be viewed in Figures S22-25.

With regards to the Pe GFP, all pipelines provided np$^{2}$ values between 0.45 and 0.55, except for wICA_all which provided np$^{2}$ = 0.29. Post-hoc testing of the interaction between each pair of pipelines and the two conditions indicated the following rank order from best performing pipeline to worst performing pipeline of the ability of the pipelines to discriminate between the experimental manipulation: MWF_only*, MWF_CCA@, wICA_ICLabel^**, MWF_wICA_AMICA^^, MWF_wICA_fastICA^^, MWF_wICA_infomax^^+, MWF_wICA_CCA**^^@@, ICA_subtract**^^++@@, MWF_ICA_subtract**^^@@ > wICA_all. With regards to the Pe TANOVA, all pipelines provided np$^{2}$ values between 0.15 and 0.20. Post-hoc testing of the interaction between each pair of pipelines and the two conditions indicating the following rank order of the ability of the pipelines to discriminate between the experimental manipulation: wICA_ICLabel*^, MWF_wICA_infomax**^, MWF_wICA_CCA^, MWF_ICA_subtract**^, ICA_subtract**^, MWF_wICA_AMICA**^, MWF_only**^, MWF_CCA**, MWF_wICA_fastICA**^, wICA_all**^^ . Visual inspection of the topoplots indicated all pipelines showed similar patterns for the ERN and for the Pe, with no obvious indication of the reason for the differences in explained variance between the pipelines (Figure S26-31).

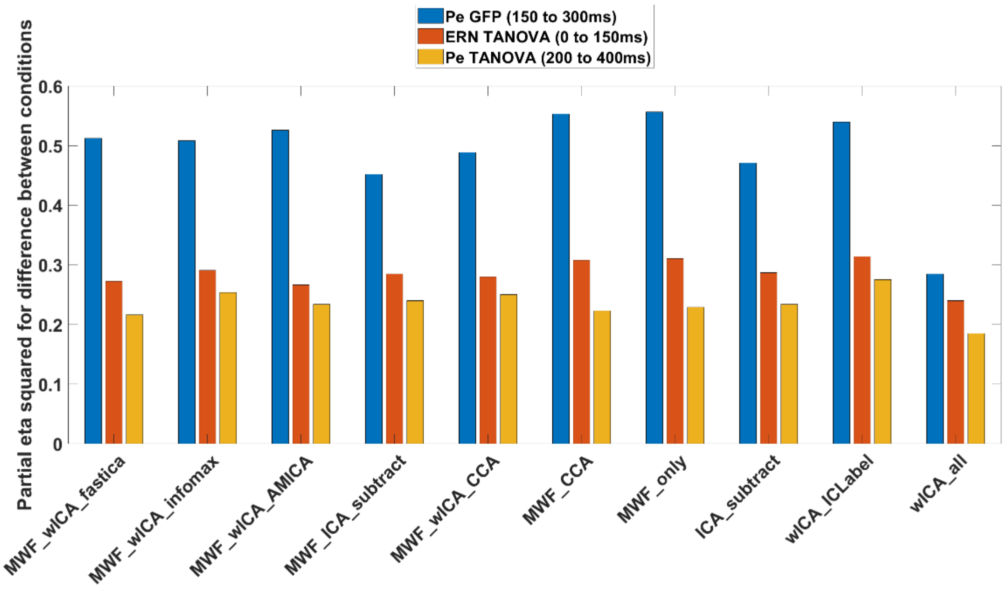

Figure S22. The variance explained by the difference between error and correct trials in the distribution (using the TANOVA) of the ERN (0 to 150ms after a response) and the Pe (200 to 400ms after a response), as well as the GFP of the Pe (150 to 300ms after a response) for each of the cleaning pipelines.

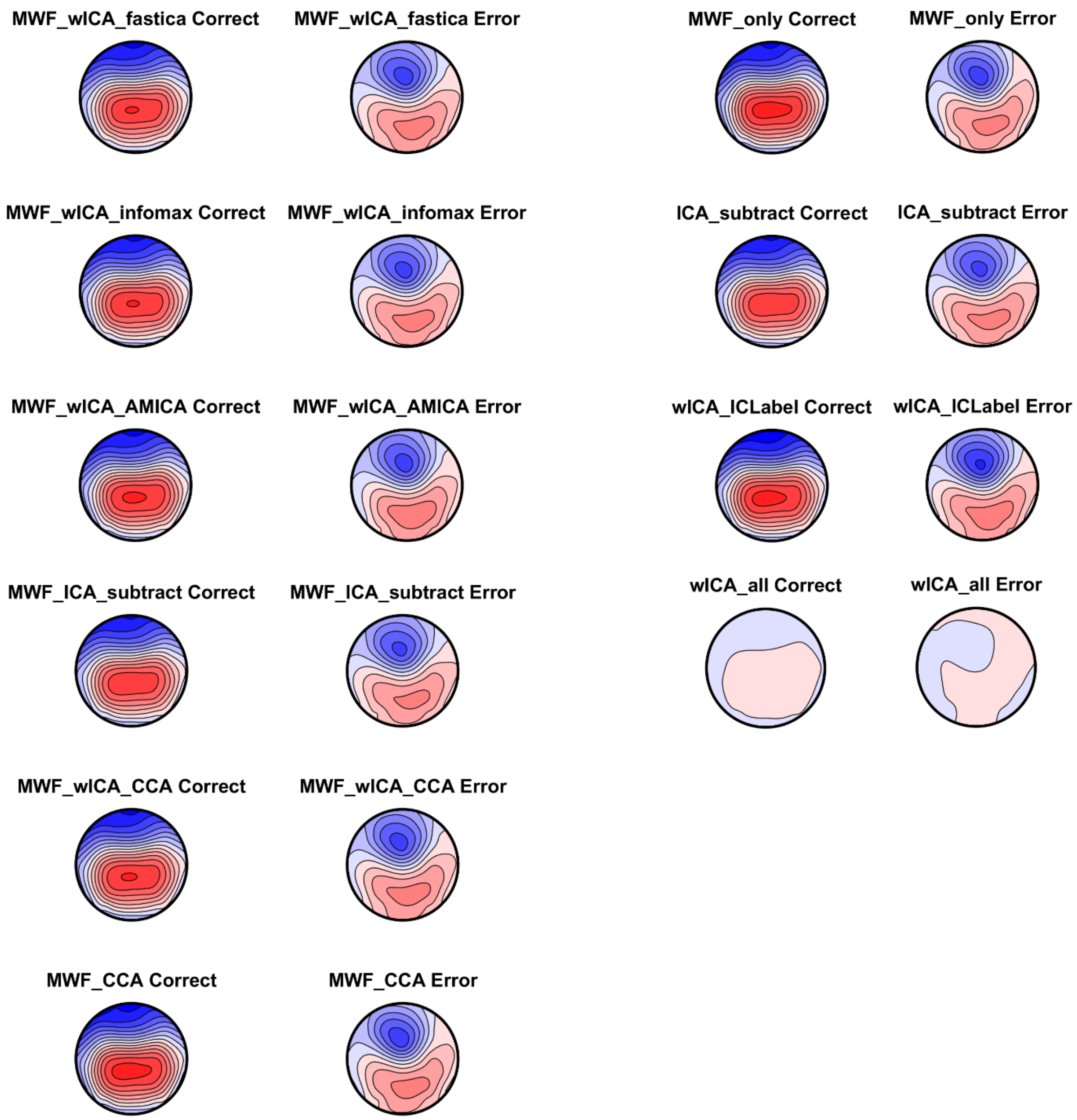

Figure S23. Topoplots of the averaged ERN window for correct and error responses for each pipeline. All plots are on the same scale so comparisons can be made across pipelines / conditions. Note the similarity across the majority of pipelines, with only wICA_all displaying much lower voltage amplitudes across the entire scalp.

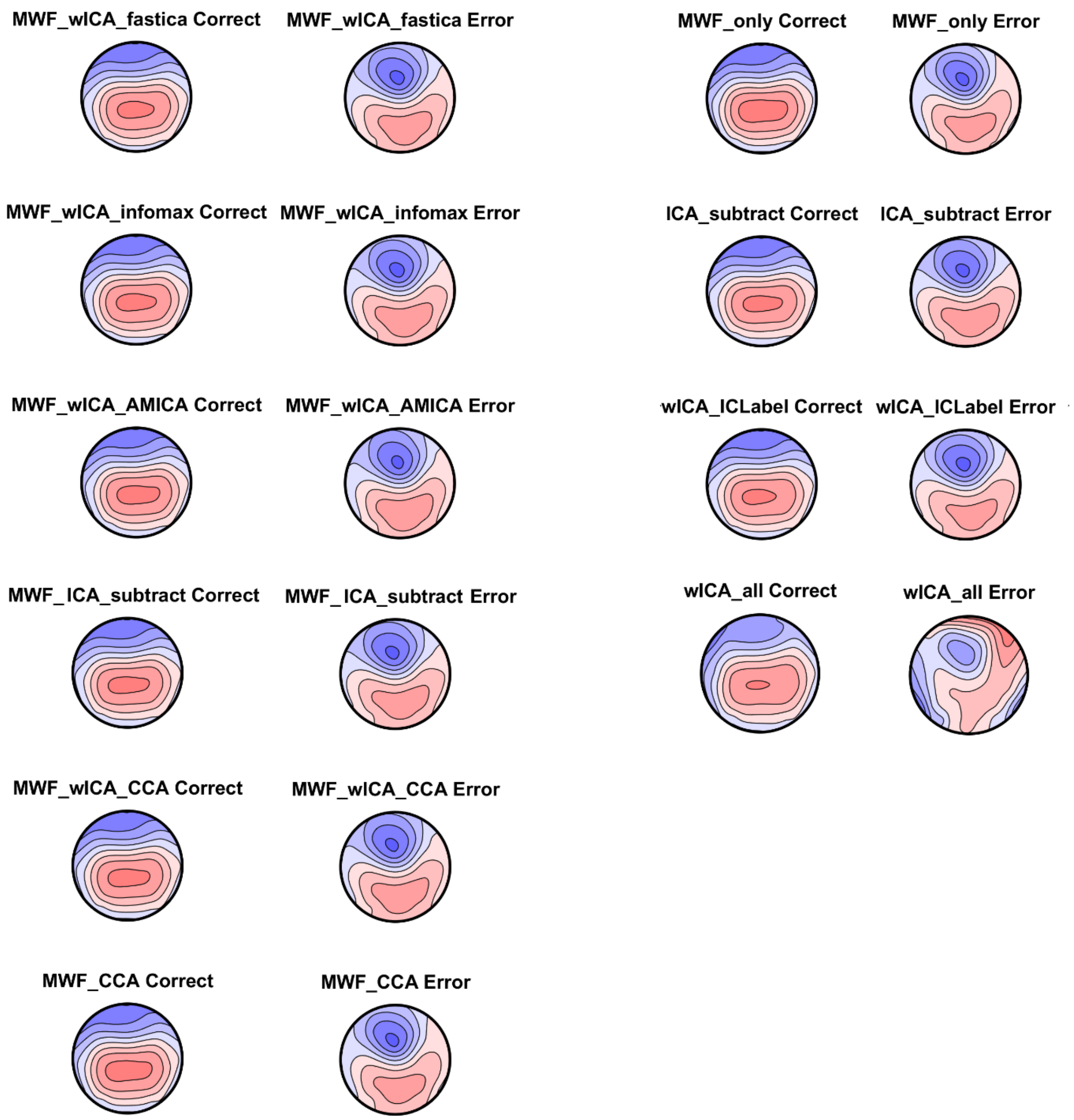

Figure S24. Topoplots of the averaged ERN window for correct and error responses for each pipeline. All plots are on their own scale so the distribution within each pipeline / condition can be viewed. Note the similarity across the majority of pipelines, with only wICA_all showing a different pattern of results.

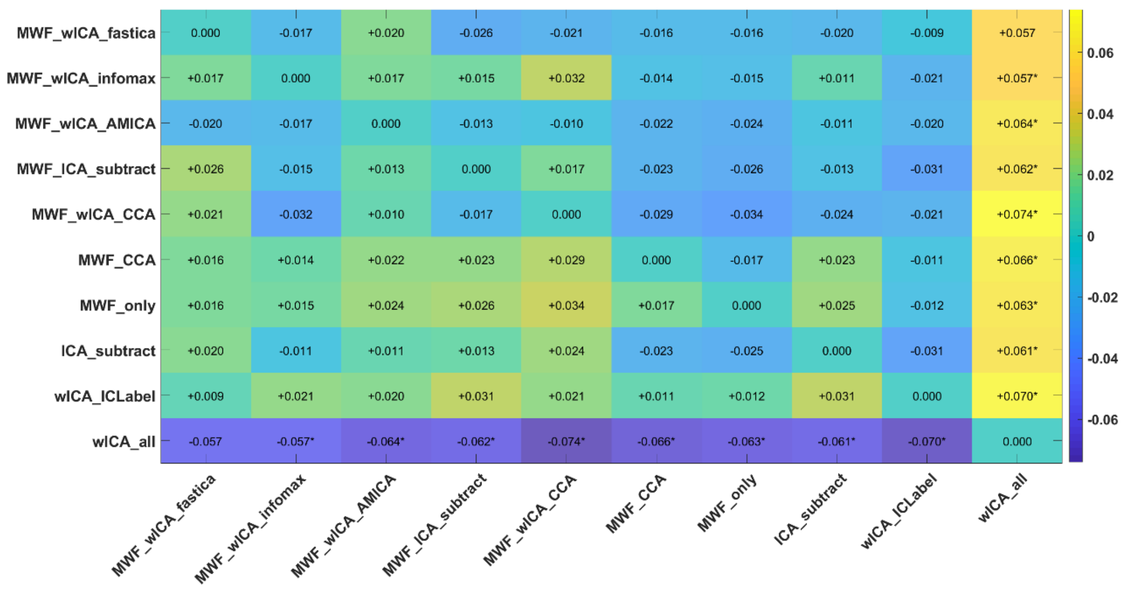

Figure S25. Heat map of the variance explained (np²) by the interaction between each pair of pipelines and averaged activity within the ERN window TANOVA test for correct vs error responses. Interactions that were significant (FDR-p < 0.05) are indicated with an *. We have also provided an indication of which pipeline of each pair provided larger values for variance explained using – and + symbols, which can be interpreted as the pipeline listed on the left of the heatmap having shown less (-) or more (+) variance explained in the comparison between the two experimental conditions than the pipeline listed at the bottom of the heatmap.

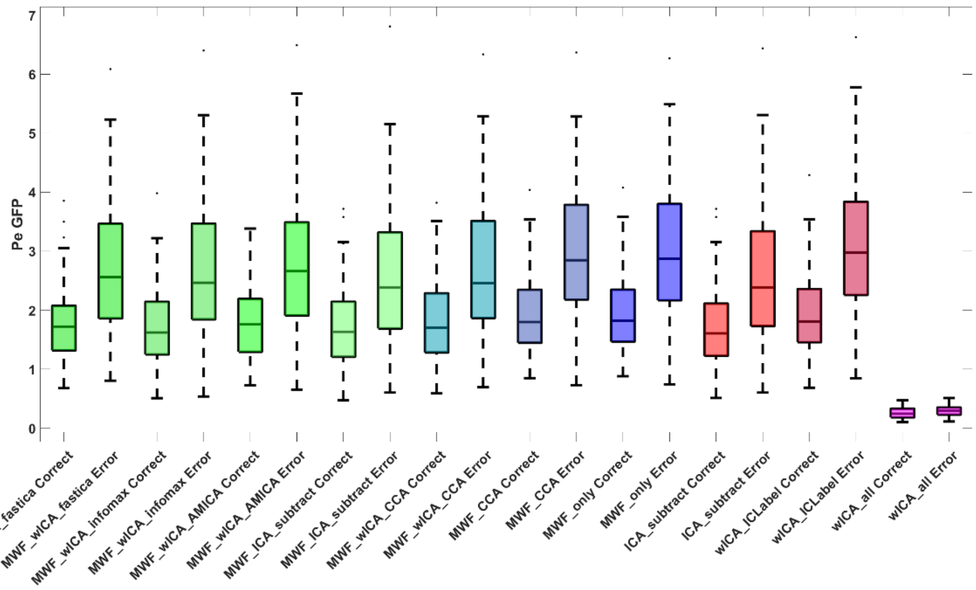

Figure S26. GFP for the Pe following correct responses and error responses for each of the cleaning pipelines.

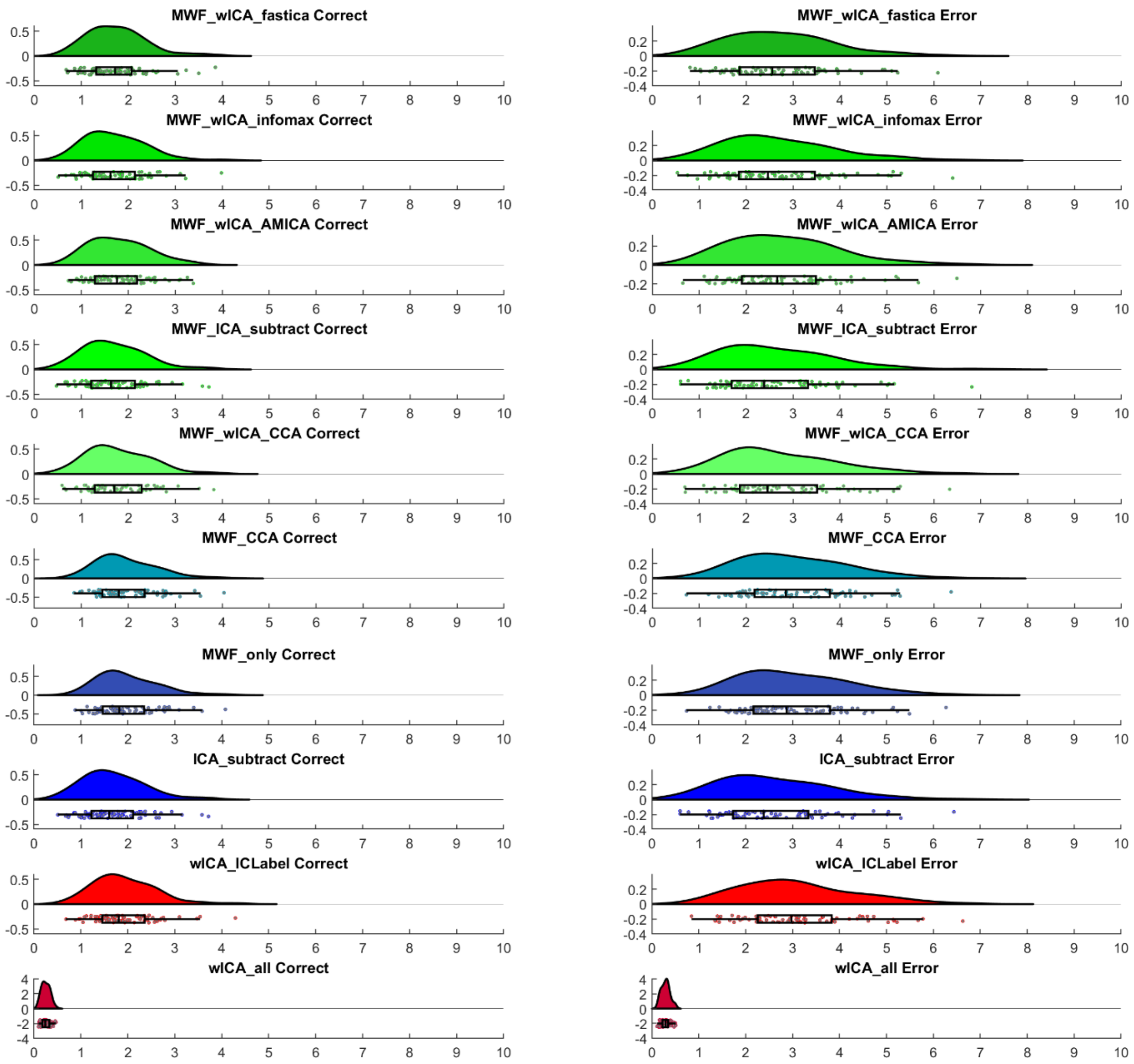

S27. Raincloud plots for the GFP of the Pe following correct and error responses for all cleaning pipelines.

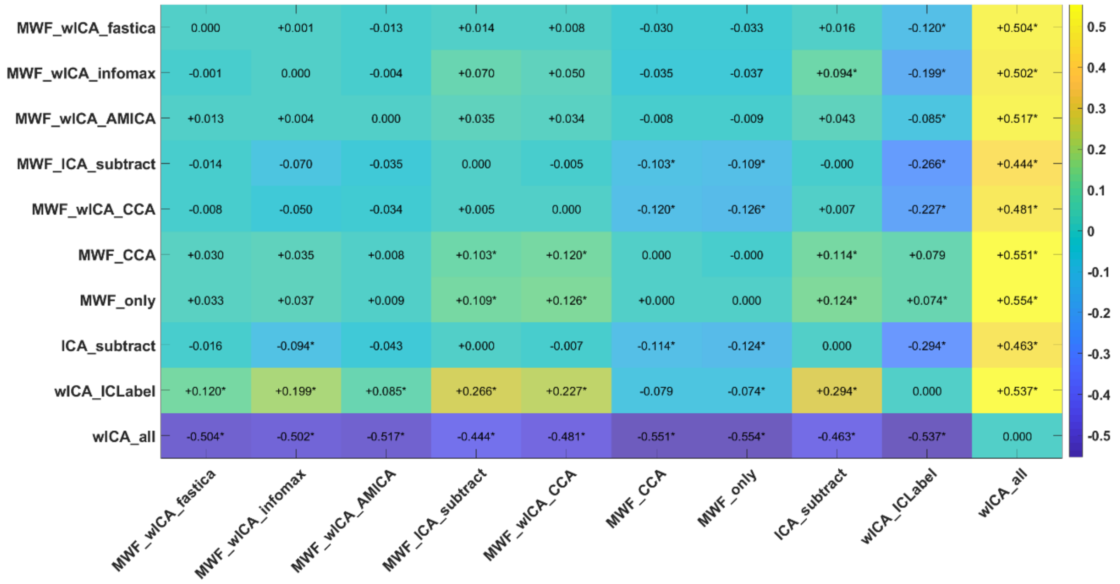

Figure S28. Heat map of the variance explained ($np^2$) by the interaction between each pair of pipelines and averaged activity within the Pe window GFP test for correct vs error responses. Interactions that were significant (FDR-p < 0.05) are indicated with an *. We have also provided an indication of which pipeline of each pair provided larger values for variance explained using – and + symbols, which can be interpreted as the pipeline listed on the left of the heatmap having shown less (-) or more (+) variance explained in the comparison between the two experimental conditions than the pipeline listed at the bottom of the heatmap.

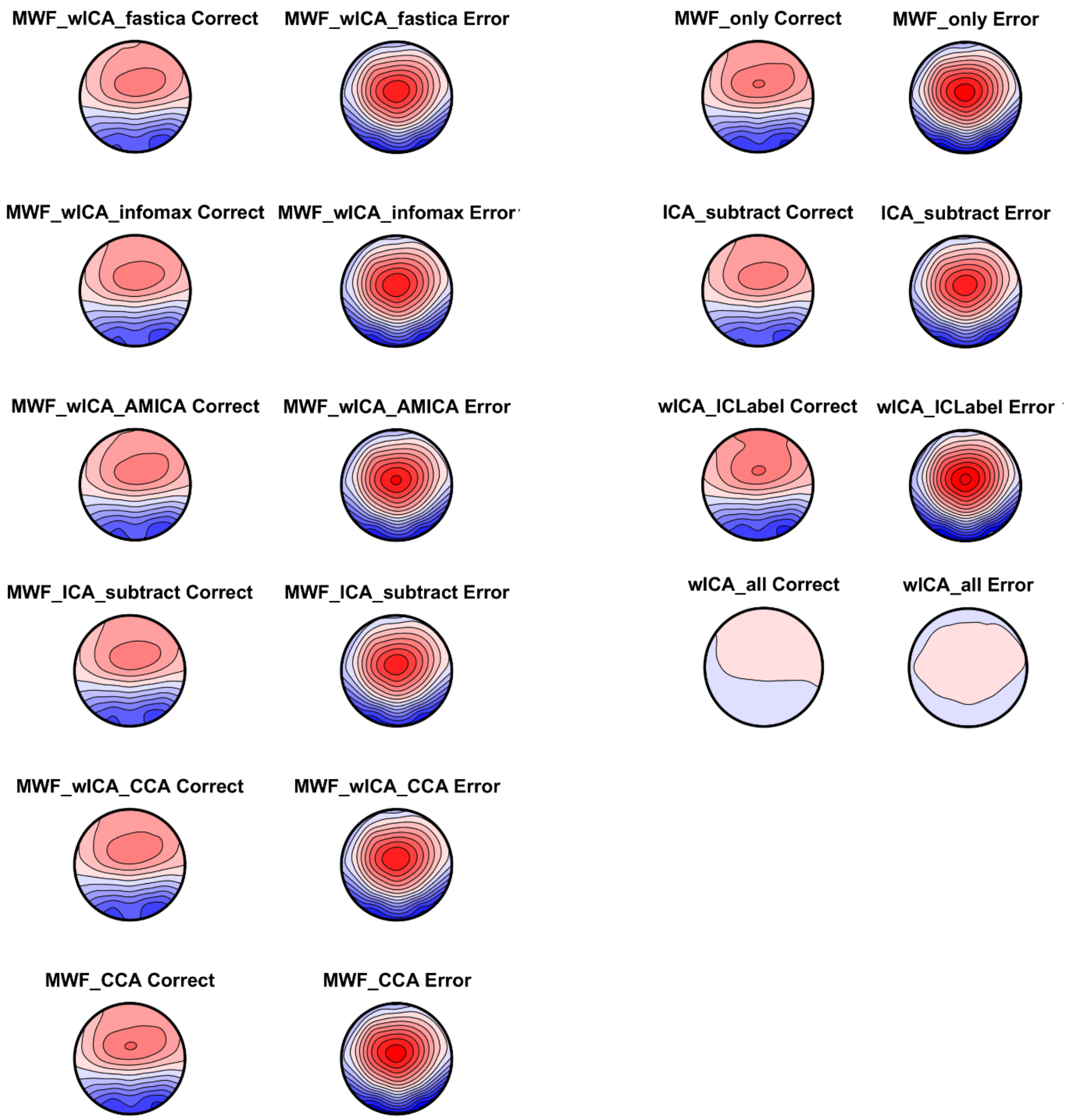

Figure S29. Topoplots of the averaged Pe window for correct and error responses for each pipeline. All plots on the same scale so the distribution within each pipeline / condition can be compared across pipelines / conditions. Note the similarity across the majority of pipelines, with only wICA_all showing a slightly different pattern of results.

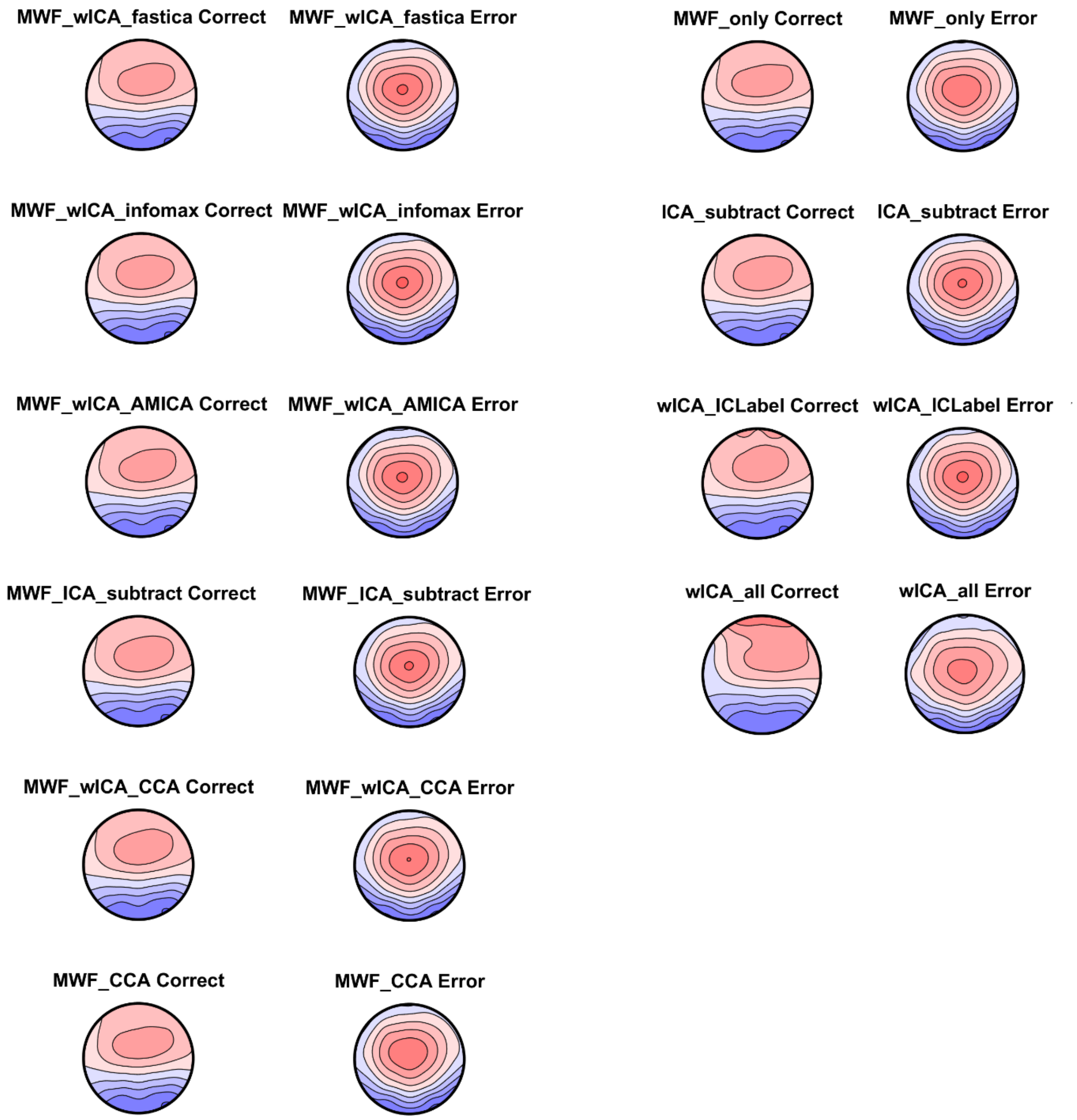

Figure S30. Topoplots of the averaged Pe window for correct and error responses for each pipeline. All plots are on their own scale so the distribution within each pipeline / condition can be viewed. Note the similarity across the majority of pipelines, with only wICA_all showing a slightly different pattern of results.

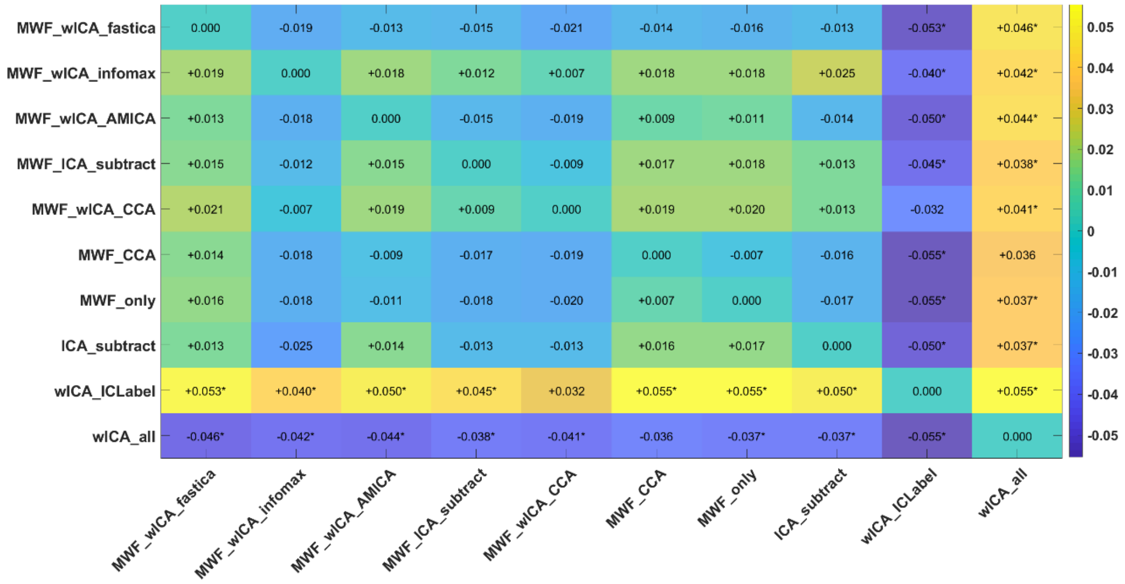

Figure S31. Heat map of the variance explained (np$^2$) by the interaction between each pair of pipelines and averaged activity within the Pe window TANOVA test for correct vs error responses. Interactions that were significant (FDR-p < 0.05) are indicated with an *. We have also provided an indication of which pipeline of each pair provided larger values for variance explained using – and + symbols, which can be interpreted as the pipeline listed on the left of the heatmap having shown less (-) or more (+) variance explained in the comparison between the two experimental conditions than the pipeline listed at the bottom of the heatmap.

#### *Variance Explained by Go vs Nogo Trials*

In addition to the amount of variance explained by the difference between response locked error and correct responses, we examined the amount of variance explained by the difference between stimulus locked Go and Nogo trials, focusing on the N2 overall neural response strength (GFP test) and distribution (TANOVA) separately, and the P3 overall neural response strength (GFP test) and distribution (TANOVA) separately (Figure S32). Here we present the post-hoc test of the interaction between each pair of pipelines and Go/Nogo trial condition for the N2 (averaged activity between 180 and 300ms) and P3 (averaged activity between 300 and 500ms) GFP and TANOVA.

Statistical comparisons of the overall interaction between pipelines and condition were highly significant for all four measures (all p < 0.001). With regards to the N2 GFP most pipelines provided np$^2$ values from 0.28 to 0.4, with post-hoc testing of the interaction between each pair of pipelines and the two conditions indicating the following rank order of the ability of the pipelines to discriminate between the experimental manipulation: wICA_ICLabel, ICA_subtract > MWF_CCA, MWF_only, MWF_wICA_fastICA, MWF_wICA_AMICA, MWF_wICA_CCA, MWF_wICA_infomax, MWF_wICA_45Hz, MWF_ICA_subtract > wICA_all (Figure S33-35). With regards to the N2 TANOVA all pipelines provided np$^2$ values between 0.15 to 0.23, with post-hoc testing of the interaction between each pair of pipelines and the two conditions indicating the following rank order of the ability of the pipelines to discriminate between the experimental manipulation: wICA_all > MWF_CCA*, MWF_only*,

MWF_wICA_AMICA**^, wICA_ICLabel$^{+}$, MWF_wICA_fastICA**, MWF_ICA_subtract**$^{\$}$, MWF_wICA_infomax**^^$^{\$}$, MWF_wICA_45Hz**^^$^{\$\$}$, MWF_wICA_CCA**$^{\$\$}$, ICA_subtract**$^{++}$. Visual inspection of the topoplots indicated all pipelines showed a similar pattern, with no obvious indication of the reason for the differences in explained variance between the pipelines (Figure S36-38). It is also worth noting that all of the methods that combined MWF and wICA showed lower performance than MWF alone. As mentioned in other sections, while wICA_all seemed to perform the best for this metric, it produced EEG data with very small amplitudes, suggesting the pipeline is likely to have cleaned neural activity as well as artifacts (even if it did preserve neural activity that provided high power to discern between the N2 distribution of Go and Nogo responses).

With regards to the P3 GFP most pipelines provided $np^2$ values from 0.05 to 0.13 except for wICA_all, which provided a value of 0.30. Post-hoc testing of the interaction between each pair of pipelines and the two conditions indicated the following rank order of the ability of the pipelines to discriminate between the experimental manipulation: wICA_all*, MWF_only**^, MWF_CCA**^, MWF_wICA_AMICA**$^{\$}$, MWF_wICA_CCA**^^, MWF_wICA_infomax**^^, MWF_ICA_subtract**, MWF_wICA_fastICA**^^$^{\$\$}$, MWF_wICA_45Hz**^^$^{\$\$}$, wICA_ICLabel**^^, ICA_subtract^^$^{\$\$}$. (Figure S39-41). Lastly, with regards to the P3 TANOVA, most pipelines provided $np^2$ values from 0.46 to 0.50, except for wICA_all which provided $np^2 = 0.20$. Post-hoc testing of the interaction between each pair of pipelines and the two conditions indicated the following rank order of the ability of the pipelines to discriminate between the experimental manipulation: ICA_subtract*, MWF_wICA_fastICA*, wICA_ICLabel$^{@}$, MWF_wICA_infomax*^, MWF_wICA_AMICA*^, MWF_wICA_45Hz*^, MWF_wICA_CCA*^, MWF_ICA_subtract*^, MWF_CCA**^^, MWF_only**$^{@@}$ > wICA_all (S42-44).

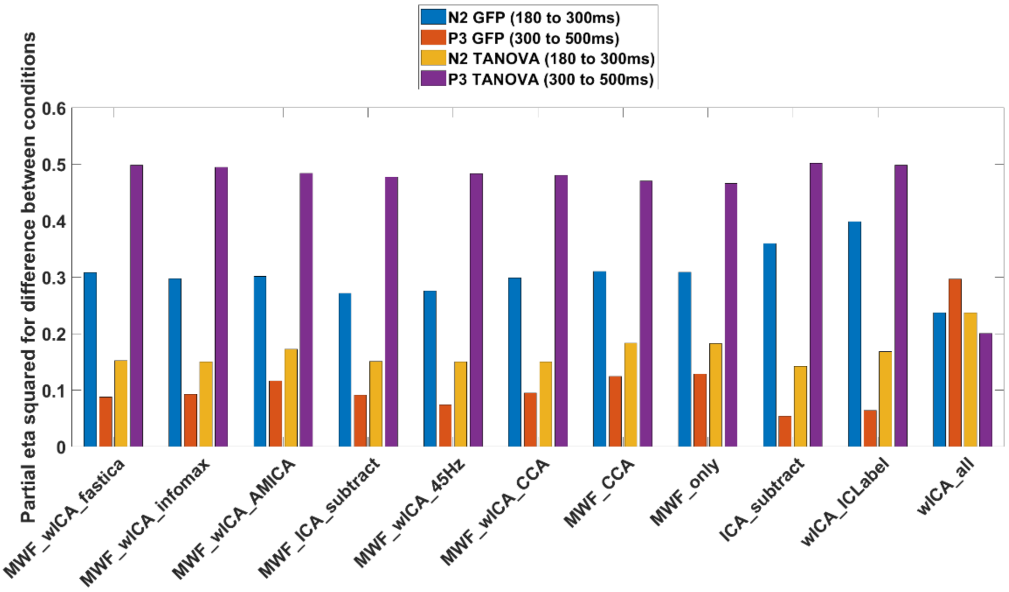

Figure S32. The variance explained ($np^2$) by the difference between Go and Nogo trials in the Go-Nogo dataset for the N2 (180 to 300ms) and P3 (300 to 500ms) GFP and TANOVA for each pipeline.

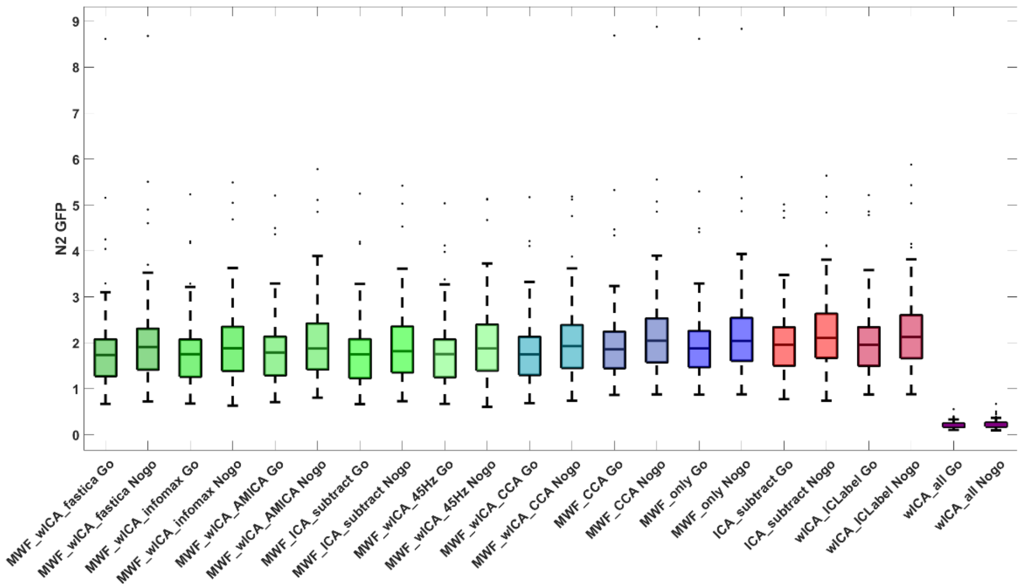

Figure S33. Box plots of the Go and Nogo N2 GFP from each of the cleaning pipelines.

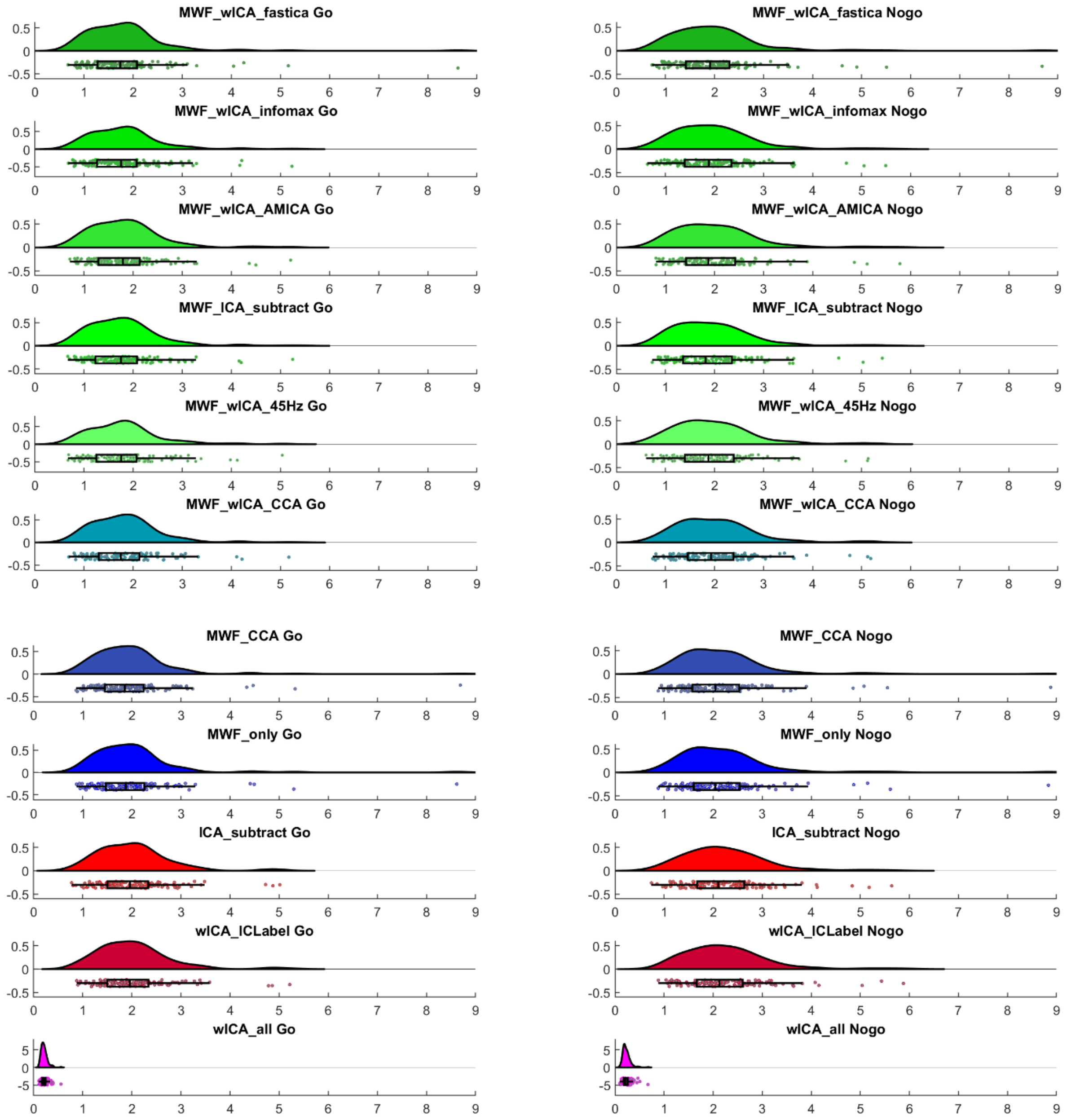

Figure S34. Raincloud plots for the N2 amplitudes from Go and Nogo trials for each of the cleaning pipelines.

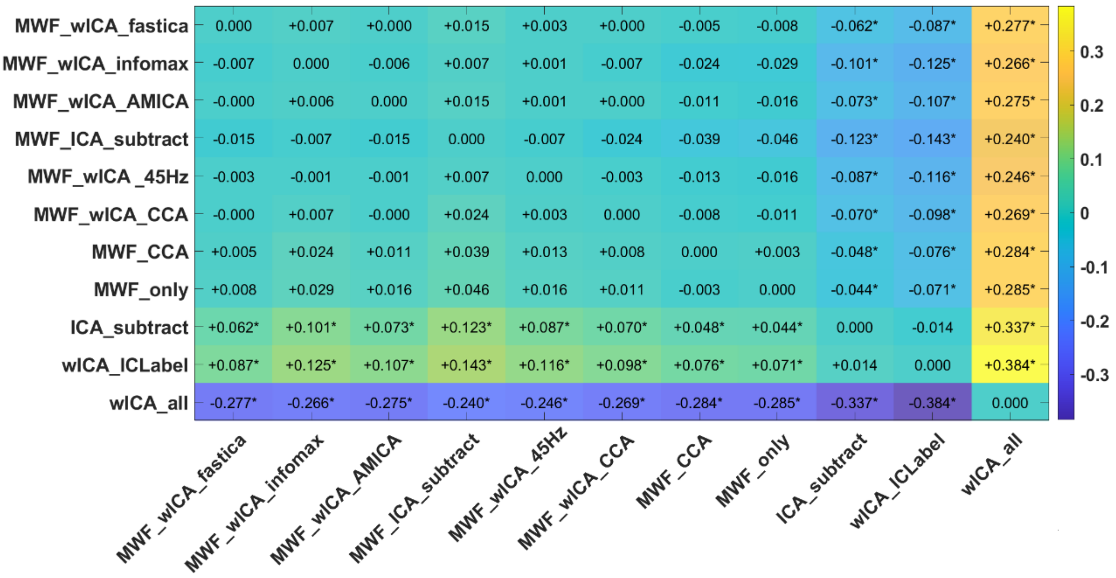

Figure S35. Heat map of the variance explained ($np^2$) by the interaction between each pair of pipelines and averaged activity within the N2 window GFP test for Go vs Nogo trials. Interactions that were significant (FDR-p < 0.05) are indicated with an *. We have also provided an indication of which pipeline of each pair provided larger values for variance explained using – and + symbols, which can be interpreted as the pipeline listed on the left of the heatmap having shown less (-) or more (+) variance explained in the comparison between the two experimental conditions than the pipeline listed at the bottom of the heatmap.

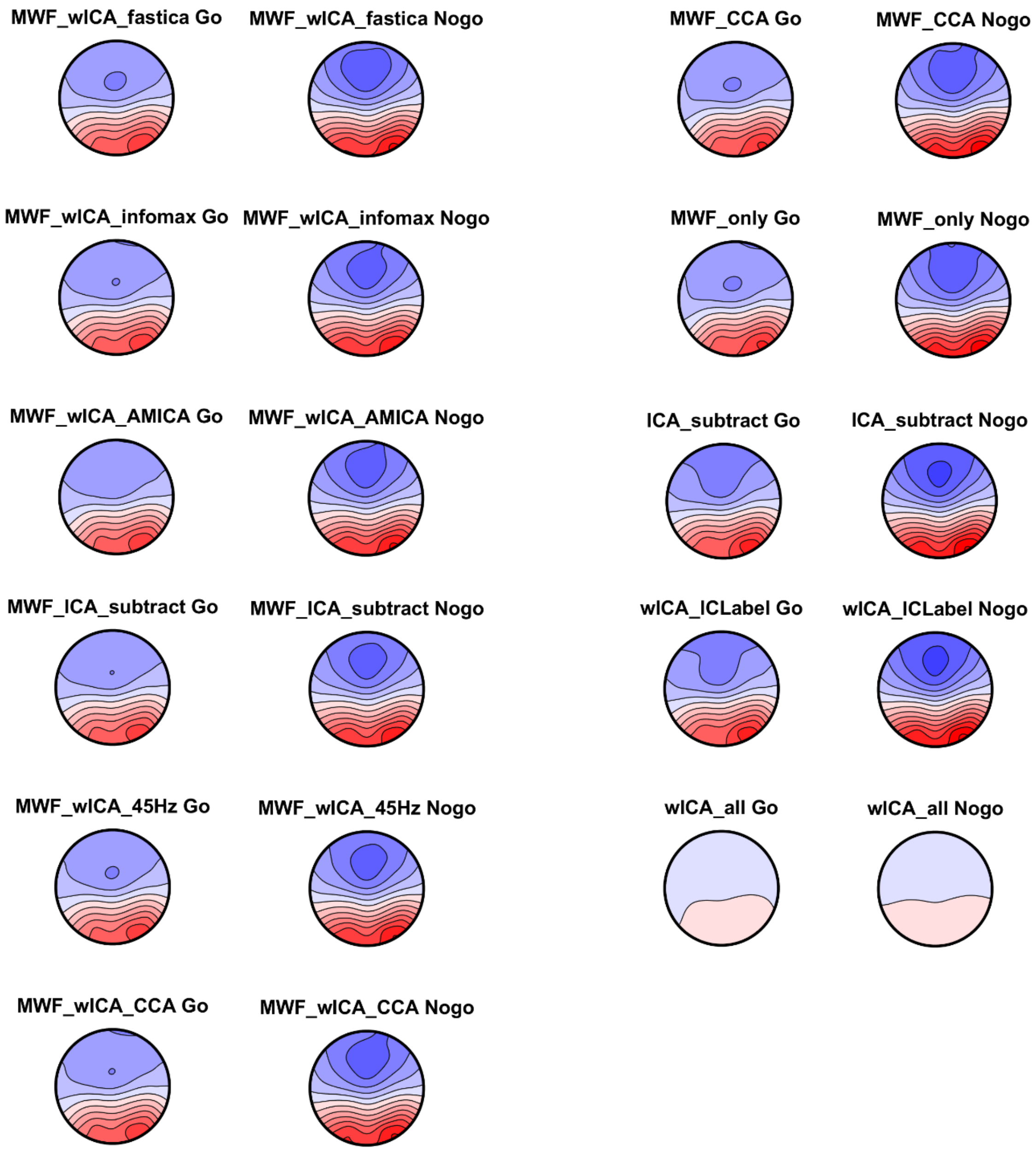

Figure S36. N2 distribution topoplots for Go and Nogo trials from each pipeline. All topoplots are depicted on the same scale so comparison is possible between each pipeline and condition. Note that all pipelines showed similar distributions and amplitudes with the exception of wICA_all which showed a much-reduced amplitude.

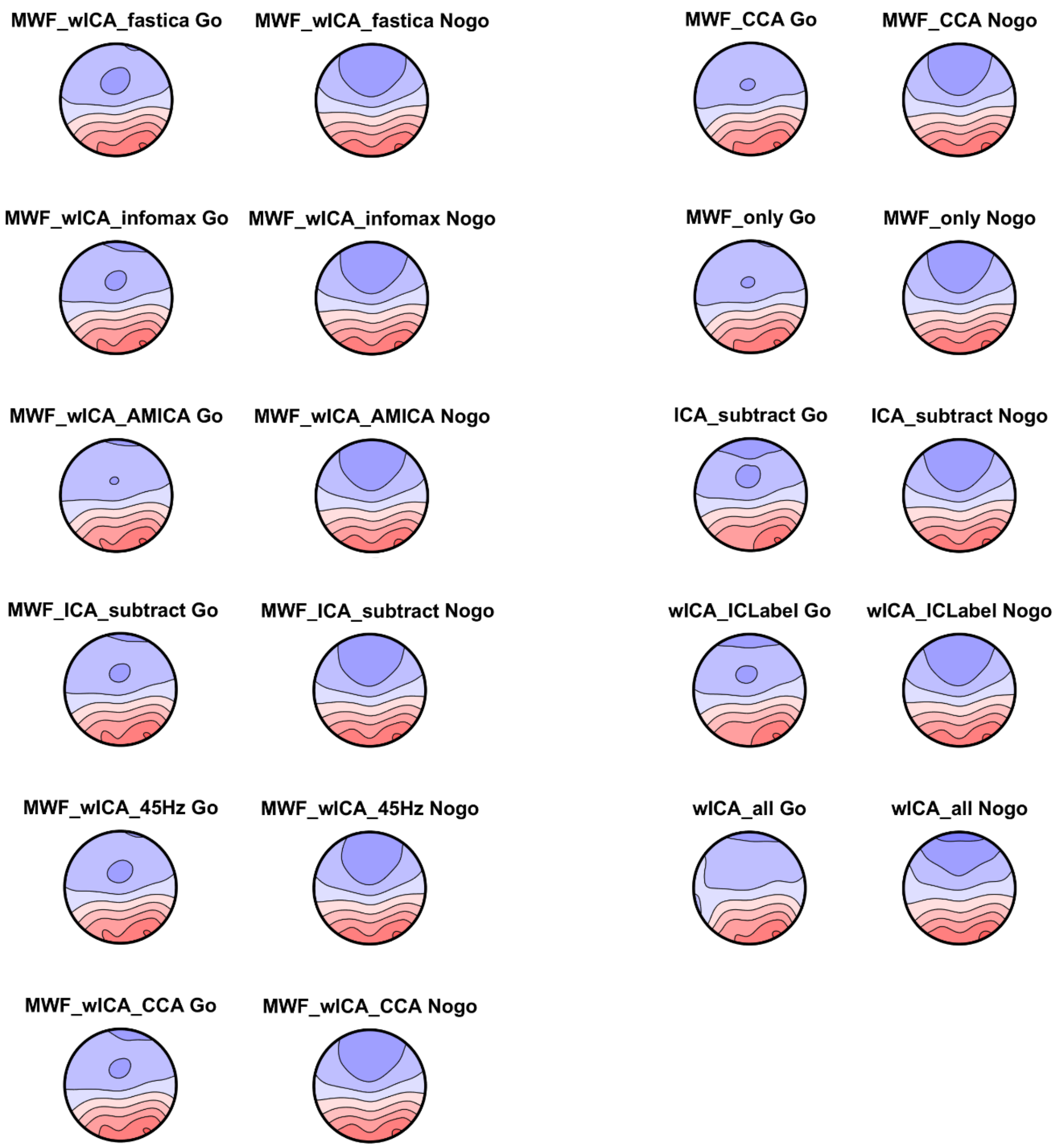

Figure S37. N2 distribution topoplots for Go and Nogo trials from each pipeline. All topoplots are depicted on their own scale so the distribution of activity is easy to understand (but comparison of amplitudes is not possible between conditions and pipelines).

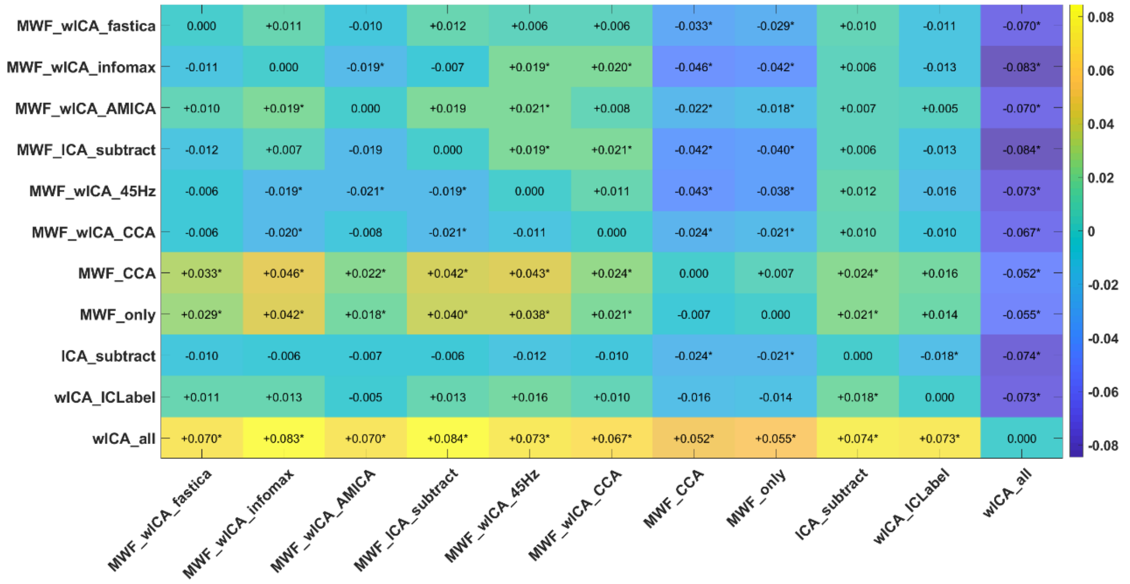

Figure S38. Heat map of the variance explained ($np^2$) by the interaction between each pair of pipelines and averaged activity within the N2 window TANOVA for Go vs Nogo trials. Interactions that were significant (FDR-p < 0.05) are indicated with an *. We have also provided an indication of which pipeline of each pair provided larger values for variance explained using – and + symbols, which can be interpreted as the pipeline listed on the left of the heatmap having shown less (-) or more (+) variance explained in the comparison between the two experimental conditions than the pipeline listed at the bottom of the heatmap.

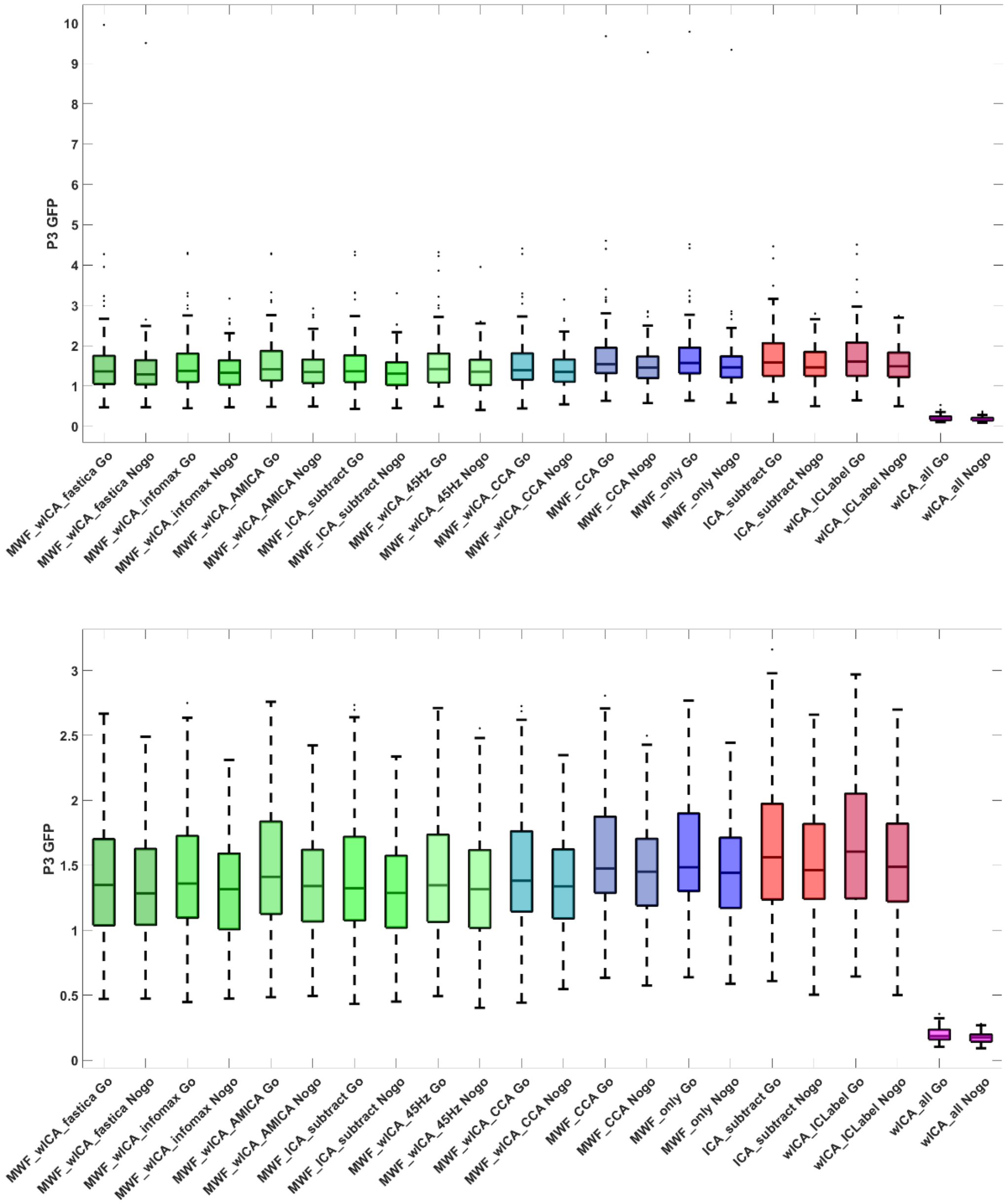

Figure S39. P3 GFP from Go and Nogo trials from each pipeline, with outliers (above) and without outliers (below) for increased discernability of the differences between pipelines.

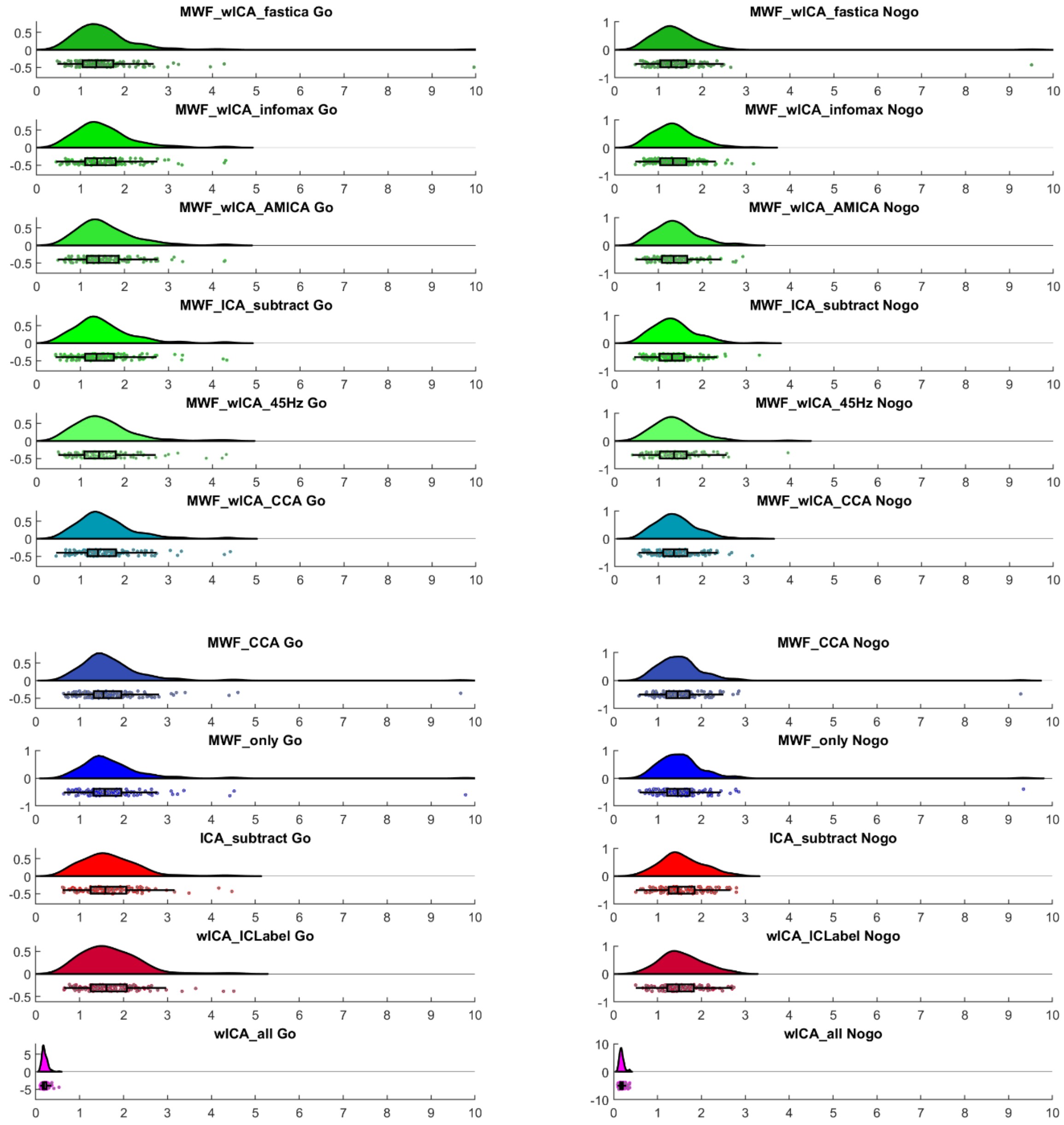

Figure S40. Raincloud plots of the P3 GFP for Go and Nogo trials from each pipeline.

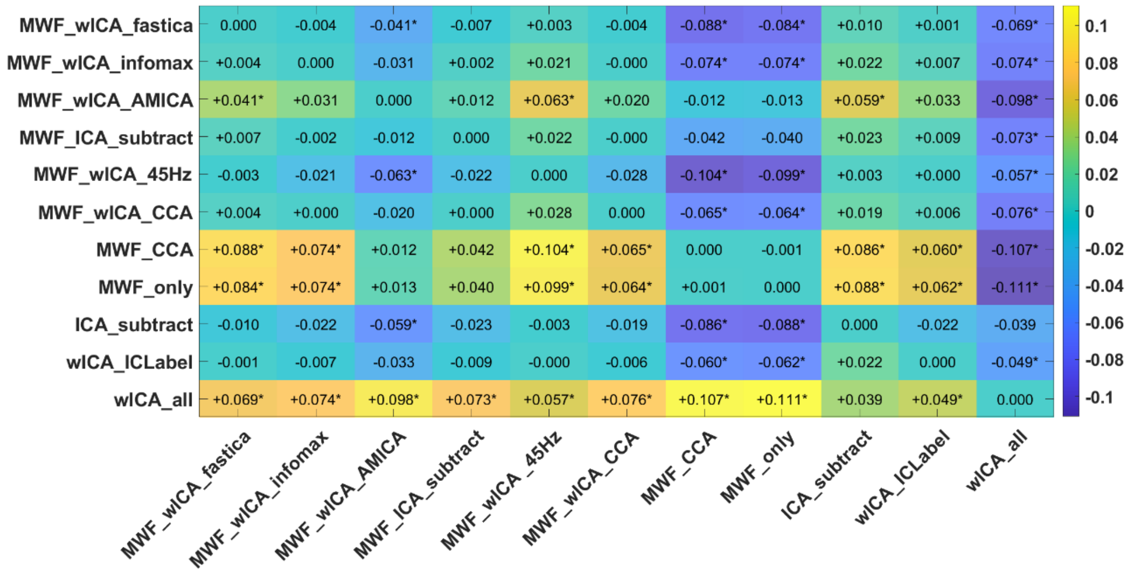

Figure S41. Heat map of the variance explained ($np^2$) by the interaction between each pair of pipelines and averaged activity within the P3 window GFP for Go vs Nogo trials. Interactions that were significant (FDR-p < 0.05) are indicated with an *. We have also provided an indication of which pipeline of each pair provided larger values for variance explained using – and + symbols, which can be interpreted as the pipeline listed on the left of the heatmap having shown less (-) or more (+) variance explained in the comparison between the two experimental conditions than the pipeline listed at the bottom of the heatmap.

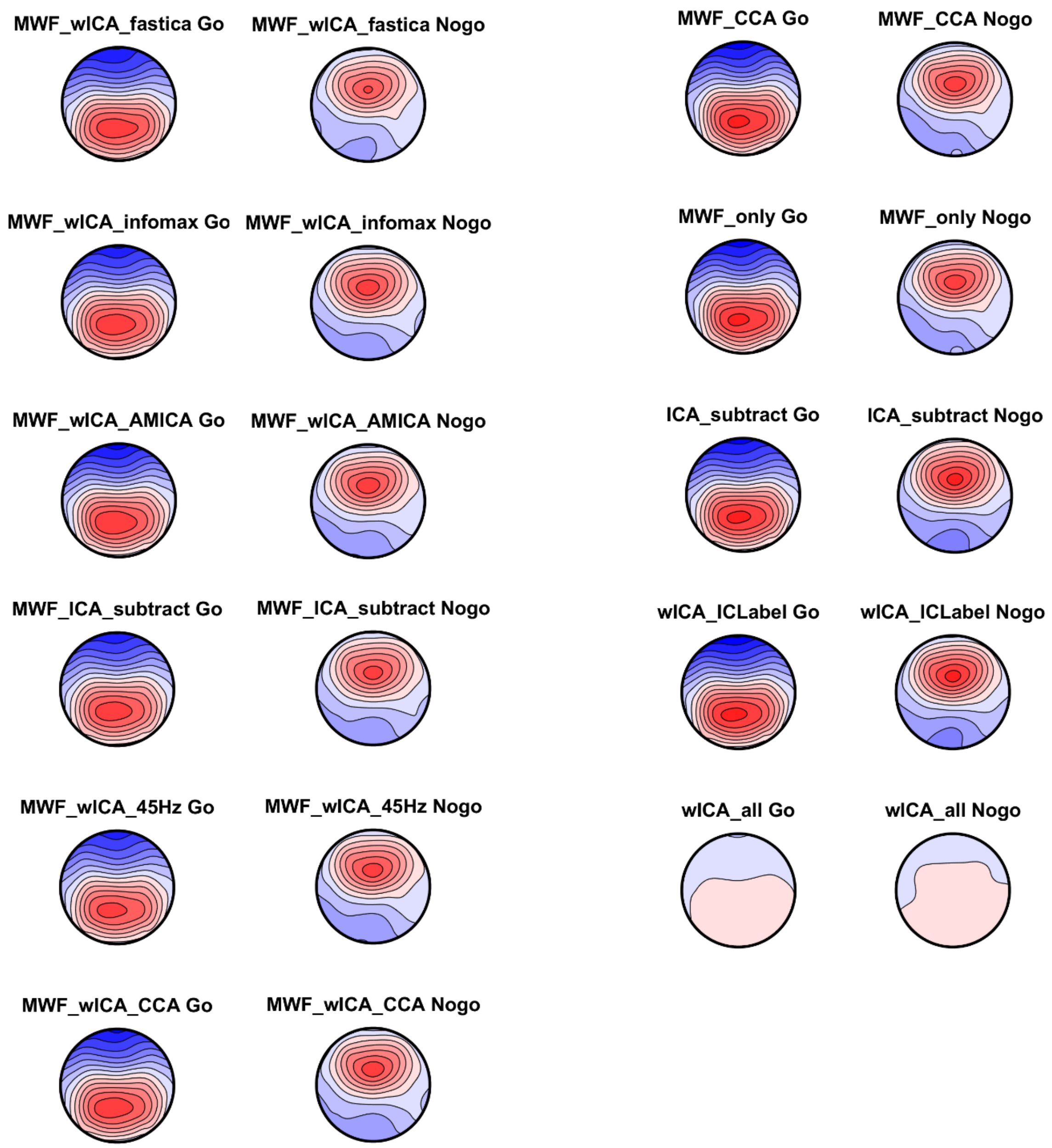

Figure S42. P3 distribution topoplots for Go and Nogo trials from each pipeline. All topoplots are depicted on their own scale so the distribution of activity is easy to understand (but comparison of amplitudes is not possible between conditions and pipelines). Note that all pipeline's produced similar amplitudes and patterns except for wICA_all, which produced a severely reduced amplitude at all electrodes.

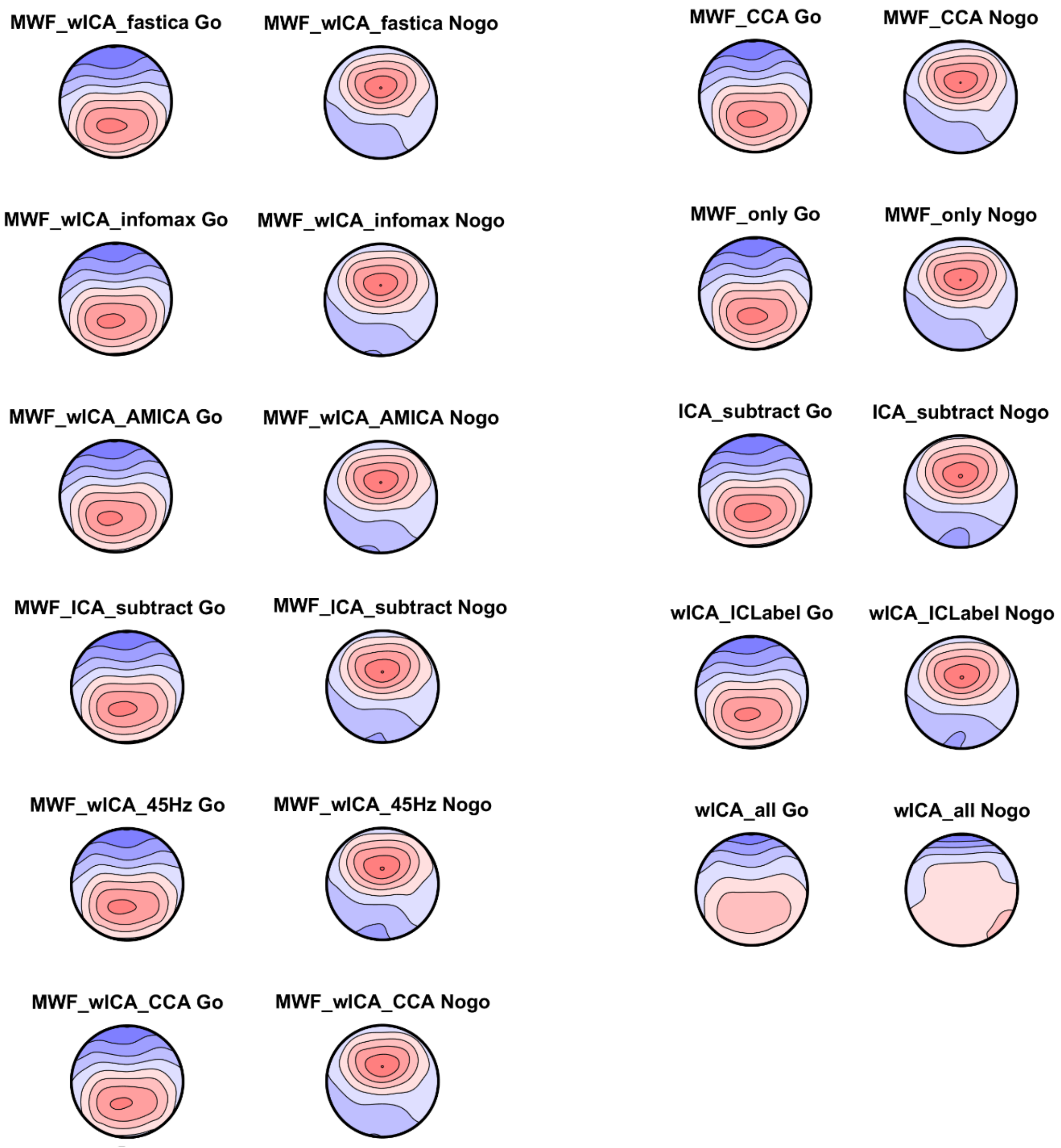

Figure S43. P3 distribution topoplots for Go and Nogo trials from each pipeline. All topoplots are depicted on the individual scale within each topoplot so comparison is possible between the pattern of each distribution without reference to a similar amplitude between the conditions / pipelines. Note that all pipelines showed similar distributions and amplitudes with the exception of wICA_all which showed an altered pattern.

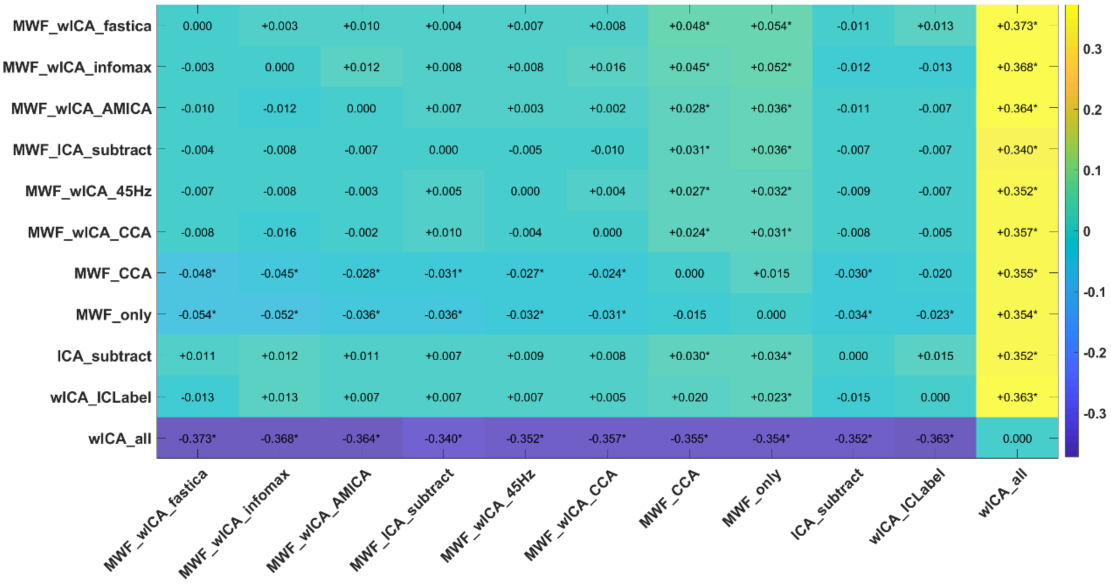

Figure S44. Heat map of the variance explained (np$^2$) by the interaction between each pair of pipelines and averaged activity within the P3 window TANOVA for Go vs Nogo trials. Interactions that were significant (FDR-p < 0.05) are indicated with an *. We have also provided an indication of which pipeline of each pair provided larger values for variance explained using – and + symbols, which can be interpreted as the pipeline listed on the left of the heatmap having shown less (-) or more (+) variance explained in the comparison between the two experimental conditions than the pipeline listed at the bottom of the heatmap.

#### *ERP Amplitude Reliability Metrics*

##### *Number of Errors Required for Dependable Analysis of the Pe*

With regards to the dependability of the Pe ERP data from error related epochs, MWF_wICA_infomax, MWF_wICA_CCA, and MWF_wICA_45Hz appeared to be the best performers, requiring only eight epochs for valid analysis and only excluding 4/76 participants. MWF_ICA_subtract, MWF_only, MWF_wICA_fastICA and MWF_wICA_AMICA all provided the same level of dependability (requiring only 8 error related epochs). However, these pipelines excluded more participants (as a larger number of epochs had to be removed by the cleaning approaches for these pipelines). ICA_subtract, MWF_CCA, and wICA_ICLabel showed less dependable Pe amplitudes across all trials and participants, requiring nine error related epochs for valid analysis (and excluding 7-8 participants). wICA_all provided the least dependable data, with 12 epochs required and 18 participants excluded. Figure S45 depicts the results of this analysis for each pipeline.

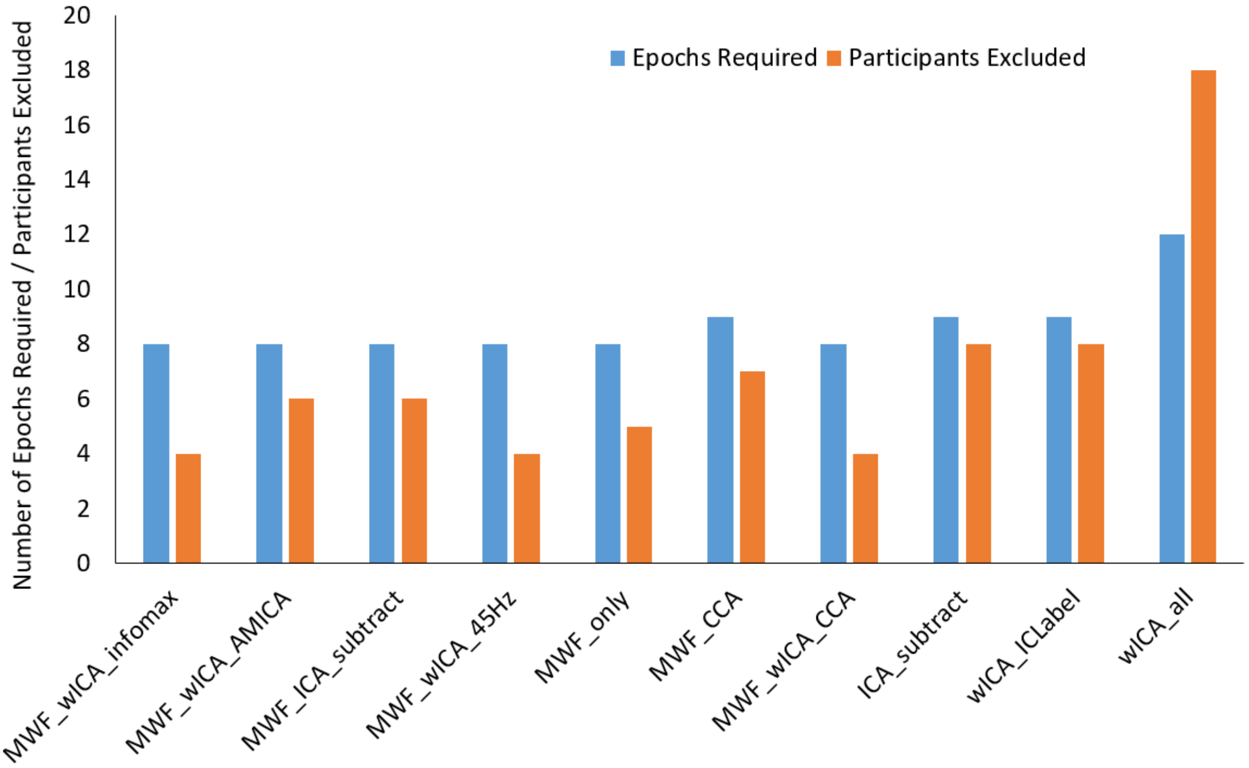

Figure S45. The minimum number of epochs required for a dependability of 0.8 in an analysis of the error related Pe at FCz, and number of participants excluded from error processing analyses based on this dependability threshold for each of the cleaning pipelines.

#### ***ERP Amplitude and Single Trial Bootstrap Standard Error of the Mean***

##### **N2 bSME**

We ran the analysis of the bSME on peak detections (rather than averaged windows of interest), as peak detection methods of measuring ERPs are more vulnerable to artifacts, since high frequency muscle artifacts can result in a spike in a small number of timepoints (which can be averaged out by average window ERP measures). For the bSME of the Nogo Peak N2 at FCz, there was a significant difference between the pipelines rANOVA $F(202.23,2.7) = 728.05$, $p < 0.0001$. The rank order of significant differences was: wICA_all > MWF_wICA_45Hz*, MWF_wICA_fastICA, MWF_ICA_subtract, MWF_wICA_infomax, MWF_wICA_AMICA, MWF_wICA_CCA** > MWF_CCA, MWF_only > ICA_subtract, wICA_ICLabel (Figure S46-48). Note that it is helpful to consider the bSME and amplitude of an ERP together, as higher variability is likely in larger amplitude ERPs, and may not be such an issue, whereas the same bSME in a low amplitude ERP is more likely to be an issue for data analysis. wICA_all showed both very small bSME values and very small amplitudes. Figure S46-47 demonstrates that although ICA_subtract and wICA_ICLabel show the highest amplitudes, they also show the highest bSME.

Because the N2 peak amplitudes all showed negative values with the exception of six files that were excluded (the same polarity for all values making ratios valid to calculate and compare), we analysed bSME : Amplitude ratios. For the bSME : Amplitude ratio of the Nogo Peak N2 at FCz, there was a significant difference between the pipelines rANOVA F(225.51,

3.18) = 208.57, p < 0.0001. The rank order from best performing pipeline to worst performing pipeline of significant differences was: wICA_all > MWF_ICA_subtract*^, MWF_wICA_CCA*^, ICA_subtract^@, MWF_wICA_fastICA*, MWF_wICA_infomax*, MWF_wICA_AMICA*, MWF_only*, wICA_ICLabel@@, MWF_CCA^^, MWF_wICA_45Hz** (Figure 49-50).

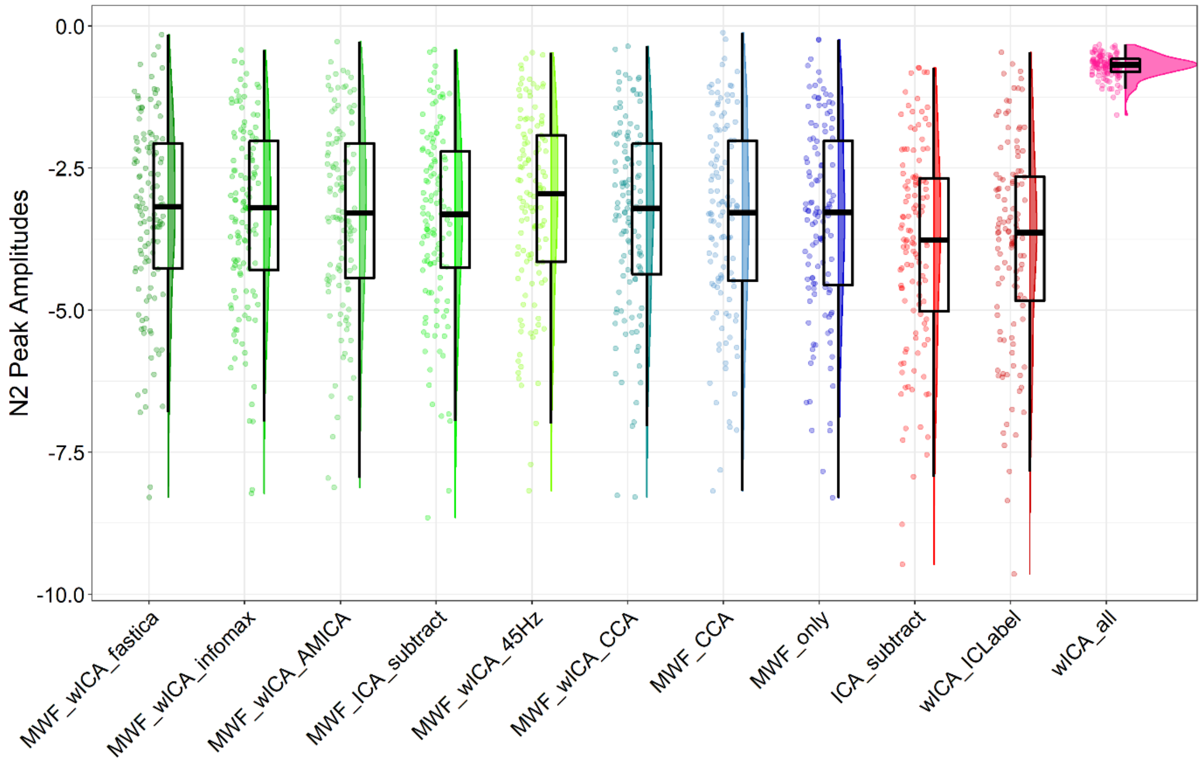

Figure S46. Raincloud plot depicting Nogo N2 peak amplitude values from FCz.

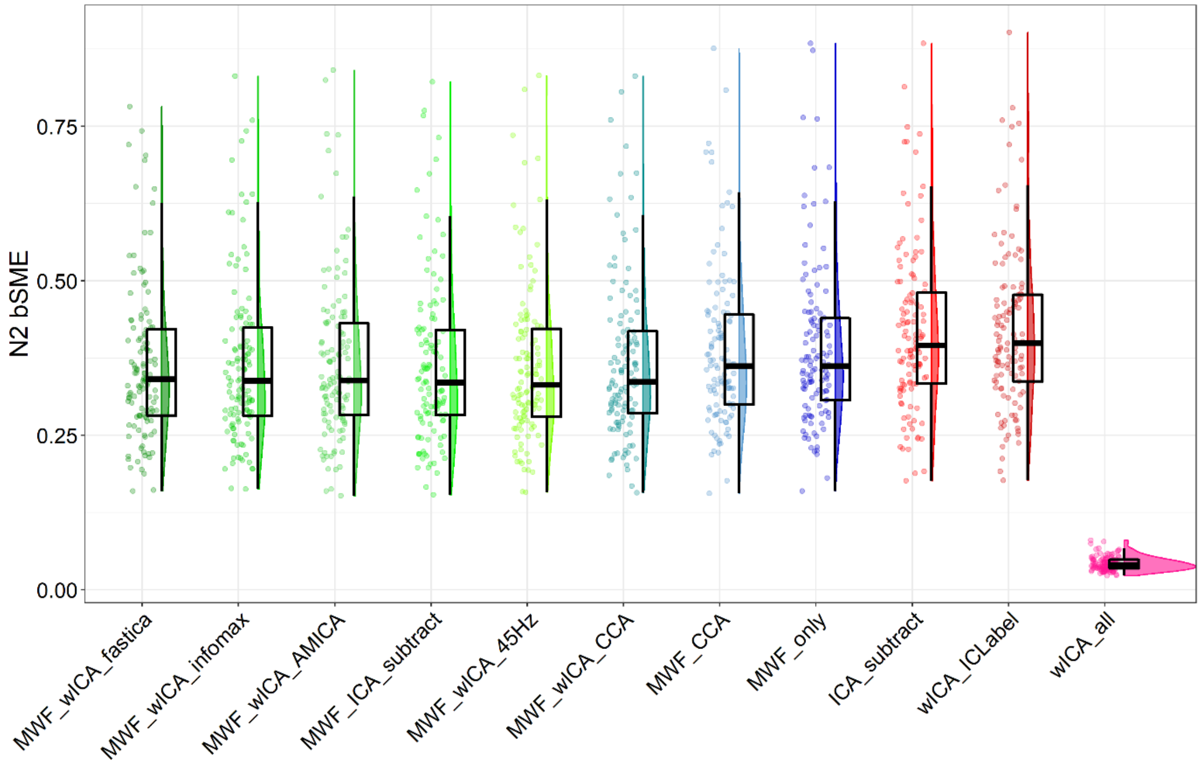

Figure S47. Raincloud plot depicting Nogo N2 peak amplitude bSME values from FCz.

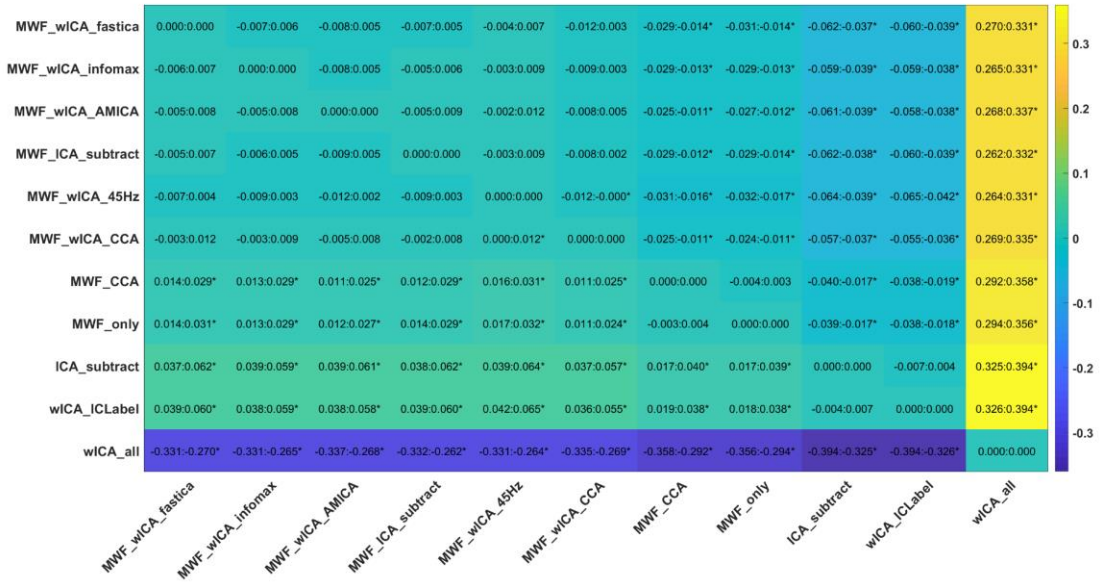

Figure S48. Post-hoc t-test results for the Nogo N2 peak amplitude bSME values from FCz.

| | N2 bSME | | N2 peak | |
|---|---|---|---|---|
| **Pipeline** | **Mean** | **SD** | **Mean** | **SD** |
| **MWF_ICA_subtract** | 0.365 | 0.134 | -3.420 | 1.640 |
| **MWF_CCA** | 0.389 | 0.129 | -3.396 | 1.729 |
| **MWF_wICA_fastICA** | 0.366 | 0.126 | -3.294 | 1.658 |
| **MWF_wICA_infomax** | 0.367 | 0.133 | -3.346 | 1.639 |
| **MWF_wICA_CCA** | 0.370 | 0.133 | -3.358 | 1.666 |
| **MWF_wICA_45Hz** | 0.364 | 0.129 | -3.154 | 1.651 |
| **MWF_only** | 0.391 | 0.132 | -3.435 | 1.733 |
| **wICA_ICLabel** | 0.420 | 0.132 | -3.827 | 1.798 |
| **wICA_all** | 0.042 | 0.011 | -0.706 | 0.199 |
| **MWF_wICA_AMICA** | 0.368 | 0.132 | -3.323 | 1.680 |
| **ICA_subtract** | 0.420 | 0.133 | -3.886 | 1.816 |

Table S6. Means and SDs for the N2 peak amplitude bSME and N2 peak amplitude from Nogo trials and FCz.

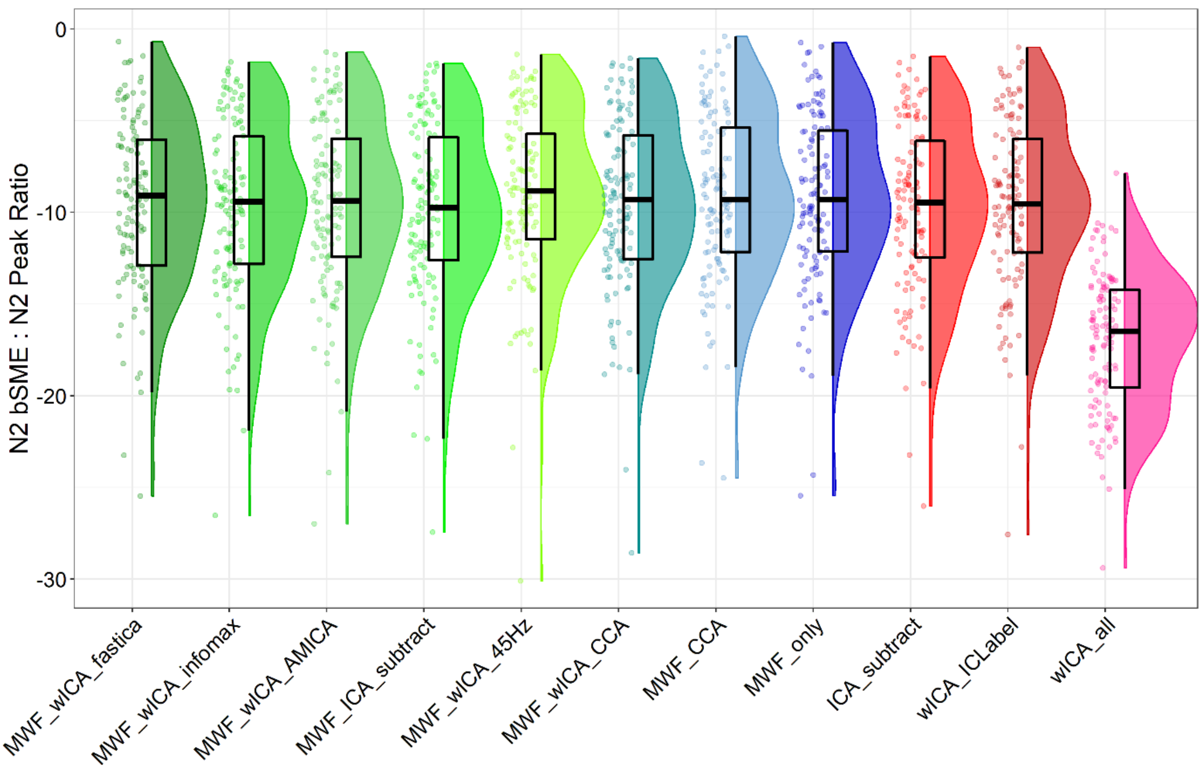

Figure S49. Raincloud plot depicting the ratio between Nogo N2 peak amplitude bSME values and Nogo N2 peak amplitudes from FCz.

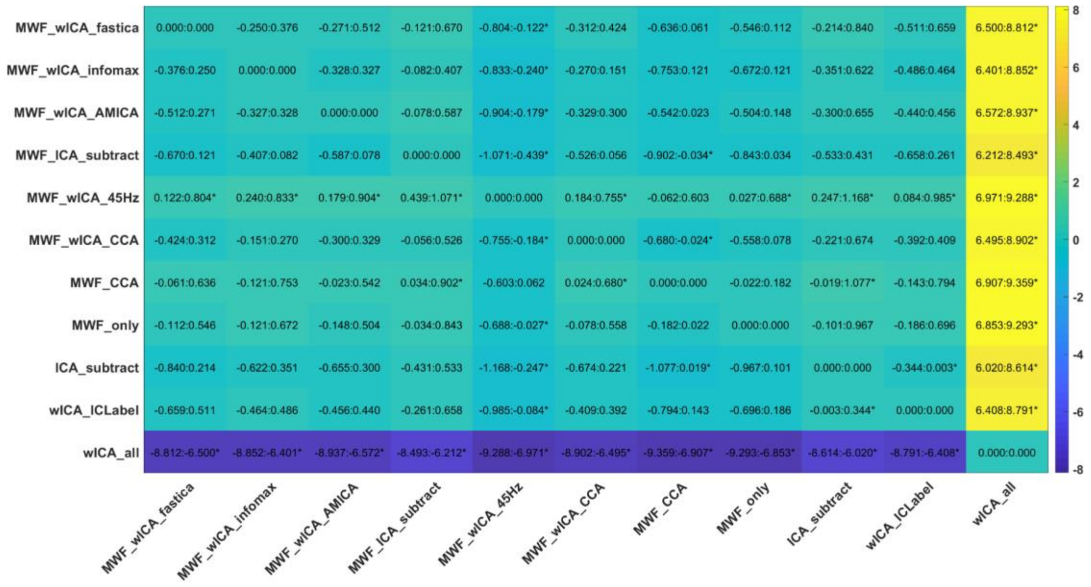

Figure S50. Post-hoc t-test results for the ratio between Nogo N2 peak amplitude bSME values and Nogo N2 peak amplitudes from FCz.

**P3 bSME**

For the bSME of the Go Peak P3 at Pz, there was a significant difference between the pipelines rANOVA $F(193.99, 2.55) = 537.05$, $p < 0.0001$. The rank order of significant differences was: wICA_all > MWF_wICA_45Hz$^{+}$, MWF_wICA_fastICA^*, MWF_wICA_infomax$^{++}$, MWF_wICA_AMICA$^{++}$^^, MWF_ICA_subtract,

MWF_wICA_CCA++^^** > MWF_CCA, MWF_only > ICA_subtract, wICA_ICLabel. Note that it is helpful to consider the bSME and amplitude of an ERP together, as higher variability is likely in larger amplitude ERPs, and may not be such an issue, whereas the same bSME in a low amplitude ERP is more likely to be an issue for data analysis. As with the N2, wICA_all showed both very small bSME values and very small amplitudes. Figure S51-53 demonstrates that although ICA_subtract and wICA_ICLabel show the highest amplitudes, they also show the highest bSME, and their peak amplitude to bSME ratio is lower than the RELAX methods.

Since all Go trial P3 peak amplitude measures showed positive values (so ratios provide a valid measure for analysis), we analysed the ratio of the bSME to P3 peak amplitude. There was a significant difference between the pipelines $F(4.37, 332.18) = 250.183$, $p < 0.0001$. The rank order of significant differences was: wICA_all > MWF_ICA_subtract+, MWF_wICA_AMICA*, MWF_wICA_fastICA*, MWF_wICA_infomax^, MWF_wICA_CCA^++, MWF_only@, MWF_CCA++**@@!, wICA_ICLabel**^^++, ICA_subtract**^^++, MWF_wICA_45Hz**^^++@@!!. Note that while wICA_all performed the best in the ratio comparison, peak amplitudes from this pipeline were on average 1.24μV, when compared to the peaks of 6.6μV to 7.6μV from other pipelines (see Figure S51). Figures S54-56 depict these ratios and the post-hoc tests.

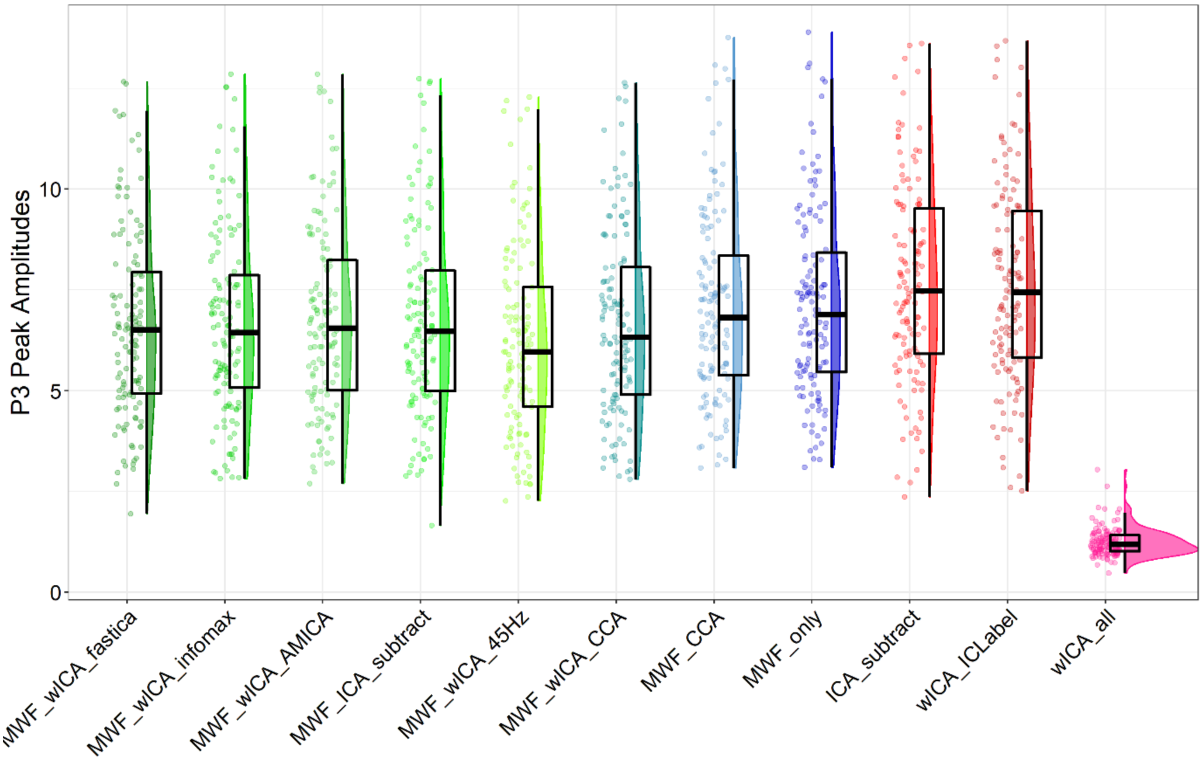

Figure S51. Peak amplitudes from the peak detection method for the P3 at Pz from Go trials.

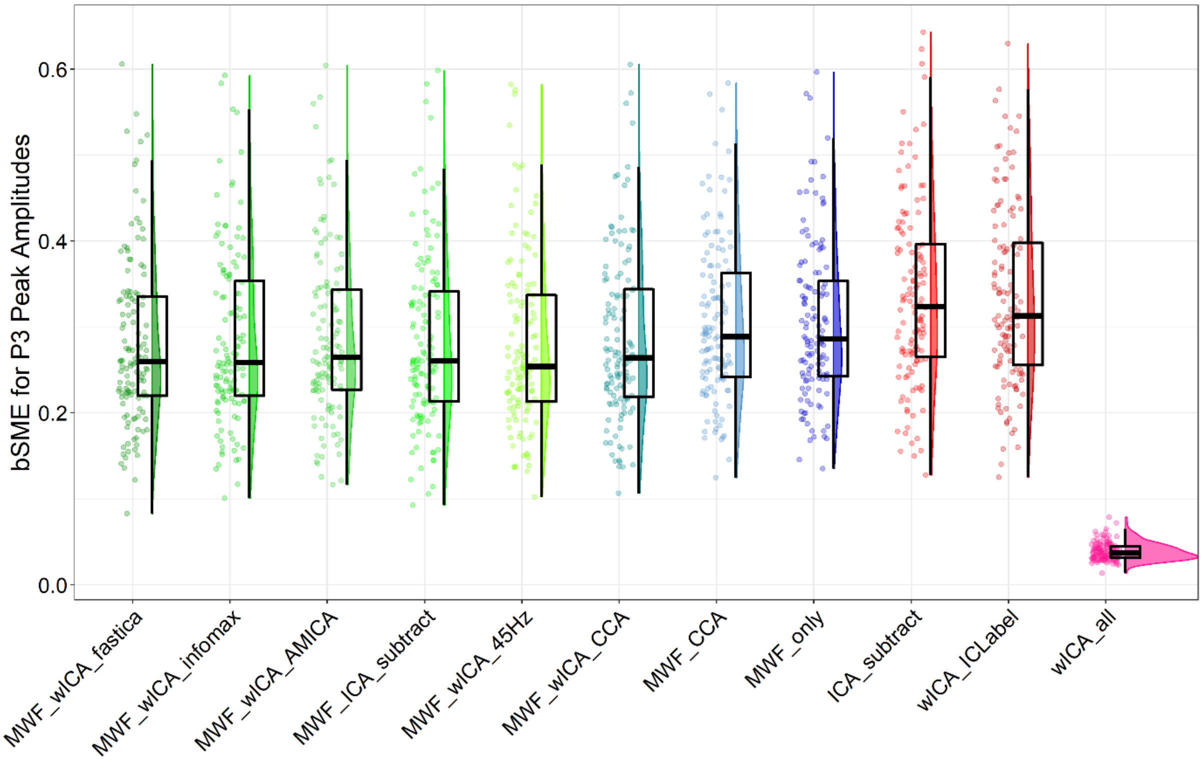

Figure S52. Raincloud plot depicting Go P3 peak amplitude bSME values from Pz.

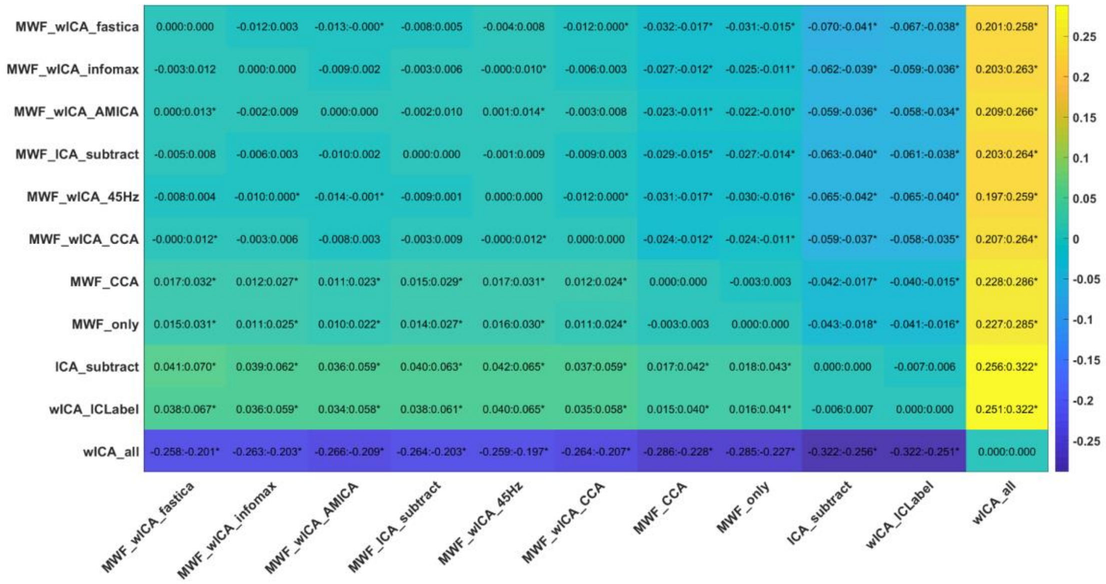

Figure S53. Post-hoc t-test results for the Go P3 peak amplitude bSME values from Pz.

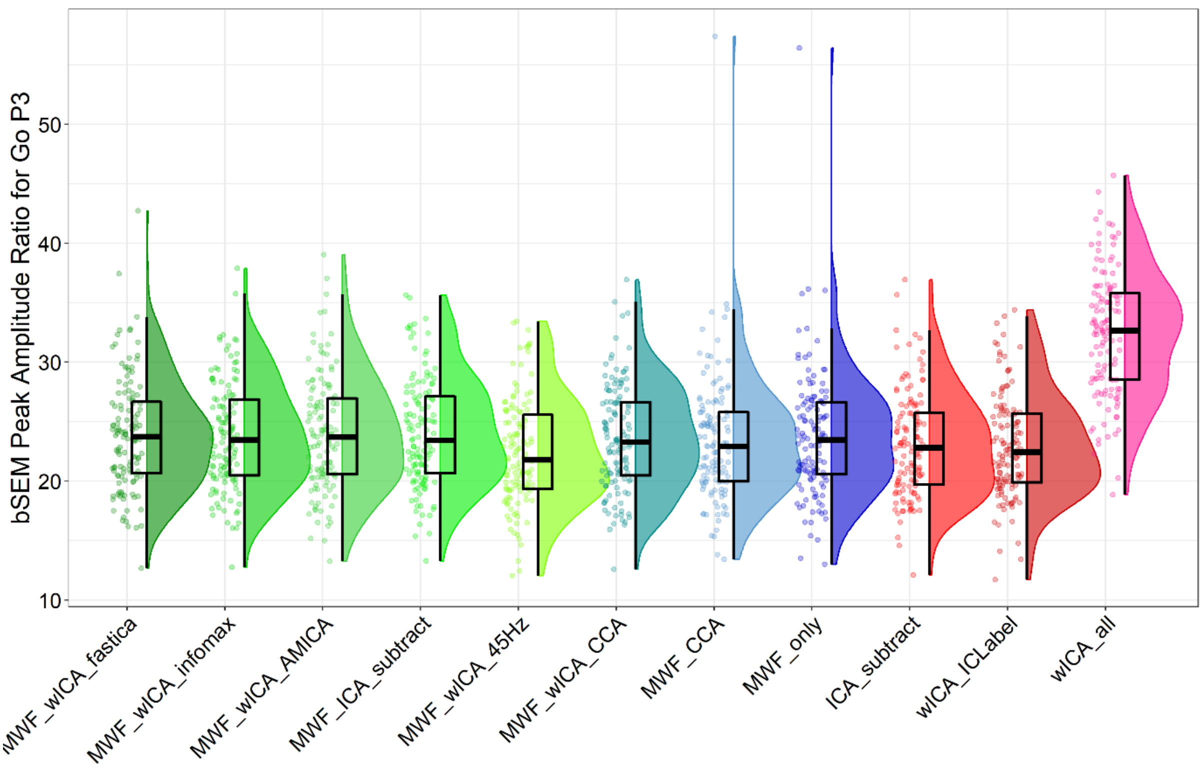

Figure S54. The ratio of peak P3 bSME values to P3 peak amplitude values for peak amplitude detection of the P3 in Go trials.

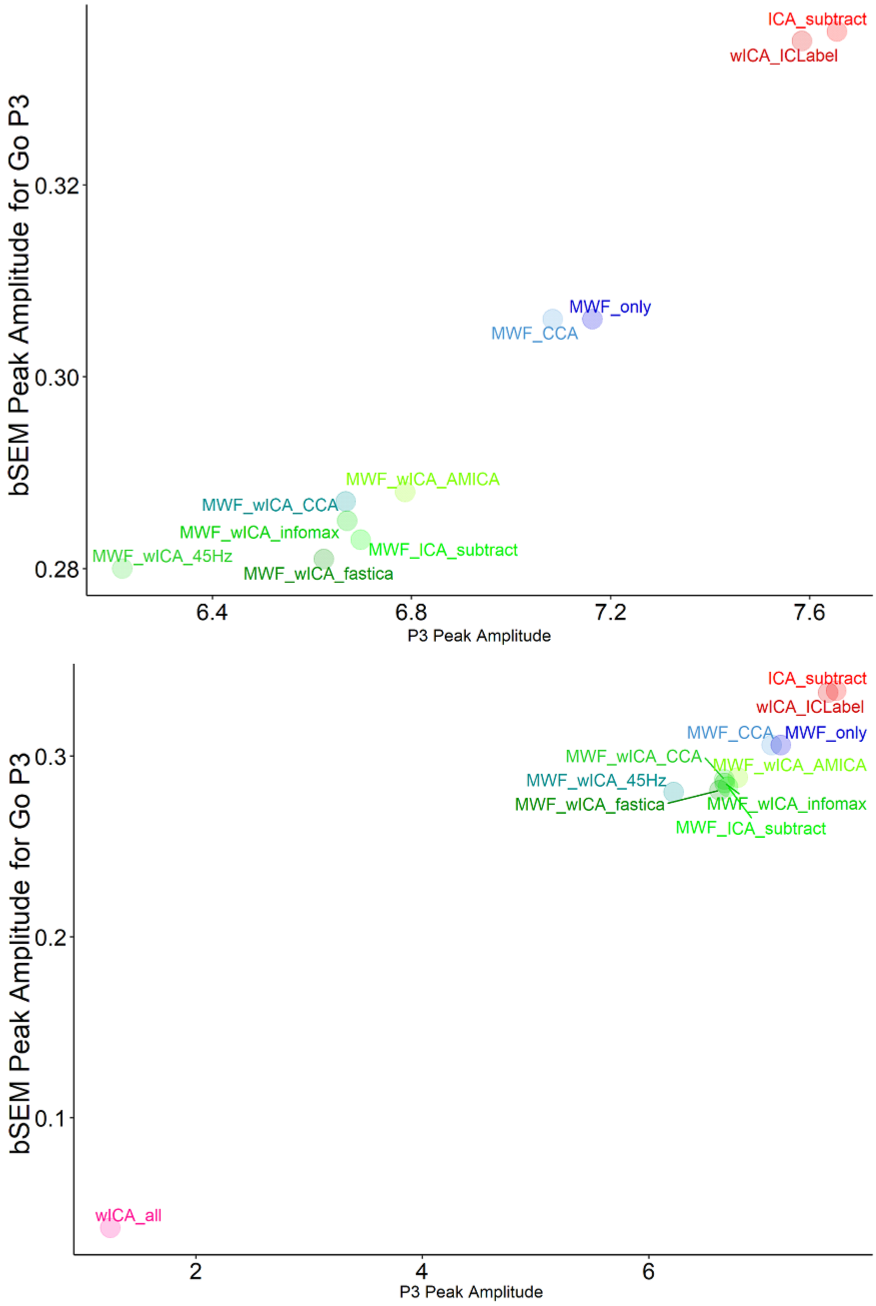

Figure S55. Scatterplot depicting mean bSME values against mean amplitude values for peak amplitude detection of the P3 in Go trials from each cleaning pipeline (above, excluding wICA_all to provide sufficient resolution to discern differences in the other pipelines, and below, including all pipelines).

| | P3 bSME | | P3 peak | |
|---|---|---|---|---|
| **Pipeline** | **Mean** | **SD** | **Mean** | **SD** |
| **MWF_CCA** | 0.306 | 0.092 | 7.084 | 2.361 |
| **MWF_wICA_infomax** | 0.285 | 0.100 | 6.671 | 2.332 |
| **MWF_wICA_45Hz** | 0.280 | 0.100 | 6.219 | 2.329 |
| **MWF_wICA_CCA** | 0.287 | 0.099 | 6.668 | 2.325 |
| **MWF_wICA_AMICA** | 0.288 | 0.099 | 6.787 | 2.307 |
| **MWF_wICA_fastICA** | 0.281 | 0.097 | 6.624 | 2.323 |
| **wICA_all** | 0.039 | 0.011 | 1.243 | 0.355 |
| **MWF_only** | 0.306 | 0.093 | 7.164 | 2.359 |
| **wICA_ICLabel** | 0.335 | 0.105 | 7.585 | 2.467 |
| **ICA_subtract** | 0.336 | 0.106 | 7.655 | 2.467 |
| **MWF_ICA_subtract** | 0.283 | 0.101 | 6.698 | 2.376 |

Table S7. Means and SDs for the P3 peak amplitude bSME and P3 peak amplitude from Go trials and Pz.

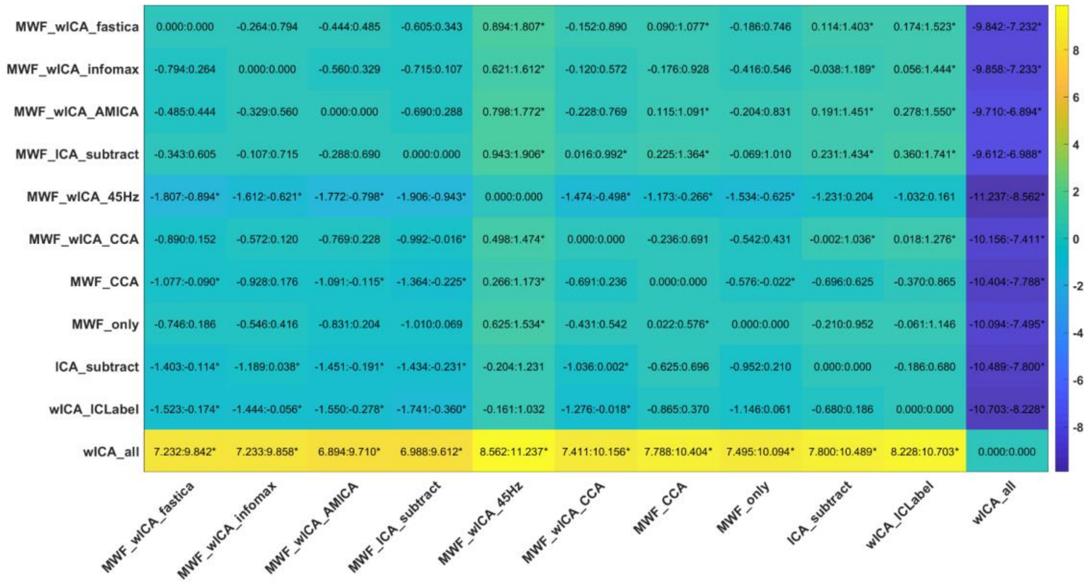

Figure S56. Post-hoc t-test results for the ratio between Go P3 peak amplitude bSME values and Go P3 peak amplitudes from Pz.

#### N1 bSME

For the bSME of the N1 Peak at FCz, there was a significant difference between the pipelines rANOVA $F(2.22, 166.6) = 766.81$, $p < 0.0001$. The rank order from best performing pipeline to worst performing pipeline of significant differences was: wICA_all > MWF_wICA_45Hz$^{+}$, MWF_wICA_fastICA, MWF_wICA_AMICA, MWF_ICA_subtract, MWF_wICA_infomax$^{++}$, MWF_wICA_CCA$^{++}$ > MWF_CCA, MWF_only > ICA_subtract, wICA_ICLabel (Figure S57-58).

Since all N1 peak amplitude measures showed negative values (all values showing the same polarity makes ratios more valid to calculate), we analysed the ratio of the bSME to N1 peak amplitude. There was a significant difference between the pipelines $F(4.1, 307.84) = 191.6258$, $p < 0.0001$. The rank order of significant differences was: wICA_all > MWF_ICA_subtract*, MWF_wICA_AMICA^, MWF_wICA_infomax$^{+}$, MWF_wICA_fastICA, MWF_wICA_CCA, MWF_only**, ICA_subtract**, wICA_ICLabel**^^, MWF_CCA**^^$^{++}$ > MWF_wICA_45Hz. Note that while wICA_all performed the best in the ratio comparison, peak amplitudes from this pipeline were on average -1.23μV, perhaps providing very little potential variation to detect between group or condition differences (as suggested by our measures of explained variance in the preceding section and in comparison to the peaks of -8.2μV to -6.8μV from other pipelines, Figure 59). Figures S60 and S63 depict these ratios and the post-hoc tests.

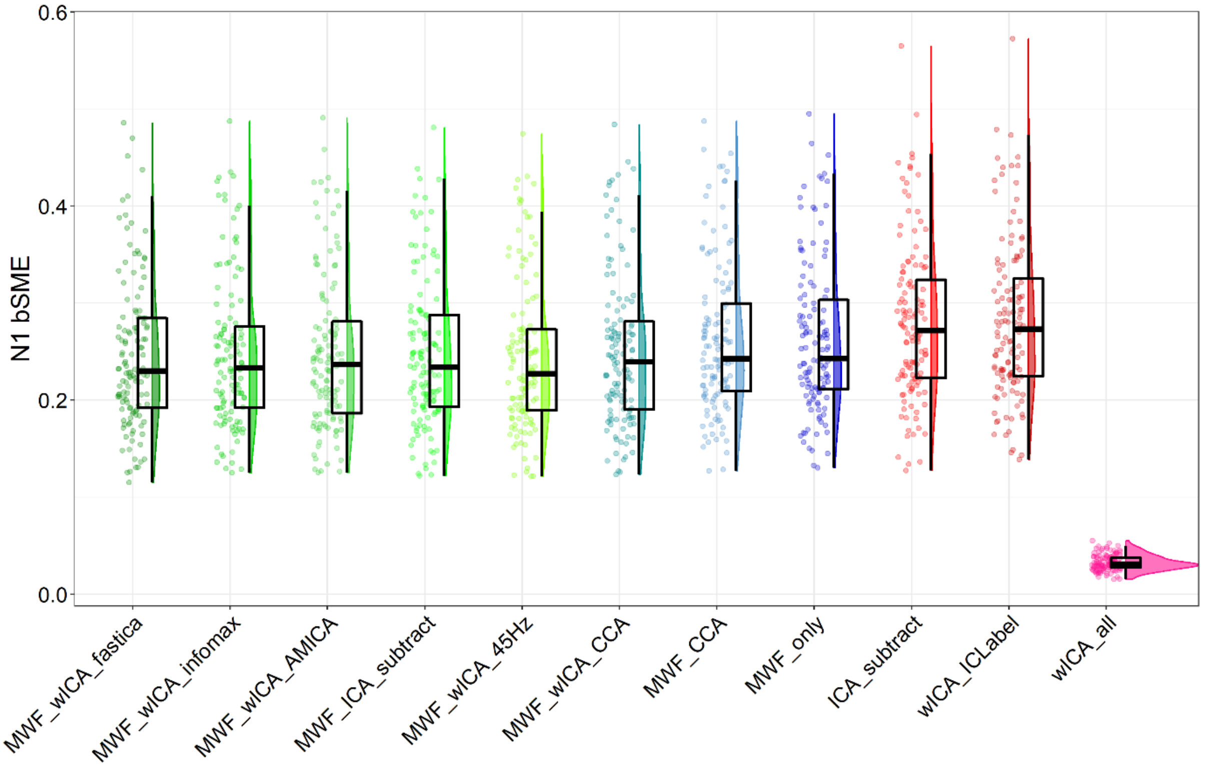

Figure S57. bSME values from the peak detection method for the N1 at FCz from the Go-Nogo dataset.

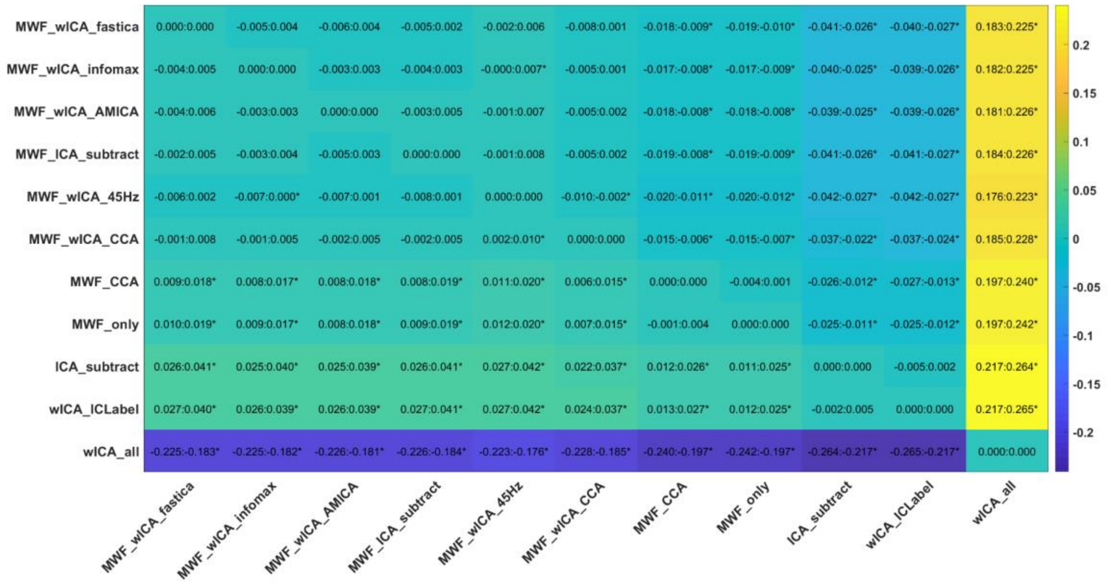

Figure S58. Post-hoc t-test results for the bSME values for N1 peak amplitudes from FCz from the Go-Nogo dataset.

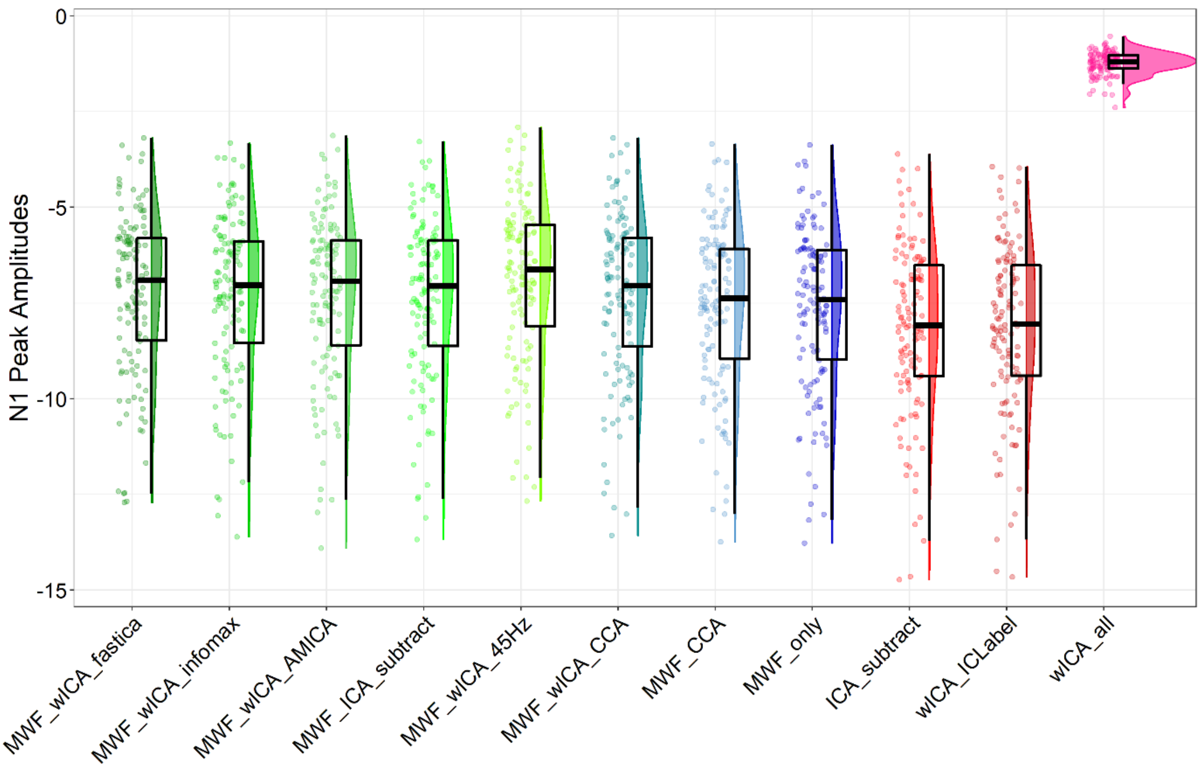

Figure S59. Peak amplitudes from the peak detection method for the N1 at FCz from the Go-Nogo dataset.

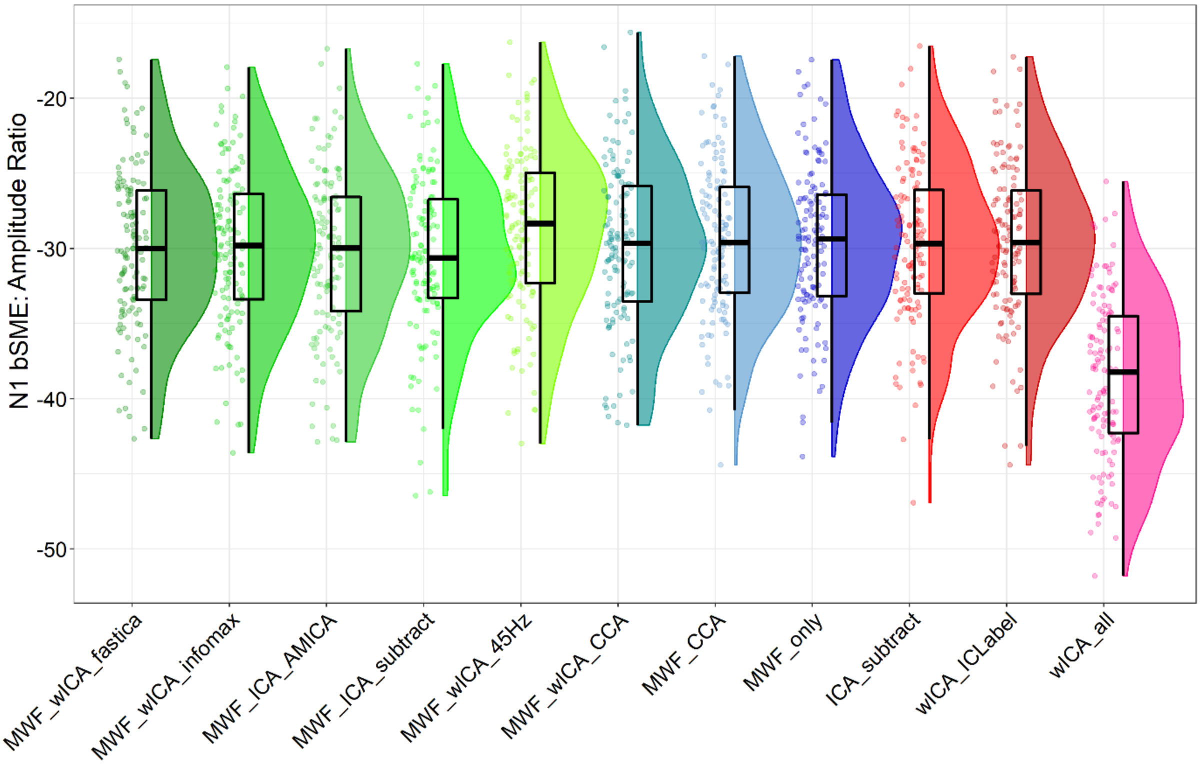

Figure S60. The ratio of peak N1 bSME values to N1 peak amplitude values for peak amplitude detection of the N1 in Go and Nogo trials.

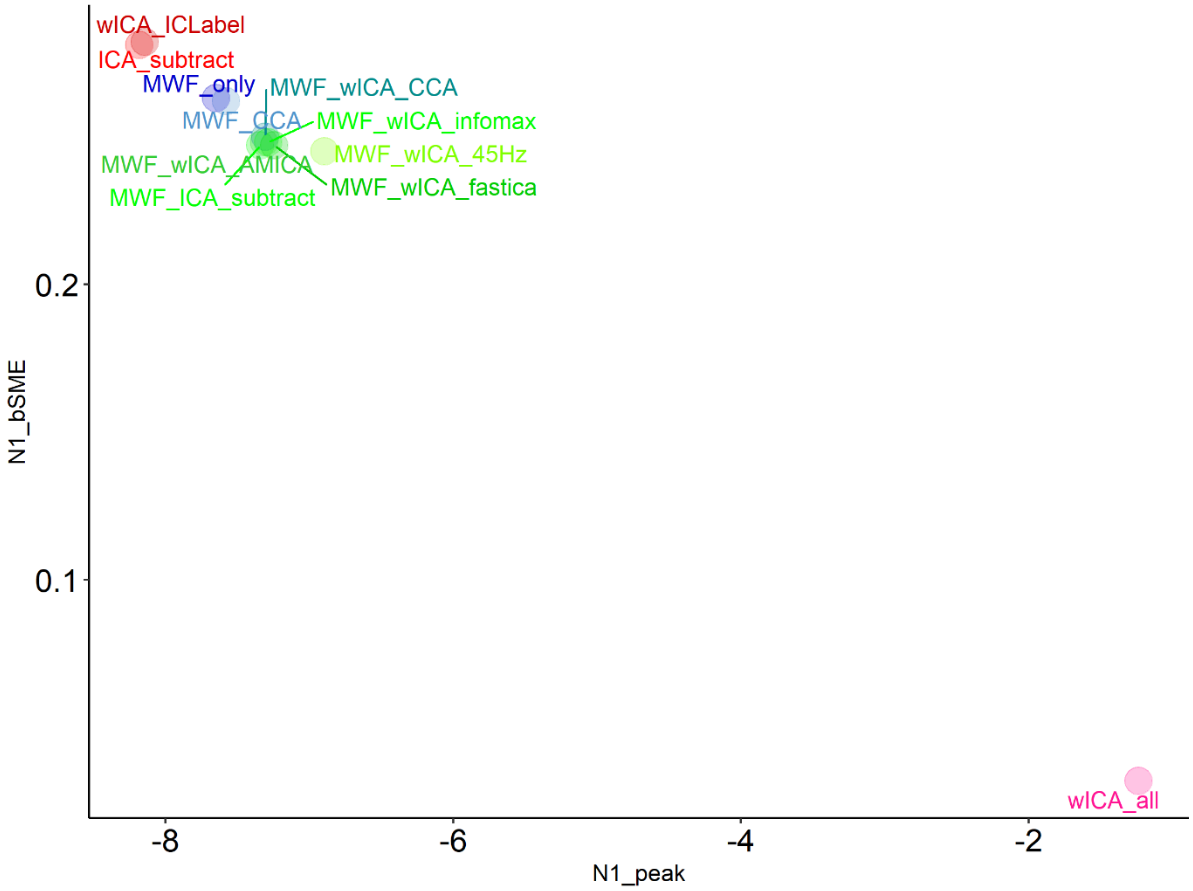

Figure S61. Scatter plot depicting N1 bSME values and N1 peak amplitudes from FCz.

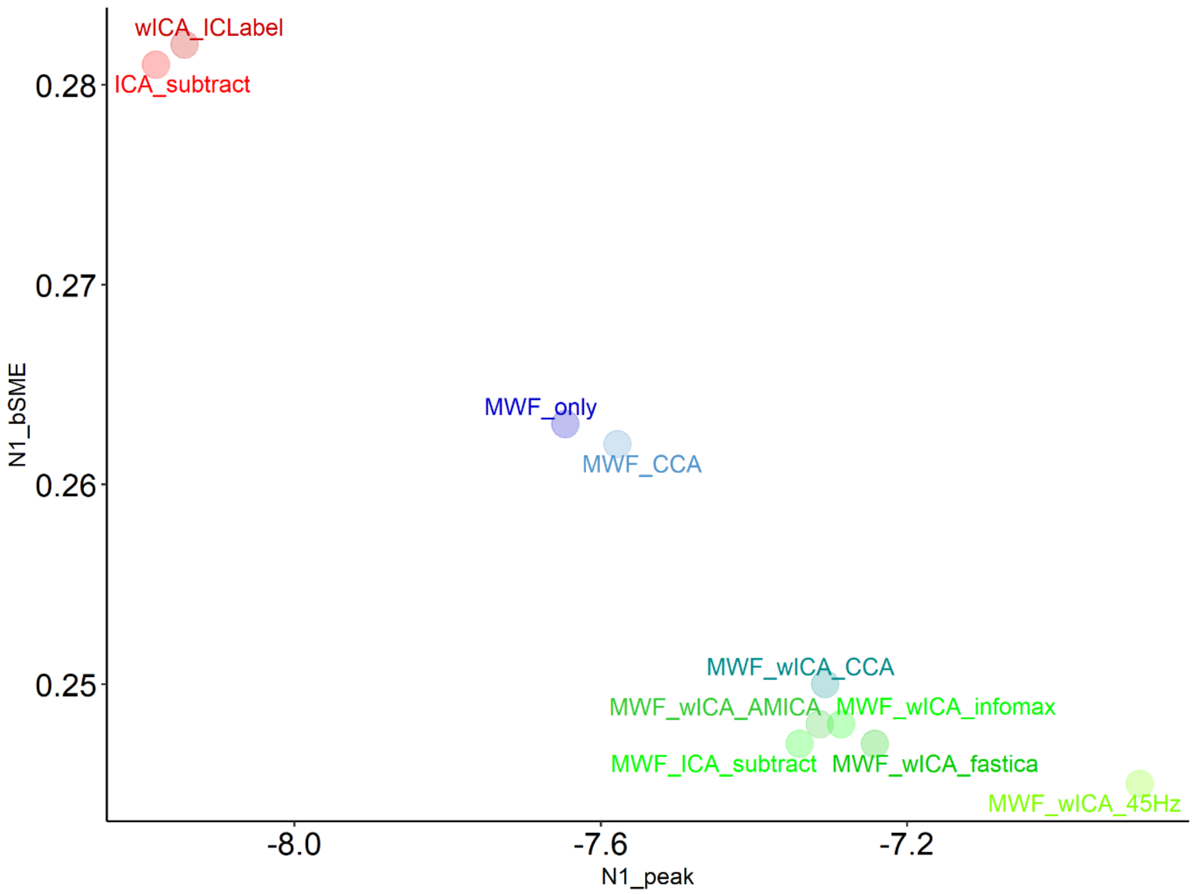

Figure S62. Scatter plot depicting N1 bSME values and N1 peak amplitudes from FCz from the Go-Nogo dataset, excluding wICA_all to improve the reader’s ability to distinguish the other pipelines.

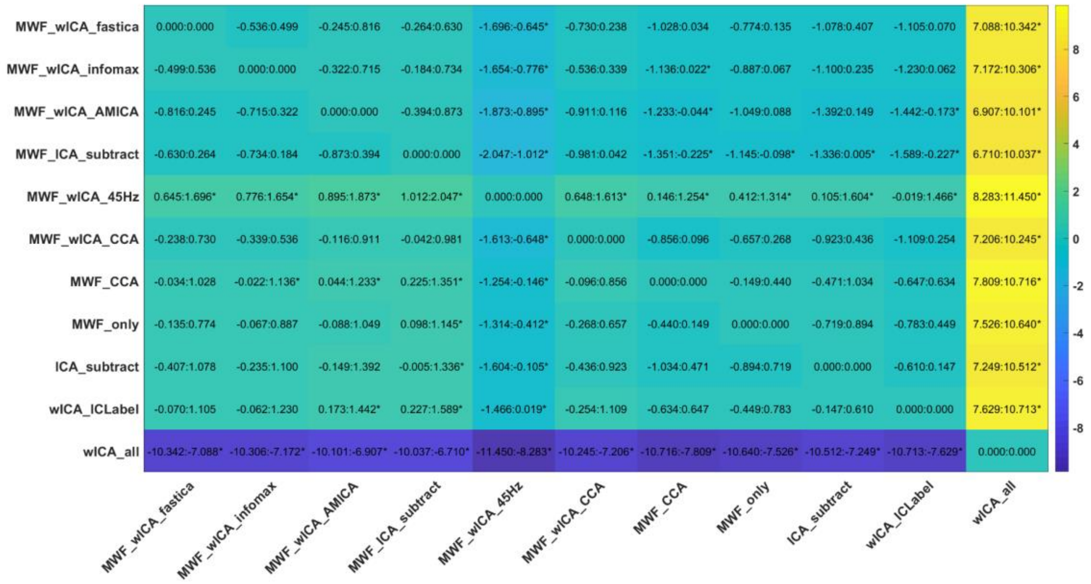

Figure S63. Post-hoc t-test results for the ratio between N1 peak amplitude bSME values and N1 peak amplitudes from FCz.

| | N1 bSME | | N1 peak | |
|---|---|---|---|---|
| **Pipeline** | **Mean** | **SD** | **Mean** | **SD** |
| **MWF_ICA_subtract** | 0.247 | 0.077 | -7.340 | 2.115 |
| **MWF_wICA_AMICA** | 0.248 | 0.080 | -7.314 | 2.165 |
| **wICA_ICLabel** | 0.282 | 0.082 | -8.144 | 2.188 |
| **MWF_wICA_45Hz** | 0.245 | 0.078 | -6.896 | 2.093 |
| **MWF_wICA_infomax** | 0.248 | 0.078 | -7.286 | 2.140 |
| **MWF_wICA_CCA** | 0.250 | 0.079 | -7.307 | 2.141 |
| **MWF_CCA** | 0.262 | 0.078 | -7.578 | 2.138 |
| **wICA_all** | 0.032 | 0.008 | -1.237 | 0.299 |
| **MWF_only** | 0.263 | 0.078 | -7.646 | 2.142 |
| **MWF_wICA_fastICA** | 0.247 | 0.078 | -7.242 | 2.123 |
| **ICA_subtract** | 0.281 | 0.081 | -8.181 | 2.222 |

Table S8. Means and SDs for the N1 peak amplitude bSME and N1 peak amplitude from both Go and Nogo trials at FCz.

### SECTION FIVE

#### RELAX Pipeline Parameter Testing

***FastICA symm vs defl Setting Comparisons within the wICA_ICLabel pipeline***

***Blink Amplitude Ratio***

A robust ANOVA revealed significant overall differences in fBAR when comparing the use of wICA_ICLabel with three different ICA methods: infomax (wICA_ICLabel_infomax), fastica symm (wICA_ICLabel_symm) and fastica defl (wICA_ICLabel_defl) methods: F(1.86, 139.22) = 3.469, p = 0.0372 (Figure S64). However, no significant difference was present between any of the pipelines in the post-hoc t-tests using rmmcp (which implements multiple comparison controls using Hochberg's approach, Figure S65). In order to determine which potential differences were driving the overall significant effect, we re-ran the post-hoc t-tests using pairdebb (bootstrap t-tests, Figure S66). This showed the following rank order of significant differences from best performance to worst performance: wICA_ICLabel_infomax > wICA_ICLabel_defl > wICA_ICLabel_symm. In contrast to the fBAR comparison, there was no significant overall difference in allBAR in the robust ANOVA between wICA_ICLabel infomax, fastica symm or fastica defl methods: F(1.82, 136.62) = 1.66, p = 0.196 (Figure S67).

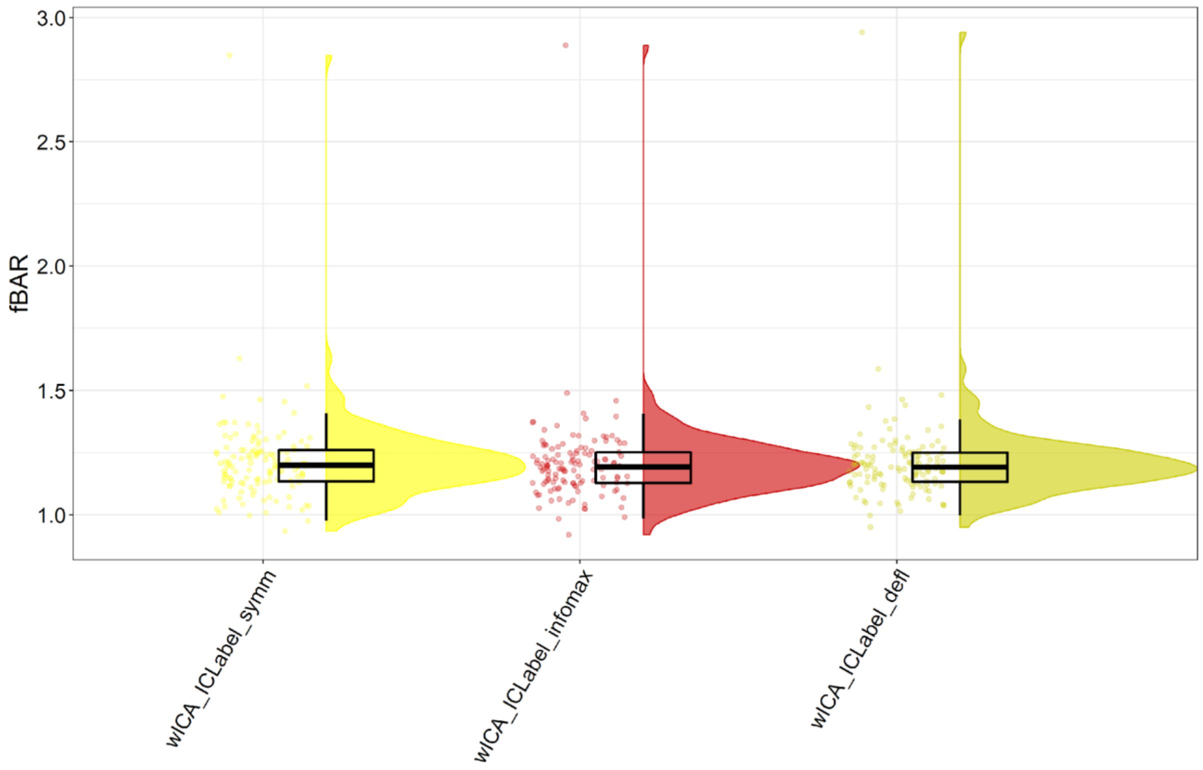

Figure S64. fBAR values from wICA_ICLabel cleaned data across the different ICA methods.

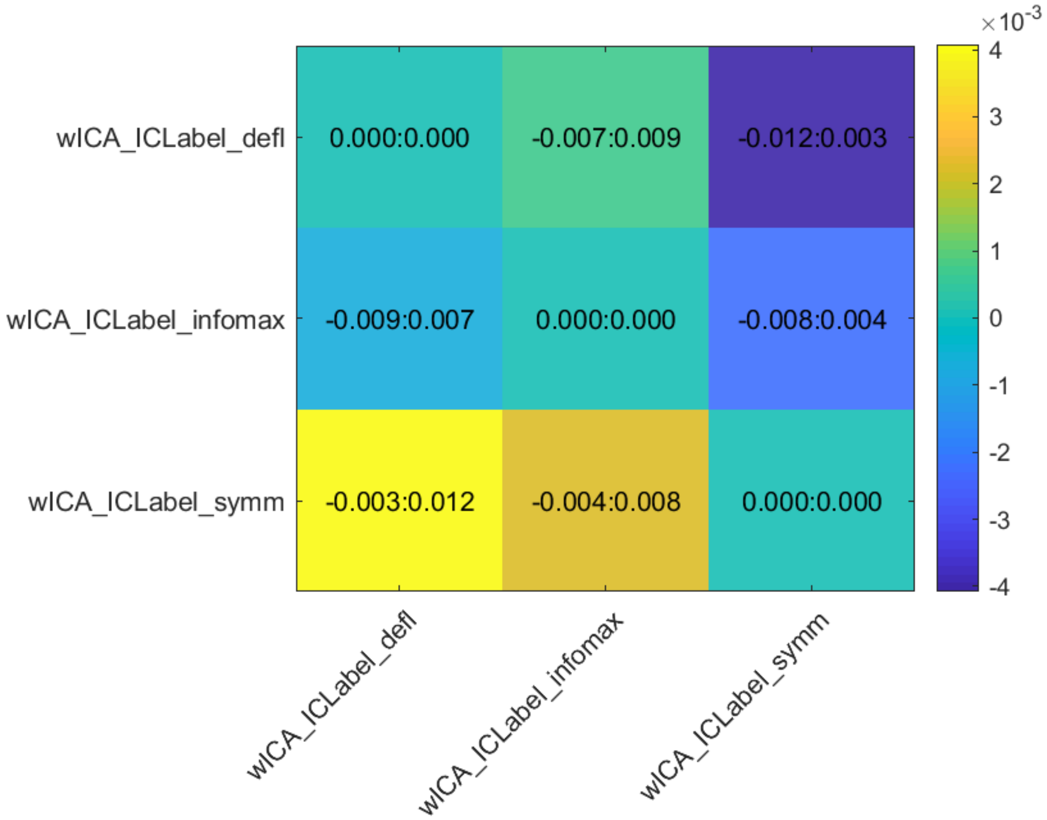

Figure S65. Post-hoc t-tests for fBAR from wICA_ICLabel cleaned data across the different ICA methods, with multiple comparison controls implemented using rmmcp (which implements multiple comparison controls using Hochberg's approach).

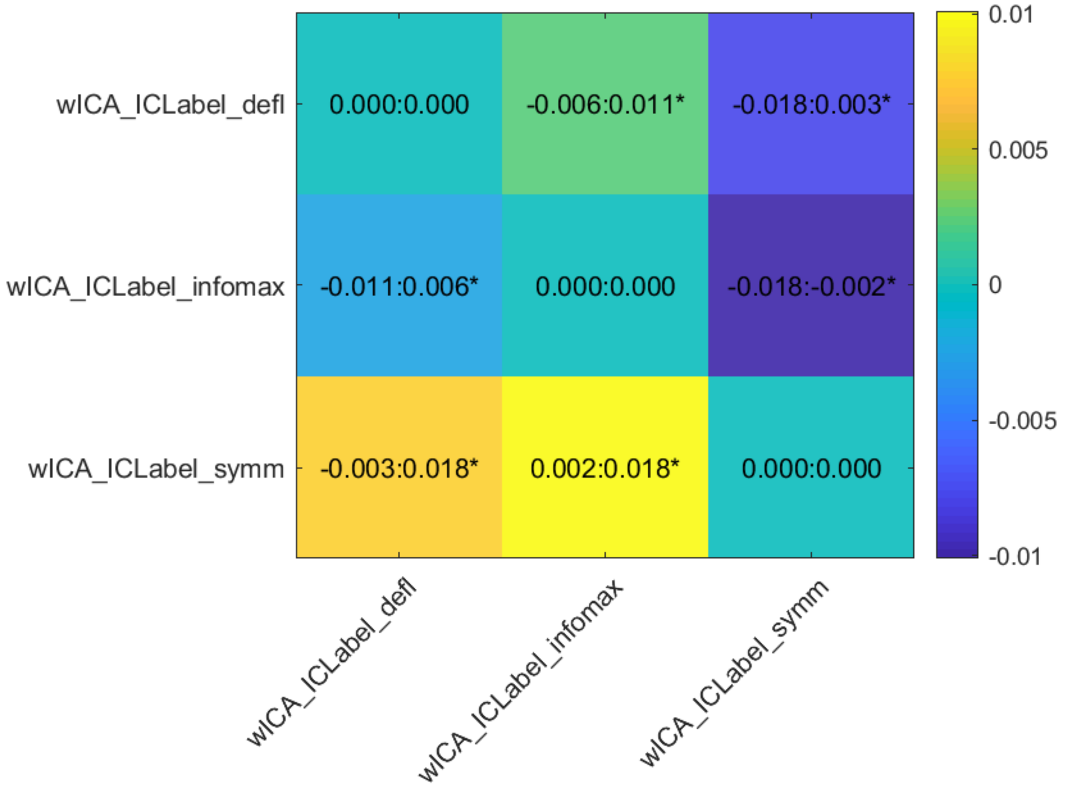

Figure S66. Post-hoc t-tests for fBAR from wICA_ICLabel cleaned data across the different ICA methods, with pairdepb (bootstrap t-tests).

| Pipeline | Mean | SD |
|---|---|---|
| **wICA_ICLabel_defl** | 1.210 | 0.185 |
| **wICA_ICLabel_infomax** | 1.205 | 0.181 |
| **wICA_ICLabel_symm** | 1.215 | 0.186 |

Table S9. Means and SDs for fBAR from wICA_ICLabel cleaned data across the different ICA methods.

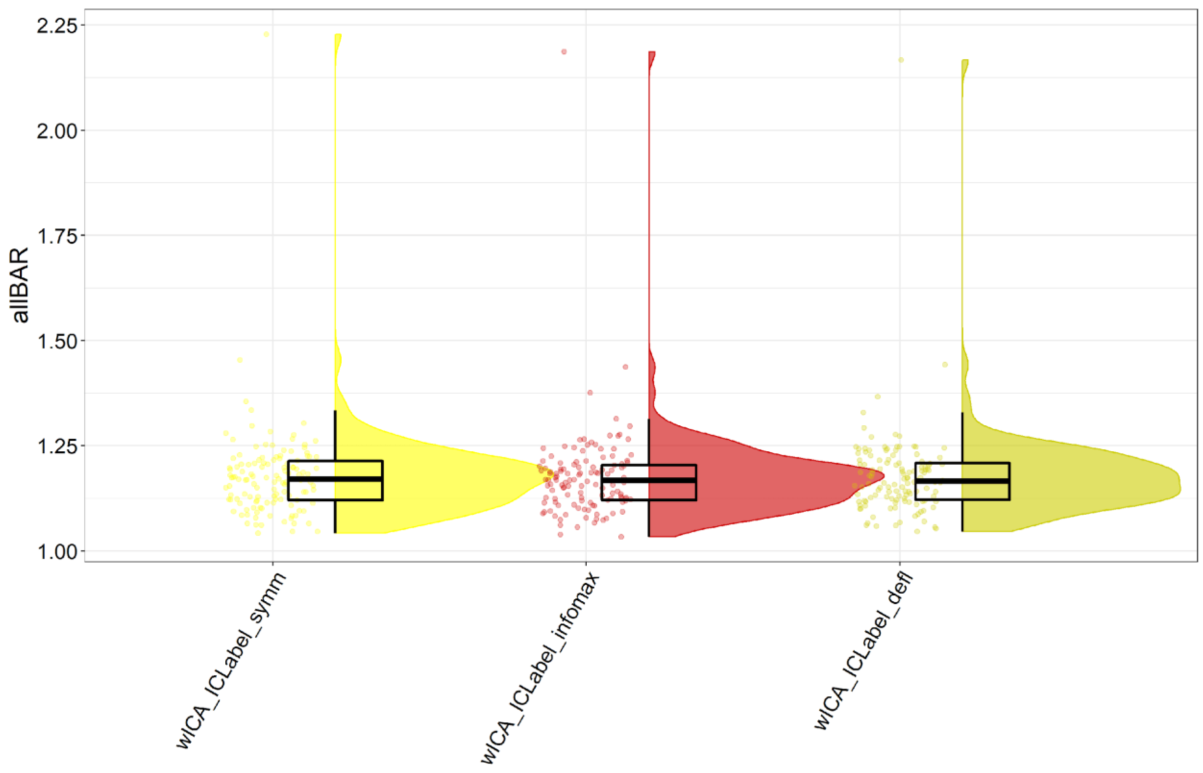

Figure S67. allBAR values from wICA_ICLabel cleaned data across the different ICA methods.

| Pipeline | Mean | SD |
|---|---|---|
| **wICA_ICLabel_symm** | 1.178 | 0.117 |
| **wICA_ICLabel_infomax** | 1.176 | 0.112 |
| **wICA_ICLabel_defl** | 1.178 | 0.110 |

Table S10. Means and SDs for allBAR from wICA_ICLabel cleaned data across the different ICA methods.

#### *Muscle Activity Remaining After Cleaning*

No significant difference was present in the number of epochs showing log-power log-frequency slopes indicative of muscle activity remaining after cleaning across the three ICA methods within the wICA_ICLabel cleaning pipeline: F(1.67, 127.11) = 1.11, p = 0.324. However, there was a significant difference for the amount by which the log-power log-frequency slope exceeded the muscle activity threshold: F(1.81, 137.62) = 4.685, p = 0.0132. Post-hoc t-testing indicated that the defl method performed better than the symm method (but that infomax did not differ from either method, see Figure S68 and S69). It is worth noting that the magnitude of this difference was very small (defl mean – symm mean = 0.009).

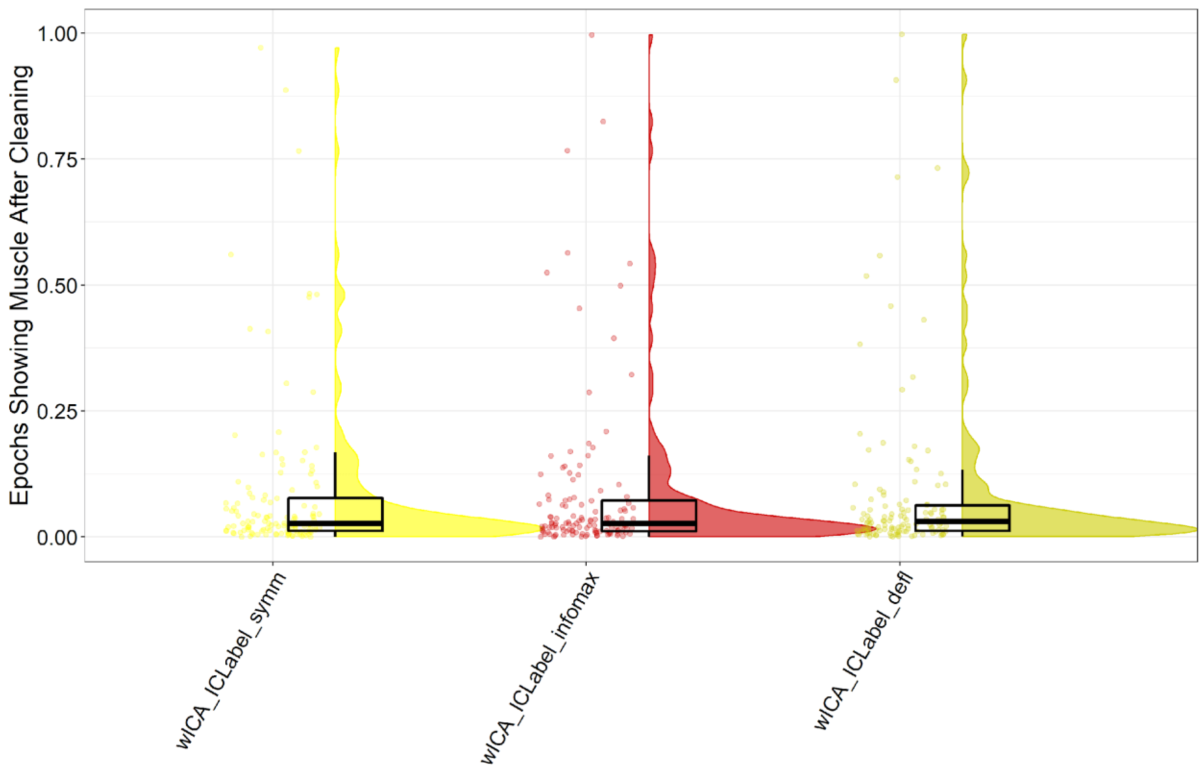

Figure S68. The proportion of epochs showing log-power log-frequency slopes indicative of muscle activity remaining after cleaning from the wICA_ICLabel cleaning approach using three different ICA methods.

| Pipeline | Mean | SD |
|---|---|---|
| wICA_ICLabel_symm | 0.087 | 0.163 |
| wICA_ICLabel_defl | 0.086 | 0.170 |
| wICA_ICLabel_infomax | 0.086 | 0.165 |

Table S11. Means and SDs for the proportion of epochs showing log-power log-frequency slopes indicative of muscle activity remaining after cleaning from the wICA_ICLabel cleaning approach using three different ICA methods.

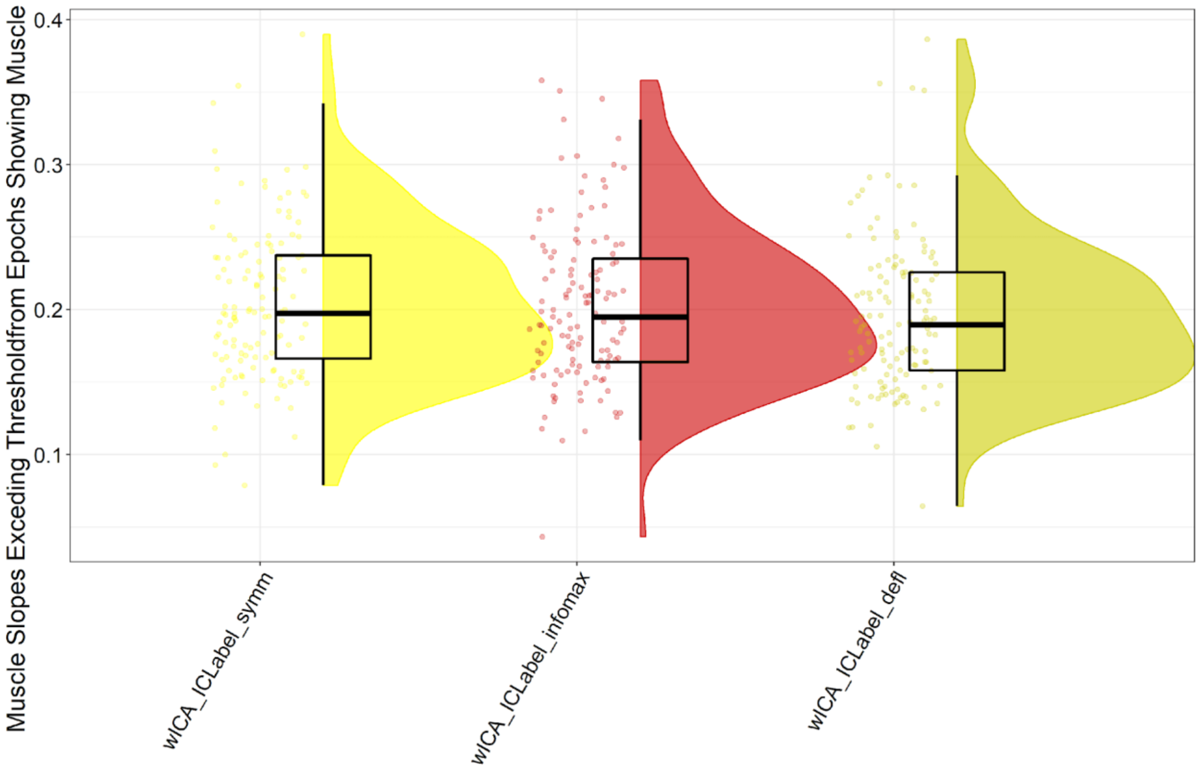

Figure S69. The amount by which log-power log-frequency slopes in epochs that showed activity indicative of muscle activity remaining after cleaning exceeded the threshold from the wICA_ICLabel cleaning approach using three different ICA methods.

| **Pipeline** | **Mean** | **SD** |
|---|---|---|
| wICA_ICLabel_infomax | 0.203 | 0.055 |
| wICA_ICLabel_defl | 0.196 | 0.053 |
| wICA_ICLabel_symm | 0.205 | 0.054 |

Table S12. Means and SDs for the amount by which log-power log-frequency slopes in epochs that showed activity indicative of muscle activity remaining after cleaning exceeded the threshold from the wICA_ICLabel cleaning approach using three different ICA methods.

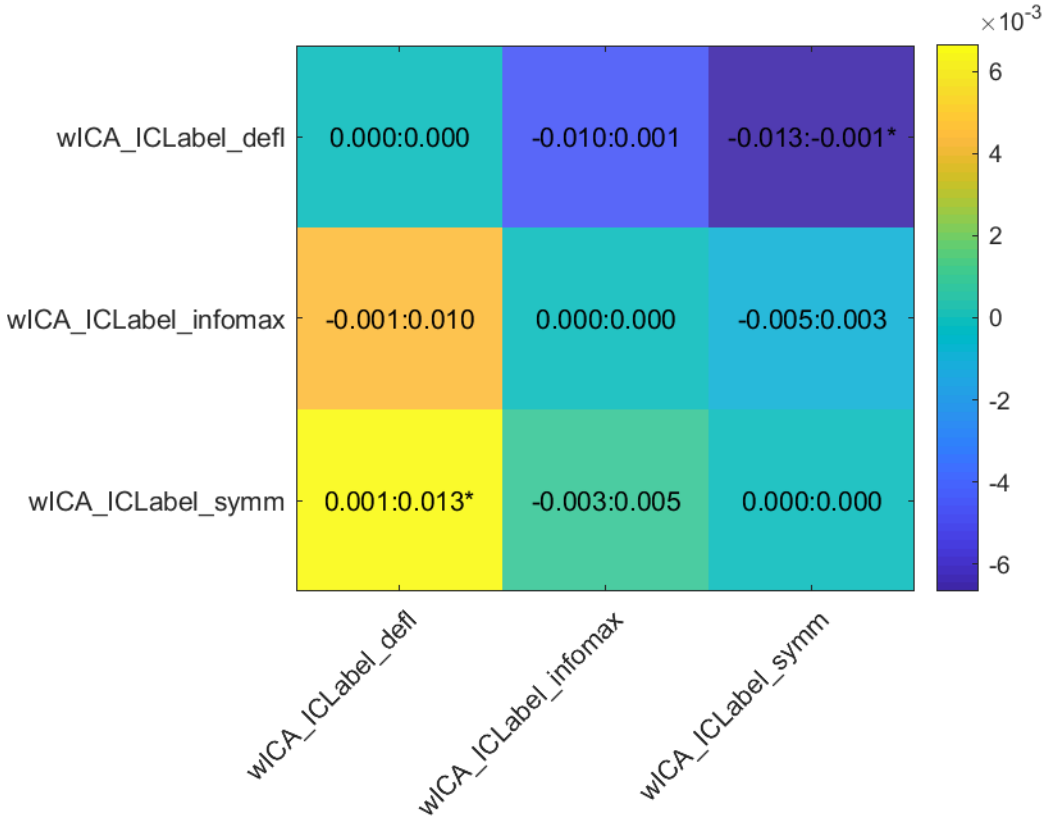

Figure S70. Post-hoc t-tests for the amount by which log-power log-frequency slopes in epochs that showed activity indicative of muscle activity remaining after cleaning exceeded the threshold from wICA_ICLabel cleaned data across the different ICA methods, with multiple comparison controls implemented using rmmcp (which implements multiple comparison controls using Hochberg's approach).

***N2 and P3 GFP***

A significant interaction was present in the between wICA_ICLabel infomax, fastica symm or fastica defl methods for the N2 GFP (p = 0.017). However, no significant interaction was present for the N2 TANOVA (p = 0.641), P3 GFP (p = 0.118), or P3 TANOVA (p = 0.967). Post-hoc tests indicated a slight benefit of infomax over defl for the N2 GFP, but no other differences (although fastica symm nearly showed better performance than fastica defl, p = 0.0505). As such, given cudaICA and fastica symm are the fastest, these methods are perhaps preferable over fastica defl, and RELAX has been set to use fastica symm by default (Figure S71-76).

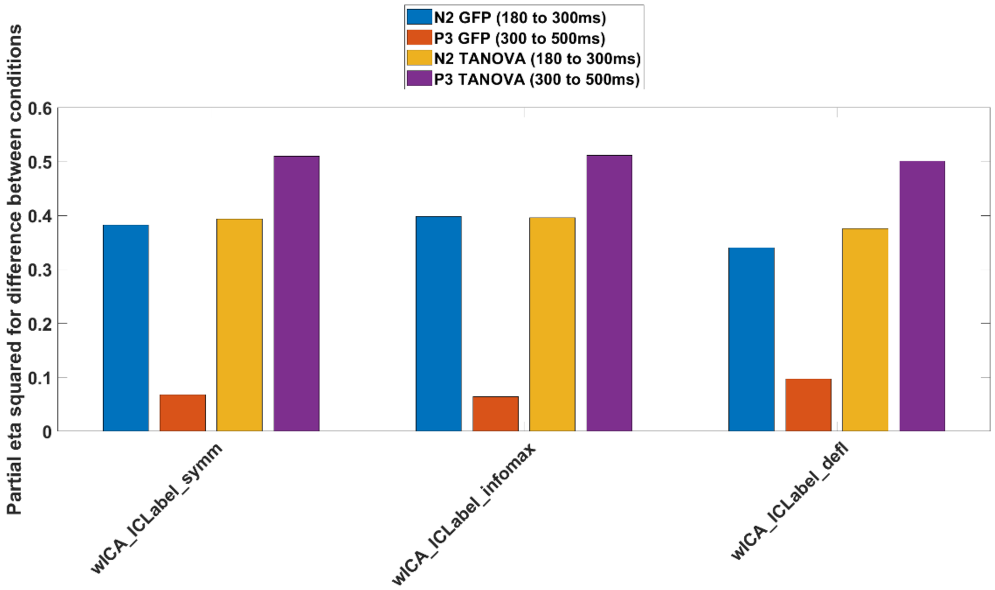

Figure S71. Variance explained by the N2 and P3 GFP and TANOVA tests between Go and Nogo conditions for each of the ICA methods used to test which ICA method was most effective for use with wICA_ICLabel.

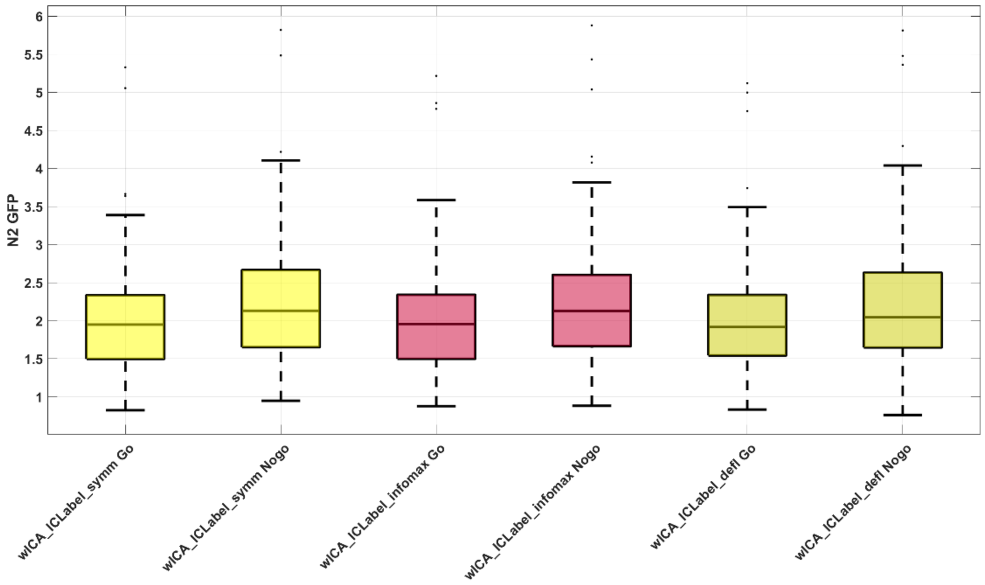

Figure S72. Box plot of N2 GFP amplitudes from the Go-Nogo dataset for a comparison between wICA_ICLabel using either the infomax, fastica symm, or fastica defl setting.

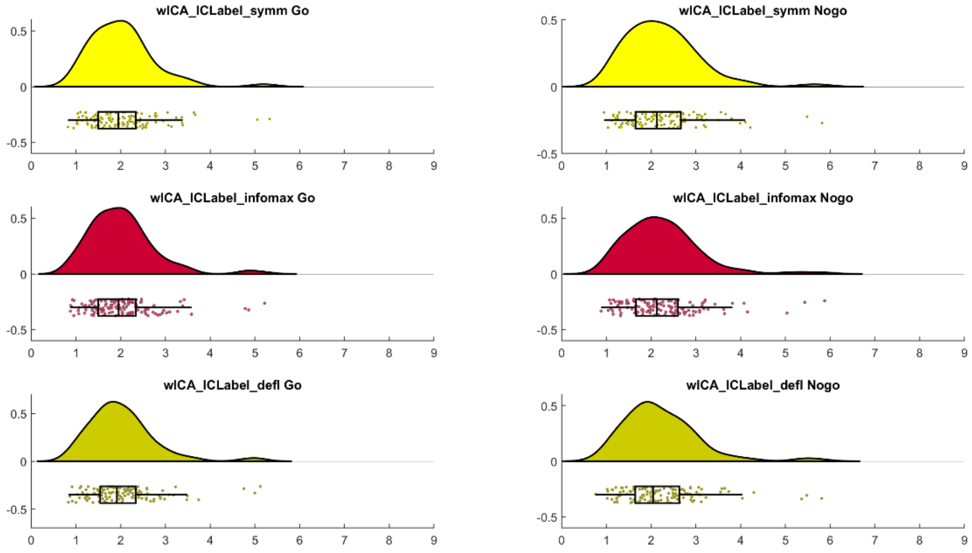

Figure S73. Raincloud plot of N2 GFP amplitudes from the Go-Nogo dataset for a comparison between wICA_ICLabel using either the infomax, fastica symm, or fastica defl setting.

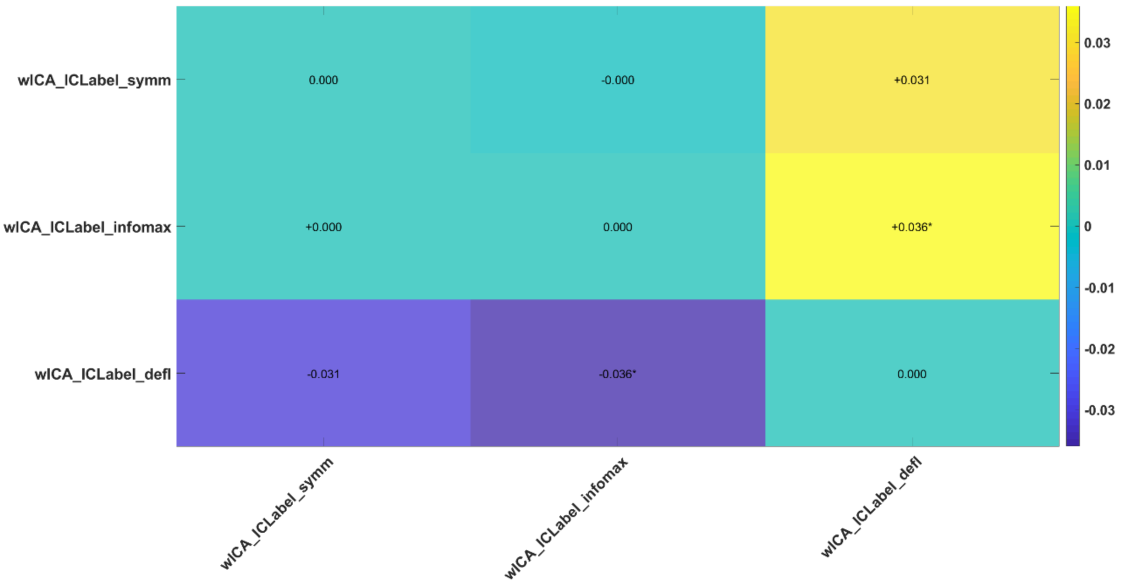

Figure S74. Post-hoc comparisons between the different ICA methods used to test wICA_ICLabel for the N2 GFP.

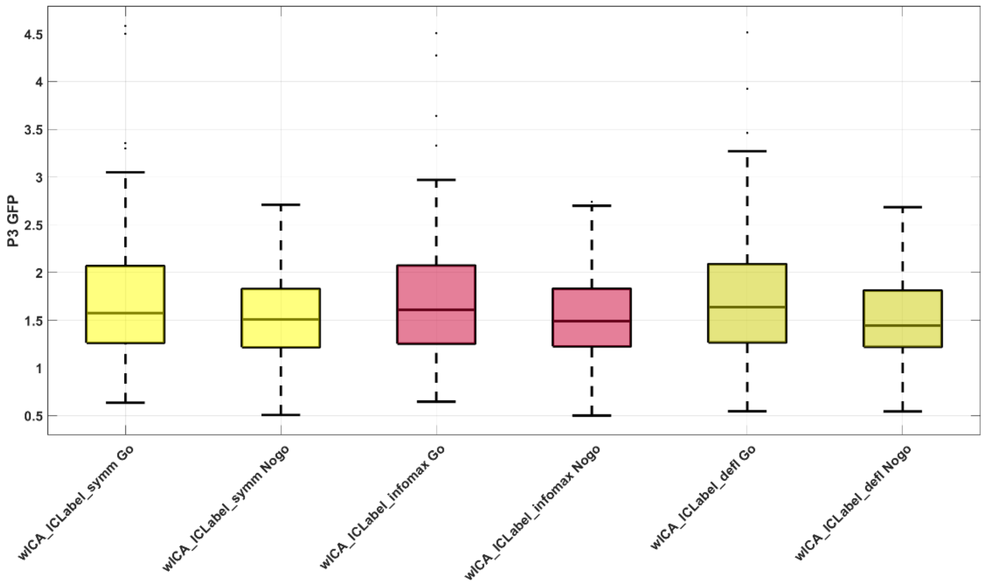

Figure S75. Box plot of P3 GFP amplitudes from the Go-Nogo dataset for a comparison between wICA_ICLabel using either the infomax, fastica symm, or fastica defl setting.

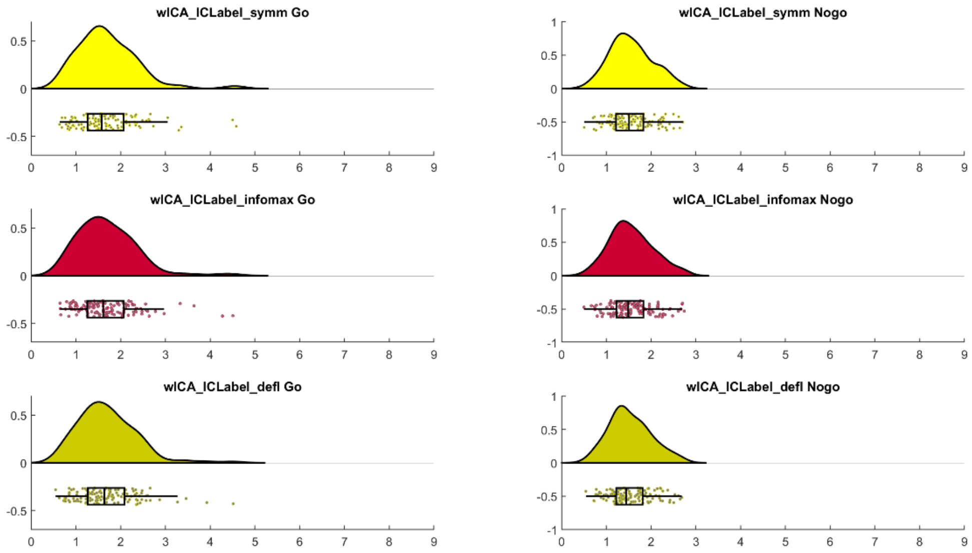

Figure S76. Raincloud plot of P3 GFP amplitudes from the Go-Nogo dataset for a comparison between wICA_ICLabel using either the infomax, fastica symm, or fastica defl setting.

Overall, it seems that infomax method performed best for blink removal (with fastica defl a close second), and the fastica defl method performed best for muscle removal. However, these results conflict with the results of Barban et al [27] who found that the fastica symm method outperformed both infomax and fastica defl for both blink and muscle removal. Additionally, the difference between the pipelines was a difference in the fBAR mean of 0.01 at largest, and 0.009 for muscle slope exceeding the threshold from muscle affected epochs after cleaning, both of which we suspect are highly unlikely to influence between condition or between group comparisons. Our “explained variance” results did not differ between the pipelines, except for in the N2, where infomax was the best approach (very closely followed by fastica symm). As such, given there was very little difference in the fBAR and muscle values across the pipelines, and cudaICA and fastica symm are the fastest, these methods are perhaps preferable over fastica defl. As such, RELAX has been set to use fastica symm by default (with instructions for implementing cudaICA if the user and users’ system can implement this approach, and fastica defl easy to implement if desired).

***Test of 1Hz Filtering Before ICA (applied to reduce artifacts in 0.25Hz filtered data)***

To test the commonly proposed solution to the problem posed by the fact that ICA performs better on 1Hz high-pass filtered data, while ERPs amplitudes are reduced by high-pass filtering above 0.3Hz, we compared data that had been 1Hz filtered prior to the ICA, and the artifact removal applied back to the 0.25Hz filtered data to our standard approach of performing the ICA decomposition on the 0.25Hz filtered data with no extra filtering. We performed this comparison for both the MWF_ICA_subtract method and the ICA_subtract method. ICLabel uses the ICA activations to determine which components are artifacts. These ICA activations can be found in the 1Hz data, or can be reconstructed in the 0.25Hz data (found in the EEG.icaact variable in EEGLAB). As such, we also tested whether it was best to use ICLabel to detect artifacts after the ICA activations had been applied back to the 0.25Hz filtered data, or whether it was best to use ICLabel to determine artifactual components on the 1Hz filtered data, then apply the ICA activations to the 0.25Hz data before removing them. In brief, our results showed that the method involving simply computing the ICA using the 0.25Hz data was best for blink removal, showed the best variance explained for the N2 GFP, and did not affect muscle activity or variance explained by the experimental manipulation for other ERP measures. It also did not make a difference if ICLabel was used to detect artifacts on ICA activations from the 1Hz data but applied back to the 0.25Hz data prior to implementing artifact detection with ICLabel (either way, computing the ICA on 1Hz filtered data was inferior to simply computing the ICA on the 0.25Hz filtered data).

***Blink Amplitude Ratio***

A significant overall difference in fBAR was present in the robust ANOVA between the different filtering approaches: $F(2.14, 160.22) = 40.01$, $p < 0.0001$. Of relevance to our question, applying 1Hz ICA activations to 0.25Hz data led to significantly larger fBAR values (worse performance) than just performing the ICA on the 0.25Hz data with no extra filtering. This was the case both for the ICA_subtract and MWF_ICA_subtract versions of the pipelines. No differences were found between the different stages at which ICLabel was used to detect artifactual components (Figure S77-S78).

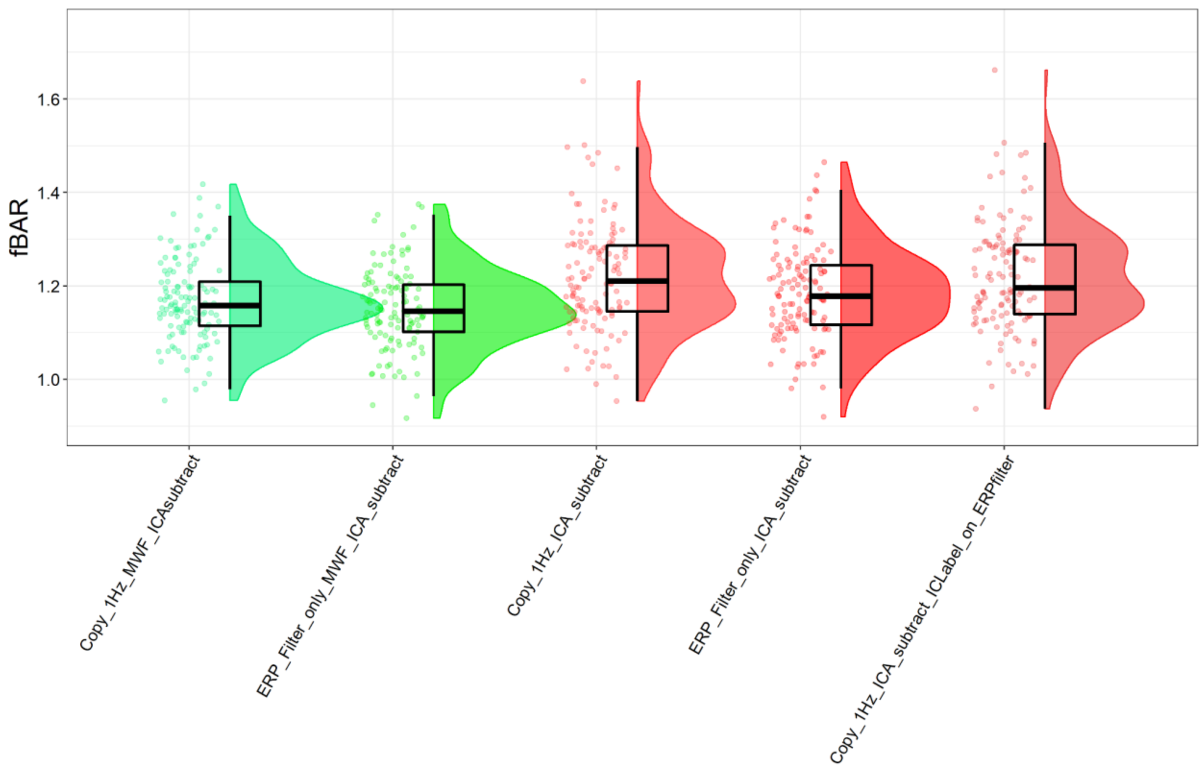

Figure S77. fBAR values for each of the different filtering approaches. The approaches labelled with Copy_1Hz first high-pass filtered at 1Hz, then computed the ICA, then subtracted the artifacts from the 0.25Hz data. Approaches labelled with ERP_Filter_only performed no additional filtering beyond the initial 0.25Hz high-pass filtering. The approach labelled with Copy_1Hz_ICA_subtract_ICLabel_on_ERPfilter applied the ICA activations back to the 0.25Hz filtered data before using ICLabel to detect artifacts (in contrast to the other 1Hz filtering approaches which used ICLabel to detect artifactual components in the 1Hz filtered data, then applied the activations to the 0.25Hz data and subtracted the artifactual components detected by ICLabel in the 1Hz data from there). Note that the scale has been reduced to allow better visualization of the data (some outliers were present from all approaches, with no observable difference in outlier frequency between the approaches).

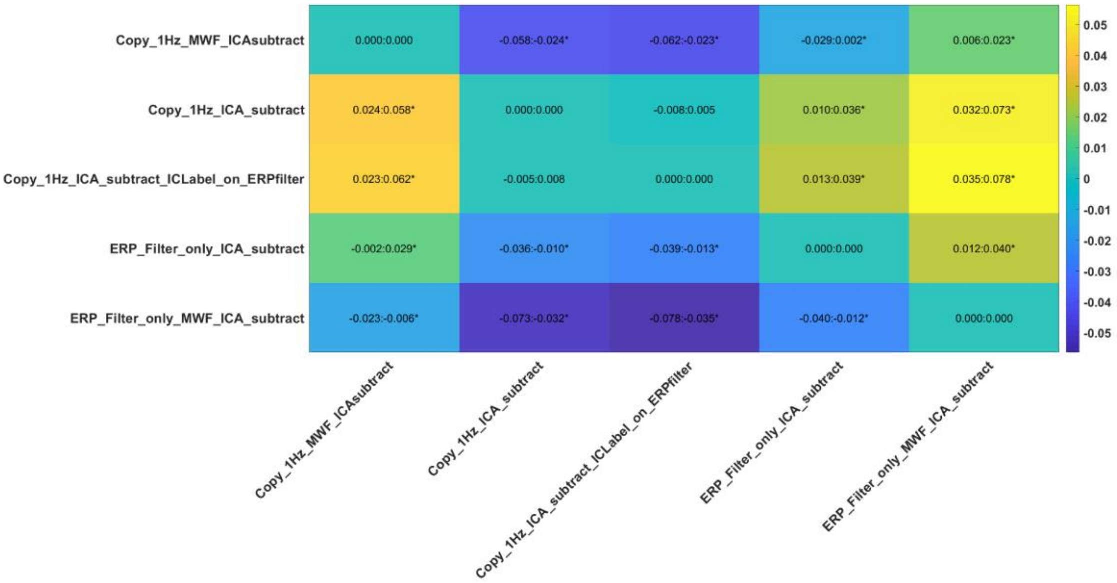

Figure S78. Post-hoc t-test comparisons of fBAR values between the different filtering approaches. Approaches labelled with Copy_1Hz first high-pass filtered at 1Hz, then computed the ICA, then subtracted the artifacts from the 0.25Hz data. Approaches labelled with ERP_Filter_only performed no additional filtering beyond the initial 0.25Hz high-pass filtering. The approach labelled with ICLabel_on_ERPfilter applied the ICA activations back to the 0.25Hz filtered data before using ICLabel to detect artifacts (in contrast to the other 1Hz filtering approaches which used ICLabel to detect artifactual components in the 1Hz filtered data, then applied the activations to the 0.25Hz data and subtracted the artifactual components detected by ICLabel in the 1Hz data from there).

#### *Proportion of Epochs Containing Muscle Activity After Cleaning*

A significant overall difference was detected in the proportion of epochs showing muscle activity after cleaning between the different filtering approaches: $F(1.87, 141.78) = 33.11$, $p < 0.0001$. Of relevance to our question, there was no difference between applying 1Hz ICA activations to 0.25Hz data compared to just performing the ICA on the 0.25Hz data with no extra filtering. This was the case both for the ICA_subtract and MWF_ICA_subtract versions of the pipelines. No differences were found between the different stages at which ICLabel was used to detect artifactual components (Figure 79-80).

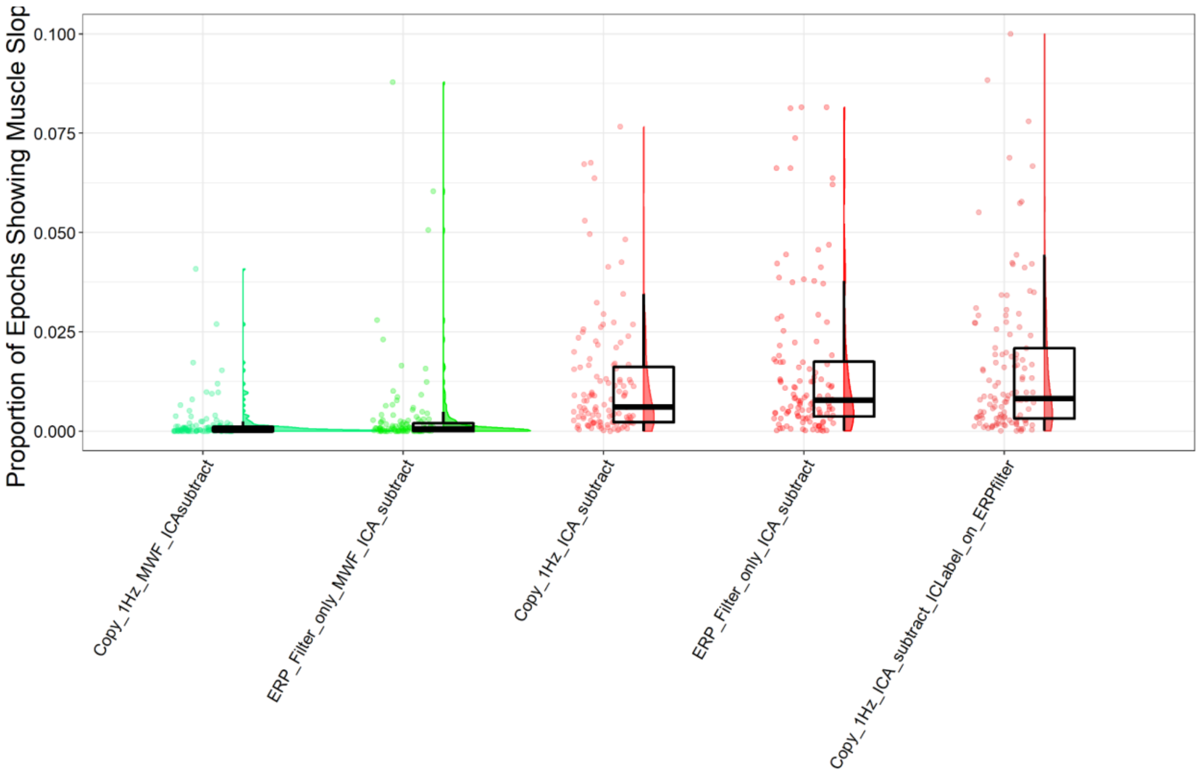

Figure S79. The proportion of epochs showing muscle activity after cleaning for each of the different filtering approaches. Approaches labelled with Copy_1Hz first high-pass filtered at 1Hz, then computed the ICA, then subtracted the artifacts from the 0.25Hz data. Approaches labelled with ERP_Filter_only performed no additional filtering beyond the initial 0.25Hz high-pass filtering. The approach labelled with Copy_1Hz_ICA_subtract_ICLabel_on_ERPfilter applied the ICA activations back to the 0.25Hz filtered data before using ICLabel to detect artifacts (in contrast to the other 1Hz filtering approaches which used ICLabel to detect artifactual components in the 1Hz filtered data, then applied the activations to the 0.25Hz data and subtracted the artifactual components detected by ICLabel in the 1Hz data from there). Note that the scale has been reduced to allow better visualization of the data (some outliers were present from all approaches, with no observable difference in outlier frequency between the approaches).

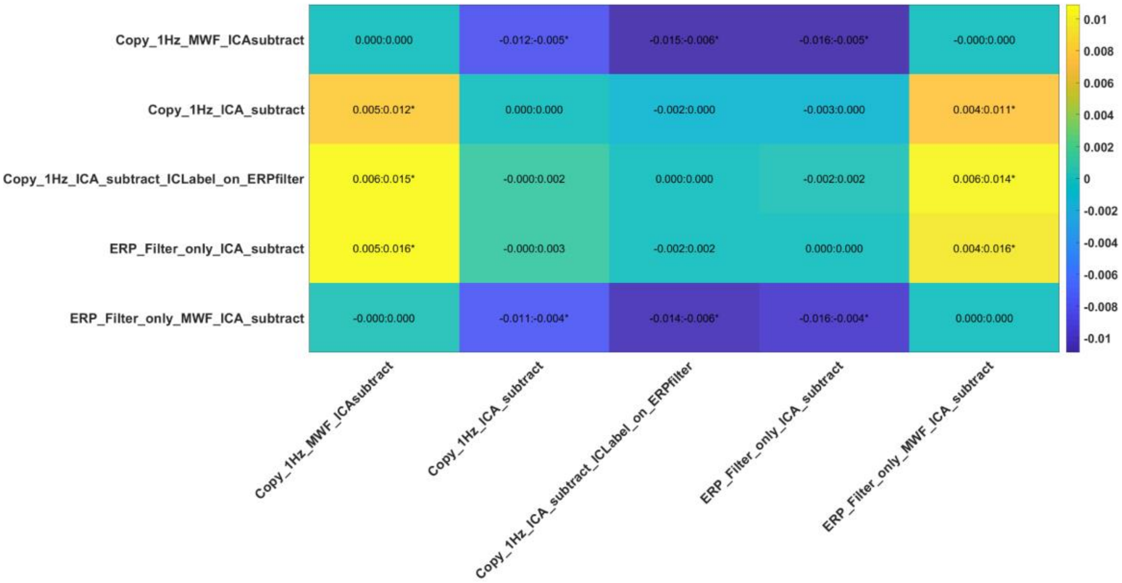

Figure S80. Post-hoc t-test comparisons of the proportion of epochs showing muscle activity after cleaning between the different filtering approaches. Approaches labelled with Copy_1Hz first high-pass filtered at 1Hz, then computed the ICA, then subtracted the artifacts from the 0.25Hz data. Approaches labelled with ERP_Filter_only performed no additional filtering beyond the initial 0.25Hz high-pass filtering. The approach labelled with Copy_1Hz_ICA_subtract_ICLabel_on_ERPfilter applied the ICA activations back to the 0.25Hz filtered data before using ICLabel to detect artifacts (in contrast to the other 1Hz filtering approaches which used ICLabel to detect artifactual components in the 1Hz filtered data, then applied the activations to the 0.25Hz data and subtracted the artifactual components detected by ICLabel in the 1Hz data from there).

#### *Variance Explained by the Experimental Manipulation*

There was a trend towards a significant interaction between the different filtering/ICA methods and the Go/Nogo trials for the N2 GFP ($p = 0.0521$). Since the effect has not been tested before and sensitivity to the detection of experimental effects with an optimal EEG pre-processing pipeline is of interest, this was explored further in a poct-hoc test (Figure S81-82). Of relevance to our question, the post-hoc test indicated that for the ICA_subtract method, simply performing the ICA on the 0.25Hz data with no extra filtering led to an improved ability to differentiate the Go and Nogo trials from the N2 GFP compared to applying 1Hz ICA activations to 0.25Hz filtered data. This was the case regardless of whether ICLabel was used to detect artifacts within the 1Hz filtered data, or whether the ICA decomposition from the 1Hz filtered data were applied to the 0.25Hz filtered data prior to ICLabel being applied to detect artifacts. No differences were found between the different stages at which ICLabel was used to detect artifactual components. There was no significant interaction between the different filtering/ICA methods and the Go/Nogo trials for the N2 TANOVA ($p = 0.8030$), nor P3 GFP ($p = 0.9622$), nor P3 TANOVA ($p = 0.6226$). When the N2 GFP result is combined with the superior blink correction from the simple 0.25Hz filtering method, our results suggest that applying ICA activations from 1Hz filtered data back to data filtered for ERP analysis inadvertently reduced data cleaning and leads to worse experimental outcomes, despite the fact that previous research has indicated that ICA performs better on data that is high-pass filtered at 1Hz [28].

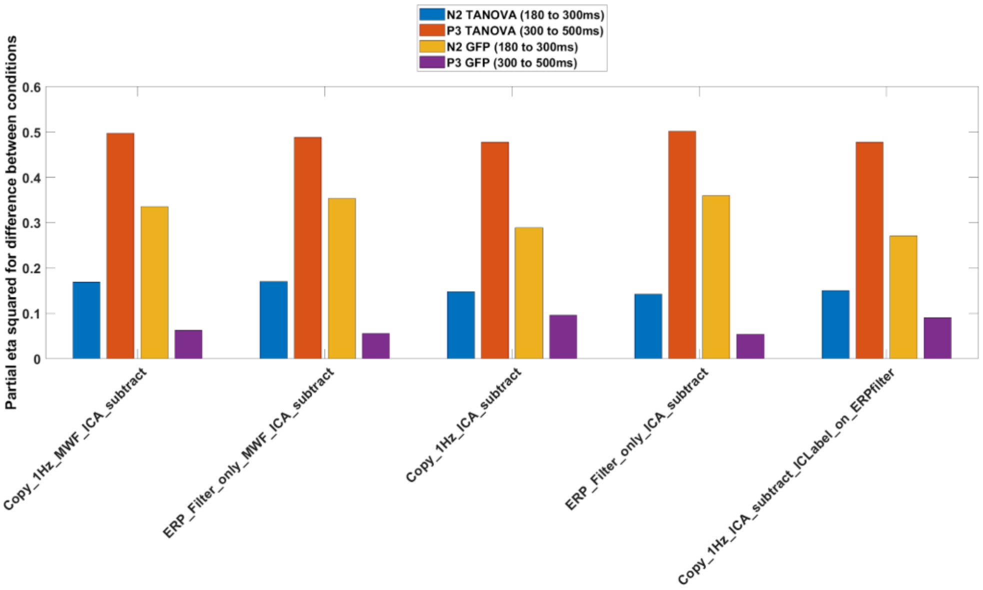

Figure S81. The variance explained by the difference between Go and Nogo trials for different filtering approaches prior to ICA. Note that applying 1Hz filtering prior to ICA, then applying the ICA subtraction to the 0.25Hz filtered data significantly reduced the variance explained for N2 GFP compared to only filtering at 0.25Hz, but no other significant differences were present between the 1Hz ICA copied to 0.25Hz approaches and 0.25Hz filtering only approach.

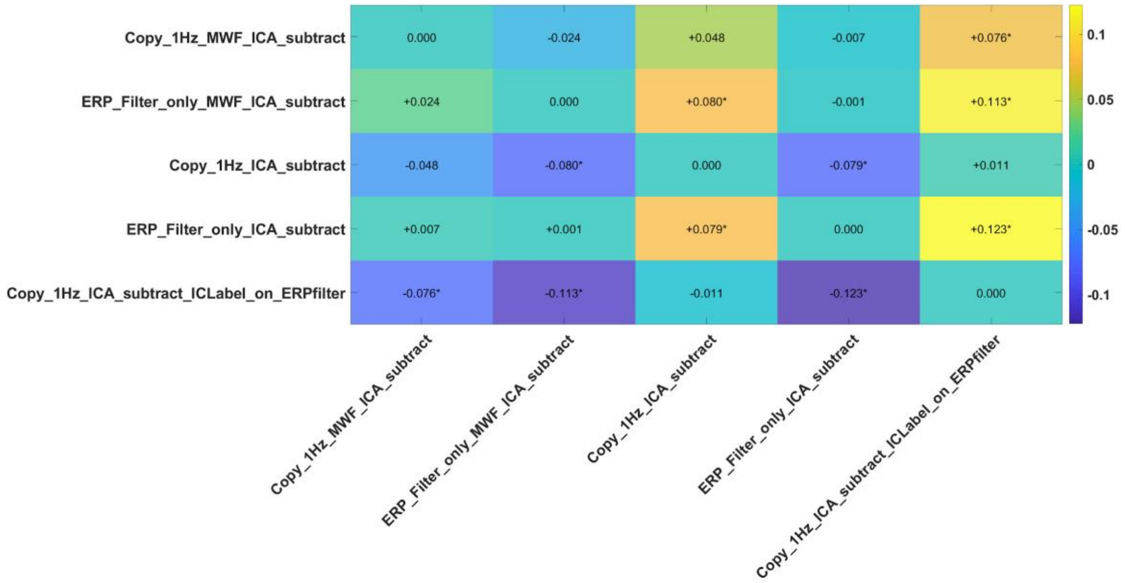

Figure S82. Post-hoc t-test comparisons for the variance explained by the interaction between Go and Nogo trials and the different filtering ICA combinations tested for the N2 GFP measure. Note the copying of 1Hz ICA to 0.25Hz data significantly reduced the variance explained for the ICA_subtract pipeline.

### SECTION SIX

#### Cleaned Dataset Examples

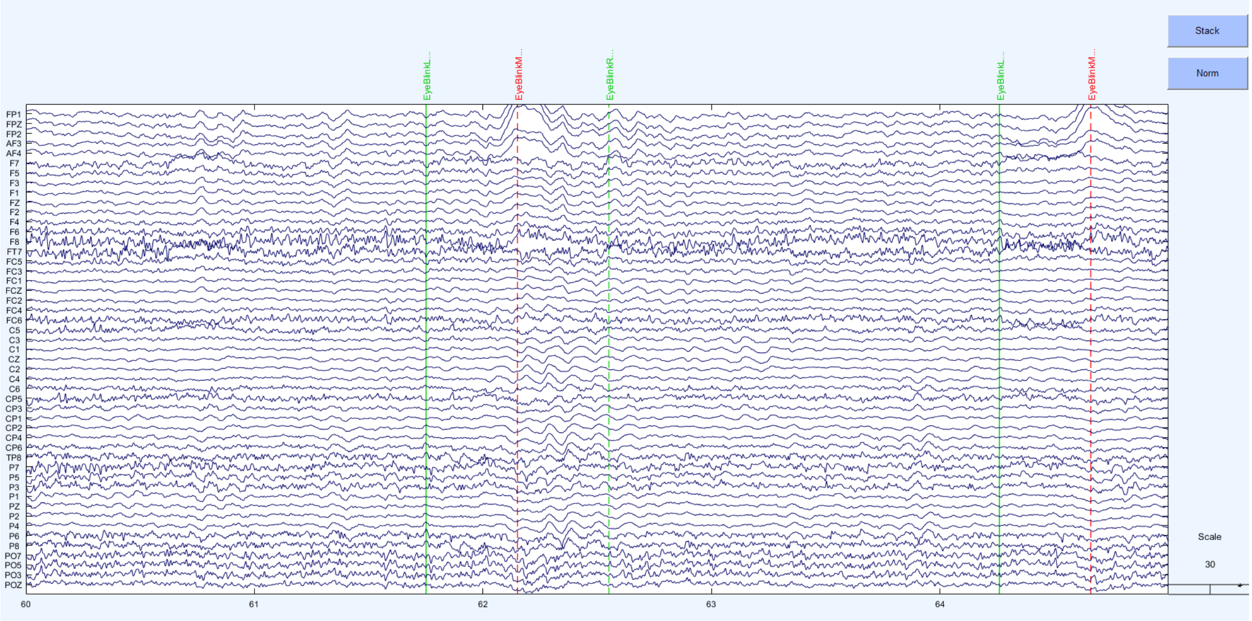

Figure S83. Raw data example from the Hard Go-Nogo dataset

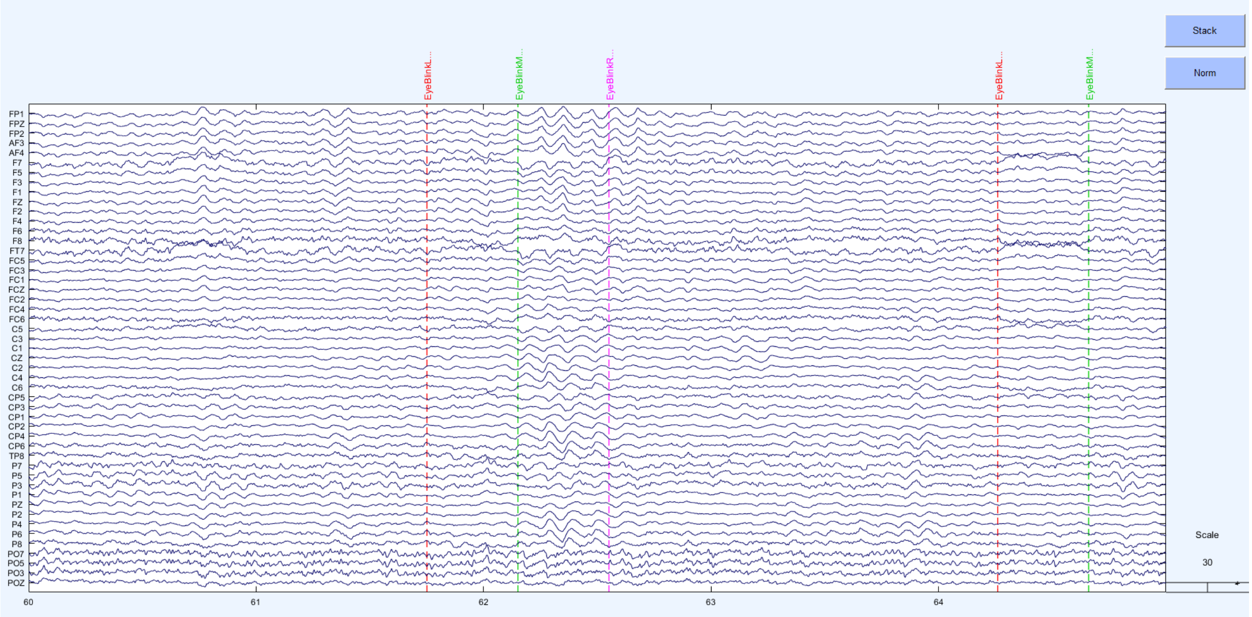

Figure S84. MWF_wICA_infomax cleaned example from the Hard Go-Nogo dataset

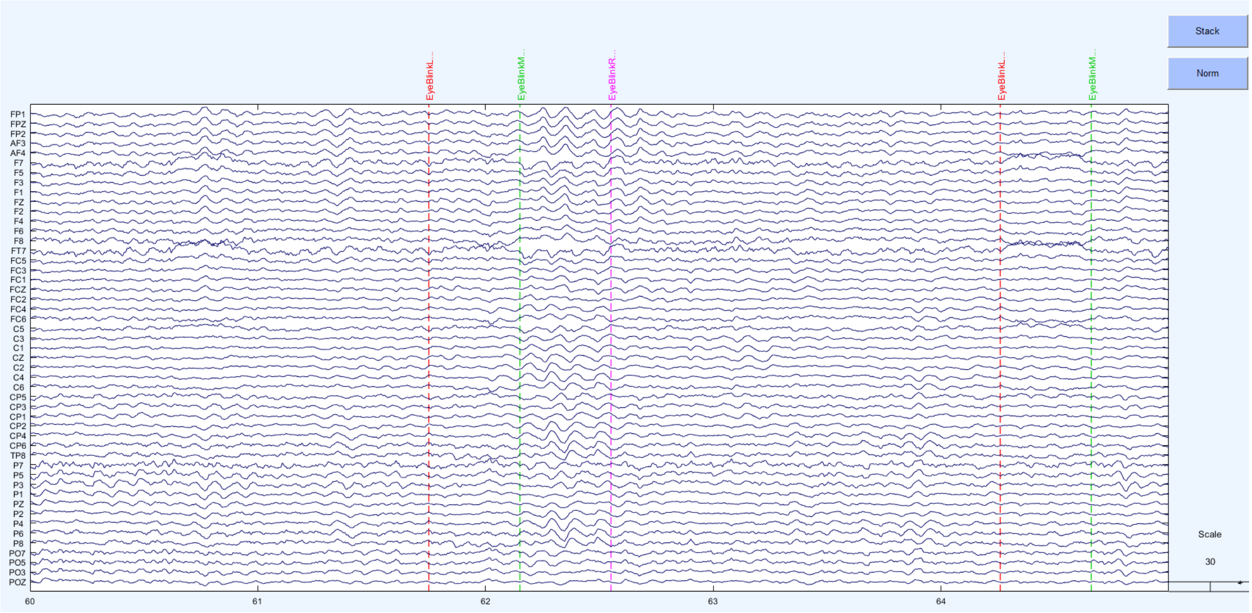

Figure S85. MWF_ICA_subtract cleaned example

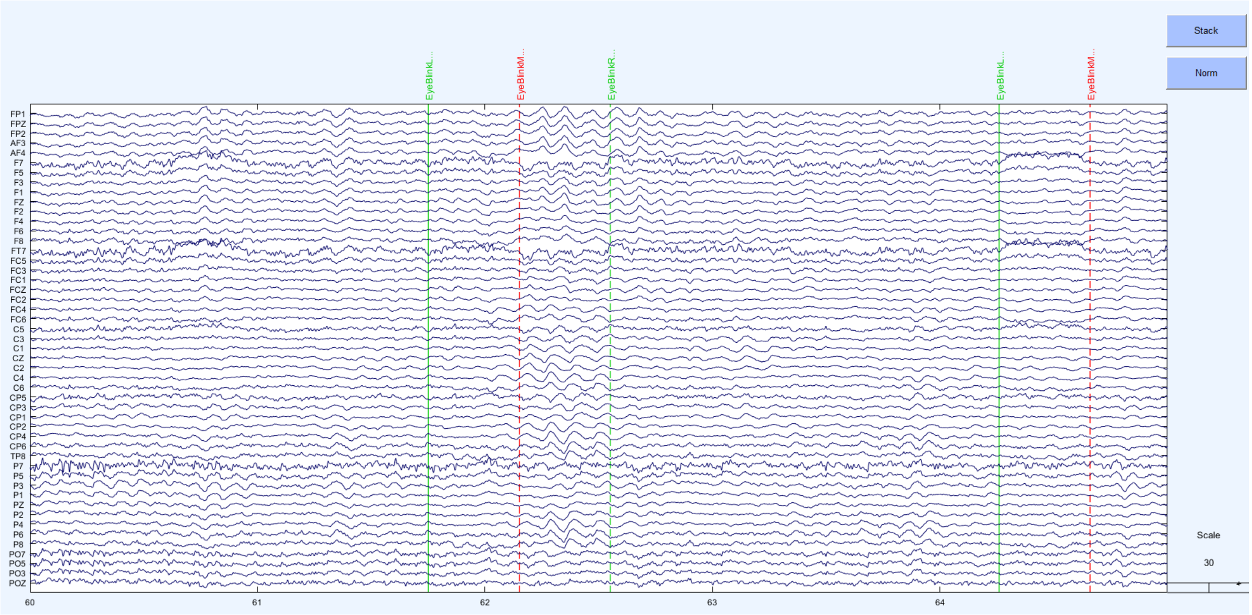

Figure S86. ICA_subtract cleaned example

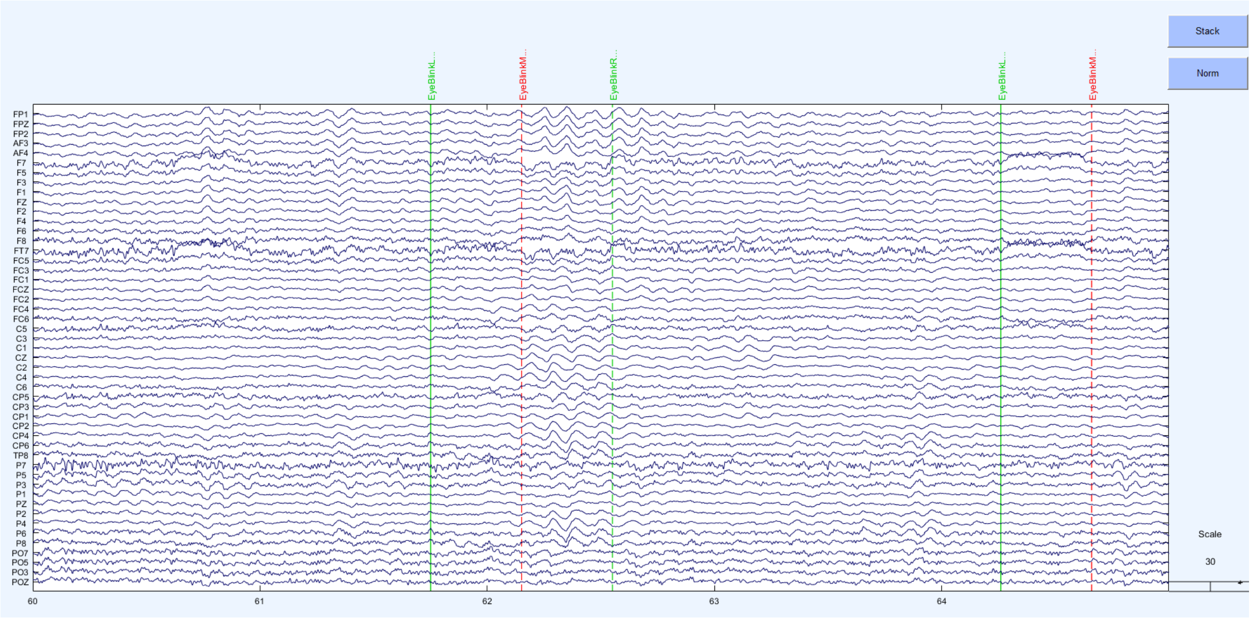

Figure S87. wICA_ICLabel cleaned example

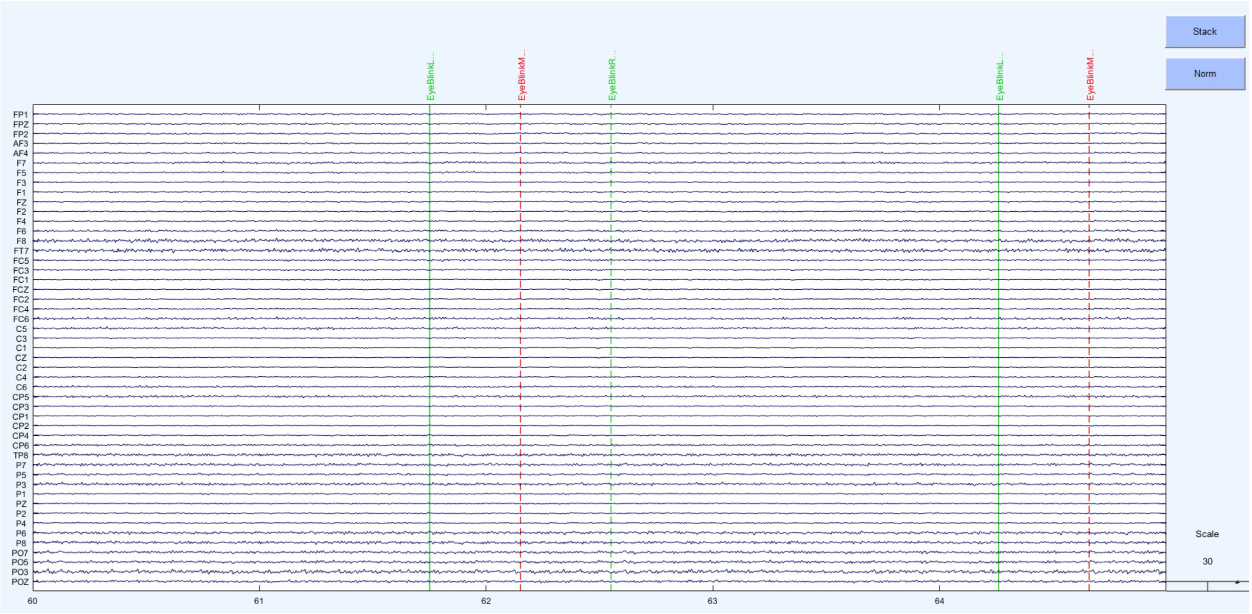

Figure S88. wICA_all cleaned dataset

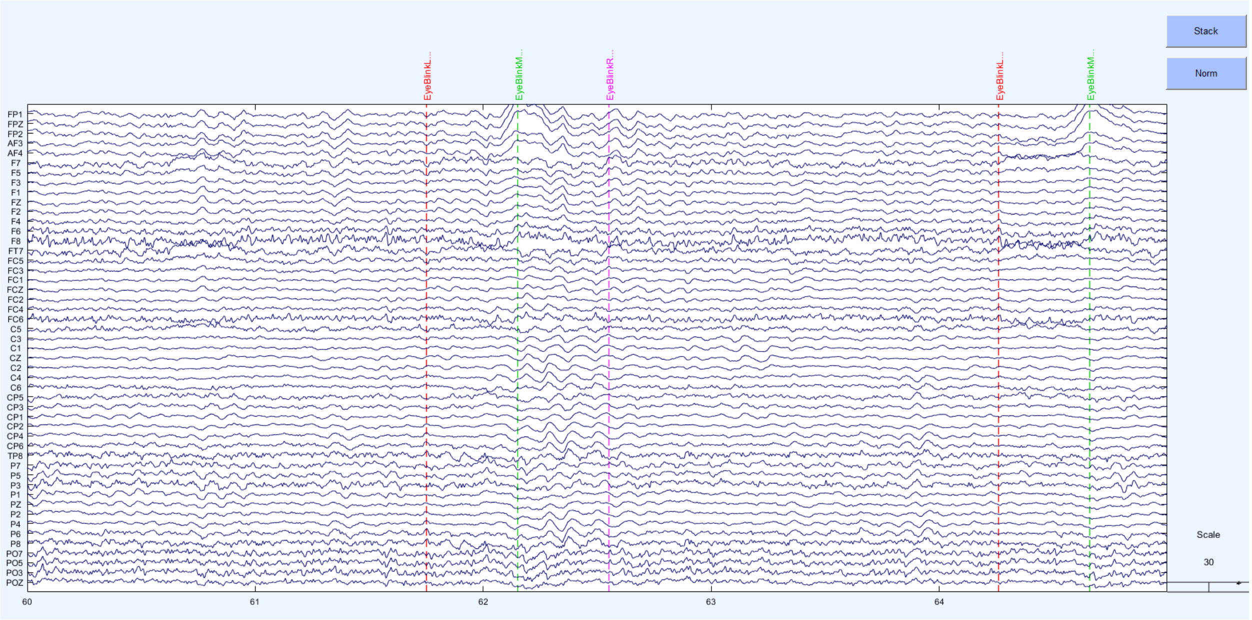

Figure S89. MWF_only cleaned example from the Hard Go-Nogo dataset

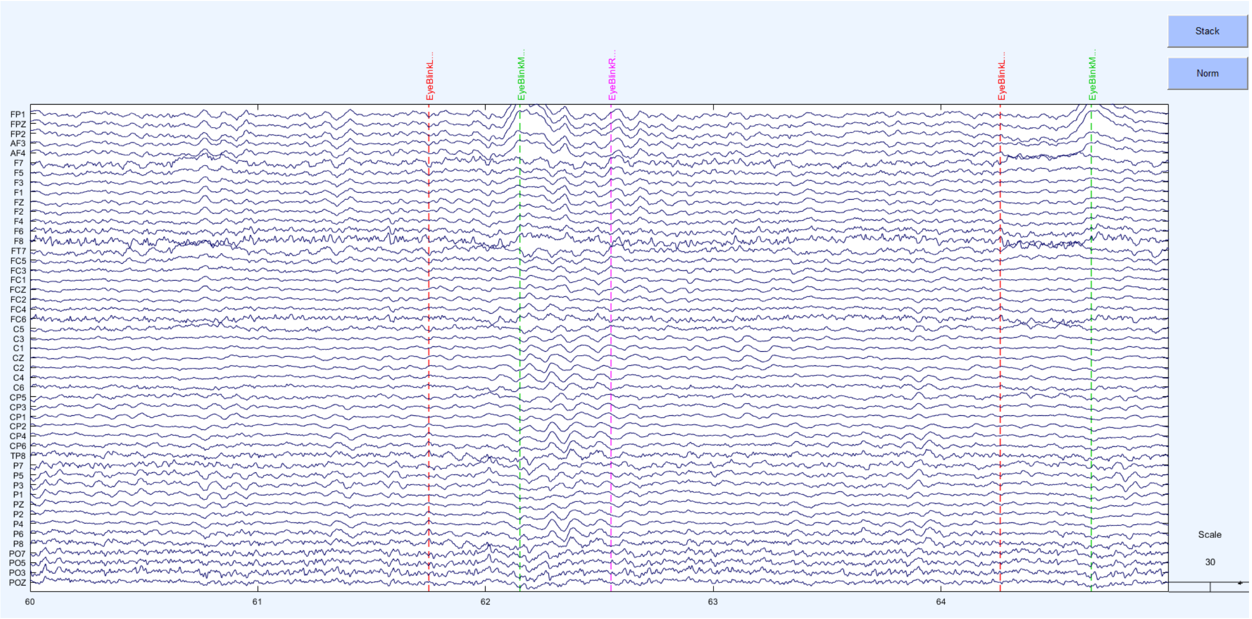

Figure S90. MWF_CCA cleaned example dataset

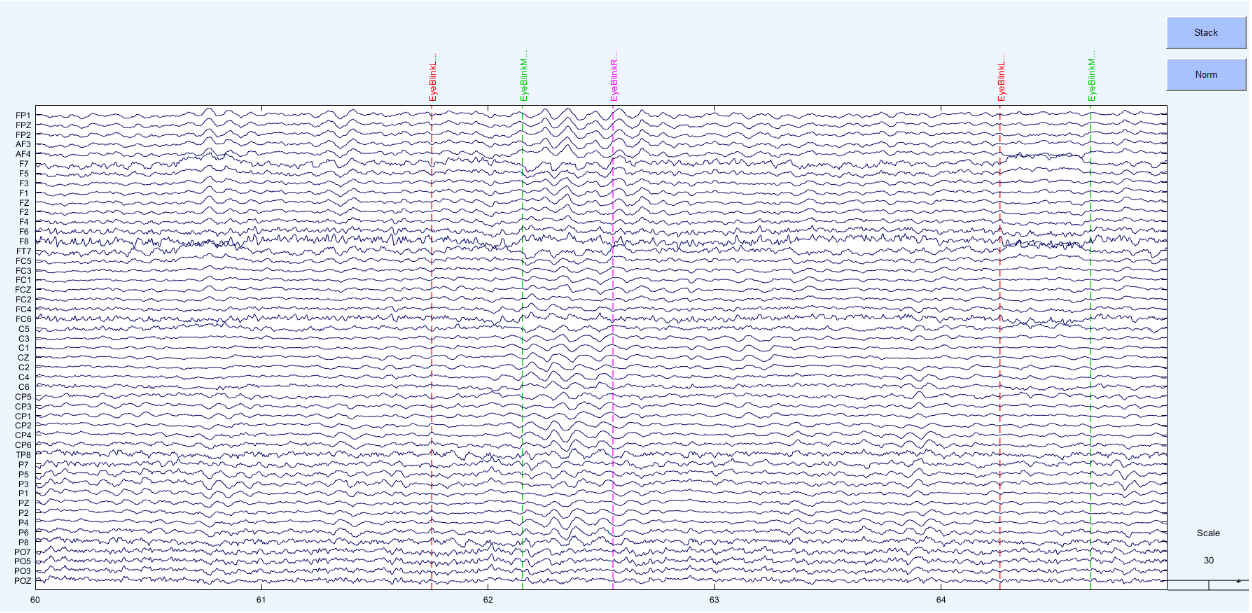

Figure S91. MWF_wICA_CCA cleaned dataset

### SECTION SEVEN

#### Supplementary Discussion Points

1) Some previous research has suggested that filtering out data above 45Hz improves ICA decomposition [13]. Filtering out data above 45Hz prior to the wICA step of RELAX seemed to lead to better performance in the amount of variance explained by brain activity from components identified by ICLabel. However, ICLabel uses a frequency metric to determine muscle artifact, so filtering out data above 45Hz may just prevent muscle being detected by ICLabel, while not necessarily eliminating the effects of artifacts on the data. Additionally, filtering out >45Hz activity also decreased the variance explained by the experimental manipulation for the P3 GFP, and decreased the SER. As such, we do not recommend filtering out data above 45Hz prior to the wICA step.

2) While our data also indicated that MWF_wICA_infomax may perform better than MWF_wICA_fastICA and MWF_wICA_AMICA, this may be a product of the infomax algorithm being used both for cleaning and for detection of brain variance in the measure, biasing results towards that cleaning pipeline.

3) Our results confirmed that RELAX could be applied to data high-pass filtered at 0.25Hz so the pipeline can be applied to ERP studies. Interestingly, SER values were lower and ARR values were higher when data were high-pass filtered at 0.25Hz compared to when data was high-pass filtered at 1Hz. We suspect this is because the “uncleaned” data for calculation of SER and ARR was taken after filtering was applied. As such, the 0.25Hz filtered data contained artifacts between 0.25 and 1Hz that were reduced (reducing the SER and increasing the ARR values), whereas data that were high-pass filtered at 1Hz did not contain artifacts in that 0.25 to 1Hz range, so the SER and ARR values were not affected by cleaning of data in those frequencies (leading to less cleaning being performed in this 1Hz high-pass filtered data, and higher SER / lower ARR values). We informally tested a potential reason for this by low-pass filtering the raw data at 1Hz. When we inspected the data after this filtering, we observed that the periods we had marked as containing a blink still showed large voltage deviations. This indicated that blinks contain influence from data between 0.25 and 1Hz, so high-pass filtering at 1Hz removes an aspect of the blink data, which is not removed when data is high-pass filtered at 0.25Hz. It also seems that both MWF and wICA were less effective at cleaning blink artifact contributions from the frequencies between 0.25 to 1Hz, as BAR values were higher for the ERP filtered dataset in all pipelines than the BAR values reported in our companion paper (where the data was filtered at 1Hz). This is an issue that we think the field still needs to address – while strong recommendations have been made not to high-pass filter data with a setting above 0.3Hz for ERP analyses (or perhaps even 0.1Hz), it seems that no cleaning method is available that can address artifacts when data is high-pass filtered at 0.25Hz as effectively as when data is high-pass filtered at 1Hz. While a common solution is to high-pass filter at 1Hz before performing ICA decomposition, then apply the 1Hz ICA decomposition to the 0.25Hz filtered data, our results indicated this resulted in worse performance than simply performing the ICA decomposition on the 0.25Hz filtered data. As such, filter settings and adequate cleaning of ERP data is still an issue that needs to be resolved by further research.

### Supplementary Materials References

1. Zeng H, Song A. Removal of EOG artifacts from EEG recordings using stationary subspace analysis. The Scientific World Journal. 2014;2014.
2. Castellanos NP, Makarov VA. Recovering EEG brain signals: Artifact suppression with wavelet enhanced independent component analysis. Journal of neuroscience methods. 2006;158(2):300-12.
3. Somers B, Francart T, Bertrand A. A generic EEG artifact removal algorithm based on the multi-channel Wiener filter. Journal of neural engineering. 2018;15(3):036007.
4. Janani AS, Grummett TS, Lewis TW, Fitzgibbon SP, Whitham EM, DelosAngeles D, et al. Improved artefact removal from EEG using Canonical Correlation Analysis and spectral slope. Journal of neuroscience methods. 2018;298:1-15.
5. De Clercq W, Vergult A, Vanrumste B, Van Paesschen W, Van Huffel S. Canonical correlation analysis applied to remove muscle artifacts from the electroencephalogram. IEEE transactions on Biomedical Engineering. 2006;53(12):2583-7.
6. Gao J, Zheng C, Wang P. Online removal of muscle artifact from electroencephalogram signals based on canonical correlation analysis. Clinical EEG and neuroscience. 2010;41(1):53-9.
7. Pion-Tonachini L, Kreutz-Delgado K, Makeig S. ICLabel: An automated electroencephalographic independent component classifier, dataset, and website. NeuroImage. 2019;198:181-97.
8. Issa MF, Juhasz Z. Improved EOG artifact removal using wavelet enhanced independent component analysis. Brain sciences. 2019;9(12):355.
9. Mammone N, La Foresta F, Morabito FC. Automatic artifact rejection from multichannel scalp EEG by wavelet ICA. IEEE Sensors Journal. 2011;12(3):533-42.
10. Raimondo F, Kamienkowski JE, Sigman M, Fernandez Slezak D. CUDAICA: GPU optimization of infomax-ICA EEG analysis. Computational intelligence and neuroscience. 2012;2012.
11. Hyvarinen A, editor Fast ICA for noisy data using Gaussian moments. 1999 IEEE international symposium on circuits and systems (ISCAS); 1999: IEEE.
12. Palmer JA, Kreutz-Delgado K, Makeig S. AMICA: An adaptive mixture of independent component analyzers with shared components. Swartz Center for Computatonal Neursoscience, University of California San Diego, Tech Rep. 2012.
13. Zakeri Z. Optimised use of independent component analysis for EEG signal processing: University of Birmingham; 2017.
14. Bertrand A. Distributed signal processing for wireless EEG sensor networks. IEEE Transactions on Neural Systems and Rehabilitation Engineering. 2015;23(6):923-35.
15. Somers B, Bertrand A. Removal of eye blink artifacts in wireless EEG sensor networks using reduced-bandwidth canonical correlation analysis. Journal of neural engineering. 2016;13(6):066008.
16. Robbins KA, Touryan J, Mullen T, Kothe C, Bigdely-Shamlo N. How sensitive are EEG results to preprocessing methods: a benchmarking study. IEEE transactions on neural systems and rehabilitation engineering. 2020;28(5):1081-90.
17. Fitzgibbon S, DeLosAngeles D, Lewis T, Powers D, Grummett T, Whitham E, et al. Automatic determination of EMG-contaminated components and validation of independent component analysis using EEG during pharmacologic paralysis. Clinical Neurophysiology. 2016;127(3):1781-93.
18. Clayson PE, Baldwin S, Rocha H, Larson MJ. The Data-Processing Multiverse of Event-Related Potentials (ERPs): A Roadmap for the Optimization and Standardization of ERP Processing and Reduction Pipelines. 2021.
19. Habermann M, Weusmann D, Stein M, Koenig T. A student's guide to randomization statistics for multichannel event-related potentials using ragu. Frontiers in neuroscience. 2018;12:355.

20. Koenig T, Kottlow M, Stein M, Melie-García L. Ragu: a free tool for the analysis of EEG and MEG event-related scalp field data using global randomization statistics. Computational intelligence and neuroscience. 2011;2011.
21. Benjamini Y, Hochberg Y. Controlling the false discovery rate: a practical and powerful approach to multiple testing. Journal of the Royal statistical society: series B (Methodological). 1995;57(1):289-300.
22. Clayson PE, Carbine KA, Baldwin SA, Olsen JA, Larson MJ. Using generalizability theory and the ERP Reliability Analysis (ERA) Toolbox for assessing test-retest reliability of ERP scores Part 1: Algorithms, framework, and implementation. International Journal of Psychophysiology. 2021.
23. Clayson PE, Miller GA. ERP Reliability Analysis (ERA) Toolbox: An open-source toolbox for analyzing the reliability of event-related brain potentials. International Journal of Psychophysiology. 2017;111:68-79.
24. Alday PM. How much baseline correction do we need in ERP research? Extended GLM model can replace baseline correction while lifting its limits. Psychophysiology. 2019;56(12):e13451.
25. Luck SJ, Stewart AX, Simmons AM, Rhemtulla M. Standardized measurement error: A universal metric of data quality for averaged event-related potentials. Psychophysiology. 2021:e13793.
26. Lopez-Calderon J, Luck SJ. ERPLAB: an open-source toolbox for the analysis of event-related potentials. Frontiers in human neuroscience. 2014;8:213.
27. Barban F, Chiappalone M, Bonassi G, Mantini D, Semprini M. Yet another artefact rejection study: an exploration of cleaning methods for biological and neuromodulatory noise. Journal of Neural Engineering. 2021.
28. Winkler I, Debener S, Müller K-R, Tangermann M, editors. On the influence of high-pass filtering on ICA-based artifact reduction in EEG-ERP. 2015 37th Annual International Conference of the IEEE Engineering in Medicine and Biology Society (EMBC); 2015: IEEE.
